# Material context and TGF-β blockade timing determine matrix retained after withdrawal in patient-derived GelMA-bioprinted keratoconus stroma

**DOI:** 10.64898/2026.11.04.764220

**Authors:** Hanyu Jiang, Chang Zhou, Fei Gao, Yuan Pang, Ying Li, Ying Jie, Huayu Yang, Yang Jiang

## Abstract

Keratoconus (KC) repair depends on host matrix, yet cultured KC cells produce less matrix. This has been interpreted as impaired synthesis, whereas tissue analyses point to the state of a thinned stroma. TGF-β induces matrix alongside myofibroblast transformation (scarring). Scarless repair separates the two in vivo, but two-dimensional (2D) culture does not. Donor-matched, patient-derived three-dimensional (3D) gelatin methacryloyl (GelMA) bioprints from six KC and six normal donors preserved the KC tissue expression state and restored this separation. KC matrix secretion per unit DNA matched that of normal cells, and lower total output arose only from fewer cells. A single TGF-β pulse induced matrix synthesis with less scarring, increasing programs low in KC tissue and the collagen genes COL5A1 and COL1A1, whose risk alleles thin and soften the cornea. Delayed receptor blockade for a finite course left more native collagen and less scarring than vehicle in 9 of 12 donors after withdrawal. Surrogate readouts used to select courses disagreed with retained host matrix. Continuous blockade suppressed scarring most deeply during treatment and produced the lowest matrix-gene expression. The course preferred in 2D produced less matrix and more scarring in 3D. Selection by retained matrix instead of readouts during treatment left 22.2% more native collagen. TGF-β-pathway courses should therefore be evaluated by the host matrix and scarring remaining after withdrawal.

## Introduction

Keratoconus (KC) progressively thins the corneal stroma and reduces its resistance to deformation [1]. Genetically predicted lower corneal resistance increases disease risk [2]. Corneal cross-linking, the most widely used treatment, forms covalent bonds within stromal collagen fibrils to stiffen the stroma and halt progression [3]. Cross-linking strengthens the remaining stroma. Restoring lost stromal tissue requires a different class of treatment. One route adds tissue, using adipose-derived stem cells and decellularized donor laminas [4,5] or a cell-free bioengineered porcine construct [6] to thicken the stroma. The other induces host cells to synthesize matrix. Eyedrops containing TGF-β3 and dexamethasone are registered for progressive KC to promote collagen formation and stiffen the cornea [7]. Both routes preserve the host cornea, whose remaining stromal cells contribute to the restored matrix. Repair depends on whether these cells can be induced to synthesize matrix and whether their induced output reaches that of normal cells. Matrix induction must also be separable from myofibroblast transformation and the resulting scarring. The course of induction then needs defined start and stop times and an endpoint by which to evaluate it.

In conventional culture, KC cells contain less fibronectin, collagen and collagen-processing protein than normal cells, even at matched cell numbers. This difference has been attributed to an intrinsic synthetic defect caused by the integrated stress response, which suppresses protein synthesis under stress [8]. Matrix content in tissue or an engineered construct depends on both cell number and synthesis per cell. KC corneas contain fewer keratocytes [9,10], and KC cells form thinner matrices in self-assembled three-dimensional constructs [11]. The contributions of cell number and synthesis per cell have not been separated. KC matrix synthesis remains inducible. Relieving the integrated stress response increased hydroxyproline, a component of collagen, by 35% after normalization to cell number [12], and TGF-β1 or β3 thickened the matrix in constructs [11]. Low inflammatory, immediate-early and TGF-β programs in KC tissue have likewise been interpreted as a matrix state with preserved repair capacity [2]. Inducibility alone does not establish normal output, because the same treatment increased hydroxyproline in normal cells by 32% [12]. Resolving this distinction requires simultaneous measurement of cell content and matrix output in the same three-dimensional material, with KC and normal cells compared before and after stimulation.

TGF-β stimulates matrix synthesis and induces fibroblasts to undergo myofibroblast transformation [13]. Because myofibroblasts contract and deposit scar-like matrix, matrix induction can lead to scarring. Repair in vivo shows that these processes can be separated. As matrix refills freeze-injured rabbit cornea, collagen and proteoglycan genes rise without increases in ACTA2 (α-SMA), a myofibroblast marker, or COL3A1 (collagen III), a scar-associated gene [14]. Whether cultured cells achieve this separation depends on their surrounding material. Substrate stiffness alone changes keratocyte activation and contraction [15]. In self-assembled three-dimensional constructs, normal corneal fibroblasts thickened their matrix in response to either TGF-β1 or β3, with markedly lower collagen III and α-SMA under β3 [16,17]. KC cells responded differently. They appeared myofibroblast-like in constructs, and TGF-β3 lowered collagen III without lowering α-SMA [11]. In printed collagen constructs, KC-cell α-SMA was higher at weeks 2 and 4 than at week 1, whereas no tested marker increased significantly in normal cells [18]. Which material directs the same stimulus toward matrix synthesis or myofibroblast transformation in KC cells remains unresolved. KC has no animal model that reproduces both the corneal phenotype and the pathophysiology [19]. A patient-derived three-dimensional system with directly comparable materials is needed to resolve this question.

Drugs that induce host matrix synthesis are given as courses, unlike the single procedure of cross-linking. The registered regimen of TGF-β3 with dexamethasone eyedrops is twice daily for 3 to 6 weeks and includes duration of effect as an objective [7]. Losartan inhibits the pathway and has been used for corneal fibrosis [20–22], including in one keratoconic cornea after cross-linking [23]. Dosing has a start and a stop, and its timing changes repair in lung injury, engineered heart tissue and dermal hydrogels [24–26]. For KC, efficacy depends on the host matrix that remains after a course ends and whether scarring stays low. Culture studies usually evaluate these drugs during dosing by the suppression of myofibroblast-associated readouts [27] or matrix protein content [12,28]. These surrogate measures must be tested against the endpoint before they can represent efficacy [29,30]. Whether measurements during dosing represent the matrix left after a course has not been tested. Pathway blockade lowers matrix synthesis along with scarring. The course that suppresses scarring most deeply may leave less matrix. Among 166 original studies of KC drugs and materials that we read and coded individually, only 4 of the 45 using repeated, continuous or stop-and-follow dosing measured deposited matrix or contraction. None measured either after the course was withdrawn.

We addressed these questions in patient-derived bioprinted stroma from six KC and six normal donors using gelatin methacryloyl (GelMA), a material already used to print corneal stroma [31]. Cells from each donor were prepared in matched two-dimensional (2D), cast and printed formats to distinguish the effects of the three-dimensional material from those of printing. Measurements of cell content and matrix output together, with and without stimulation, identify the source of low KC matrix output and test whether synthesis can be induced and whether induced KC output remains below normal.

Comparison of the increases in matrix formation and scarring after one TGF-β1 stimulus across the three formats tests whether the material separates them. We compare the start and stop of a course by delaying receptor blockade, stopping it after a finite course and directly measuring native collagen and scarring after withdrawal. These experiments test whether surrogate readouts during dosing represent the host matrix retained after withdrawal and whether the two select the same courses.

## Results

### Patient-derived GelMA stroma reveals preserved matrix-forming capacity in KC

Lower construct DNA accounted for the lower total procollagen output of KC cultures. We compared matrix production per construct and per unit DNA in primary cells cultured on plastic, cast in GelMA or printed as six-layer grids in 5.0% GelMA at 5 × 10⁶ cells/mL. Cast and printed constructs shared ink composition, thermal history and photocrosslinking (Fig. 1A). Viable cells were present throughout the sampled filament depth, and a 14-day concentration screen supported the 5% formulation (Fig. 1B and Fig. S2M–O). The 8%, 10% and 12% formulations reduced live-signal occupancy at multiple time points, whereas 4% and 6% were never lower than 5% (Fig. S2M). Scanning electron micrographs of the day-7 constructs are in Fig. 1C and Fig. S1G,H, and rheology and compression are in Fig. 1D and Fig. S1A–F.

**Fig. 1.**
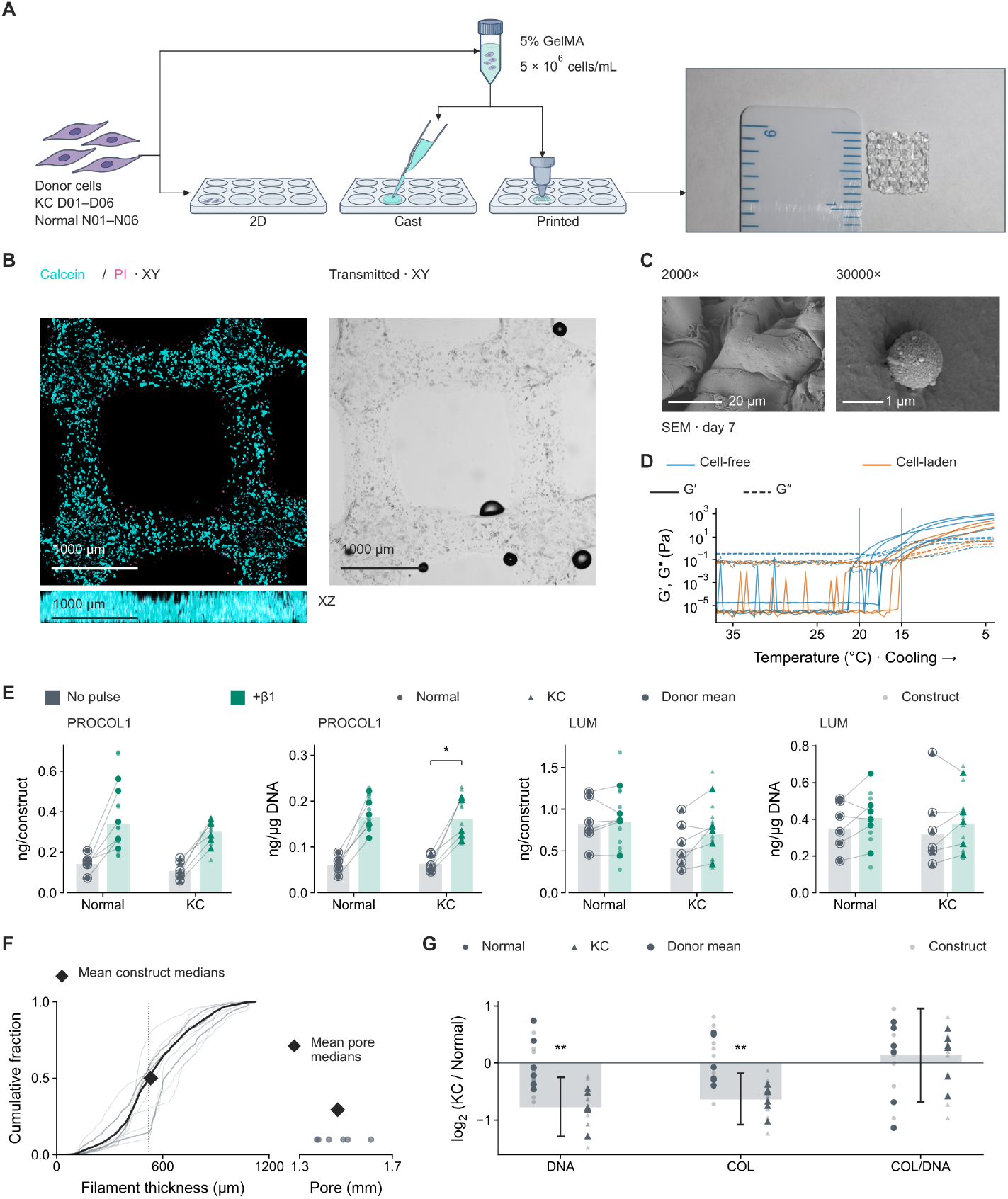
Patient-derived GelMA constructs preserve matrix production per unit DNA in keratoconus. (A) Donor-matched culture design and the printed construct. Primary stromal cells from six keratoconus (KC) and six normal donors were cultured on plastic in two dimensions (2D), cast in gelatin methacryloyl (GelMA) or printed as six-layer grids. Cast and printed cultures shared 5% GelMA, 5 × 10⁶ cells/mL, thermal history and photocrosslinking. (B) Transmitted-light and Calcein/propidium iodide (PI) imaging of one representative construct. Transmitted light is the center-z plane, Calcein and PI XY views are z-maximum projections and the XZ view is a y-maximum projection. Linear display ranges were 0–4095 for transmitted light and Calcein and 0–1761 for PI. Scale bars, 1000 μm. The depth distribution of live and dead signals and the 14-day GelMA concentration screen used to select the 5% formulation are in Fig. S2. (C) Scanning electron microscopy (SEM) at day 7, with two images of one field in one sample at 2,000× and 30,000×. Scale bars, 20 and 1 μm, respectively. (D) Storage modulus G′ and loss modulus G″ (Pa) during cooling from 37 to 4 °C in seven independent traces, comprising four cell-free and three cell-laden recordings. The modulus axis is logarithmic, and vertical references mark the syringe and platform temperatures of 20 and 15 °C. (E) Procollagen I (PROCOL1) and lumican (LUM) secretion during 48–72 h, expressed in ng per construct and ng per μg DNA, in six KC and six normal donors. No pulse denotes a vehicle pulse and +β1 a 4 h transforming growth factor-β1 (TGF-β1) pulse. Each assay includes 36 constructs, 12 vehicle and 24 pulsed. Bars are geometric means of six donor summaries per group and treatment. Donor summaries are arithmetic means of construct amounts or construct-specific DNA-normalized values, and lines join paired donor summaries. Rings mark donor means that coincide with a construct value. An asterisk denotes *p* < 0.05 for KC PROCOL1 per μg DNA after pulse versus vehicle (two-sided exact donor sign-flip test). (F) Empirical cumulative distributions of filament thickness (μm) for seven fields (light gray), three constructs (dark gray) and pooled positions (black), based on 7,159 measured positions. The pooled median (dotted line) was 521 μm and the mean of the three construct medians (diamond) was 532 μm. Inset, six complete pores from three constructs, with a mean construct median equivalent diameter of 1.46 mm (diamond). (G) Construct DNA, native type I collagen (COL) per construct and collagen per μg DNA at 336 h after a β1 pulse followed by vehicle, in 24 matched constructs from six KC and six normal donors. Matched log₂ quantities were averaged within donors and centered on the normal donor mean. Bars are log₂(KC/normal) differences between donor means, with Welch 95% confidence intervals. The horizontal zero line denotes the normal reference. Double asterisks denote *p* < 0.01 for KC versus normal DNA and collagen (exact donor-label permutation tests). Full rheology, compression and scanning electron microscopy data are in Fig. S1.

KC constructs secreted procollagen I and lumican at normal-donor levels per unit DNA, both without stimulation and after a TGF-β1 pulse (procollagen I, normal/KC 0.95 and 1.02; lumican, 1.09 and 1.08, Fig. 1E and Supplementary Data 1). Unstimulated normal constructs contained 1.40 times the DNA of KC constructs. Their 1.33-fold greater total procollagen output equaled the product of 1.40 for DNA and 0.95 for secretion per μg DNA. A 4 h pulse increased procollagen secretion per μg DNA 2.6-fold in all six KC donors (*p* = 0.031).

Native collagen retained in the material showed the same relationship to cell content. At 336 h after pulse onset, vehicle-treated KC constructs contained 65% of the collagen and 59% of the DNA of normal constructs, but 1.10 times the collagen per μg DNA (Fig. 1G and Table S4). KC cells retained matrix-producing capacity that the pulse could induce.

### Three-dimensional GelMA increases matrix induction and lowers baseline contractile expression

Three-dimensional GelMA increased matrix induction and reduced the contractile-gene response to the same TGF-β pulse. We compared donor-matched 2D, cast and printed cultures, defining the induced increase as expression after the pulse minus matched vehicle expression at 72 h (Table 1). In six KC donors, printed culture increased the response of FN1, COL1A1 and COL5A1 to 2.25 times that in 2D and reduced that of ACTA2, CCN2 and TAGLN to 39% of that in 2D (Fig. 2A and Supplementary Data 2). These contractile genes read out myofibroblast transformation. Every donor showed both effects. Cast GelMA reproduced the separation, with matrix and contractile responses of 2.01 times and 27% of those in 2D, placing the principal effect in the shared three-dimensional GelMA environment (Fig. 2A).

**Fig. 2.**
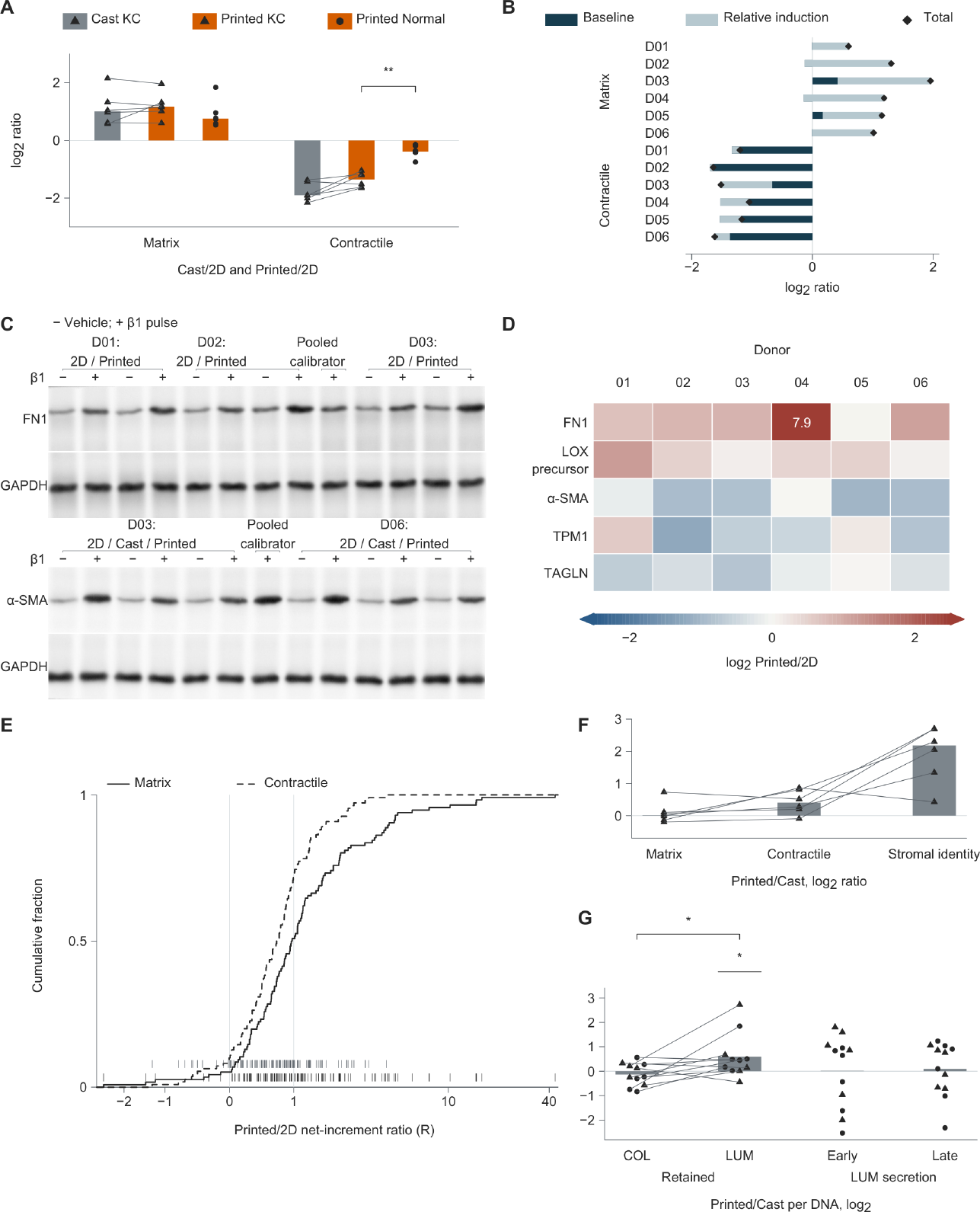
Three-dimensional GelMA separates matrix and contractile responses, and printing adds stromal identity and lumican retention. (A) Log₂ ratios of the transforming growth factor-β1 (TGF-β1)-induced increase in cast or printed gelatin methacryloyl (GelMA) cultures relative to two-dimensional (2D) cultures at 72 h after pulse onset. The induced increase is pulse-treated abundance minus matched vehicle abundance. Matrix genes are FN1, COL1A1 and COL5A1, and contractile genes are ACTA2, CCN2 and TAGLN. Printed comparisons include six keratoconus (KC) and six normal donors, and cast comparisons include the six paired KC donors. Points are 36 donor–program–comparison values and bars are medians. The printed KC–normal contractile comparison used an exact independent donor-label permutation test of the difference in group means (*p* = 0.0022, \*\**p* < 0.01). (B) Decomposition of printed/2D log₂ ratios into baseline and baseline-relative induction terms, whose sum is the total format effect. The 12 rows represent six KC donors in each of two programs. (C) Representative FN1 and α-smooth muscle actin (α-SMA) blots from three and two KC donors, respectively, with matched GAPDH strips. Each of the four continuous strips contains 13 lanes, including a pooled calibrator. Full membranes and lane assignments are in Supplementary Data 3. (D) Log₂ printed/2D ratios of induced protein increases in five protein rows and six KC donor columns (30 cells). Quantification uses DNA-equivalent loading and a same-membrane pooled calibrator. Colors saturate at −2.5 and 2.5, with out-of-range values printed in cells. (E) Exact empirical cumulative distributions and gene-level rugs (matrix, dark lower row; contractile, light upper row) of printed/2D response ratios for 116 matrix and 110 contractile genes in RNA sequencing from three KC donors. R is the printed induced increase divided by the positive 2D induced increase. The horizontal axis uses an inverse hyperbolic sine (asinh) scale labeled in signed R units. Vertical guides mark R = 0 (no printed increase) and R = 1 (equal increases). (F) Log₂ printed/cast ratios of induced increases in the matrix genes, the contractile genes and the stromal identity genes DCN, LUM and CHST6 in six paired KC donors. Points are 18 donor–program values, lines join the three programs within each donor, and bars are log₂-transformed medians of the linear ratios. (G) Log₂ printed/cast ratios of retained native type I collagen (COL) and lumican (LUM) per μg DNA at 336 h after pulse onset in constructs receiving vehicle after the pulse, in 11 paired donors (five KC and six normal), and of lumican secreted into the medium per μg DNA during 48–72 h (early) and 312–336 h (late) after pulse onset, in 12 paired donors (six KC and six normal). Bars are arithmetic means of donor log₂ ratios. Two-sided exact sign-flip tests compare retained LUM with zero (*p* = 0.017) and retained LUM minus COL with zero (*p* = 0.027), with \**p* < 0.05. Construct-level amounts in ng/construct (42 lumican and 44 collagen observations) are in Fig. S7D. Lines in (G) join the retained COL and LUM values of the same donor. In (F) and (G), triangles denote KC and circles normal donors. Zero reference lines in A, B, F and G denote no format difference.

**Table 1.**
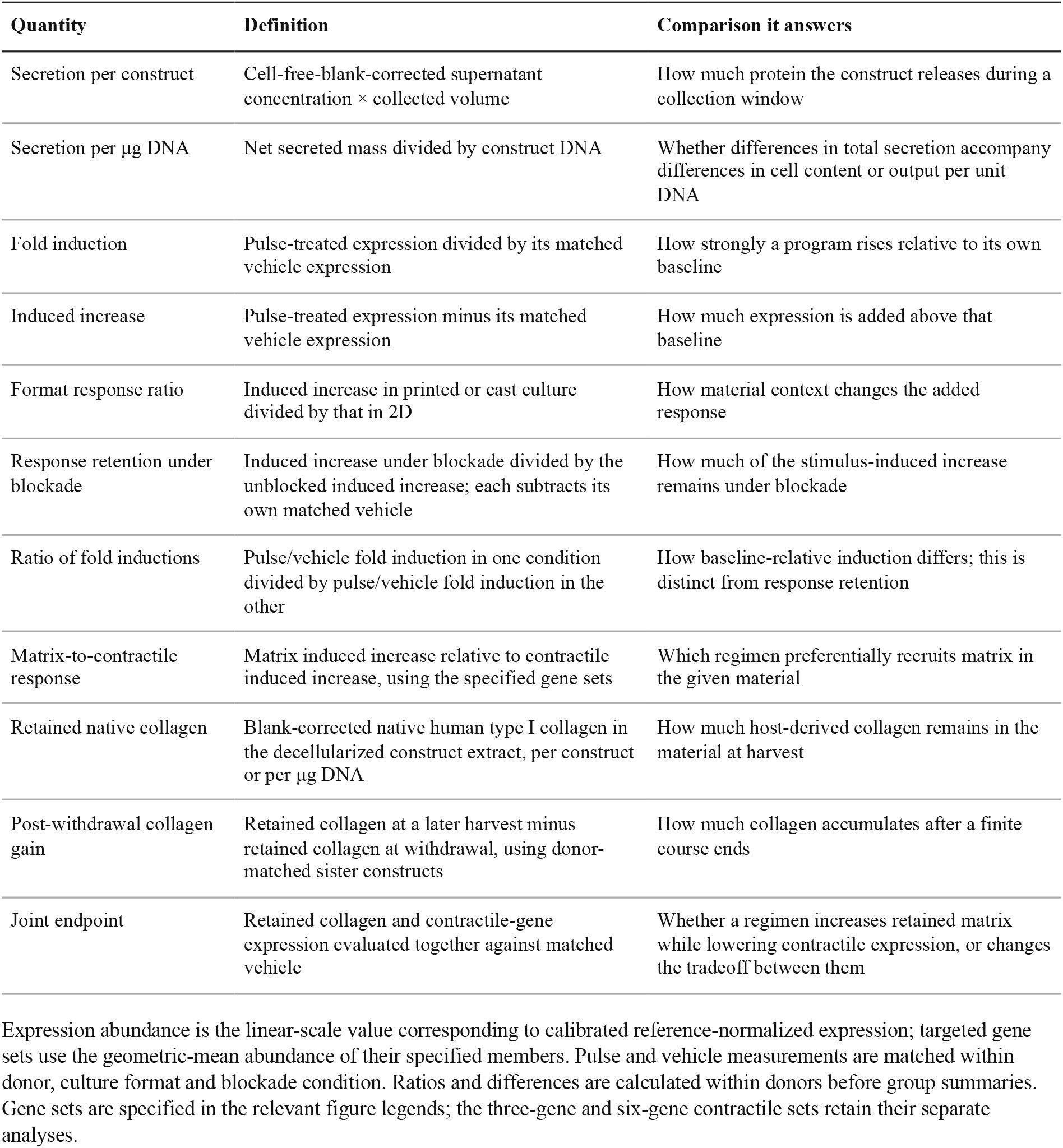
Quantities used to distinguish cell content, induced responses and retained matrix.

| Quantity | Definition | Comparison it answers |
|---|---|---|
| Secretion per construct | Cell-free-blank-corrected supernatant concentration × collected volume | How much protein the construct releases during a collection window |
| Secretion per µg DNA | Net secreted mass divided by construct DNA | Whether differences in total secretion accompany differences in cell content or output per unit DNA |
| Fold induction | Pulse-treated expression divided by its matched vehicle expression | How strongly a program rises relative to its own baseline |
| Induced increase | Pulse-treated expression minus its matched vehicle expression | How much expression is added above that baseline |
| Format response ratio | Induced increase in printed or cast culture divided by that in 2D | How material context changes the added response |
| Response retention under blockade | Induced increase under blockade divided by the unblocked induced increase; each subtracts its own matched vehicle | How much of the stimulus-induced increase remains under blockade |
| Ratio of fold inductions | Pulse/vehicle fold induction in one condition divided by pulse/vehicle fold induction in the other | How baseline-relative induction differs; this is distinct from response retention |
| Matrix-to-contractile response | Matrix induced increase relative to contractile induced increase, using the specified gene sets | Which regimen preferentially recruits matrix in the given material |
| Retained native collagen | Blank-corrected native human type I collagen in the decellularized construct extract, per construct or per µg DNA | How much host-derived collagen remains in the material at harvest |
| Post-withdrawal collagen gain | Retained collagen at a later harvest minus retained collagen at withdrawal, using donor-matched sister constructs | How much collagen accumulates after a finite course ends |
| Joint endpoint | Retained collagen and contractile-gene expression evaluated together against matched vehicle | Whether a regimen increases retained matrix while lowering contractile expression, or changes the tradeoff between them |

Expression abundance is the linear-scale value corresponding to calibrated reference-normalized expression; targeted gene sets use the geometric-mean abundance of their specified members. Pulse and vehicle measurements are matched within donor, culture format and blockade condition. Ratios and differences are calculated within donors before group summaries. Gene sets are specified in the relevant figure legends; the three-gene and six-gene contractile sets retain their separate analyses.

Matrix and contractile-gene expression changed in different ways (Fig. 2B). Matrix induction increased above a largely unchanged baseline. The smaller contractile increase arose predominantly from lower resting abundance of ACTA2, CCN2 and TAGLN, with little change in their baseline-relative induction. Both effects had the same direction in all six KC donors, and cast GelMA showed the same pattern. These changes also explained the stronger separation in KC. The printed/2D contractile-response ratio was 0.39 in KC and 0.77 in normal donors, with every KC donor below every normal donor (exact permutation *p* = 0.0022). Printed culture lowered the contractile baseline more in KC while preserving its matrix baseline; normal cells had a lower matrix baseline in printed culture (Table S4).

Protein measurements supported the gene-expression results (Fig. 2C,D and Supplementary Data 3). Printed culture increased the pulse-induced increments in FN1 and LOX precursor in all six KC donors, whereas the median printed/2D ratios for α-SMA, TPM1 and TAGLN were below 1 (Fig. 2D). The format effect on fold induction was greater for FN1 than for α-SMA (*p* = 0.031, Fig. S6B). Pulse-end SMAD2 phosphorylation was lower in printed culture in all three assayed donors (Fig. S6A), with greater matrix induction coexisting with a smaller SMAD2 response at that time point.

Transcriptome-wide comparisons also showed stronger responses in matrix-annotated than contractile gene sets (Fig. 2E). TGF-β2 and TGF-β3 gave the same overall pattern (Fig. S6E,F). Under TGF-β1, 1,238 genes showed a significant format-by-response interaction, of which 701 responded with opposite sign in 2D and printed culture (Table S1). GelMA changed both baseline expression and the response to stimulation.

### Printing adds a stromal identity response and preferential lumican retention

Printing increased the TGF-β1-induced response of DCN, LUM and CHST6, representing corneal proteoglycan and keratan sulfate synthesis, 4.54-fold over casting in six paired KC donors (Fig. 2F). Cast GelMA reproduced the principal separation between matrix and contractile responses, and comparison with the same material formed by printing identified the additional contribution of fabrication. The core matrix response remained at 101% of the cast response (Fig. 2F), indicating that printing added a stromal identity program to the matrix response supported by GelMA.

Printing also changed the composition of the retained matrix. At 336 h, printed constructs retained 1.51 times the lumican per μg DNA of paired cast constructs (10/11 donors, *p* = 0.017, Fig. 2G), although DNA-normalized lumican secretion was similar in the early and final collection windows (Table S4). Within donors, the printed/cast gain in retained lumican was 1.66 times the corresponding gain in native type I collagen (*p* = 0.027, Fig. 2G). The stromal identity response was accompanied by preferential lumican retention.

### The induced response runs opposite to KC tissue-associated expression and includes structural collagen genes

In GelMA, the TGF-β pulse shifted expression away from the KC tissue state and induced structural collagen genes. All six tissue-derived signatures of genes downregulated in KC were also lower in KC than in normal printed constructs, and the core matrisome was reduced (Fig. 3B and Table S1). An independently derived signature of proteins reduced in KC corneal cones showed the same direction. The constructs retained this broad KC-associated expression state even though resting abundance of the targeted matrix genes FN1, COL1A1 and COL5A1 was not lower in printed KC than normal constructs (1.39-fold, Table S4). Printed culture increased all six KC-downregulated tissue signatures and the cone-protein signature relative to 2D (Fig. 3B).

**Fig. 3.**
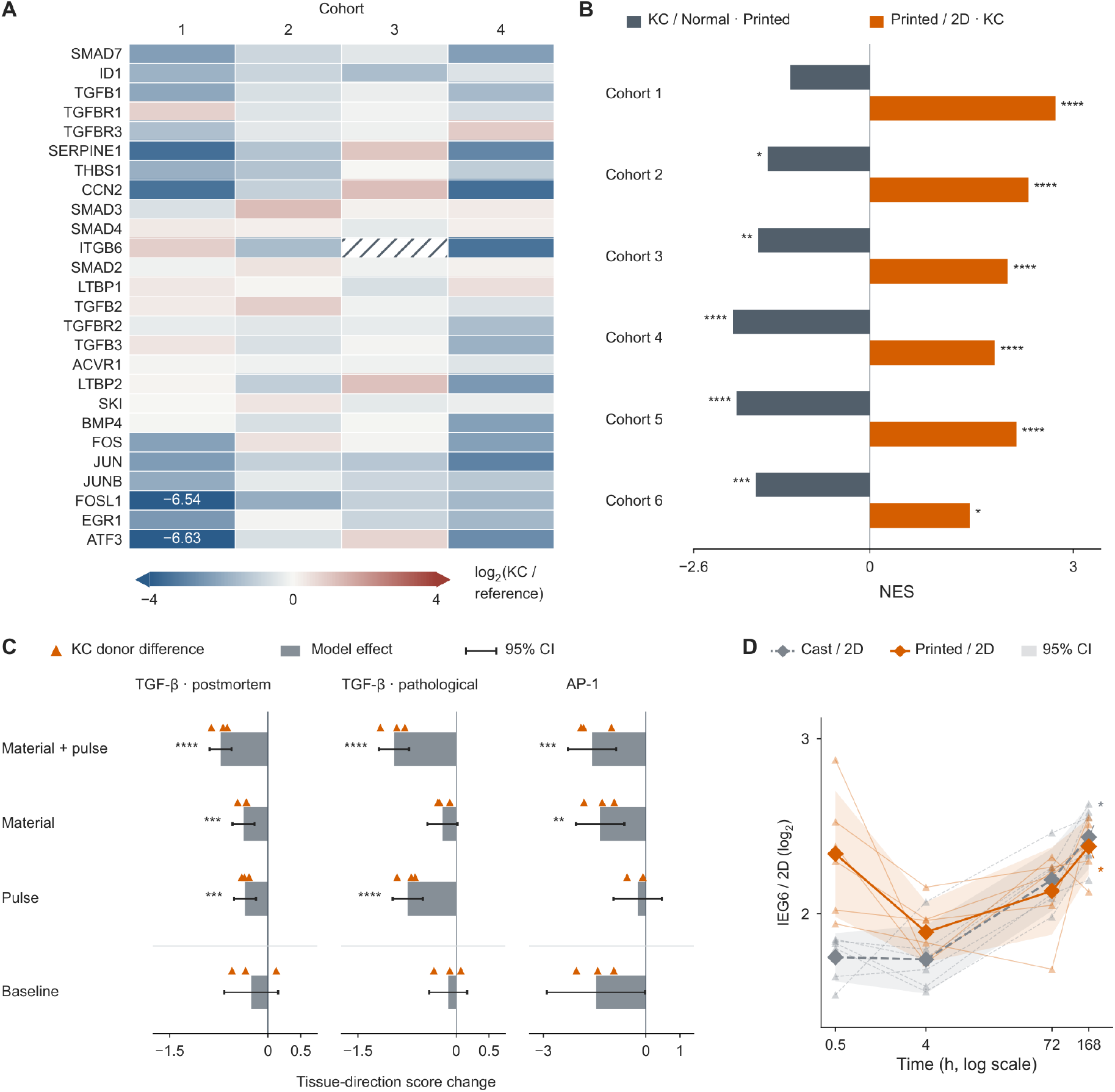
Patient-derived constructs retain keratoconus (KC)-associated expression that shifts in the opposite direction with material context and stimulation. (A) KC-versus-reference log₂ fold changes for 26 transforming growth factor-β (TGF-β)-pathway and immediate-early genes (rows) across four cohorts (columns). KC/reference sample counts are 19/7 for GSE151631, 8/7 for GSE204791 stroma, 6/8 for PRJNA1184491 stroma and 25/25 for GSE77938. The first three cohorts use postmortem controls and GSE77938 pathological controls. Colors span −4 to 4, centered at zero and capped, without row standardization. Hatching marks missing ITGB6 in PRJNA1184491. (B) Normalized enrichment scores (NES) for six KC-downregulated tissue signatures in two comparisons (12 bars), with three baseline donors per disease group. Cohorts 1–4 correspond to (A), and cohorts 5–6 are GSE303036 and GSE241715. Stars denote Benjamini–Hochberg-adjusted q values from preranked gene set enrichment analysis. (C) Tissue-direction score changes for TGF-β (20 genes in stimulation, 19 at baseline) and the six-gene activator protein-1 (AP-1)/immediate-early program, with 12 model contrasts and 36 donor differences (three KC donors per contrast). Material + pulse compares printed transforming growth factor-β1 (TGF-β1) with two-dimensional (2D) vehicle, Material compares printed vehicle with 2D vehicle, and Pulse compares TGF-β1 with vehicle in printed culture. Baseline is printed versus 2D in a separate unstimulated batch. Models use three donors for stimulation and six for baseline. Error bars are model-based 95% t confidence intervals (CIs). Stars denote two-sided donor-adjusted linear-model t tests with Benjamini–Hochberg adjustment. In (B, C), \**q* < 0.05, \*\**q* < 0.01, \*\*\**q* < 0.001 and \*\*\*\**q* < 0.0001. Vertical lines mark zero in (B, C). Negative values indicate movement away from the KC tissue state. The TGF-β blocks share a scale, and AP-1 has its own scale. The genetic and injury-repair comparisons are given in Table 2 and Fig. S9, respectively. (D) Mean log₂ expression differences for six immediate-early genes (IEG6: ATF3, EGR1, FOS, FOSL1, JUN and JUNB) relative to paired 2D cultures at 0.5, 4, 72 and 168 h after fabrication without TGF-β. Six KC donors provide 48 differences. Triangles and thin lines represent donors, diamonds represent eight means, and bands are their 95% t CIs. Time is logarithmic. At 168 h, *p* = 0.03125 for each format versus paired 2D (two-sided exact sign-flip test, \**p* < 0.05).

**Table 2.**
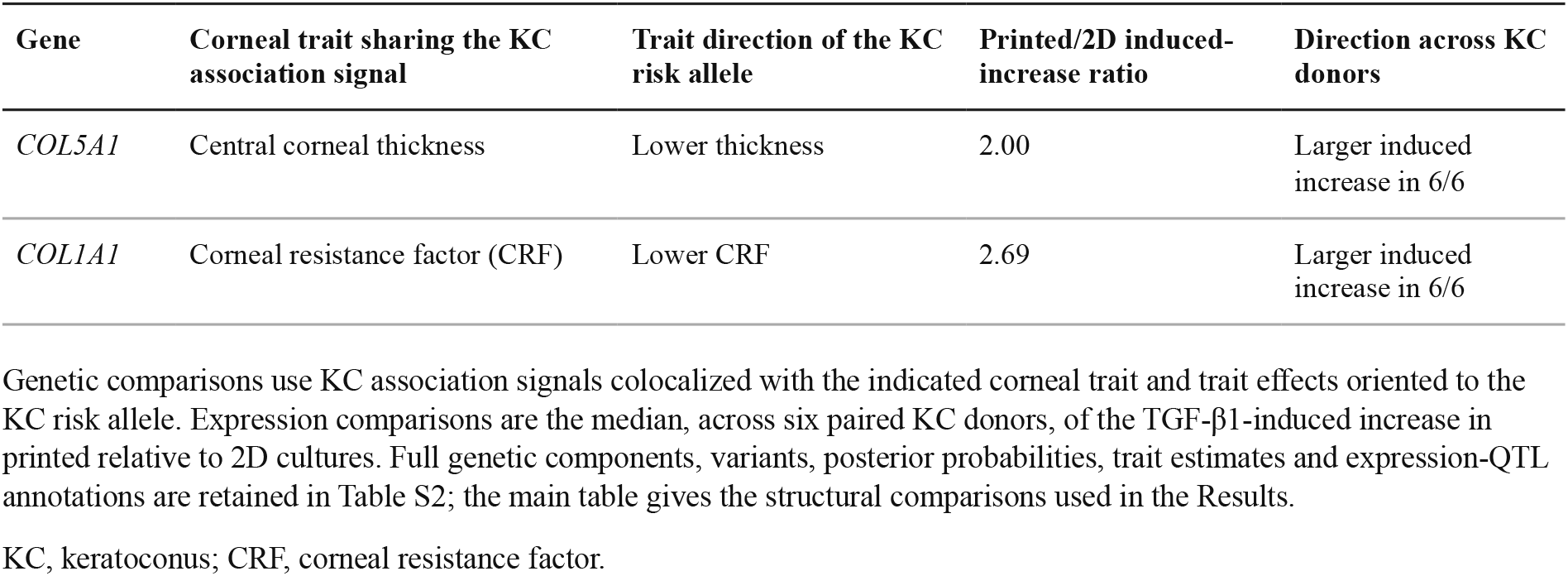
Structural collagen genes link the material-induced response to corneal traits associated with KC.

| Gene | Corneal trait sharing the KC association signal | Trait direction of the KC risk allele | Printed/2D induced-increase ratio | Direction across KC donors |
|---|---|---|---|---|
| *COL5A1* | Central corneal thickness | Lower thickness | 2.00 | Larger induced increase in 6/6 |
| *COL1A1* | Corneal resistance factor (CRF) | Lower CRF | 2.69 | Larger induced increase in 6/6 |

Genetic comparisons use KC association signals colocalized with the indicated corneal trait and trait effects oriented to the KC risk allele. Expression comparisons are the median, across six paired KC donors, of the TGF-β1-induced increase in printed relative to 2D cultures. Full genetic components, variants, posterior probabilities, trait estimates and expression-QTL annotations are retained in Table S2; the main table gives the structural comparisons used in the Results.
KC, keratoconus; CRF, corneal resistance factor.

After the pulse, TGF-β and AP-1 tissue-direction scores shifted away from the KC tissue state in all three sequenced KC donors, with much of the AP-1 shift already produced by the material (Fig. 3C). TGF-β-pathway and immediate-early genes were predominantly lower in the patient-tissue cohorts (Fig. 3A). Cast and printed GelMA increased immediate-early expression from the first fabrication time point through day 7, and the additional early effect of printing over casting subsided during culture (Fig. 3D and Fig. S3). Acute targets and feedback genes remained inducible. SMAD7, PMEPA1 and SERPINE1 were induced at 4 h in all six KC donors, and SMAD7 remained induced at 72 h (Table S4). Patient-derived constructs remained responsive despite the lower pathway-associated expression in KC tissue.

The pulse induced genes at structural susceptibility loci. COL5A1 and COL1A1 had KC association signals colocalized with central corneal thickness and corneal resistance factor, respectively. Their median pulse-induced increases were 2.00 and 2.69 times those in 2D, respectively, with the printed increase larger in all six KC donors. Their KC risk alleles reduced these structural traits (Table 2 and Table S2), linking the induced collagens to structural traits impaired in KC.

Public repair references distinguished matrix production from contractile activation. Fibroblasts repopulating freeze-injured rabbit stroma increased matrix programs without the contractile activation of fibrotic remodeling cells (Fig. S9A). Printed KC constructs retained more of this repopulating program after the pulse than 2D cultures did (Fig. S9 and Table S3). Matrix and stromal identity programs were also elevated in printed constructs before stimulation, and collagen-processing enzymes were more strongly induced than in 2D (Fig. S8B and Table S1). These comparisons gave the enhanced matrix response a structural role in KC distinct from the accompanying contractile program.

### Delaying receptor blockade until 24 h preserves matrix induction and late secretion

Delayed receptor blockade preserved more matrix induction while suppressing contractile-gene expression. GelMA alone reduced the contractile response but did not abolish its induction. Both COL1A1 and ACTA2 were already induced in all six KC donors by 24 h (Table S4). After the 4 h TGF-β1 pulse, we applied SB431542 immediately after washout, from 4 to 72 h, or after a delay, from 24 to 72 h [32] (Fig. 4A). SB431542 inhibited signaling during a TGF-β1 challenge in both 2D and printed culture (Fig. S6C) and reduced p-SMAD2/total SMAD2 to 68% of its predose level within about 1 h of delayed dosing (Fig. S6D).

**Fig. 4.**
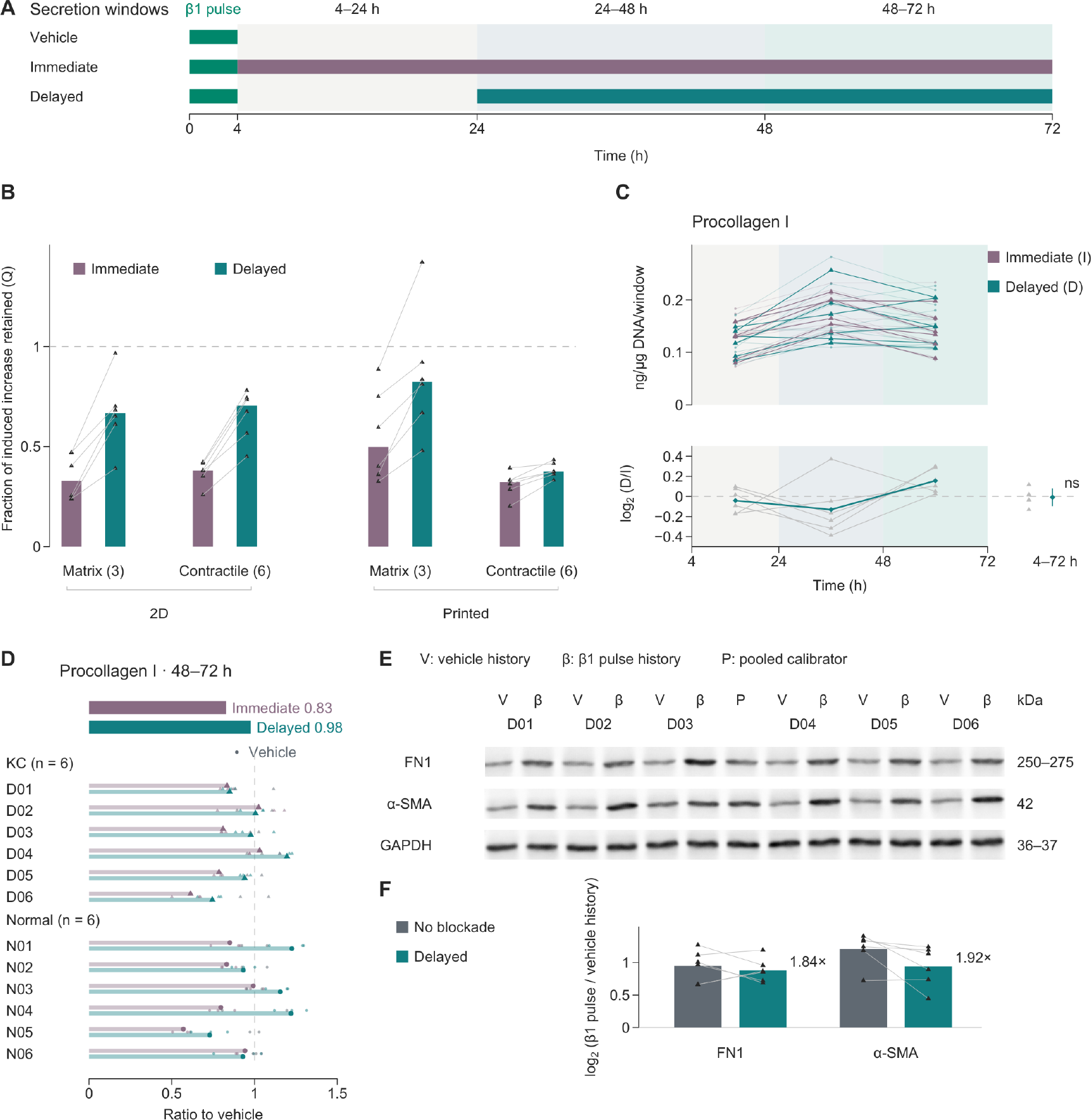
Delaying receptor blockade preserves matrix induction and late procollagen secretion. (A) Treatment schedules after a 4 h transforming growth factor-β1 (TGF-β1) pulse: vehicle, immediate SB431542 at 4–72 h and delayed SB431542 at 24–72 h. All times are from pulse onset. (B) Fraction of the unblocked TGF-β1-induced increase retained (Q) under immediate or delayed blockade in paired two-dimensional (2D) and printed cultures from six keratoconus (KC) donors. The numerator and denominator each subtract their blockade-matched vehicle. Matrix (3) comprises FN1, COL1A1 and COL5A1 and Contractile (6) comprises ACTA2, TAGLN, TPM1, CCN2, LOX and CCN1, with gene counts in parentheses. A value of 1 (dashed line) denotes full retention of the unblocked increase. Bars show the median Q for each format, gene set and blockade schedule, triangles show the six KC donors, and lines join the two schedules within each donor. (C) Procollagen I secreted per μg DNA during the 4–24, 24–48 and 48–72 h windows in 24 matched printed constructs from six KC donors (upper), with delayed/immediate (D/I) ratios for each window and for cumulative 4–72 h secretion (lower and right). Amounts are per window, not rates. Small symbols are constructs, triangles are donor means and diamonds are six-donor mean log₂ ratios. The cumulative mean log₂ ratio is −0.008 (95% CI −0.094 to 0.078, two-sided exact sign-flip *p* = 0.75, ns). (D) Procollagen I secretion per μg DNA during 48–72 h in printed cultures from six KC and six normal donors, relative to each donor’s own vehicle mean. Treatments are paired within donors, small symbols are constructs and large symbols are donor means. The summary bars give the geometric means of the 12 paired donor ratios, 0.83 for immediate and 0.98 for delayed blockade. (E) FN1, α-smooth muscle actin (α-SMA) and GAPDH immunoblots after vehicle (V) or β1 pulse (β) histories followed by delayed blockade in six KC donors, with a pooled calibrator (P). Continuous native membrane strips are shown with the expected molecular masses (kDa) from the antibody datasheets. Full membranes and lane assignments are in Supplementary Data 3. (F) Pulse-induced FN1 and α-SMA protein in the same six KC donors without blockade or with delayed blockade. Abundance is normalized to the pooled calibrator after DNA-equivalent loading, bars show the mean log₂ ratio of β1-history to vehicle-history lanes (geometric-mean inductions without and with delayed blockade: 1.93-fold and 1.84-fold for FN1, 2.31-fold and 1.92-fold for α-SMA), triangles show donors and lines join the two conditions within each donor. Because the vehicle is the one matched to each condition, induction here differs from the induced-increase retention in (B).

In printed constructs, immediate and delayed blockade retained 32% and 38% of the expanded six-gene contractile response (the three genes above plus TPM1, LOX and CCN1), respectively (Fig. 4B). Immediate blockade retained 50% of the matrix-gene induced increase, whereas delayed blockade retained 82% (Fig. 4B). In 2D, delayed blockade retained 67% of the matrix and 70% of the contractile response, suppressing the two together (Fig. 4B). Under the same delayed regimen, printed cultures produced 3.22 times the matrix increase and 46% of the contractile increase of 2D, with both effects in all six donors (Table S4). Earlier blockade gained additional contractile suppression at the cost of matrix induction.

Delayed blockade also preserved late protein secretion. Serial collections from the same KC constructs showed that the delayed-over-immediate advantage developed in the late window, while cumulative secretion was unchanged (Fig. 4C). In printed constructs from six KC and six normal donors, procollagen I secretion per μg DNA during 48–72 h was 98% of vehicle after delayed blockade and 83% after immediate blockade (Fig. 4D). Both regimens included blockade throughout this collection window. The regimens differed only in blockade during 4–24 h, yet subsequent matrix output differed under the same late-window drug exposure.

Matrix transcription continued during delayed blockade. Pre-FN1 and pre-COL1A1 retained their induced increases at 72 h (Table S4), and FN1 protein remained inducible, increasing 1.84-fold with delayed blockade compared with 1.93-fold without blockade (Fig. 4E,F and Fig. S6B). α-SMA protein, which had a higher resting level under blockade, also remained inducible (Table S4). A 20 h unblocked interval after pulse washout preserved matrix production and ongoing transcription during subsequent receptor blockade. Across both disease groups, delayed treatment produced 1.36 times (cast) and 1.18 times (printed) the late-window secretion of immediate treatment (*p* = 0.003 in printed constructs, Fig. S7E).

### After withdrawal, delayed finite blockade preserves matrix expression and collagen accumulation

Delayed blockade redistributed procollagen secretion toward the late window without raising cumulative secretion (Fig. 4C), and we measured what it left in the matrix after the course ended. A separate batch supplemented with ascorbic acid 2-glucoside compared vehicle after the pulse, immediate blockade stopped at 72 h, delayed blockade stopped at 72 h and delayed blockade continued to 336 h (Fig. 5A and Fig. S5). Eleven days after withdrawal, delayed finite blockade had left more native collagen and lower contractile-gene expression than vehicle in 9/12 donors, including four KC and five normal donors (Table S4, Four_arm_dual_endpoint). Across donors, collagen was 1.25-fold higher (*p* = 0.054), and the geometric-mean expression of ACTA2, CCN2 and TAGLN was 67% of vehicle (each gene *p* ≤ 0.0054).

**Fig. 5.**
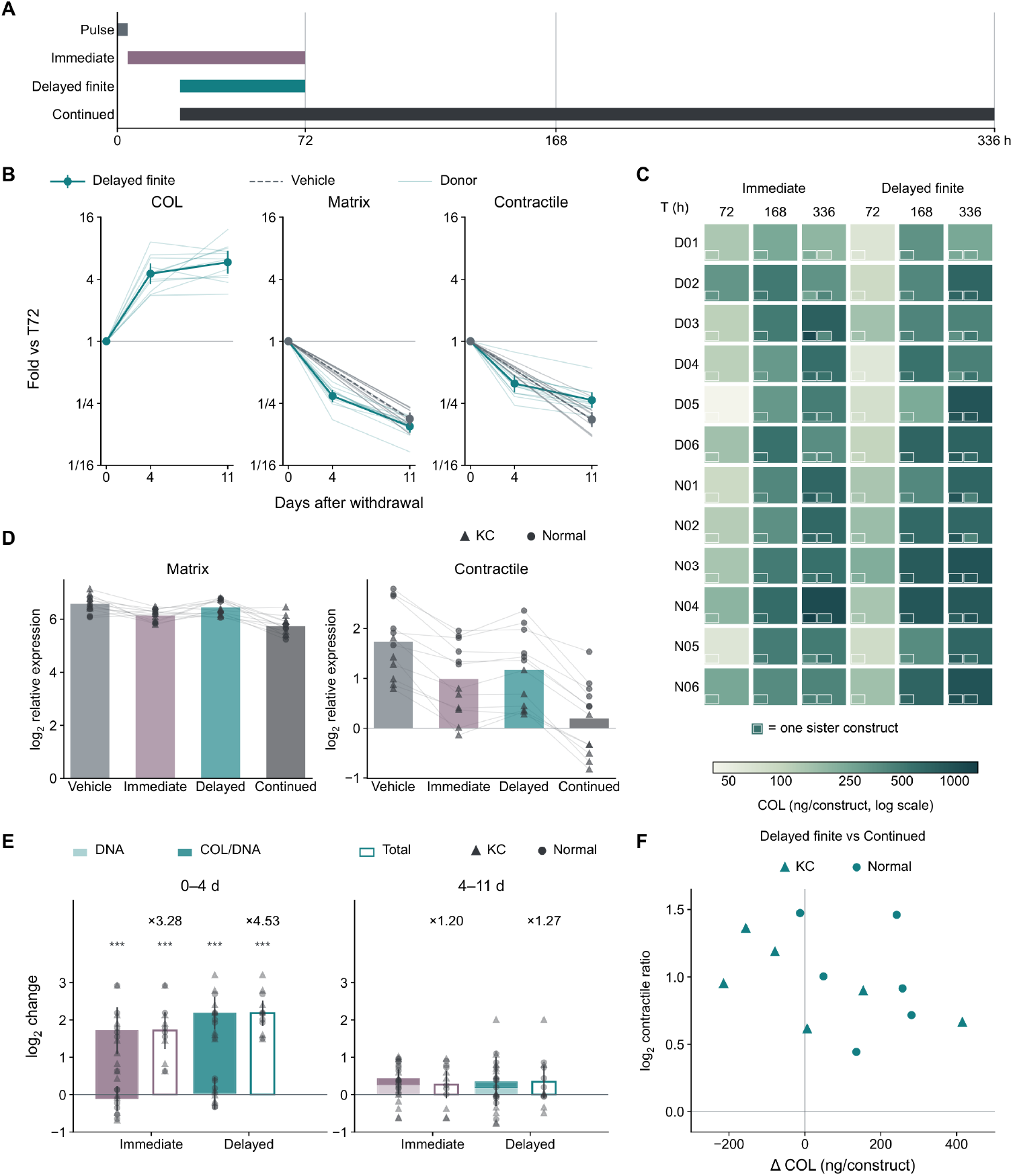
Delayed finite blockade preserves matrix expression and collagen accumulation after withdrawal. (A) Treatment schedules after a 4 h transforming growth factor-β1 (TGF-β1) pulse. Immediate blockade was given at 4–72 h, delayed finite blockade at 24–72 h and continued blockade at 24–336 h. T denotes hours from pulse onset. T72, T168 and T336 correspond to 0, 4 and 11 days after withdrawal for the finite courses. Continued treatment was still present at T336. (B) Native type I collagen (COL) per construct, matrix-gene expression (FN1, COL1A1 and COL5A1) and contractile-gene expression (ACTA2, CCN2 and TAGLN), relative to each arm’s own T72 level on logarithmic y axes. The 12 donors comprise six keratoconus (KC) and six normal donors. Each time point uses sister constructs. Points show geometric means with 95% confidence intervals, and fine lines show paired donor trajectories. The horizontal reference denotes the T72 level. Vehicle expression is shown at days 0 and 11. (C) Retained COL in 12 donor rows (D01–D06, KC; N01–N06, normal) and six regimen–harvest columns, with 72 arithmetic donor means and 96 sister-construct tiles. Columns show immediate and delayed finite blockade at T72, T168 and T336. Both layers use a logarithmic color scale spanning 40.76–1369.53 ng/construct without capping. Collagen per μg deoxyribonucleic acid (DNA) is shown in Fig. S10E. (D) T336 expression under the four regimens, as mean calibrated log₂ expression across the three genes in each set. The eight bars show unweighted means of 12 donor values, with lines joining paired donors. Zero is the assay calibration reference. (E) Decomposition of total COL changes into DNA and COL/DNA changes during 0–4 and 4–11 days after withdrawal under immediate and delayed finite blockade, in the same 12 donors. Construct-level log₂ values were averaged within donors before paired endpoint differences were taken. Components add on the log₂ scale. Bars show equal-donor means with 95% Student’s t confidence intervals for each component, offset by the mean DNA change for COL/DNA. Donor points show DNA, cumulative DNA plus COL/DNA and total COL changes. Outlined bars show total COL change, with the fold change labeled above. The horizontal reference denotes no change. For total COL and COL/DNA during 0–4 days in each regimen, *** denotes two-sided exact sign-flip *p* = 0.00048828125 versus no change. (F) Joint comparison of delayed finite versus continued blockade at T336, with one point per donor (n = 12). Horizontal positions are delayed finite minus continued COL (ng/construct), and vertical positions are log₂ delayed finite/continued ratios of contractile-gene expression. Zero lines denote equality. The comparison of delayed finite blockade with vehicle is in Fig. 8D.

Native collagen continued to accumulate after withdrawal, principally during the first four days (Fig. 5B,C). Over this interval, collagen per construct increased 4.53-fold after delayed blockade and 3.28-fold after immediate blockade, with increases in all 12 donors (Fig. 5B). Collagen accumulation was 140 ng greater per construct after delayed than after immediate blockade (*p* = 0.031, Fig. 5C). At 336 h, delayed treatment retained 1.43 times the collagen of immediate treatment per μg DNA and 1.29 times per construct (Fig. 5C and Fig. S10E). Over the full eleven days, collagen per construct after delayed blockade rose 5.86-fold from its level at withdrawal (95% CI 4.53–7.57, 12/12 donors, Fig. 5B). Matrix- and contractile-gene expression fell to 15% and 27% of their withdrawal-time levels, each in all 12 donors, while vehicle expression fell to a similar degree (18% and 17%). The delayed-to-vehicle ratio of contractile expression was 0.44 at withdrawal and 0.67 at day 11, below 1 in 12/12 and 11/12 donors (Table S4, Post_withdrawal_trajectory). These within-donor comparisons used sister constructs collected at each time point.

Withdrawal of delayed blockade left more collagen than its continuation in eight donors and less in four, while contractile expression was higher after withdrawal in all 12 (Fig. 5F). Continued blockade produced the lowest contractile expression and the lowest matrix-gene expression of the four arms, with reduced stromal identity transcription (Fig. 5D and Fig. S10A). Relative to continuation, delayed finite treatment retained 1.63 times the matrix-gene expression, including 2.00 times the COL5A1 expression, together with higher LUM and CHST6 expression (Fig. 5D, Fig. S10 and Table S4).

COL1A1, COL5A1, FN1 and LUM remained at vehicle levels after delayed finite treatment, while ACTA2, CCN2 and TAGLN were lower. Immediate treatment reduced COL1A1 and COL5A1, and continued treatment reduced all three matrix genes (Fig. S10). Delayed onset preserved matrix induction, whereas a finite course preserved matrix and stromal identity expression after withdrawal. Contractile expression remained below vehicle after the finite delayed course. Over the first four days, collagen per unit DNA accounted for the collagen increase, with little change in construct DNA (Fig. 5E).

### Additional collagen extends beyond individual nuclear neighborhoods

Additional collagen after delayed blockade extended from nuclear neighborhoods into the surrounding gel. Whole-mount confocal imaging at 336 h compared matched vehicle and delayed-blockade constructs from six donors (Fig. 6A–E and Supplementary Data 4). Delayed blockade increased collagen coverage of the sampled gel volume, and the 15–50 μm layers contributed 60.4% of that gain (Fig. 6C,D).

**Fig. 6.**
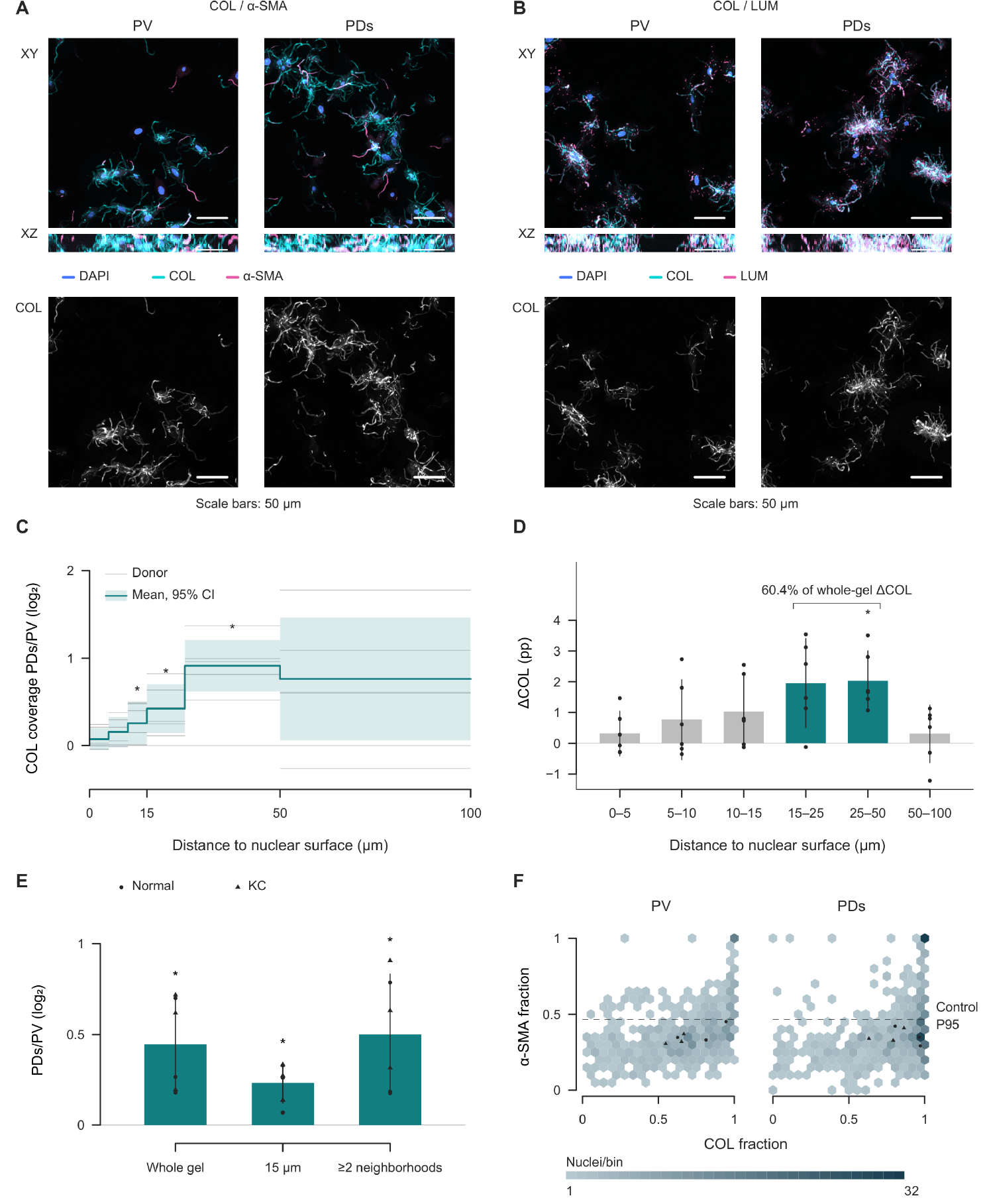
Collagen retained after delayed blockade extends beyond individual nuclear neighborhoods. (A) Whole-mount confocal images of collagen I and α-smooth muscle actin (α-SMA) from one keratoconus (KC) donor (D03) at 336 h after onset of a transforming growth factor-β1 (TGF-β1) pulse followed by vehicle (PV) or delayed blockade (PDs) withdrawn at 72 h, with one field per condition. (B) Collagen I and lumican in sister constructs from the same donor, with one field per condition. In (A, B), XY and XZ are maximum-intensity projections along Z and Y, respectively, and the lower row shows collagen alone. Display ranges are 0–4095 detector counts, except 0–1997 for α-SMA in (A) and 0–3950 for collagen in (B), with linear clipping and gamma = 1. Scale bars, 50 μm. Individual channels are shown in Fig. S11D. (C) Collagen coverage ratios (log₂ PDs/PV) in six shells from the nearest nuclear surface (0–5, 5–10, 10–15, 15–25, 25–50 and 50–100 μm), with 36 donor segments from six paired donors (three KC and three normal). (D) Additive contribution of each shell to the whole-gel collagen coverage increase in percentage points, with six bars and six donor values per bar. The 15–25 and 25–50 μm shells account for 60.4% of the whole-gel coverage increase (3.99 of 6.60 percentage points). (E) PDs/PV ratios (log₂) for collagen-positive gel volume fraction, median collagen coverage of extranuclear neighborhoods within 15 μm of the nearest nuclear surface, and gel volume fraction occupied by collagen-positive components spanning at least two nuclear neighborhoods (three bars, six paired donor contrasts each). In (C–E), the staining sets are pooled within donors, and steps and bars are arithmetic donor means with 95% Student’s t confidence intervals (band in C, whiskers in D and E). Donors are the statistical units. Zero reference lines denote equal PV and PDs values in (C, E) and no additive contribution in (D). Asterisks denote two-sided exact sign-flip tests of paired donor contrasts against zero, with *p* = 0.03125 for every marked comparison. (F) Collagen and α-SMA fractions in the same neighborhoods in PV and PDs, comprising 1,225 nuclei from 12 constructs and six donors. Color gives nuclei per bin in 346 occupied hexagons (shared linear scale, 1–32, capped at 32), and symbols show 12 construct medians (triangles, KC; circles, normal). The α-SMA gate is the no-primary-control 95th percentile (0.4654). Both axes show linear fractions from 0 to 1, with view limits of −0.04 and 1.04 for complete boundary hexagons. The volume distribution of collagen-positive connected components, lumican–collagen spatial measurements and additional α-SMA readouts are provided in Fig. S11.

Whole-gel coverage rose 1.36-fold and coverage within 15 μm of nuclear surfaces 1.18-fold, with both increases in every donor (*p* = 0.031, Fig. 6E). Collagen-positive components spanning at least two nuclear neighborhoods occupied 1.42-fold more gel volume after delayed blockade (*p* = 0.031, Fig. 6E, component size distribution in Fig. S11F). Lumican overlapped the retained collagen, and collagen coverage increased around nuclei in both α-SMA-defined states (Figs. 6B,F and S11B,C,H). The greater biochemically measured collagen retention included continuous collagen-positive regions beyond individual nuclear neighborhoods.

### Choosing a regimen in 2D selects a different course than the printed matrix does

The preferred regimen changed from unblocked TGF-β3 in 2D to delayed TGF-β1 blockade in printed culture. Courses are usually selected from responses measured during dosing in formats that differ from the repair material. These surrogate readouts differ from the repair goal, the host collagen retained after the course, in culture format, readout time and the object measured. Each step toward that goal changed the chosen regimen.

TGF-β1 followed by vehicle, immediate blockade or delayed blockade was compared with unblocked TGF-β3, which supports non-fibrotic matrix assembly in earlier three-dimensional corneal models [16,17]. Preference was based on matrix induction relative to contractile induction. The change in group preference produced a regimen-by-format interaction of 2.36-fold across six KC donors (95% CI 1.37–4.08, exact *p* = 0.031, Fig. 7A,B and Table S4).

**Fig. 7.**
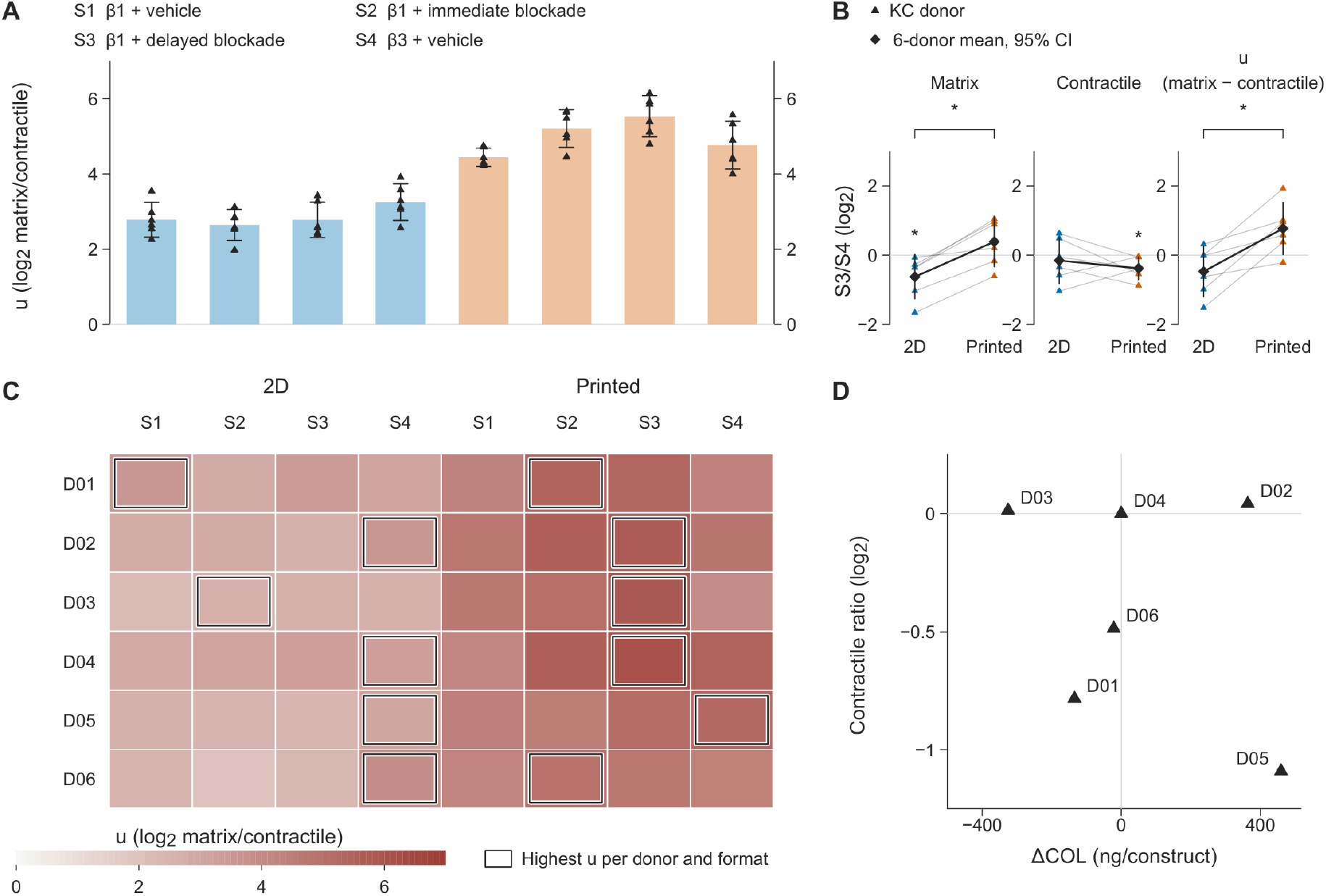
Culture format changes the preferred regimen and its subsequent collagen and contractile outcomes. (A) Matrix-to-contractile response ratio (u) on a log₂ scale in two-dimensional (2D) and printed cultures from six keratoconus (KC) donors under four regimens. S1 is a 4 h pulse of transforming growth factor-β1 (TGF-β1) followed by vehicle, S2 and S3 are the same pulse followed by SB431542 blockade at 4–72 h (immediate) and 24–72 h (delayed), respectively, and S4 is a 4 h TGF-β3 pulse followed by vehicle. Eight bars show unweighted means for S1–S4 in 2D (blue) and then printed cultures (orange), with triangles marking donors. The format and regimen labels identify both the bars in (A) and the columns in (C). Error bars in (A) and (B) are two-sided 95% t confidence intervals across six donors. The response ratio u compares induced increases in the three matrix genes FN1, COL1A1 and COL5A1 with those in the six contractile genes ACTA2, TAGLN, TPM1, CCN2, LOX and CCN1, using each regimen’s matched vehicle. (B) S3 versus S4 contrasts for matrix induction, contractile induction and u, with six paired donors per format and metric (36 donor-level contrasts). Values are donor-level log₂ S3/S4 ratios for each gene set and their difference for u. Thin lines join the same donor across formats, and thick lines join the group means. The horizontal zero lines denote equal responses to S3 and S4. Two-sided paired exact sign-flip tests compare S3 with S4 within formats and the S3/S4 contrast between formats. *p* < 0.05 (*) marks the matrix contrast in 2D, the contractile contrast in printed cultures and the bracketed regimen-by-format interactions for matrix and u. Per-donor responses and all S3/S4 estimates with their intervals are in Fig. S12C and D. (C) Donor-level u values for the same six donors, with one donor per row and four regimens per format across eight columns (48 cells). Colors show log₂ ratios on a common 0–7 scale, with no values clipped. Outlines mark the highest u within each donor and culture format. (D) Native type I collagen differences at 336 h (ng/construct) and contractile-expression ratios on a log₂ scale after applying the format-specific choices among the three TGF-β1 regimens (S1–S3) to the post-withdrawal batch. Each of six points represents a within-donor paired comparison. Collagen differences are calculated as the value for the regimen selected in printed culture minus that for the regimen selected in 2D. Contractile-expression ratios are calculated by dividing values for these regimens in the same order. Zero reference lines denote no collagen difference and a contractile-expression ratio of one. Here, the contractile set is ACTA2, CCN2 and TAGLN. D04 selects the same regimen in both formats. The three-regimen selection analysis is shown in Fig. S12.

The matrix response accounted for the interaction. In 2D, delayed TGF-β1 blockade produced 65% of the matrix increase of unblocked TGF-β3 in all six donors, whereas in printed culture it produced 1.31 times that increase (matrix interaction 2.02-fold, all six donors, Fig. 7B and Table S4). The 2D-preferred regimen applied in printed culture gave 23.7% less matrix-gene induction and 30.2% more contractile-gene induction than the printed-culture preference (Fig. S12D). The greater contractile increase occurred in all six donors. Regimen preference changed in five of six donors across all four regimens (Fig. 7C) and also in five of six when selection was restricted to the three TGF-β1 regimens (Fig. S12A,E).

Retained-matrix measurements carried the choices among the three TGF-β1 regimens through to the final endpoint. Selection by printed rather than 2D response changed native collagen by −325 to +459 ng per construct across the six KC donors at 336 h (Fig. 7D). Contractile expression was 73% of the 2D-selected value among the five switching donors, while matrix expression across all six remained at 103% (Table S4). The material used to select the regimen changed the collagen and contractile expression left after treatment.

### Measures taken during treatment and collagen retained after withdrawal select different courses

Selection solely by the depth of contractile suppression while the drug is present would choose continued blockade, which is still dosed at 336 h, in all 12 donors. That regimen produced the lowest contractile expression but reduced matrix expression relative to delayed finite treatment. The delayed course, once stopped, left 1.63 times the matrix-gene expression of continued blockade (Fig. 5D).

Secretion during treatment and collagen retained in the material also selected different schedules. Within the same post-withdrawal batch, 48–72 h procollagen secretion and native collagen retained at 336 h selected different immediate or delayed schedules in 6/12 donors per construct, comprising three KC and three normal donors. The corresponding count per μg DNA was 7/12 (Fig. 8A). All three switching KC donors moved from immediate to delayed blockade. Summed over the 12 donors, measured donor-level mean retained native collagen increased from 7,348 ng with secretion-selected schedules to 8,980 ng with retention-selected schedules, an additional 1,632 ng, or 22.2% (Fig. 8B). The increase was 28.8% in KC donors and 17.5% in normal donors (Fig. 8B).

**Fig. 8.**
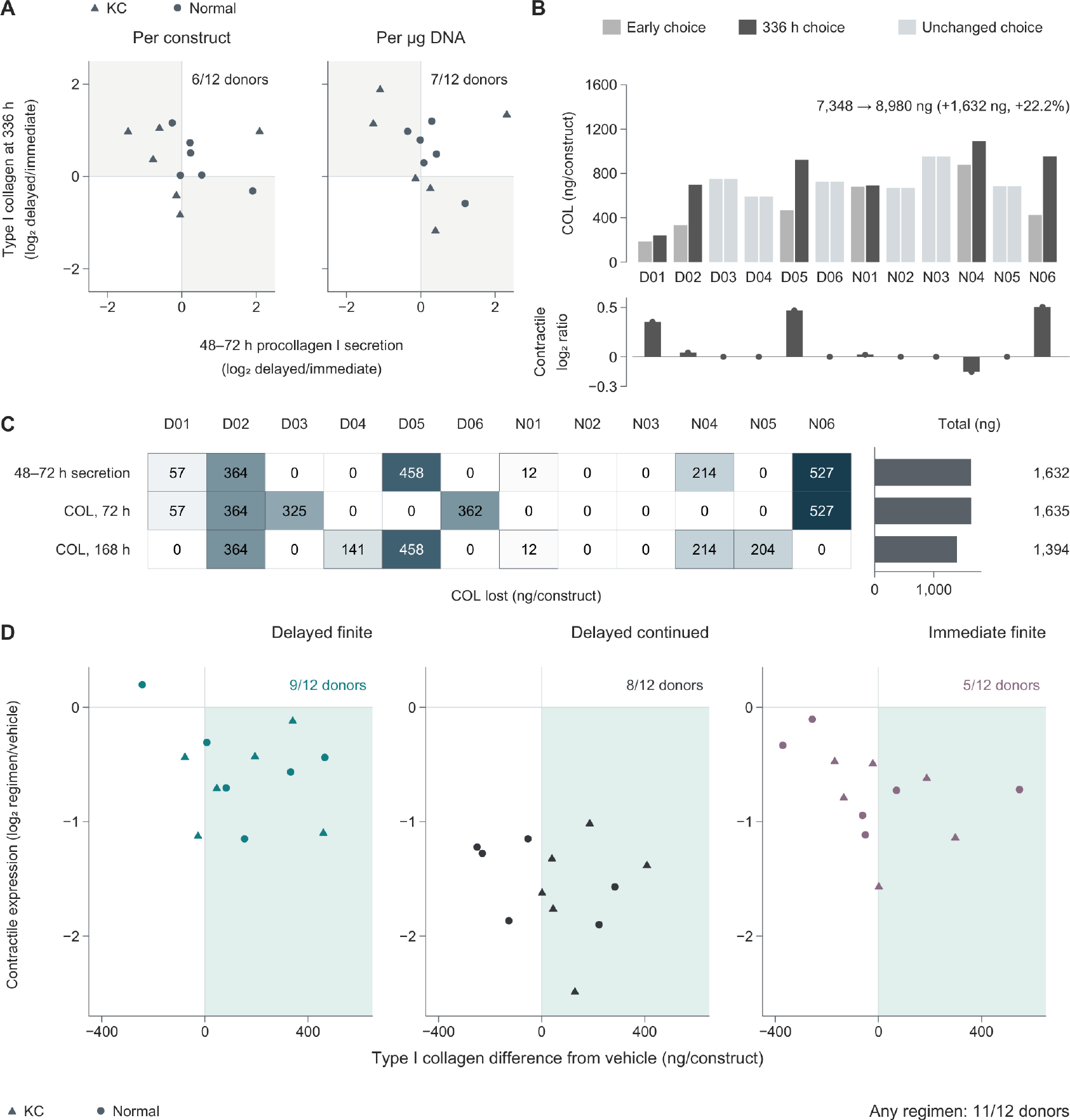
Retained collagen and contractile expression reveal consequences missed by earlier readings. (A) Within-donor delayed/immediate ratios for 48–72 h procollagen I secretion and retained native type I collagen (COL) at 336 h, each expressed per construct and per μg DNA on log₂ axes. Each scatter plot contains 12 points from six keratoconus (KC) and six normal donors. Immediate blockade spans 4–72 h and delayed finite blockade 24–72 h after pulse onset. The shaded quadrants mark discordant choices in 6/12 donors per construct and 7/12 per μg DNA. Zero reference lines denote equal values under the two regimens. (B) Native COL at 336 h under regimens selected by 48–72 h secretion or by COL at 336 h. The upper plot contains 24 bars showing paired donor means in ng/construct for the same 12 donors (D01–D06, KC; N01–N06, normal). The lower bars and dots show 12 log₂ ratios of contractile expression under the 336 h choice relative to the early choice. Contractile expression is the geometric mean abundance of ACTA2, CCN2 and TAGLN. The horizontal zero line denotes equal expression. The sums of the 12 donor means are 7,348 and 8,980 ng, respectively, a difference of 1,632 ng or 22.2%. Retained COL under immediate and delayed finite blockade is shown in Fig. 5C. The same two selections at withdrawal, day 4 and day 11 are in Fig. S10F. (C) COL at 336 h forfeited by selection from earlier readings, in ng/construct. The three rows represent selection from 48–72 h secretion, COL at 72 h and COL at 168 h, and the 12 columns represent donors, giving 36 cells. Each cell is the donor mean under the regimen selected at 336 h minus that under the regimen selected from the earlier reading, rounded to whole ng/construct. The choice is between immediate and delayed finite blockade. Colors scale linearly from zero to the maximum observed loss (527 ng/construct, rounded), without clipping. Dark borders identify changed choices, and zeros denote no loss. The three right-hand bars are sums of the 12 donor means, 1,632, 1,635 and 1,394 ng in row order. Source data are in Table S4. (D) Joint comparison of COL and contractile expression at 336 h between each blockade regimen and pulse followed by vehicle. Horizontal positions are paired differences in COL (ng/construct), and vertical positions are log₂ ratios of contractile expression, both relative to the same donor’s vehicle. Each plot contains 12 donor points, giving 36 donor–regimen pairs across delayed finite, delayed continued and immediate finite blockade. Zero reference lines denote equality with vehicle. The shaded quadrant marks more COL and lower contractile expression in 9/12, 8/12 and 5/12 donors, respectively, and 11/12 donors under at least one regimen. The continued regimen remains on treatment at 336 h. Donors are the statistical unit throughout.

With each donor’s two choices held fixed, the retention-selected schedules held 258 ng less native collagen than the secretion-selected schedules at withdrawal and 44 ng less four days later, but 1,632 ng more at 336 h (Fig. S10F). The retention advantage developed after the course ended. An earlier reading of the same analyte gave the same loss. The choice by native collagen at withdrawal differed from the 336 h choice in 5/12 donors and left 7,345 ng, and the choice four days later differed in 6/12 and left 7,587 ng, against 7,348 ng for the secretion choice and 8,980 ng for the 336 h choice (Fig. 8B,C and Table S4, Readout_time_choice). The secretion choice matched the collagen choice at withdrawal in 7/12 donors, and the loss came from when the course was read and not from the molecule measured.

### Retained collagen and contractile expression together identify courses that serve both repair goals

Against each donor’s vehicle, retention-selected schedules gave more native collagen and lower contractile expression in 10/12 donors, compared with 8/12 for secretion-selected schedules. Among the six donors whose choice changed, the counts were 5/6 and 3/6, and contractile expression under the retention-selected schedule remained below vehicle in all six (Table S4, Four_arm_dual_endpoint). Collagen alone did not settle the choice, because five of these six donors had higher contractile-gene expression under the retention-selected than under the secretion-selected schedule and one had lower expression (Fig. 8B). Joint evaluation of retained collagen and contractile expression extended this comparison to the full set of blockade regimens. At 336 h, 11/12 donors, including all six KC donors, had at least one regimen with more native collagen and lower contractile expression than vehicle (Fig. 8D and Table S4). Delayed finite treatment achieved both in four KC and five normal donors; continued treatment, still present at this endpoint, did so in six KC and two normal donors; and immediate finite treatment did so in five donors. 10/12 donors, five KC and five normal, had at least one of the two finite regimens, which end before the endpoint, with more native collagen and lower contractile expression than vehicle, and D01, the one KC donor outside them, reached it only under continued blockade. The regimen retaining the most collagen differed among donors.

The course chosen by retained collagen differed from those chosen by suppression or secretion during treatment. Joint measurement of collagen and contractile expression identified regimens that improved both and exposed the tradeoff when they did not improve together.

### Published KC-treatment studies do not assess matrix or contraction after withdrawal of repeated dosing

Across 166 human-coded original studies, no repeated or continuous dosing regimen was evaluated or selected using deposited matrix or contraction after dosing stopped (Table S4 companion files). The survey examined whether published KC-treatment evaluations connected the course to these measurements after withdrawal.

Of 45 studies with repeated, continuous or stop-and-follow dosing, four measured deposited matrix or contraction. These examined TGF-β3 with dexamethasone in keratocytes and corneal tissue [33], ISRIB in KC stromal cells [12], topical retinoid treatment in mouse corneas [34], and nintedanib in three-dimensional KC cultures [28]. None assessed deposited matrix or contraction after withdrawal of a repeated course. Fourteen studies included follow-up after repeated dosing stopped. Their subsequent readouts included clinical geometry, imaging, visual function, safety, RNA, enzyme activity, biomechanics and secreted protein, but no deposited-matrix or contraction endpoint. The ten studies that read deposited matrix or contraction after dosing stopped used single or brief exposures, such as cross-linking, matrix or tissue implants and short topical, soaking or culture applications, with no repeated course to withdraw (Table S4 companion files).

A separate coding of 40 full-text studies showed the same pattern in individual comparisons. None of 72 comparisons from 11 corneal pathway-inhibition and myofibroblast-reversal studies continued culture after inhibitor withdrawal before the outcome was assessed, and two studies counted lower type I collagen as an antifibrotic effect [35,36]. This reading indicates efficacy in fibrosis but points the opposite way in the thin KC stroma, where matrix has to be restored (Table S4). Published evaluations left the connection between a repeated course and the matrix or contraction remaining after withdrawal unmeasured. In our constructs, choosing schedules from secretion during dosing retained less native collagen than choosing them from the post-withdrawal measurement.

## Discussion

Low matrix output in KC reflects an inducible disease state with preserved synthetic capacity. Repair that retains the host cornea should stimulate its stromal cells to make matrix while limiting scarring. Our patient-derived three-dimensional GelMA model preserved the KC expression state and directed a single TGF-β stimulus toward matrix synthesis. With delayed, finite receptor blockade, the induced matrix remained as the host cells’ own native collagen after withdrawal. Surrogate readouts commonly used to select courses disagreed with the host matrix retained after treatment. Selection by secretion during dosing and by collagen retained after withdrawal differed in half of the donors. Across the 12 donors, selection by retained collagen left 22.2% more native collagen.

Low matrix synthesis in KC is a state of the diseased cells and not a capacity they have lost. The patient-derived model preserved this state. Tissue-derived signatures of downregulated genes [2] remained lower in KC than in normal constructs, yet procollagen I and lumican secretion per microgram of DNA matched that of normal cells. Normal constructs contained 1.40 times as much DNA, accounting for the lower total output of KC constructs. This agrees with the reduced keratocyte number in KC corneas [9,10]. The per-cell synthetic defect described in 2D culture [8] was absent here. The disease state can be reversed. The material and the pulse increased the TGF-β and immediate-early programs that are low in KC tissue [2] and induced COL5A1 and COL1A1, whose KC risk alleles thin and soften the cornea [37,38]. Induction ran opposite to the KC state. Drug-induced repair of KC stroma has a cellular basis, with the amount of repair limited by the cells that remain. A treatment course must then preserve the matrix those cells are induced to make.

The material directs the balance between TGF-β-induced matrix synthesis and scarring. Repair in vivo can add matrix without scarring [14], whereas cultured KC cells often appear myofibroblast-like [11,18]. The surrounding material accounts for this difference. In donor-matched printed GelMA, a single pulse produced a matrix-gene increase 2.25 times that in 2D culture. The corresponding ratio for ACTA2, CCN2, TAGLN and related genes, which read out myofibroblast transformation, was only 0.39. Separation was stronger in KC than in normal cells (0.39 versus 0.77). Cast GelMA produced the same separation, identifying the shared three-dimensional hydrogel environment as its source, consistent with the effect of substrate stiffness on corneal cell activation [15]. Printing additionally changed the identity of the induced matrix. DCN, LUM and CHST6 induction was 4.54 times that in cast constructs, and retained lumican was 1.51 times that in cast constructs. Lumican is required for regular collagen fibril arrangement and corneal transparency [39].

Treatment should induce the host matrix program while limiting scarring. Uniform suppression downstream of TGF-β reduces both. Immediate receptor blockade retained 50% of the pulse-induced matrix-gene increase, whereas blockade delayed to 24 h retained 82%. The retained contractile-gene increases were similar at 32% and 38%, respectively. The same delayed blockade suppressed both responses in 2D culture, showing that delay preserved the matrix bias established by the three-dimensional material. Dosing time also changes repair in lung injury, engineered heart tissue and dermal hydrogels [24–26]. Greene and colleagues expected deposited collagen to remain after treatment unless degraded [33]. Here, collagen continued to accumulate after withdrawal. At 11 days after withdrawal, 9 of 12 donors retained more native collagen and lower contractile-gene expression than vehicle, with the retained matrix supplied by the host cells themselves.

The further a surrogate lies from the repair goal, the less reliable the course it selects. The failure of surrogates to represent endpoints is a recurring problem in clinical trials [30]. We tested this directly in KC stromal repair. Pathway drugs in culture are usually evaluated during dosing by suppression of myofibroblast-related genes or proteins [27,35,36] and matrix protein content [12,28], mostly in 2D monolayers [12,27,35]. These readouts differ from the repair goal, the host collagen retained after a course, in culture format, measurement time and the object measured. Each step closer changed the selected course. In the printed matrix, which more closely reproduced the KC tissue state, the 2D-selected course produced 23.7% less matrix induction and 30.2% more contractile-gene induction. Regimen choice changed in 5 of 6 donors. Selection by contractile-gene suppression during dosing favored continuous blockade in all 12 donors. After withdrawal, the delayed finite course retained 1.63 times the matrix-gene expression of continuous blockade. Selection by secretion during 48–72 h and by retained collagen after withdrawal differed in 6 of 12 donors. The retention-selected courses held 258 ng less native collagen at withdrawal and overtook the secretion-selected courses only as collagen accumulated afterwards. A choice by native collagen itself at withdrawal or four days later left the 336 h choice 22.3% and 18.4% ahead, against 22.2% for secretion, and the time of reading, not the molecule measured, set the loss. Courses should be evaluated as close to the repair goal as possible, by native host collagen retained after withdrawal together with persistently low contractile-gene expression in a patient-derived three-dimensional model. This joint measurement is the closest in vitro readout of the repair goal.

Surrogate readouts are widely treated as efficacy measures in this field. In 11 studies of corneal pathway inhibition and myofibroblast reversal, none of the 72 comparisons continued culture after the inhibitor was withdrawn, and two studies counted lower collagen I as antifibrotic efficacy [35,36]. Collagen reduction indicates efficacy in fibrosis but opposes the need to restore matrix in KC. Among 166 studies of KC drugs and materials, none of the 45 using repeated, continuous or stop-and-follow dosing measured deposited matrix or contraction after withdrawal. Our joint evaluation identified at least one tested regimen that retained more native collagen and lower contractile-gene expression than vehicle in 11 of 12 donors, including all 6 KC donors. Every KC donor had a selectable regimen. The registered study of TGF-β3 with dexamethasone eyedrops lists duration of effect as an objective [7]. Losartan has been used for corneal fibrosis [20–22], including after cross-linking in a keratoconic cornea [23], and a placebo-controlled trial is registered for stable corneal scars [40]. The start, stop and duration of both treatments can be compared using these joint measurements in patient-derived three-dimensional material. KC lacks an animal model that reproduces both its corneal phenotype and pathophysiology, leaving treatment development dependent on clinical studies, ex vivo donor corneas and imperfect animal models [19]. A three-dimensional material that preserves the patient’s disease state fills this preclinical gap. After withdrawal, joint measurement identifies courses with the additional native collagen needed by a thin KC stroma and the lower contractile-gene expression needed by a scarred stroma.

## Conclusions

KC stromal cells retain inducible matrix-forming capacity in three-dimensional GelMA. The material enhances matrix induction and lowers baseline contractile expression. Printing adds stromal identity and lumican retention. The induced response opposes KC tissue-associated expression and includes structural collagen genes. Delayed, finite receptor blockade turns this capacity into a course that leaves more native collagen and lower contractile expression than vehicle in 9/12 donors eleven days after withdrawal. At the final endpoint, every KC donor has at least one regimen achieving both effects among all tested blockade regimens, including continued treatment. Culture format changes regimen preference. Selection by retained collagen instead of readouts during treatment leaves 22.2% more native collagen across the 12 donors. Courses of TGF-β-pathway treatment for keratoconic and scarred corneas should therefore be evaluated by the host matrix and contractile expression remaining after withdrawal. This patient-derived material system allows the start and stop of treatment to be controlled and both to be assessed together.

## Materials and methods

### Donors and primary corneal stromal cells

Keratoconus (KC) stromal tissue was obtained from host corneal buttons removed at keratoplasty, and normal stromal tissue from refractive-surgery corneas without apparent disease. Six KC donors (D01–D06) and six normal donors (N01–N06) contributed one isolate each. Printing, protein assays and sequencing used passage 3. RNA sequencing used the donor-derived isolates described above. The study received institutional ethics approval, donors gave informed consent, and procedures followed the Declaration of Helsinki. Fabrication-batch assignments are provided in the Supplementary Methods.

### GelMA–LAP ink, printing and rheology

The ink contained 5.0% GelMA, 0.25% lithium phenyl-2,4,6-trimethylbenzoylphosphinate (LAP) and 5 × 10⁶ cells/mL; the concentration screen used 4%–12% GelMA and one normal isolate. Ascorbic acid 2-glucoside (AA2G) was not added to the ink. Six-layer grids of nominal 8 × 8 × 1.2 mm were crosslinked once after deposition. Rheology compared cell-free and cell-laden ink, with the gel point defined as the last G′ = G″ crossing. Compressive modulus was measured over 10%–20% strain in molded cylinders and compressive strength in printed constructs. Fabrication and testing parameters are given in the Supplementary Methods.

### Culture formats

Cells were compared on tissue-culture plastic (2D), in cast GelMA and in printed GelMA. Paired cast and printed constructs shared ink composition, thermal history and crosslinking. The fabrication clock M0 was the end of crosslinking or of the paired 2D sham treatment; stimulation began on day 7, designated T0, with T denoting hours from pulse onset. The functional batch comprised 184 constructs from 12 donors, including 16 cell-free blanks, in DMEM/F-12 with 10% fetal bovine serum. The post-withdrawal batch comprised 536 constructs, including 44 cell-free blanks, and used a single serum lot with 0.5 mM AA2G from the first medium addition. Parallel bridging arms omitted AA2G. The imaging batch used the post-withdrawal medium and AA2G and ended at T336. Plate layouts, cell numbers and medium-change schedules are given in the Supplementary Methods.

### TGF-β pulses and receptor blockade

On day 7, cultures received 10 ng/mL TGF-β1, TGF-β2 or TGF-β3 for 4 h or matched vehicle. SB431542 (10 μM) was applied immediately after washout (4–72 h) or after a delay (24–72 h). All arms received matched 0.1% DMSO from 4 h. Functional-batch arms comprised no pulse with vehicle, a β1 pulse with vehicle, and β1 followed by immediate or delayed blockade, in printed and cast constructs. Supernatants were collected completely at 4–24, 24–48 and 48–72 h (W1–W3), and endpoint constructs supplied DNA.

The post-withdrawal batch compared vehicle after the pulse in printed and cast constructs, immediate blockade stopped at T72, delayed blockade stopped at T72, and delayed blockade continued to T336. Constructs were harvested at T72, T168 and T336 for collagen, lumican, RNA and DNA, corresponding to 0, 4 and 11 days after withdrawal for the finite regimens; the continued regimen remained on treatment at T336. Late-window secretion was collected from printed vehicle, immediate and delayed finite-treatment arms, with the final 24 h of secretion also collected at T336. The imaging batch comprised no-pulse printed constructs, printed vehicle, delayed finite blockade and cast vehicle at T336. Abbreviations appearing in the figures are mapped in the corresponding legends.

Supplementary qPCR experiments supplied the paired-format comparisons in Figs. 2A,B,F, 4B and 7A–C: six KC donors contributed 72 T72 samples, with 24 vehicle samples shared with the main arms, plus β3 and T24 arms. Signaling assays comprised pulse-end p-SMAD2/total SMAD2 at 4 h, a TGF-β1 challenge at 24 h in which SB431542 or solvent was applied for 1 h before 10 ng/mL β1 for about 45 min, and cultures with TGF-β1 history sampled before delayed SB431542 dosing and about 1 h afterward. Sample inventories and renewal schedules are provided in the Supplementary Methods.

### RNA extraction and quantitative PCR

RNA from whole constructs and 2D cultures was purified with the same silica-column chemistry and on-column DNase treatment. SYBR Green assays used primers designed with Primer3 [41] on MANE Select transcripts [42] and checked by Primer-BLAST [43] (Table S5). The supplementary batch comprised 276 samples, including 252 KC and 24 normal samples, across 51 assays. Plates included duplicate wells, no-template and no-RT controls, and a pooled-cDNA calibrator (Fig. S5).

Detection required Cq ≤ 35 and a single melt peak within 1.0 °C of the assay reference, with additional no-RT separation for JUN, JUNB and pre-mRNA assays. Passing duplicates were averaged, and absent passing wells were missing. Plate offsets were calibrator Cq minus the gene’s mean calibrator Cq. Assay efficiency E came from seven-level standard curves (10⁷–10¹ copies; R² ≥ 0.98 and E 0.80–1.20; otherwise E = 1). Calibrated log₂ abundance was −Cq_adj × log₂(1 + E); normalized expression Y subtracted the mean calibrated log₂ abundance of TBP, HPRT1 and RPLP0, all required to be measurable. Reference stability was assessed using geNorm [44] and a NormFinder-style variance decomposition [45]. Same-format KC–normal comparisons used matched vehicle and TGF-β1 conditions at 72 h; acute canonical-target and feedback-gene responses were assessed within KC donors. Pre-FN1 and pre-COL1A1 assays measured precursor transcripts; their induced increases and response retention used the same blockade-matched vehicle comparisons as mature transcripts (Table S4). Derived quantities are defined in Table 1. Reaction inputs, calibration equations and assay criteria are detailed in the Supplementary Methods.

### Western blot

GelMA interference with protein assays was addressed by loading 100 ng DNA equivalent per lane, with a six-donor 2D β1 T72 lysate pool on every membrane. Eighty-four independent lysates were distributed over 13 membranes, and antibodies are listed in Table S6. Background-corrected signals were normalized to the same-membrane pool for each target, giving abundance per DNA equivalent. Phosphorylated SMAD2 was normalized to total SMAD2, LOX precursor and mature bands were quantified separately, and the two TPM1 bands were summed. Band-integration parameters are provided in the Supplementary Methods.

### Secreted protein, DNA and retained matrix assays

Supernatant procollagen type I α1 and lumican were measured by ELISA, lactate dehydrogenase by CytoTox 96, and endpoint DNA by PicoGreen with matrix-matched standards. Samples were measured in duplicate. Net secretion was the sample concentration minus the matched cell-free concentration, multiplied by collected volume, retaining negative values; window masses were summed for cumulative output. Per-DNA output was calculated for each construct before averaging its replicates.

Post-withdrawal constructs were decellularized without DNase. Decellularization solution and washes were pooled as D1; 10% of the homogenized matrix supplied DNA (D2) and 90% supplied protein. Collagen and lumican were measured in independent sister constructs by native human type I collagen and lumican ELISAs. Net retained mass was (dilution-corrected sample − matched same-age cell-free extract) × final volume/0.90. Construct DNA was reconstructed as D1 plus D2/0.10 from RPP30 qPCR. Post-withdrawal collagen gain was endpoint mass minus mass at withdrawal for each finite regimen, calculated from sister constructs within donors. The difference between these gains compared delayed with immediate blockade. The same calculation was used for the T72-to-T168 and T72-to-T336 intervals. Standard curves, control samples and calculation formulas are given in the Supplementary Methods.

### Imaging

The concentration screen used Calcein-AM and propidium iodide (PI) confocal z-stacks on days 1, 3, 5, 7, 11 and 14, with one intrafilament field in each of three independent constructs per concentration and day. Architecture imaging comprised 11 volumes from three constructs, and day-7 scanning electron microscopy comprised 16 micrographs from two specimens.

Whole-mount immunofluorescence at T336 used KC donors D01–D03 and normal donors N01–N03. Set A labeled DAPI, collagen I and α-SMA in no-pulse printed, printed vehicle and delayed-blockade constructs; set B labeled DAPI, collagen I and lumican in printed vehicle, cast vehicle and delayed-blockade constructs. Each arm and set contained six constructs, each stained for one set without sectioning. Each set included a no-primary control and a fully stained cell-free gel (Table S6). Confocal settings were fixed within sets with unsaturated printed-vehicle images. Two fields per experimental construct were chosen by position before imaging, yielding 76 volumes from 40 constructs. Acquisition parameters are provided in the Supplementary Methods.

### Image analysis

Filament regions of interest (ROIs) were segmented from transmitted-light texture. Closed-pore dimensions and local filament thickness were measured from these ROIs. Calcein and PI thresholds used the median plus 4 × 1.4826 × MAD. Live-signal volume fraction was positive voxel volume divided by sampled ROI-column volume, and concentration-screen occupancy was positive area divided by ROI area. PI object density was expressed per 100,000 ROI pixels. Fields were combined within constructs by summing numerators and denominators.

Whole-mount images were filtered without pixel-wise background subtraction. The collagen threshold was the control mean plus three standard deviations, and the 561 nm threshold was the larger corresponding value from no-primary and cell-free controls. DAPI nuclei were segmented by Otsu thresholding and watershed. Collagen coverage was positive gel volume divided by segmented gel volume. Nuclear neighborhoods comprised extranuclear gel within 15 μm of the nearest nuclear surface, and collagen and α-SMA fractions used neighborhood volume as denominator. The α-SMA-positive gate was the no-primary-control 95th percentile (0.4654). Radial coverage used distance bins from the nearest nuclear surface. Connected collagen components spanning neighborhoods intersected at least two 15 μm neighborhoods, and lumican overlap was expressed as a fraction of collagen-positive volume. Two fields were pooled per construct, and delayed/vehicle ratios were formed within donors on the log₂ scale with sets A and B weighted equally. Full segmentation, connectivity and radial-bin specifications are given in the Supplementary Methods.

### RNA sequencing

The baseline batch comprised three KC and three normal donors cultured unstimulated in 2D or printed GelMA (12 libraries). The stimulation batch comprised three KC donors, each contributing all eight format × ligand conditions: 2D or printed GelMA under vehicle, TGF-β1, TGF-β2 or TGF-β3 (24 libraries). HTSeq [46] gene counts were filtered for at least 10 reads in at least three samples, retaining 15,989 genes for stimulation and 16,661 for baseline analysis.

DESeq2 [47] used donor-adjusted models: ∼ donor + group for stimulation and ∼ donor + format + KC-in-3D for baseline; within-format disease contrasts used ∼ disease. Analyses used unshrunken log₂ fold changes, Wald statistics and Benjamini–Hochberg (BH) adjustment per contrast. A likelihood-ratio test assessed heterogeneity of format changes among isoforms. For each donor, the induced increase was normalized abundance after TGF-β minus vehicle in the same format; 3D/2D ratios required a positive 2D increase. Gene-set abundance summed member counts with at least three measurable members; the set-level ratio was the median across donors with a positive 2D increase. MSigDB [48] and curated sets were classified as matrix, contractile or other (Table S1). Preranked fgsea used Wald statistics, with BH adjustment within collections; matrix and contractile classes were compared by Mann–Whitney U test. Model contrasts, set definitions, scoring settings and software versions are provided in the Supplementary Methods.

### Patient-tissue datasets, stromal proteome and program scores

Patient-tissue transcriptomes included GSE151631 (19 KC, 7 control) [49], GSE204791 (8 KC, 7 normal) [50], PRJNA1184491 (6 KC, 8 control), GSE77938 (25 KC, 25 non-KC) [51], GSE303036 (8 KC, 5 control sections) [52], GSE285060 (5 KC, 3 control donors) [53] and GSE241715 (4 KC, 7 healthy) [54]. Each dataset was compared with its own controls, retaining the distinctions among postmortem-donor, living-enucleation, pathological and healthy refractive-surgery controls, as applicable; GSE204791 was reprocessed with limma [55]. Patient signatures used the 200 highest-ranked genes in each direction at BH-adjusted *p* < 0.05 and were tested by fgsea in construct contrasts. TGF-β and AP-1 tissue-direction scores used inverse-variance-pooled tissue effect signs to weight standardized expression. Models adjusted for donor and culture group, with BH correction within test families.

The stromal proteome used SWATH-MS data from cone and non-cone regions of four KC corneas and two normal corneas [56]. Limma compared cone with normal tissue and paired cone with non-cone tissue. Downregulated proteins formed the construct-enrichment signature, and UniProt accessions were mapped to gene symbols. Fig. S8A displays matrix, stromal-identity and contractile proteins across the ten samples.

Matrix, contractile and stromal identity programs were scored with singscore [57] on variance-stabilized construct expression. Matrix and contractile sets comprised 64 and 47 genes, respectively, intersecting a 642-gene TGF-β consensus-induced signature [58–60] with functional annotations, including the core matrisome [61]. Consensus genes were upregulated at BH-adjusted *p* < 0.05 and log₂ fold change > 1 in at least two of three public datasets. The 121-gene stromal identity set combined keratocyte-specific genes shared by at least two of six independent references [53,58,60,62,63]. Full cohort sizes, gene-selection rules, memberships and score definitions are provided in the Supplementary Methods and Tables S1 and S3.

The in vivo repopulation reference was GSE328779: injured and contralateral uninjured eyes of four rabbits after full-thickness freeze injury, at day 7 or day 28 (eight libraries). Injury-induced fibroblasts were compared with uninjured keratocytes using within-animal program-score differences. Fibrosis was assessed in GSE302936, comparing stromal remodeling cells from three alkali-burned corneas at day 14 with keratocytes from three uninjured corneas, using library-level means and Welch intervals. Human and rabbit fibroblast-state and myofibroblast-state signatures came from GSE308374 [58] and GSE287915 [60]. The injury-repair signature was also scored after removing contractile genes. Construct pulse effects and their format difference used the donor-adjusted stimulation model. In GSE337743, serum-free rabbit keratocytes in 3D fibrillar collagen were compared with 2D collagen cultures (six libraries per format, Fig. S9 and Table S3). Cluster selection, species mapping and signature construction are detailed in the Supplementary Methods.

### Genetic analysis

KC genome-wide association statistics came from the European analyses of Hardcastle et al. (2,116 cases, 24,626 controls) [64] and the Million Veteran Program (573 cases, 450,622 controls) [65], and FinnGen release 13 (921 cases, 471,319 controls) [66]. Central corneal thickness and corneal resistance factor (CRF) statistics came from published studies [67,68], and CRF adjusted for thickness was genetically residualized on thickness. Colocalization used coloc.susie [69], retaining the highest shared-causal-variant posterior probability within each trait for each KC credible-set component. Expression QTLs used GTEx v8 [70] through the eQTL Catalogue [71] in cultured fibroblasts (n = 483), unexposed skin (n = 517) and sun-exposed lower-leg skin (n = 602). Trait effects were matched to the same lead KC variant and oriented to its risk allele; QTL annotations were matched by gene, KC source and anchor variant (Table S2). Cohort sizes, accessions, variants, tissues and priors are provided in the Supplementary Methods.

### Coding of published stromal intervention studies

The KC-treatment evaluation analysis used human-coded study-level and endpoint-level tables. Two readers independently coded the available reports and a third adjudicated disagreements. Eligible original studies tested drugs, biologics or growth factors, supplements or antioxidants, cells or exosomes, gene or RNA agents, or engineered stromal materials in a KC-related system. The study table contained 191 screened records, of which 166 were coded eligible, linked to 2,116 endpoint records. Study-level fields recorded the model, treatment class, dosing schedule and authors’ benefit claim. Endpoint-level fields recorded the treated system, readout category, sample compartment, timing relative to dosing, direction of effect and whether the authors called the result beneficial. Supporting quotations and their source locations were retained (Table S4 companion files).

Aggregations used the adjudicated categorical codes and counted each study once for each study-level question. Repeated or continuous courses combined continuous/repeated exposure through the readout with repeated dosing followed by withdrawal and follow-up. The post-withdrawal matrix/contraction analysis required a deposited-matrix or contraction endpoint coded as measured after dosing stopped within those dosing designs. Single- or brief-exposure studies were tabulated separately. The absence of a qualifying post-withdrawal endpoint established that no regimen was evaluated or selected using that measured outcome. Definitions, screened records and the reproducible study-level aggregation are supplied in the Supplementary Methods and the Table S4 companion files.

A separate purposive set of 40 full-text studies comprised 39 corneal stromal studies and one non-corneal comparator. Studies were selected from a PubMed-based candidate pool, preferring those that reported expression or secretion together with a deposited-matrix, contraction or functional readout. Two readers independently coded each study and a third adjudicated disagreements. For every comparison supporting an author conclusion, readers recorded candidate conditions, the supporting readouts and their compartment and normalization, the exposure and withdrawal design, and the authors’ selection statement with its source location. Its pathway-inhibition comparisons were retained separately from the KC-treatment denominator (Table S4).

### Statistical analysis

The donor was the statistical unit. Masses and amounts per μg DNA were calculated per construct, averaged arithmetically within donor × condition × readout and then log₂ transformed. For the KC–normal cell-content decomposition at T336, matched log collagen, log DNA and log collagen/DNA measurements were averaged within donors to preserve their additive identity. Expression replicates were averaged on the linear scale before log₂ transformation. Targeted qPCR gene sets used the mean normalized log₂ abundance across member genes within each donor and condition, corresponding to their geometric-mean abundance. The induced increase was stimulated abundance minus the matched vehicle abundance. A format response ratio divided this increase in printed or cast culture by the increase in 2D. Response retention under blockade divided the induced increase in the blocked arm by that in the unblocked arm, using each arm’s own vehicle. This quantity was distinct from the ratio of fold inductions, which compared stimulated/vehicle fold changes. Matrix-to-contractile comparisons used the gene sets specified in the relevant figure and Table 1, and the three- and six-gene contractile sets were analyzed separately.

The log₂ printed/2D ratio of induced increases was decomposed into a baseline term and a baseline-relative induction term. The baseline term was log₂(vehicle abundance in printed culture/vehicle abundance in 2D). The induction term was log₂[(fold induction in printed culture − 1)/(fold induction in 2D − 1)]. Their sum equaled the total format effect. The same decomposition was applied separately to KC and normal donors; same-format disease contrasts compared baseline abundance, fold induction and stimulated abundance separately. Ratios were reported as donor medians on the linear scale or donor means and intervals on the log₂ scale, as specified, with logarithms restricted to positive ratios. Full equations and gene-level aggregation rules are provided in the Supplementary Methods.

Paired effects used two-sided exact sign-flip tests over all donor sign assignments. Confidence intervals were 95% *t* intervals on donor effects; geometric-mean ratios and their intervals were back-transformed from log₂. KC and normal groups used Welch intervals and exact label permutations. The four-regimen analysis shown in Fig. 7A–C included all six KC donors (D01–D06, Table S4). The three β1 regimens were analyzed separately in Fig. S12. Regimen ties were averaged, ranking agreement used Kendall’s τ-b, and endpoint choice followed the within-donor sign of delayed minus immediate blockade. Endpoint-based regimen choices and their collagen differences were calculated from the same donor-level arm means. In the readout-time comparison, the immediate or delayed finite schedule was chosen within donor by the higher native collagen per construct at 72, 168 or 336 h, and the collagen of the chosen arm was read at 336 h. The outcomes of secretion-selected and retention-selected schedules were also compared with each donor’s vehicle using the joint endpoint of more native collagen and lower ACTA2, CCN2 and TAGLN expression.

The concentration screen used balanced two-way ANOVA with interaction (six concentrations, six days, three constructs), on raw and transformed occupancy and dead-nucleus density. For each metric at each day, the five concentrations other than 5% GelMA were compared with 5% GelMA using Welch tests on the raw scale, with Holm correction within that set of five comparisons (60 comparisons in total). Welch 95% confidence intervals were unadjusted. Within-construct architecture contrasts used exact Wilcoxon tests. Cross-layer donor associations used Spearman or Pearson correlations, exact permutation tests and percentile bootstrap intervals from 20,000 donor resamples. Differential-expression tests were BH-adjusted across genes per contrast and enrichment tests within collections. Analyses used Python 3 and R 4.5.0. Transformations, permutation counts, bootstrap settings and package versions are detailed in the Supplementary Methods.

## Supporting information

Table S4 companion files

## Data and code availability

The gene count matrices, normalized expression and differential-expression results of the RNA sequencing libraries, processed experimental readouts, analysis code, microscopy images (lossless OME-TIFF), and plate-reader, qPCR and Western blot instrument exports are deposited at Zenodo (doi:10.5281/zenodo.23616198). Native qPCR and plate-reader project files are held under restricted access at Zenodo (doi:10.5281/zenodo.23617766) and are released to researchers on request. Raw sequencing reads from patient-derived cells are not publicly available because the ethical approval for the study does not permit their public release; they can be requested from the corresponding author and will be considered by the authors subject to institutional and ethical approval. Source data for all figures and Supplementary Data 1–4 are provided with this paper, and primers, antibodies and reagents, sample inventories, and lane and plate maps are listed in Tables S5–S8. Full protocols are given in the Supplementary Methods.

## Declarations

### CRediT authorship contribution statement

Hanyu Jiang: Conceptualization, Data Curation, Formal analysis, Investigation, Methodology, Software, Validation, Visualization, Writing - Original Draft, Writing - Review & Editing. Chang Zhou: Data Curation, Investigation, Methodology, Writing - Review & Editing. Fei Gao: Data Curation, Investigation, Writing - Review & Editing. Yuan Pang: Resources, Supervision, Writing - Review & Editing. Ying Li: Supervision, Writing - Review & Editing. Ying Jie: Funding acquisition, Supervision, Writing - Review & Editing. Huayu Yang: Supervision, Writing - Review & Editing. Yang Jiang: Data Curation, Investigation, Project administration, Resources, Supervision, Writing - Review & Editing.

### Ethics approval and consent to participate

All procedures were performed in compliance with relevant laws and institutional guidelines and were approved by the Institutional Ethics Committee of the Chinese Academy of Medical Sciences and Peking Union Medical College Hospital (approval number: I-24YSB0340, 6 March 2024). Informed consent was obtained from all donors.

### Declaration of generative AI and AI-assisted technologies in the manuscript preparation process

During the preparation of this work, the authors used Claude (Anthropic) and ChatGPT (OpenAI) in order to improve the readability and language of the manuscript and to assist with writing analysis code. After using this tool/service, the authors reviewed and edited the content as needed and take full responsibility for the content of the published article.

## References

[1] P. Padmanabhan, A. Elsheikh, Keratoconus: A Biomechanical Perspective, Curr Eye Res 48 (2023) 121–129. 10.1080/02713683.2022.2088798.

2. H. Jiang, F. Gao, W. Zhang, Y. Jie, Y. Li, Y. Jiang, Reinterpreting Keratoconus Gene Expression in Light of Inherited Stromal Weakness, bioRxiv (2026), version 2, doi:10.64898/2026.10.03.754609. [Preprint].

[3] N. Papachristoforou, A. Ueno, K. Ledwos, J. Bartuś, A. Nowińska, I. Karska-Basta, A Review of Keratoconus Cross-Linking Treatment Methods, J Clin Med 14 (2025) 1702. 10.3390/jcm14051702.

[4] M. El Zarif, K. A Jawad, J.L. Alió Del Barrio, Z. A Jawad, A. Palazón-Bru, M.P. de Miguel, et al., Corneal stroma cell density evolution in keratoconus corneas following the implantation of adipose mesenchymal stem cells and corneal laminas: an in vivo confocal microscopy study, Invest Ophthalmol Vis Sci 61 (2020) 22. 10.1167/iovs.61.4.22.

[5] M. El Zarif, J.L. Alió, J.L. Alió Del Barrio, K. Abdul Jawad, A. Palazón-Bru, Z. Abdul Jawad, et al., Corneal stromal regeneration therapy for advanced keratoconus: long-term outcomes at 3 years, Cornea 40 (2021) 741–754. 10.1097/ICO.0000000000002646.

[6] M. Rafat, M. Jabbarvand, N. Sharma, M. Xeroudaki, S. Tabe, R. Omrani, et al., Bioengineered corneal tissue for minimally invasive vision restoration in advanced keratoconus in two clinical cohorts, Nat Biotechnol 41 (2023) 70–81. 10.1038/s41587-022-01408-w.

[7] ClinicalTrials.gov, NCT07388069: Study of TN-001 topical eyedrops for keratoconus, https://clinicaltrials.gov/study/NCT07388069 (accessed 25 October 2026).

[8] J.W. Foster, V. Shinde, U.S. Soiberman, G. Sathe, S. Liu, J. Wan, et al., Integrated Stress Response and Decreased ECM in Cultured Stromal Cells From Keratoconus Corneas, Invest Ophthalmol Vis Sci 59 (2018) 2977–2986. 10.1167/iovs.18-24367.

[9] J.C. Erie, S.V. Patel, J.W. McLaren, C.B. Nau, D.O. Hodge, W.M. Bourne, Keratocyte density in keratoconus. A confocal microscopy study, Am J Ophthalmol 134 (2002) 689–695. 10.1016/s0002-9394(02)01698-7.

[10] J.Y. Ku, R.L. Niederer, D.V. Patel, T. Sherwin, C.N. McGhee, Laser scanning in vivo confocal analysis of keratocyte density in keratoconus, Ophthalmology 115 (2008) 845–850. 10.1016/j.ophtha.2007.04.067.

[11] D. Karamichos, R. Zareian, X. Guo, A.E. Hutcheon, J.W. Ruberti, J.D. Zieske, Novel in Vitro Model for Keratoconus Disease, J Funct Biomater 3 (2012) 760–775. 10.3390/jfb3040760.

[12] U.S. Soiberman, A.E.M. Shehata, M.X. Lu, T. Young, Y.J. Daoud, S. Chakravarti, et al., Small Molecule Modulation of the Integrated Stress Response Governs the Keratoconic Phenotype In Vitro, Invest Ophthalmol Vis Sci 60 (2019) 3422–3431. 10.1167/iovs.19-27151.

[13] S.E. Wilson, L.P. Sampaio, T.M. Shiju, G.S.L. Hilgert, R.C. de Oliveira, Corneal Opacity: Cell Biological Determinants of the Transition From Transparency to Transient Haze to Scarring Fibrosis, and Resolution, After Injury, Invest Ophthalmol Vis Sci 63 (2022) 22. 10.1167/iovs.63.1.22.

[14] K. Borner, Z. Yu, C. Xing, W.M. Petroll, Single cell RNA-seq characterization of non-fibrotic stromal wound repopulation in the rabbit, Exp Eye Res 264 (2026) 110816. 10.1016/j.exer.2025.110816.

[15] D.P. Maruri, M. Miron-Mendoza, P.B. Kivanany, J.M. Hack, D.W. Schmidtke, W.M. Petroll, et al., ECM Stiffness Controls the Activation and Contractility of Corneal Keratocytes in Response to TGF-β1, Biophys J 119 (2020) 1865–1877. 10.1016/j.bpj.2020.08.040.

[16] D. Karamichos, A.E. Hutcheon, J.D. Zieske, Transforming growth factor-β3 regulates assembly of a non-fibrotic matrix in a 3D corneal model, J Tissue Eng Regen Med 5 (2011) e228–e238. 10.1002/term.429.

[17] D. Karamichos, A.E. Hutcheon, J.D. Zieske, Reversal of fibrosis by TGF-β3 in a 3D in vitro model, Exp Eye Res 124 (2014) 31–36. 10.1016/j.exer.2014.04.020.

18. A. Choi, S.E. Nicholas, J. Kim, D. Karamichos, Exploring Bioprinted Keratoconus Corneal Stroma: A Comparative Study, Invest Ophthalmol Vis Sci 66 (2025) 1029 (ARVO Annual Meeting abstract).

[19] R. Hadvina, A. Estes, Y. Liu, Animal Models for the Study of Keratoconus, Cells 12 (2023) 2681. 10.3390/cells12232681.

20. A.L. Pereira-Souza, R. Ambrósio, F. Bandeira, M.Q. Salomão, A. Souza Lima, S.E. Wilson, Topical losartan for treating corneal fibrosis (haze): first clinical experience, J Refract Surg 38 (2022) 741–746. 10.3928/1081597X-20221018-02.

[21] J.L. Domene-Hickman, L. Haro-Morlett, A. Lichtinger, A. Hernandez-Solis, A. Ramirez Miranda, A. Navas, et al., Topical losartan for established corneal fibrosis with machine learning-based predictors, Ther Adv Ophthalmol 17 (2025) 25158414251378123. 10.1177/25158414251378123.

[22] B. Burgos-Blasco, G. Moloney, Topical losartan for corneal fibrosis: a case series with densitometry analysis, Cornea 45 (2026) 168–173. 10.1097/ICO.0000000000003783.

[23] F. Barzaghi, M. Kronschläger, S. Amir-Asgari, O. Findl, Topical losartan to prevent stromal scarring during post-cross-linking infection in pediatric keratoconus: a case report, Am J Ophthalmol Case Rep 42 (2026) 102549. 10.1016/j.ajoc.2026.102549.

[24] W. Wei, B. Ma, H.Y. Li, Y. Jia, K. Lv, G. Wang, et al., Biphasic effects of selective inhibition of transforming growth factor beta1 activin receptor-like kinase on LPS-induced lung injury, Shock 33 (2010) 218–224. 10.1097/shk.0b013e3181aef736.

[25] P. Yang, L. Zhu, S. Wang, J. Gong, J.N. Selvaraj, L. Ye, et al., Engineered model of heart tissue repair for exploring fibrotic processes and therapeutic interventions, Nat Commun 15 (2024) 7996. 10.1038/s41467-024-52221-9.

[26] D.C. Zanin-Silva, N.J.T. van Kooten, T.I. Papadimitriou, D.N. Dorst, B. Walgreen, E.L. Vitters, et al., Monocytes strongly induce (myo)fibroblast contraction in a 3D skin model to understand inflammation-fibrosis crosstalk, Sci Rep 16 (2026) 28481. 10.1038/s41598-026-62996-0.

[27] X. Guo, S. Sriram, J.A. Tran, A.E.K. Hutcheon, J.D. Zieske, Inhibition of Human Corneal Myofibroblast Formation, Invest Ophthalmol Vis Sci 59 (2018) 3511–3520. 10.1167/iovs.18-24239.

[28] Y. Moradi, P. Shrestha, S. Mabry, P. Sharma, K.S. Multani, K.M. Riaz, et al., In Vitro Assessment of Nintedanib in Keratoconus Corneal Stromal Microenvironment, Biomolecules 16 (2026) 1137. 10.3390/biom16081137.

[29] R.L. Prentice, Surrogate endpoints in clinical trials: definition and operational criteria, Stat Med 8 (1989) 431–440. 10.1002/sim.4780080407.

[30] T.R. Fleming, D.L. DeMets, Surrogate end points in clinical trials: are we being misled?, Ann Intern Med 125 (1996) 605–613. 10.7326/0003-4819-125-7-199610010-00011.

[31] C. Kilic Bektas, V. Hasirci, Cell loaded 3D bioprinted GelMA hydrogels for corneal stroma engineering, Biomater Sci 8 (2020) 438–449. 10.1039/c9bm01236b.

[32] G.J. Inman, F.J. Nicolás, J.F. Callahan, J.D. Harling, L.M. Gaster, A.D. Reith, et al., SB-431542 is a potent and specific inhibitor of transforming growth factor-beta superfamily type I activin receptor-like kinase (ALK) receptors ALK4, ALK5, and ALK7, Mol Pharmacol 62 (2002) 65–74. 10.1124/mol.62.1.65.

[33] C.A. Greene, C.R. Green, M.E. Dickinson, V. Johnson, T. Sherwin, Keratocytes are induced to produce collagen type II: A new strategy for in vivo corneal matrix regeneration, Exp Cell Res 347 (2016) 241–249. 10.1016/j.yexcr.2016.08.010. PMID: 27539660.

[34] J. Wu, J. Wang, L. Wang, Y. Huang, Topical retinoic acid induces corneal strengthening by upregulating transglutaminase 2 in murine cornea, Exp Eye Res 214 (2022) 108850. 10.1016/j.exer.2021.108850. PMID: 34861212.

[35] E.F. Nelson, C.W. Huang, J.M. Ewel, A.A. Chang, C. Yuan, Halofuginone down-regulates Smad3 expression and inhibits the TGFβ-induced expression of fibrotic markers in human corneal fibroblasts, Mol Vis 18 (2012) 479–487.

[36] S. Sriram, P. Robinson, L. Pi, A.S. Lewin, G. Schultz, Triple combination of siRNAs targeting TGFβ1, TGFβR2, and CTGF enhances reduction of collagen I and smooth muscle actin in corneal fibroblasts, Invest Ophthalmol Vis Sci 54 (2013) 8214–8223.

[37] T. Shuijing, W. Jiakang, Y. Tingting, J. Ma, Biomechanical and inflammatory pathways underlying the genetic architecture of keratoconus: a genomic SEM study, Exp Eye Res 263 (2026) 110770. 10.1016/j.exer.2025.110770.

[38] X. Jiang, T. Boutin, V. Vitart, Colocalization of corneal resistance factor GWAS loci with GTEx e/sQTLs highlights plausible candidate causal genes for keratoconus postnatal corneal stroma weakening, Front Genet 14 (2023) 1171217. 10.3389/fgene.2023.1171217.

[39] S. Chakravarti, T. Magnuson, J.H. Lass, K.J. Jepsen, C. LaMantia, H. Carroll, Lumican regulates collagen fibril assembly: skin fragility and corneal opacity in the absence of lumican, J Cell Biol 141 (1998) 1277–1286. 10.1083/jcb.141.5.1277.

[40] ClinicalTrials.gov, NCT07449000: Losartan for corneal fibrosis, https://clinicaltrials.gov/study/NCT07449000 (accessed 25 October 2026).

[41] A. Untergasser, I. Cutcutache, T. Koressaar, J. Ye, B.C. Faircloth, M. Remm, et al., Primer3--new capabilities and interfaces, Nucleic Acids Res 40 (2012) e115. 10.1093/nar/gks596.

[42] J. Morales, S. Pujar, J.E. Loveland, A. Astashyn, R. Bennett, A. Berry, et al., A joint NCBI and EMBL-EBI transcript set for clinical genomics and research, Nature 604 (2022) 310–315. 10.1038/s41586-022-04558-8.

[43] J. Ye, G. Coulouris, I. Zaretskaya, I. Cutcutache, S. Rozen, T.L. Madden, Primer-BLAST: a tool to design target-specific primers for polymerase chain reaction, BMC Bioinformatics 13 (2012) 134. 10.1186/1471-2105-13-134.

44. J. Vandesompele, K. De Preter, F. Pattyn, B. Poppe, N. Van Roy, A. De Paepe, et al., Accurate normalization of real-time quantitative RT-PCR data by geometric averaging of multiple internal control genes, Genome Biol 3 (2002) RESEARCH0034. 10.1186/gb-2002-3-7-research0034.

[45] C.L. Andersen, J.L. Jensen, T.F. Ørntoft, Normalization of real-time quantitative reverse transcription-PCR data: a model-based variance estimation approach to identify genes suited for normalization, applied to bladder and colon cancer data sets, Cancer Res 64 (2004) 5245–5250. 10.1158/0008-5472.CAN-04-0496.

[46] S. Anders, P.T. Pyl, W. Huber, HTSeq--a Python framework to work with high-throughput sequencing data, Bioinformatics 31 (2015) 166–169. 10.1093/bioinformatics/btu638.

[47] M.I. Love, W. Huber, S. Anders, Moderated estimation of fold change and dispersion for RNA-seq data with DESeq2, Genome Biol 15 (2014) 550. 10.1186/s13059-014-0550-8.

[48] A. Liberzon, C. Birger, H. Thorvaldsdóttir, M. Ghandi, J.P. Mesirov, P. Tamayo, The Molecular Signatures Database (MSigDB) hallmark gene set collection, Cell Syst 1 (2015) 417–425. 10.1016/j.cels.2015.12.004.

[49] V. Shinde, N. Hu, A. Mahale, G. Maiti, Y. Daoud, C.G. Eberhart, et al., RNA sequencing of corneas from two keratoconus patient groups identifies potential biomarkers and decreased NRF2-antioxidant responses, Sci Rep 10 (2020) 9907. 10.1038/s41598-020-66735-x.

[50] T. Stachon, M. Nastaranpour, B. Seitz, E. Meese, L. Latta, S. Taneri, et al., Altered Regulation of mRNA and miRNA Expression in Epithelial and Stromal Tissue of Keratoconus Corneas, Invest Ophthalmol Vis Sci 63 (2022) 7. 10.1167/iovs.63.8.7.

[51] M. Kabza, J.A. Karolak, M. Rydzanicz, M.W. Szcześniak, D.M. Nowak, B. Ginter-Matuszewska, et al., Collagen synthesis disruption and downregulation of core elements of TGF-β, Hippo, and Wnt pathways in keratoconus corneas, Eur J Hum Genet 25 (2017) 582–590. 10.1038/ejhg.2017.4.

[52] A. Wysocka, K. Jaskiewicz-Rajewicz, J. Wozniak, M. Rydzanicz, R. Ploski, M. Udziela, et al., “Eye-Conic” Spatial Transcriptomics Reveals the Layer-Specific Molecular Alterations in Corneas of Patients With Keratoconus, Invest Ophthalmol Vis Sci 67 (2026) 7. 10.1167/iovs.67.4.7.

[53] S. Luo, J. Li, Y. Yang, Y. Jiang, Y. Jie, W. Ge, Spatial transcriptomics and single-cell RNA-sequencing revealed dendritic cell-mediated inflammation in keratoconus, Ocul Surf 36 (2025) 134–150. 10.1016/j.jtos.2025.01.008.

[54] T. Lapp, P. Kammrath Betancor, G. Schlunck, C. Auw-Hädrich, P. Maier, C. Lange, et al., Transcriptional profiling specifies the pathogen-specific human host response to infectious keratitis, Front Cell Infect Microbiol 13 (2023) 1285676. 10.3389/fcimb.2023.1285676.

[55] M.E. Ritchie, B. Phipson, D. Wu, Y. Hu, C.W. Law, W. Shi, et al., limma powers differential expression analyses for RNA-sequencing and microarray studies, Nucleic Acids Res 43 (2015) e47. 10.1093/nar/gkv007.

[56] G.H. Yam, M. Fuest, L. Zhou, Y.C. Liu, L. Deng, A.S. Chan, et al., Differential epithelial and stromal protein profiles in cone and non-cone regions of keratoconus corneas, Sci Rep 9 (2019) 2965. 10.1038/s41598-019-39182-6.

[57] M. Foroutan, D.D. Bhuva, R. Lyu, K. Horan, J. Cursons, M.J. Davis, Single sample scoring of molecular phenotypes, BMC Bioinformatics 19 (2018) 404. 10.1186/s12859-018-2435-4.

[58] M. Kim, K. Choi, D. Križaj, J. Kim, Regulation of corneal stromal cell behavior by modulating curvature using a hydraulically-controlled organ chip array, Nat Commun 16 (2025) 9944. 10.1038/s41467-025-64889-8.

[59] R. Kumar, R. Tripathi, N.R. Sinha, R.R. Mohan, RNA-Seq Analysis Unraveling Novel Genes and Pathways Influencing Corneal Wound Healing, Invest Ophthalmol Vis Sci 65 (2024) 13. 10.1167/iovs.65.11.13.

[60] K. Poole, K.S. Iyer, D.W. Schmidtke, W.M. Petroll, V.D. Varner, Corneal Keratocytes, Fibroblasts, and Myofibroblasts Exhibit Distinct Transcriptional Profiles In Vitro, Invest Ophthalmol Vis Sci 66 (2025) 28. 10.1167/iovs.66.3.28.

[61] A. Naba, K.R. Clauser, S. Hoersch, H. Liu, S.A. Carr, R.O. Hynes, The matrisome: in silico definition and in vivo characterization by proteomics of normal and tumor extracellular matrices, Mol Cell Proteomics 11 (2012) M111.014647. 10.1074/mcp.M111.014647.

[62] J. Collin, R. Queen, D. Zerti, S. Bojic, B. Dorgau, N. Moyse, et al., A single cell atlas of human cornea that defines its development, limbal progenitor cells and their interactions with the immune cells, Ocul Surf 21 (2021) 279–298. 10.1016/j.jtos.2021.03.010.

[63] P. Català, N. Groen, J.A. Dehnen, E. Soares, A.J.H. van Velthoven, R.M.M.A. Nuijts, et al., Single cell transcriptomics reveals the heterogeneity of the human cornea to identify novel markers of the limbus and stroma, Sci Rep 11 (2021) 21727. 10.1038/s41598-021-01015-w.

[64] A.J. Hardcastle, P. Liskova, Y. Bykhovskaya, B.J. McComish, A.E. Davidson, C.F. Inglehearn, et al., A multi-ethnic genome-wide association study implicates collagen matrix integrity and cell differentiation pathways in keratoconus, Commun Biol 4 (2021) 266. 10.1038/s42003-021-01784-0.

[65] A. Verma, J.E. Huffman, A. Rodriguez, M. Conery, M. Liu, Y.L. Ho, et al., Diversity and scale: Genetic architecture of 2068 traits in the VA Million Veteran Program, Science 385 (2024) eadj1182. 10.1126/science.adj1182.

[66] M.I. Kurki, J. Karjalainen, P. Palta, T.P. Sipilä, K. Kristiansson, K.M. Donner, et al., FinnGen provides genetic insights from a well-phenotyped isolated population, Nature 613 (2023) 508–518. 10.1038/s41586-022-05473-8.

[67] H. Choquet, R.B. Melles, J. Yin, T.J. Hoffmann, K.K. Thai, M.N. Kvale, et al., A multiethnic genome-wide analysis of 44,039 individuals identifies 41 new loci associated with central corneal thickness, Commun Biol 3 (2020) 301. 10.1038/s42003-020-1037-7.

[68] X. Jiang, N. Dellepiane, E. Pairo-Castineira, T. Boutin, Y. Kumar, W.A. Bickmore, et al., Fine-mapping and cell-specific enrichment at corneal resistance factor loci prioritize candidate causal regulatory variants, Commun Biol 3 (2020) 762. 10.1038/s42003-020-01497-w.

[69] C. Wallace, A more accurate method for colocalisation analysis allowing for multiple causal variants, PLoS Genet 17 (2021) e1009440. 10.1371/journal.pgen.1009440.

[70] GTEx Consortium, The GTEx Consortium atlas of genetic regulatory effects across human tissues, Science 369 (2020) 1318–1330. 10.1126/science.aaz1776.

[71] N. Kerimov, J.D. Hayhurst, K. Peikova, J.R. Manning, P. Walter, L. Kolberg, et al., A compendium of uniformly processed human gene expression and splicing quantitative trait loci, Nat Genet 53 (2021) 1290–1299. 10.1038/s41588-021-00924-w.

