## Supplementary Information for "Material context and TGF-β blockade timing determine matrix retained after withdrawal in patient-derived GelMA-bioprinted keratoconus stroma"

### Supplementary Methods

#### Donors and primary corneal stromal cells

Keratoconus (KC) stromal tissue was obtained from host corneal buttons removed at keratoplasty, and normal stromal tissue from refractive-surgery corneas without apparent disease. Six KC donors (D01–D06) and six normal donors (N01–N06) were used; each donor yielded one independent isolate. Tissue was minced and digested in DMEM/F-12 containing 1 mg/mL type I collagenase at 37 °C for 6–8 h. Cells were cultured in DMEM/F-12 with 10% fetal bovine serum, 100 U/mL penicillin, 100 μg/mL streptomycin and 2 mM L-glutamine at 37 °C in 5% $CO_2$, with medium changed every 2–3 days, and were passaged at 80%–90% confluence with 0.25% trypsin–EDTA. All printing, protein assays and sequencing used passage 3. Fabrication batch B1 comprised D02, D05 and D06, batch B2 comprised D01, D03 and D04, and the normal donors were split in the same way (N01–N03 in B1, N04–N06 in B2). The baseline transcriptomes used KC isolates D02, D05 and D06 and normal isolates N01, N02 and N05; the TGF-β-stimulated transcriptomes used KC isolates D03, D04 and D05. The study received institutional ethics approval; donors gave informed consent and procedures followed the Declaration of Helsinki.

#### GelMA–LAP ink and rheology

Cell suspension in PBS, 15% (w/v) gelatin methacryloyl (GelMA) and 2.5% (w/v) lithium phenyl-2,4,6-trimethylbenzoylphosphinate (LAP; both from SUNP Biotech) were combined at 17:10:3 (v/v) to give 5.0% GelMA, 0.25% LAP and $5 \times 10^6$ cells/mL. The ink was loaded into a sterile 3 mL syringe, cooled at 4 °C for 10 min and rewarmed at 20 °C for 10 min before use. Ascorbic acid 2-glucoside (AA2G) was never added to the ink. For the concentration screen, GelMA was prepared at 4%, 5%, 6%, 8%, 10% and 12% with cells of one normal isolate, and 5% was used for all later work.

Rheology was conducted on an Anton Paar rheometer with a PP25 parallel plate (gap 0.199 mm) at 1.5 Hz and 1% strain under TruStrain control. Cell-free precursor and cell-laden ink of matched composition were cooled from 37 to 4 °C at 5 °C/min (400 s, 100 points per curve; measured rate 4.96–5.00 °C/min). The gel point was the temperature of the last crossing from $\ln(G'/G'') < 0$ to $> 0$, linearly interpolated between adjacent points. The print thermal sequence was reproduced in a time sweep of 10 min at 4 °C and 10 min at 20 °C. Compression used molded cylinders (height 3.50 mm, diameter 5.50 mm, area 23.758 mm²), photocrosslinked as below; the modulus was the slope of a linear regression of stress on strain over 10%–20% strain after averaging in 0.05 mm displacement bins, and compressive strength was read from printed constructs.

The recorded G′–G″ cooling crossover temperatures were 17.7 ± 1.8 °C for cell-free precursor and 15.6 ± 1.6 °C for cell-laden ink (Fig. S1A). Conditioning and rewarming recordings are shown in Fig. S1B.

**Bioprinting and photocrosslinking**

Constructs were printed on a BIOMARKER 3D cell bioprinting system through a 21G conical nozzle into 24-well plates as six-layer grids of nominal 8 × 8 × 1.2 mm at a head speed of 8 mm/s and a nominal volumetric rate of 2 $mm^3$/s. The syringe was held at 4 °C for 10 min and at 20 °C for 10 min, the print head at 20 °C and the platform at 15 °C. After deposition of the whole construct, one exposure to 405 nm light at 15 mW/$cm^2$ for 15 s (225 mJ/$cm^2$ at the incident surface) crosslinked the gel, and constructs were transferred to complete medium.

**Culture formats**

Three formats were compared. Two-dimensional (2D) cultures were cells on tissue-culture plastic. Cast constructs were ink of the same final composition, thermal history, pre-cooled collection tube, dwell time and wide-bore tip transfer as the paired printed construct, delivered from a needleless Luer outlet into a Press-to-Seal silicone frame (P24744, Invitrogen; diameter 9 mm, depth 1.0 mm; 64 µL and nominally $3.2 \times 10^5$ cells per frame) on the 15 °C platform and crosslinked together with the paired print. A nozzle-pass recast reference was extruded through the 21G nozzle at print parameters into the same frames. Frames were trimmed to fit the well bottom, sterilized, dried and applied adhesive side down; medium was added along the wall until the gel was covered, and the frame was lifted vertically. The fabrication clock M0 was the end of crosslinking of a three-dimensional (3D) construct or the end of the paired sham treatment of a 2D culture, and the stimulation clock T0 was the start of dosing on day 7 after fabrication (Md7). Samples collected at M30 min were stopped 25–35 min after M0 by lysis or snap freezing in liquid nitrogen.

The functional batch (184 constructs on eight 24-well plates; 12 donors with 14 constructs each, and 16 cell-free GelMA blanks) was cultured in 1.00 mL DMEM/F-12 with 10% fetal bovine serum without serum withdrawal or ascorbate, with medium changed on days 3 and 5. The post-withdrawal batch (536 constructs, including 44 cell-free) was cultured in 1.00 mL medium with 10% fetal bovine serum from a single lot, with 0.5 mM AA2G from the first medium addition after the first crosslinking, replenished at every change, and bridging arms without AA2G (PIn, PDn) were run in parallel. The imaging batch used the post-withdrawal batch medium and AA2G, with the endpoint at T336.

**TGF-β pulses and receptor blockade**

Recombinant human TGF-β1, TGF-β2 and TGF-β3 (Proteintech) were reconstituted in PBS with 0.1% BSA at 10 µg/mL and stored at −20 °C. On Md7, cultures received 10 ng/mL of an isoform for 4 h (0–4 h), or matched vehicle, with dosing staggered so that every sample received a 4 h exposure. At 4 h cells were washed once with warm PBS and returned to their original complete medium; the 4 h sample was collected before the wash. SB431542 (10 µM) was given immediately after washout (4–72 h) or after a 20 h delay (24–72 h), all arms contained 0.1% (v/v) DMSO from 4 h, delayed arms received matched DMSO before 24 h, and medium with drug or solvent was renewed at 4, 24 and 48 h. In the functional batch, U was no pulse with vehicle, V a β1 pulse with vehicle, I a β1 pulse with blockade from 4 h, and D a β1 pulse with blockade from 24 h, in printed and cast constructs. Supernatant from each construct was collected in three windows (W1 4–24 h, W2 24–48 h, W3 48–72 h), removing all medium at each collection, and the construct was homogenized in 0.50 mL at 72 h for DNA.

In the retention batch, printed (P) and cast (C) constructs received vehicle after the pulse (PV, CV), blockade from 4 h to 72 h and then no drug (PIs), blockade from 24 h to 72 h and then no drug (PDs) or blockade from 24 h continued to day 14 (PDc). After 72 h, constructs were washed twice gently and medium with AA2G was exchanged at T120, T168, T216, T264 and T312, with SB431542 renewed only for PDc; the supernatant of the last 24 h (T312–T336) was collected at T336. In the post-withdrawal batch, supernatant of the 48–72 h window (W3) was collected from the PV, PIs and PDs arms. At T72, T168 and T336 constructs were collected for collagen, lumican, RNA and DNA. For the finite regimens, T168 is 4 days and T336 is 11 days after withdrawal at 72 h; PDc remained on treatment at T336. The imaging batch comprised no pulse and no blockade (PU), PV, PDs and cast vehicle (CV), each collected at T336. In the supplementary quantitative PCR (qPCR) experiment, six KC donors in 2D and printed culture received vehicle or β1 followed by vehicle, immediate or delayed blockade (72 T72 samples, of which the 24 vehicle samples were shared with the main arms); further arms covered β3 pulses in 2D and printed culture and a T24 memory arm. For the signaling readout, constructs were pre-incubated with SB431542 or solvent for 1 h before 10 ng/mL β1 for about 45 min.

#### RNA extraction and quantitative PCR

Three-dimensional constructs were lysed whole after a brief PBS wash and snap freezing, disrupted mechanically in the frozen state in lysis reagent, cleared by centrifugation and purified on silica columns with on-column DNase I; 2D samples were processed with the same chemistry. RNA was eluted in 14 µL. RNA concentration, A260/280 and A260/230 were measured by NanoDrop. Reverse transcription used 200 ng RNA in 20 µL (post-withdrawal batch) or an input scaled to the number of assays (supplementary batch; 5 ng RNA equivalent per reaction well, two technical wells per assay). Quantitative PCR used SYBR Green with 400 nM primers and 2 µL of 1:5-diluted cDNA in 20 µL on a Bio-Rad CFX96 (95 °C for 30 s; 40 cycles of 95 °C for 5 s and 60 °C for 30 s; melt 55–95 °C in 0.5 °C steps). Primers (Table S5) were designed with Primer3 on MANE Select transcripts (GRCh38; Tm 58–62 °C, GC 40%–60%, amplicons 70–180 bp, worst-case dimer $\Delta G > -9$ kcal/mol) and checked by Primer-BLAST against human RefSeq mRNAs (≤2 mismatches, non-target products ≤1000 bp); high-GC targets used 70–90 bp amplicons.

The supplementary batch comprised 227 96-well sample plates (276 samples: 252 KC, 24 normal; 51 assays; 14,472 sample wells and 897 control wells), each with a single gene, same-donor paired conditions together, duplicate wells, one no-template control, one pooled-cDNA calibrator well and at least one no-RT well, and 13 seven-point standard-curve plates carrying three or four assays each with triplicate wells per level. The post-withdrawal batch comprised 36 expression plates (three plates per assay for the target genes COL1A1, COL5A1, FN1, LUM, DCN, CHST6, ACTA2, CCN2 and TAGLN and the reference genes TBP, HPRT1 and RPLP0; four complete donors × 11 conditions × duplicates per plate) and 24 RPP30 genomic DNA plates. Cq values were read from baseline-corrected quantification results and melt peaks from the instrument export (CFX Manager 3.1). A sample was detected at Cq ≤ 35 with a single melt peak within 1.0 °C of the assay's reference peak; for JUN, JUNB, pre-FN1, pre-COL1A1 and pre-ACTA2, the earliest finite no-RT Cq on the plate was used as a floor and the sample had to be more than 5 cycles earlier. Duplicate wells were averaged when both passed (a difference above 0.5 cycles was flagged), a single passing well was kept, and a sample with no passing well was left missing. Plate offsets were the plate's calibrator Cq minus the mean calibrator Cq of that gene. Amplification efficiency E came from seven-level standard curves of $10^7$–$10^1$ copies in triplicate (at least three consecutive levels with all Cq ≤ 35, $R^2 \geq 0.98$, E between 0.80 and 1.20;

otherwise E = 1; E 0.845–1.121 for 44 assays). Calibrated $\log_2$ abundance was −Cq_adj × $\log_2(1 + E)$. Reference-normalized expression, Y, was calculated by subtracting the mean calibrated $\log_2$ abundance of TBP, HPRT1 and RPLP0, all of which had to be measurable; reference stability was assessed with geNorm M and a NormFinder-style variance decomposition. Induced increases, format effects, response retention under blockade and matrix-to-contractile response ratios are defined in the Statistical analysis section.

The normal-donor qPCR panel comprised FN1, COL1A1, COL5A1, ACTA2, CCN2, TAGLN, TPM1, LOX and CCN1 at 72 h, in 2D and printed cultures with vehicle or a TGF-β1 pulse. Within each format, disease-group comparisons used vehicle abundance, pulse/vehicle fold induction and abundance after the pulse. The matrix program comprised FN1, COL1A1 and COL5A1; the three-gene contractile program comprised ACTA2, CCN2 and TAGLN, and the expanded contractile program also included TPM1, LOX and CCN1. Program abundance was calculated as the geometric mean of the corresponding normalized gene abundances within each donor and condition. Acute canonical-target, feedback-gene and immediate-early responses were analyzed within the KC donors by pairing pulse and vehicle samples from the same donor, format and time; $\log_2$ fold induction was summarized by its donor median and the number of donors with induction above zero. These targets and early time points were measured in KC donors only.

Pre-FN1 and pre-COL1A1 were assayed as precursor transcripts alongside mature matrix transcripts. Their fold induction and response retention used the same donor- and blockade-matched vehicle comparisons. Summary values for these assays are provided in Table S4, Ongoing_transcription.

#### Western blot

Constructs and 2D cultures were washed once in ice-cold PBS, snap frozen (the 64 μL gel was kept whole) and pulverized under liquid nitrogen with a pre-cooled pestle. Lysis used RIPA (50 mM Tris-HCl pH 7.4, 150 mM NaCl, 1% NP-40, 0.5% sodium deoxycholate, 0.1% SDS) with 1× protease inhibitor, 1× PhosSTOP, 10 mM NaF and 1 mM $Na_3VO_4$, 120 μL per construct on ice for 10 min followed by 30 min of sonication in a 4 °C water bath and 14,000 × g for 15 min at 4 °C. DNA of the homogenate was measured by Quant-iT PicoGreen (P7589; Invitrogen) in a BioTek Synergy H1 reader (Ex 480/Em 520 nm) after at least 100-fold dilution in TE with matrix-matched standards. Because GelMA interferes with protein assays, each lane was loaded with lysate containing 100 ng DNA (diluted 3:1 with 4× Laemmli buffer containing reducing agent, 95 °C for 5 min), and a calibration pool of six-donor 2D β1 T72 lysate (100 ng DNA) was on every membrane. Lane 1 of each gel carried a prestained protein marker (10–250 kDa, 3–5 μL), and the kDa positions in the figures are marker bands of the same membrane. Hand-cast 12% (membranes 1–6) or 10% (7, 8, 11, 13) gels or 4%–15% precast gels (9, 10, 12) were run in a Mini-PROTEAN Tetra system at 80 V for about 30 min and 120 V thereafter, transferred to 0.45 μm PVDF in wet format at 100 V for 90 min at 4 °C and blocked in 5% non-fat milk/TBST (5% BSA/TBST for p-SMAD2) for 1 h. Membranes were incubated with primary antibodies (Table S6) overnight at 4 °C and HRP-conjugated secondary antibodies (1:5000) for 1 h at room temperature, and signal was developed with ECL and imaged on a Tanon 5200 system with serial exposures saved as TIFF. Membranes were stripped and reprobed in the order α-SMA → GAPDH with the lower segment for TAGLN, CCN2 → TPM1 → GAPDH, p-SMAD2 → total SMAD2/3, and the upper segment for FN1 with the lower for LOX. Eighty-four independent lysates were distributed over 13 membranes.

Band intensity was integrated from 1100 × 1344 8-bit images as the sum of (background − pixel) over a box of 0.88 × the median lane spacing in width and 28 pixels in height (double bands shared the height by band spacing, minimum 10 pixels), with background from the median of flanking regions above and below the lane box excluding other bands. For each membrane and target, the longest exposure with no saturated pixels in any loaded lane box (≤ 2 gray level in ≥ 0.5% of pixels defined saturation) was used; for TAGLN on two membranes, TPM1 on two membranes and LOX on two membranes no exposure met this criterion, and the shortest exposure was used. The primary scale was the $\log_2$ of the signal relative to the calibration pool on the same membrane for the same target, which gives abundance per DNA equivalent at 100 ng DNA per lane; p-SMAD2 was normalized to total SMAD2, LOX precursor (50 kDa) and mature (32 kDa) bands were integrated separately, and the two TPM1 bands were summed.

#### Secreted protein and DNA assays

Procollagen type I α1 and lumican in supernatants were measured by DuoSet ELISA (DY6220-05 and DY2846-05; R&D Systems; standards 31.2–2000 and 125–8000 pg/mL) read at 450 nm on a BioTek Synergy H1; lactate dehydrogenase was measured with CytoTox 96 (G1780; Promega; 490 nm). Each plate carried an eight-level standard curve with zero in duplicate, a technical pool and known-concentration controls, and samples were assayed in duplicate. Standards were fitted with a four-parameter logistic function $y = d + (a - d)/(1 + (x/c)^b)$ with equal weights, concentrations were back-calculated per well, duplicate wells were averaged and the mean was multiplied by the dilution. A sample below the lowest non-zero standard was below the limit of detection and one above the top standard was above range; the latest within-range reread was used where a higher dilution had been run. The net window output of a construct was (sample concentration − matched cell-free concentration of the same batch, format, arm and window) × collected volume (1.00 mL), with negative values retained; the masses from the three windows were added. Total DNA of the end-point homogenate (0.50 mL) was quantified by PicoGreen after at least 100-fold dilution in TE with matrix-matched standards; blank-subtracted signal was fitted by linear regression with intercept, and DNA mass per construct was concentration × dilution × homogenate volume. Secretion per µg DNA was the net mass of the construct divided by its DNA, and replicate constructs were averaged after the ratio.

#### Retained collagen and lumican

After the last supernatant collection, post-withdrawal batch constructs were washed three times with PBS and decellularized in 0.5% Triton X-100 with 20 mM $NH_4OH$ for 3 min at room temperature, without DNase, and washed again (decellularization solution and washes pooled, D1). The remaining matrix was homogenized cold, 10% of the suspension was taken for DNA (D2) and 90% for protein, with collagen and lumican taken from independent sister constructs. For collagen, matrix was extracted in 0.05 M acetic acid with 0.1 mg/mL pepsin at 4 °C for 24 h, neutralized in ELISA-compatible buffer and brought to 250 µL. For lumican, matrix was extracted in 4 M guanidine-HCl with 50 mM Tris pH 7.4 and protease inhibitors at 4 °C overnight and exchanged into DuoSet diluent on desalting columns. Native human type I collagen was measured by the Chondrex 6021 ELISA (0.08–5 µg/mL; OPD, 490 nm) and lumican by DY2846-05; extracts were mixed 1:1 with diluent (twofold dilution), samples were assayed in duplicate with a seven-level standard curve plus zero, and above-range samples were rerun at higher dilution. Net retained mass per construct was (dilution-corrected sample − matched same-age cell-free extract) × final volume/0.90, where 0.90 is the share of the homogenate sent to the protein route. DNA of D1, D2 and whole constructs was purified on columns and quantified by qPCR of the single-copy RPP30 locus against a seven-point four-fold standard curve (33,000–8.06 ng/reaction from genomic DNA at 0.0033 ng/copy; $Cq = a + b \log_{10}$ ng/reaction); duplicate wells were averaged, tube DNA was ng/reaction × elution volume/template volume (50/2 µL), and construct DNA was the D1 total plus D2/0.10. Post-withdrawal accumulation was the retained native collagen at T336 minus that at T72 within each blockade schedule. The difference in accumulation between schedules subtracted the immediate-blockade gain from the delayed-blockade gain. The time points were measured in sister constructs compared within donors. The first post-withdrawal interval used T168 instead of T336.

#### Live/dead imaging, scanning electron microscopy and architecture imaging

For the GelMA concentration screen, constructs were stained on days 1, 3, 5, 7, 11 and 14 with 1 µM Calcein-AM and 2 µM propidium iodide (PI) for 20 min at room temperature in the dark, washed in PBS and imaged as confocal z-stacks, with one field inside a filament in each of three independent constructs per concentration and day. Live-signal occupancy was the Calcein-positive area divided by the area of the filament region of interest (ROI), and PI object density was the red nuclear load per 100,000 ROI pixels; grid voids were excluded from the ROI.

Printed-construct architecture was imaged as Olympus OIR volumes of 640 × 640 pixels (11 volumes from three independent constructs: seven 4× fields and four 10× fields) with a 4× objective (UPLXAPO4X, NA 0.16, working distance 13.0 mm; pixel size 4.97 µm) or a 10× objective (UPLXAPO10X, NA 0.4, working distance 3.1 mm; pixel size 1.99 µm), at a z step of 18.43 µm (21–25 planes at 4× and 17–21 planes at 10×). Channel 1 recorded Calcein (excitation 500 nm, emission 540 nm), channel 2 PI (570/620 nm) and channel 3 transmitted light; excitation lines were 488 and 561 nm and the pinhole was 127 µm.

For scanning electron microscopy, day-7 constructs were washed in PBS, fixed in 2.5% glutaraldehyde in 0.1 M phosphate buffer at 4 °C overnight and postfixed in 1% osmium tetroxide for 1.5 h, dehydrated through graded ethanol and isoamyl acetate, critical-point dried and sputter-coated with gold. Sixteen micrographs (two specimens, four fields each, imaged at 2.00k and 30.0k with the same field in each pair) were acquired at 3.00 kV with a Hitachi field-emission microscope.

#### Whole-mount immunofluorescence

The imaging batch used KC donors D01–D03 and normal donors N01–N03 at T336. Set A (DAPI, collagen I, α-SMA) comprised PU, PV and PDs constructs and set B (DAPI, collagen I, lumican) PV, CV and PDs constructs, six constructs per arm and set; each construct was stained for one set and was not sectioned. PV and PDs constructs of sets A and B were sister constructs. Each set contained one no-primary-antibody control (secondary antibodies and DAPI only) and one fully stained cell-free gel. Constructs were fixed in 4% paraformaldehyde (diluted from 16% formaldehyde, 28908, Thermo Fisher Scientific) for 30 min at room temperature, washed three times in PBS, permeabilized in 0.3% Triton X-100/PBS for 30 min and blocked in 5% normal goat serum (005-000-121, Jackson ImmunoResearch) with 1% BSA for 2 h. The collagen I primary antibody (ab317789, recombinant rabbit–mouse chimeric antibody with a mouse IgG1 constant region, Abcam; 1 µg/mL) was applied overnight at 4 °C, followed by three 30 min washes in PBS-T, an Alexa Fluor 488 secondary antibody (115-545-205, Fcγ1-specific, rabbit-adsorbed, Jackson ImmunoResearch; 1:400) for 2 h at room temperature and three further washes. Directly labeled rabbit primary antibodies were then applied overnight at 4 °C: α-SMA–Alexa Fluor 555 (ab202509, Abcam; 5 μg/mL) in set A and lumican–Alexa Fluor 555 (ab319968, Abcam; 1:250) in set B. After three 30 min washes, nuclei were stained with DAPI (1 μg/mL) for 30 min, and constructs were stored in PBS in glass-bottom dishes without mounting medium.

Images were acquired on a Nikon C2plus confocal microscope with three PMTs and sequential channel scanning (405 nm excitation with 450/50 emission for DAPI, 488 nm with 525/50 for collagen I and 561 nm with 595/50 for α-SMA or lumican), a Plan Fluor 20×/NA 0.5 objective, 1024 × 1024 pixels at 0.313 μm per pixel (320 × 320 μm) and 16 planes at 2 μm, starting 15 μm below the construct surface, at 12 bit depth. Acquisition settings were fixed within a set so that the brightest PV field was unsaturated. Two non-overlapping fields were chosen per construct by position before imaging, at least 1 mm from the construct edge with the construct center on a filament, giving 76 volumes from 40 constructs (36 experimental constructs with two fields each and four controls with one field each).

**Image analysis**

Filament ROIs of the low-magnification volumes were derived from the channel 3 z-mean texture (Gaussian high-pass σ = 15 μm, RMS smoothing σ = 40 μm, Li threshold within the central 90% of both axes, closing radius 60 μm, opening radius 10 μm, removal of components below 10,000 $\mu m^2$ and of holes below 100,000 $\mu m^2$, and a 32-pixel crop on each edge). Closed pores were measured by connected-component area (equivalent diameter $2\sqrt{(A/\pi)}$) and by x and y chords through the centroid, and local filament thickness was the diameter of the largest inscribed circle on the ROI medial axis, excluding radii limited by the crop frame. Calcein and PI were thresholded per file on the maximum-intensity projection at the median plus 4 × 1.4826 × MAD (MAD = 0 taken as 1); Calcein objects had to cover at least 20 pixels per plane and PI objects 8–220 pixels with a short bounding-box side of at least 2 pixels and an aspect ratio of at most 3.5. The live-signal volume fraction was the Calcein-positive voxel volume divided by the sampled column volume of the filament ROI across all z planes, and PI planes were joined by 26-connectivity into three-dimensional PI objects, reported as object density and positive volume fraction. Fields were pooled within a construct by summing numerators and denominators.

In the seven low-magnification fields from three constructs quantified in Fig. 1F (representative image, Fig. 1B), filament thickness was measured at 7,159 positions with a pooled median of 521 μm, and the mean of the three construct medians was 532 μm (95% CI 370–694 μm); the mean of the three construct median pore equivalent diameters was 1.46 mm (95% CI 1.28–1.65 mm).

Whole-mount volumes were analyzed on the channel identified by its excitation wavelength, without pixel-wise background subtraction, after Gaussian filtering ($\sigma xy$ = 0.5 μm, $\sigma z$ = 1 μm) and removal of positive blocks below 1 $\mu m^3$ (26-connectivity). The collagen threshold was the mean plus three population standard deviations of all filtered voxels of the no-primary controls of the set, and the 561 nm threshold was the larger of the two analogous candidates from the no-primary control and the cell-free gel. Nuclei were segmented by an Otsu threshold on the pooled DAPI histogram of each set with control voxels included, hole filling and a watershed seeded by h-maxima (h = 2 μm) of the Euclidean distance transform in physical units, keeping small and border nuclei. The gel mask was the sum of the 488 and 561 nm images, each clipped at its non-zero median, low-pass filtered (Gaussian $\sigma xy$ = 5 μm, $\sigma z$ = 2 μm), thresholded at 10% of the non-zero low-frequency median and closed and opened with a radius of about 1 μm per plane. Collagen coverage was the collagen-positive voxels divided by gel-mask voxels. The 15 μm neighborhood of a nucleus was the extranuclear gel within 15 μm of its surface by Euclidean distance, with each voxel assigned uniquely to the nearest nucleus; the collagen fraction of a neighborhood was its collagen-positive voxels divided by its voxels, and the α-SMA fraction s was the α-SMA-positive voxels divided by its voxels. The α-SMA gate NP95 was the 95th percentile (linear interpolation) of s over neighborhoods of the no-primary control ($s > 0.4654$ was α-SMA positive). Radial bins were (0, 5], (5, 10], (10, 15], (15, 25], (25, 50], (50, 100] and > 100 μm from the nearest nuclear surface, with the denominator the extranuclear gel voxels of the bin and the numerator the collagen-positive voxels of the same domain. Collagen components were labeled in the collagen-positive gel by 26-connectivity, with volume as voxel count × voxel volume, and a component spanned neighborhoods when it intersected at least two nuclear 15 μm neighborhoods. For lumican, coverage was the lumican-positive gel volume divided by gel volume, and f(LUM|COL) the overlap of collagen and lumican divided by the collagen-positive volume. Two fields of a construct were combined by summing numerators and denominators, nucleus counts were added, and the PDs/PV ratio was formed at the donor level on the $\log_2$ scale, with the A and B sets combined with equal weights.

Scanning electron microscopy (SEM) micrographs were cropped to x = 0–2560 and y = 0–1790 pixels to remove the annotation band, thresholded at the Otsu value to define the dark phase (8-connectivity) without smoothing, and used to measure local thickness of the bright phase as the diameter of the largest inscribed circle, averaged over covered pixels.

**RNA sequencing**

Two sets of libraries were sequenced. The baseline set comprised three KC and three normal donors cultured unstimulated in 2D or as printed constructs (12 libraries). The stimulation set comprised three KC donors in 2D or printed GelMA under vehicle, TGF-β1, TGF-β2 or TGF-β3 (24 libraries, with each donor contributing all eight format × ligand conditions). Total RNA was sequenced on an Illumina NovaSeq 6000 platform (paired-end, 150 bp), and gene-level counts were generated with HTSeq. Genes with at least 10 reads in at least three samples were kept (15,989 genes in the stimulation model and 16,661 in the paired-format baseline model).

Differential expression used DESeq2 (R 4.5.0, DESeq2 1.48.2). The stimulation model was ~ donor + group, with group the eight format × ligand combinations and 2D vehicle as the reference; normalized counts supplied linear abundance, and variance-stabilized values supplied principal-component analyses (blind = FALSE) and program scores (blind = TRUE). The baseline model was ~ donor + format + KC-in-3D, with KC-versus-normal contrasts within each format from ~ disease. Results used independent filtering, unshrunken $\log_2$ fold changes and Wald statistics, and Benjamini–Hochberg (BH) adjustment per contrast. The effect common to the three isoforms was the equal-weight mean of the TGF-β1, TGF-β2 and TGF-β3 contrasts. Heterogeneity of the 2D-to-3D change among isoforms was tested by likelihood ratio between the full model (~ donor + group) and a reduced model with a single 3D-by-stimulation term (~ donor + culture + ligand + 3D-stimulated).

For each gene and donor, the TGF-β-induced increase was the normalized count under TGF-β minus that under vehicle in the same format, and the 3D/2D ratio was the 3D increase divided by the 2D increase, defined when the 2D increase was positive. A gene set's abundance was the sum of the normalized counts of its members (at least three measurable members), and the set-level ratio was computed per donor and summarized as the median over donors with a positive 2D increase. Gene sets came from MSigDB through msigdbr (Hallmark, KEGG, Reactome, canonical pathways, Gene Ontology biological process, cellular component and molecular function, and cell-type signatures; 8,025 sets) plus seven curated sets (the three matrix genes, seven structural matrix genes, the six contractile genes, a myofibroblast contractile core, collagen biosynthesis and modification, fibrillar matrix assembly and YAP/TAZ mechano-effectors), and were classed as matrix, contractile or other (Table S1). Preranked gene set enrichment used fgsea on the Wald statistic (set size 10–500, seed 1) with BH adjustment within each collection, and over-representation analysis used clusterProfiler on genes with BH-adjusted $p < 0.05$, with all tested genes of the contrast as the universe. The matrix and contractile classes were compared with the Mann–Whitney U test.

**Patient-tissue datasets and tissue-direction scores**

Patient-tissue transcriptomes were taken from public datasets: GSE151631 (central cornea, 19 KC and 7 control), GSE204791 (isolated stromal layer, 8 KC and 7 normal), PRJNA1184491 (stromal subset, 6 KC and 8 control), GSE77938 (whole cornea, 25 KC and 25 non-KC), GSE303036 (spatial transcriptomics, stromal layer of 8 KC and 5 control sections), GSE285060 (10x Visium spatial transcriptomics, keratocyte-cluster spots pooled to pseudobulk per donor, 5 KC and 3 control) and GSE241715 (formalin-fixed tissue, 4 KC and 7 healthy). Control types were kept separate: postmortem donor corneas (GSE151631, GSE204791 stroma, PRJNA1184491), pathological corneas (GSE77938), and corneas from living patients undergoing enucleation for choroidal melanoma (GSE241715). Gene-level effects ($\log_2$ fold change, standard error and statistic) were taken from the comparison of each dataset against its own controls, with the genes tested in that dataset as background. The GSE204791 array was reprocessed from the Agilent green channel (normexp background correction with offset 16, quantile normalization, probes detected above background in at least 25% of samples, averaging by gene symbol; limma with empirical Bayes, ~ disease).

Patient gene sets were built per dataset from genes with BH-adjusted $p < 0.05$, ordered by statistic, taking the 200 highest in each direction. Their enrichment in the ranked KC-versus-normal and 2D-versus-3D contrasts of the constructs was analyzed using fgsea (set size 10–5,000, eps $1 \times 10^{-50}$, seed 20261002) with the adjusted $p$ value reported as q. The tissue-direction score for TGF-β pathway and AP-1/immediate-early genes pooled the $\log_2$ fold change of each gene across control classes by inverse variance and used its sign as weight; the project's $\log_2$(normalized count + 1) was $z$-scored per gene across samples, and the score was the equal-weight mean over direction-weighted genes (donor-adjusted models ~ donor + group for stimulation and ~ donor + format + KC-in-3D for baseline; BH within each family of tests).

**Stromal proteome and RNA program scores**

The stromal proteome of keratoconic corneas came from the SWATH-MS data of Yam et al. (Supporting Data File S1, stromal proteins; 799 protein rows), which cover the cone and non-cone regions of four KC corneas and two normal corneas (ten samples; 100 μg protein per sample, normalized to total ion current). Abundances were $\log_2$ transformed and rows with finite values in at least eight of the ten samples were kept (all 799). Cone-versus-normal effects were fitted with limma and empirical Bayes moderation in the six samples concerned (four KC cone regions and two normal corneas) and the cone-versus-non-cone effect in the eight KC samples with donor and region terms. Proteins with nominal $p < 0.05$ and a negative $t$ statistic formed the downregulated set (31 proteins for cone versus normal), mapped to gene symbols through their UniProt accessions (org.Hs.eg.db); their enrichment in the Wald-statistic-ranked contrasts of the constructs was analyzed using fgsea (minimum set size 10). Fig. S8A shows the centered $\log_2$ abundance of 26 matrix, stromal-identity and contractile proteins across all ten samples.

Program scores were computed with singscore (single-set, centered, known direction) on DESeq2 variance-stabilized expression (blind = TRUE; one row per gene symbol, the one with the highest mean) for the 12 baseline and 24 stimulation-batch libraries, and a set with fewer than 10 scorable genes returned no score. The matrix gene set (64 genes) is the intersection of a TGF-β consensus-induced set (642 genes) with matrix annotations (Gene Ontology extracellular matrix organization, collagen fibril organization, collagen-containing extracellular matrix and collagen metabolic process, and the NABA core matrisome; 413 genes); the contractile gene set (47 genes) is the intersection of the same TGF-β set with contractile annotations (Gene Ontology actomyosin structure organization, muscle and smooth muscle contraction, contractile fiber, stress fiber, actin filament bundle and its assembly; 658 genes). The TGF-β consensus-induced set comprised genes upregulated (BH-adjusted $p < 0.05$, $\log_2$ fold change > 1) in at least two of three public datasets: GSE260476 (TGF-β versus control), GSE308374 and rabbit GSE287915 (myofibroblasts versus keratocytes). The stromal identity gene set (121 genes) comprised keratocyte-specific genes shared by at least two of six independent references: GSE121922 (keratocyte mean at least 7.5 TPM, epithelial and endothelial means below 7.5 TPM and a keratocyte-to-maximum ratio of at least 4), GSE308374 and GSE287915 (keratocyte versus fibroblast and myofibroblast, BH-adjusted $p < 0.05$ and $\log_2$ fold change > 1, top 150 by mean Wald statistic), and the keratocyte clusters of the single-cell datasets GSE155683 and GSE186433 and the keratocyte spot cluster of the spatial dataset GSE285060 (highest-AUC markers, 100–150 per cluster). Gene sets were read from MSigDB 2026.1.

The in vivo reference used the single-cell dataset GSE328779 (rabbit cornea after full-thickness freeze injury; injured and uninjured contralateral eyes of four rabbits at day 7 or day 28; eight libraries; 17 clusters). The injury-induced fibroblast cluster (cluster 8; 2,615 of 2,635 cells from injured eyes) was compared with the two keratocyte clusters of highest keratocyte-marker score (clusters 1 and 3). For each animal, the mean normalized expression (expm1 of the normalized data, cells weighted equally; clusters 1 and 3 pooled by cell number) of cluster 8 cells of the injured eye and of cluster 1 and 3 cells of the uninjured eye was scored with the sets above on the complete measured gene universe, rabbit genes being matched to human symbols of identical name (59 of 64 matrix, 43 of 47 contractile and 100 of 121 identity genes), and the within-animal difference was averaged over the four animals with a t-distribution 95% confidence interval (3 degrees of freedom); a second reference, clusters 1 and 3 of the injured eye, gave the same sign of the mean difference for every set (Table S3). The fibroblast-state and myofibroblast-state signatures were the 150 genes per state higher (BH-adjusted $p < 0.05$, $\log_2$ fold change > 1) than in both other states of GSE308374 (human) or GSE287915 (rabbit), ranked by mean Wald statistic; the human reference differences are myofibroblast minus keratocyte scores averaged over the four curvature conditions of GSE308374. The injury-repair program (715 genes) comprised the genes higher in cluster 8 than in clusters 1 and 3 (Wilcoxon test, BH-adjusted $q < 0.01$, $\log_2$ fold change > 0.5, expressed in at least 25% of cells in one group) and was also scored without the contractile genes (ACTA2, TAGLN, POSTN, CNN1, MYH11, CCN2 and the members of the contractile set). In the stimulation batch, the pulse effects in 2D and in printed constructs and their difference were contrasts of the donor-adjusted model (~ donor + group; 24 libraries, 14 degrees of freedom). In GSE302936 (rabbit cornea 14 days after alkali burn; three injured and three uninjured corneas, six libraries, 38,710 cells; author clusters), the stromal clusters labeled ECM-remodeling cells and activated keratocytes (clusters 2 and 7 of the injured corneas; 7,523 cells) were compared with the keratocyte cluster of the uninjured corneas (cluster 0; 9,028 cells); the clusters were fixed before scoring and the cluster labeled persistent myofibroblasts (cluster 8) was not included. Each library's mean normalized expression (expm1 of the log-normalized layer, equal weight per cell) was scored with the sets above (rabbit genes matched to human symbols of identical name: 60 of 64 matrix, 46 of 47 contractile, 103 of 121 identity, 137 of 150 myofibroblast-state genes), the difference between the injured libraries and the mean of the three uninjured libraries is the library-level difference, and the difference of group means is given with an unpaired Welch t-distribution 95% confidence interval (the GEO record does not pair injured and uninjured corneas). Marker-positive fractions are the fractions of cells with at least one UMI count of the gene, pooled over libraries. In GSE337743 (rabbit keratocytes, serum-free; 15 libraries), 3D fibrillar collagen (50,000 and 100,000 cells per matrix; six libraries) was compared with 2D collagen-coated and 2D fibrillar culture (six libraries) in a linear model on group (10 degrees of freedom).

#### Genetic analysis

KC association statistics came from three genome-wide association studies: Hardcastle et al. (European, GCST90013442; 2,116 cases and 24,626 controls), the Million Veteran Program European analysis (GCST90477661) with 573 cases and 450,622 controls, and FinnGen release 13 (921 and 471,319). Summary statistics for central corneal thickness (CCT, GCST90000654) and corneal resistance factor (CRF, GCST90100568) came from published genome-wide association studies; CRF adjusted for CCT was the CRF signal genetically residualized on CCT. Colocalization of the KC and trait signals used coloc.susie: for each locus, gene, KC source, trait and KC credible-set component, the table gives the posterior probability of a shared causal variant (PP4) for the component with the highest value within that trait, without taking a maximum across traits. Expression quantitative trait loci were from GTEx v8 as processed in the eQTL Catalogue (cultured fibroblasts, QTD000216, n = 483; unexposed skin, QTD000311, n = 517; sun-exposed lower leg skin, QTD000316, n = 602; prior probabilities $p1 = p2 = 1 \times 10^{-4}$ and $p12 = 5 \times 10^{-6}$). Effects for the lead KC variant (rs3118518-A, rs1536483-T and rs1536482-A at COL5A1 for the three sources, and rs2075556-G at COL1A1) were taken for the same variant in each trait dataset and oriented to the KC risk allele using the original effect and other alleles. QTL annotations were matched on gene, KC source and anchor variant, and opposite tissue directions were kept as reported (Table S2).

#### Coding of published treatment evaluation

Two human-coded literature sets addressed different questions. The KC-treatment set supplied the study-level denominator for evaluation practice. The purposive stromal set supplied detailed comparison-level records of how particular readouts supported an author's intervention choice. The sets were analyzed separately.

For the KC-treatment set, two readers independently coded reports and a third reader adjudicated disagreements. Original studies were eligible when they tested a drug, biologic or growth factor, supplement or antioxidant, cell or exosome treatment, gene or RNA agent, or engineered stromal material with a KC or ectasia treatment aim. Systems included KC-patient clinical studies, KC patient-derived cells or tissue, KC animal or model systems, and other corneal systems with a KC treatment aim. The final study table contained 191 records. Eligibility codes identified 166 eligible studies and

25 excluded records: 16 with no treatment tested, three studying a procedure or device alone, one without a KC link, one that was not original research, and four coded unclear. The endpoint table contained 2,128 records; joining by PMID to the eligible study set retained 2,116. Full text was coded for 157 of the 166 eligible studies and title and abstract for nine, and source-coverage notes and coding decisions are preserved in the supplied tables.

The study table recorded PMID, bibliographic identifiers, eligibility, KC-related system, agent class and name, dosing schedule and whether benefit was claimed, together with quotations and notes. The endpoint table recorded the treated system, readout category, sample compartment, timing relative to dosing, treated-versus-comparator direction, whether the author called the outcome beneficial, and the supporting result quotation. The readout vocabulary distinguished transcripts, cellular or lysate proteins, secreted proteins, deposited matrix, contraction or traction, biomechanics, imaging, clinical geometry, visual function, cell behaviour, safety, and enzyme or metabolite measures. These categories kept a change in transcript or whole-construct lysate protein distinct from a measurement of deposited matrix.

All study counts used unique PMIDs. Repeated-course studies were the union of the schedule codes continuous_or_repeated_through_readout and repeated_then_stopped_then_followed (31 and 14 studies). A study contributed to the matrix/contraction count if at least one of its endpoints had readout_category equal to deposited_matrix or contraction_or_traction. It contributed to the post-withdrawal count only when such an endpoint was also coded after_dosing_stopped. This deterministic join yielded four repeated-course studies with matrix or contraction endpoints and zero with either endpoint after withdrawal. Because no repeated-course study measured this post-withdrawal outcome, none could have evaluated or selected its regimen using it.

Single- or brief-exposure studies remained in the full 166-study denominator and were tabulated separately from repeated courses. Their later matrix or contraction outcomes did not enter the repeated-course withdrawal count. The complete schedule distribution was 102 single/brief exposures, 31 continuous/repeated courses through the readout, 14 repeated courses followed by stopping and follow-up, 13 not stated, five other and one blank. The blank schedule belonged to PMID 10703124, whose source note reported abstract-based coding, and its adjudicated eligible classification and blank fields were retained.

The four repeated-course studies with deposited-matrix or contraction endpoints were also checked against their supplied full texts. Greene et al. (PMID 27539660) measured deposited collagen during TGF-β3/dexamethasone treatment and followed collagen transcripts after withdrawal; the latter was an RNA endpoint. Soiberman et al. (PMID 31390655) measured cell-number-normalized hydroxyproline during ISRIB exposure. Wu et al. (PMID 34861212) measured collagen fibril density at the end of repeated topical retinoid treatment. Moradi et al. (PMID 42650804) described a contraction pilot without a stated measurement time and used whole-construct lysates for protein measurements after four weeks of continuous treatment; its immediate wash-and-stain viability assay was distinct from matrix or contraction after a post-withdrawal culture period. Original endpoint timing codes were retained.

The study and endpoint tables of the KC-treatment evaluation are supplied as KC_treatment_studies.csv (191 screened records) and KC_treatment_endpoints.csv (2,128 endpoint records), companion files to Supplementary Table S4 (Supplementary_Table_4_companion_files.zip).

KC_treatment_summary.csv gives the deterministic aggregates, KC_treatment_source_records.csv identifies the source records, and KC_treatment_codebook.md supplies the complete category definitions. Data-record numbers count parsed CSV records after the header, and quoted multiline fields do not create additional records. The Table S4 workbook itself holds the experimental sheets and the coding sheets of the purposive set of 40 full-text studies described below.

The purposive set comprised 40 full-text studies, with 39 corneal stromal studies and one non-corneal comparator, selected from a PubMed-based candidate pool with preference for studies reporting expression or secretion together with deposited matrix, contraction or a functional outcome. Two readers independently extracted candidate interventions, comparisons, readout compartments and normalizations, treatment and withdrawal histories, and verbatim author selection statements; a third reader adjudicated disagreements. Eleven corneal pathway-inhibition or myofibroblast-reversal studies contributed 72 comparisons. None included continued culture after inhibitor withdrawal before outcome assessment: 65 comparisons used continuous exposure, six comparisons involved washing before readout only, and one did not state exposure timing. Two studies counted lower type I collagen as an antifibrotic effect. These comparison-level observations are given in the Table S4 coding sheets Study_coding_sources, Study_coding_comparisons and Study_coding_summary. They are not combined numerically with the 166-study KC-treatment denominator.

**Statistical analysis**

The statistical unit was the donor. Construct replicates were summarized within each donor × condition × readout before any comparison: masses and amounts per µg DNA were computed per construct, averaged arithmetically within donor, and converted to $\log_2$; expression was averaged on the linear scale (2^Y, with Y the reference-normalized $\log_2$ expression) before conversion to $\log_2$. For the KC–normal cell-content decomposition at T336, matched log collagen, log DNA and log collagen/DNA measurements were averaged within donors to preserve their additive identity. For qPCR programs, gene expression was averaged on the sample $\log_2$ scale before conversion to linear abundance. The TGF-β-induced increase was abundance after the pulse minus matched vehicle abundance in the same format and blockade regimen. The format effect was the ratio of the induced increase in 3D to that in 2D, defined when the 2D increase was positive. Response retention under blockade was the induced increase with blockade divided by the unblocked induced increase, defined when the unblocked increase was positive. Relative preservation of matrix and contractile responses was compared by the ratio of their response-retention fractions. Fold induction was pulse abundance divided by matched vehicle abundance; the $\log_2$ ratio of fold induction with versus without blockade was calculated separately and compared between the gene programs. Ratios of fold inductions and retention of induced increases were therefore distinct quantities.

The baseline/induction decomposition separated the format effect on resting abundance from the format effect on the increase relative to that baseline. On the $\log_2$ scale, the baseline term was the printed/2D vehicle-abundance ratio. The baseline-relative induction term was $\log_2$[(printed fold induction − 1)/(2D fold induction − 1)]. Their sum equaled the $\log_2$ printed/2D ratio of the pulse-induced increase. The format interaction in fold induction was also calculated as $\log_2$(printed fold induction) minus $\log_2$(2D fold induction). Format-response ratios were summarized by donor medians on the linear scale and by donor means and intervals on the $\log_2$ scale when positive. KC and normal donors were compared for each component and for vehicle abundance, fold induction and abundance after the pulse within the same format. These disease-group comparisons used two-sided exact label permutations and Welch 95% confidence intervals on the $\log_2$ scale; their reported p values were unadjusted.

Paired effects were tested with a two-sided exact sign-flip test on the donor-level differences, enumerating all $2^n$ sign assignments (64 for six donors, 32 for five and 4,096 for twelve), with *p* the share of assignments at least as extreme as the observed statistic. The smallest attainable *p* is therefore 0.0625 for five donors, 0.031 for six donors and 0.0005 for twelve. Intervals were 95% *t* intervals on donor-level effects (mean ± $t_{0.975,n-1}$ × SD/√n), and ratios were summarized as geometric means with intervals back-transformed from the $\log_2$ scale. KC and normal donors were compared as independent groups by the difference of group means with a Welch *t* interval and an exact label permutation preserving group sizes (924 assignments for six versus six). The four-regimen analysis and the three-β1-regimen analysis both used all six KC donors (D01–D06) and are reported separately.

Regimen ranks averaged ties; the three targets were the largest matrix-gene induced increase, the smallest contractile-gene induced increase and the largest matrix-to-contractile ratio. Agreement between format rankings used Kendall's τ-b, and a donor's endpoint choice was delayed or immediate blockade according to the sign of the difference between the two arms. The spatial PDs/PV contrast combined imaging sets A and B with equal weights on the $\log_2$ scale at the donor level. For the concentration screen, a balanced two-way analysis of variance with interaction (six concentrations, six days, three constructs; 72 residual degrees of freedom) was applied to live-signal occupancy (raw and logit-transformed after clipping to 0.0001–0.9999) and to dead-nucleus density (raw and log(1 + x)); the concentration-versus-5% comparisons used Welch tests on the raw scale. Within each metric and day, Holm correction was applied to the five comparisons with 5% GelMA (60 comparisons in total across both metrics and all six days). Welch 95% confidence intervals were unadjusted. Within-construct filament architecture contrasts used two-sided exact Wilcoxon tests; SEM micrographs were summarized per specimen over four fields. Cross-layer associations were Spearman or Pearson correlations of donor-level effects with exact two-sided permutation *p* values (720 permutations for six donors) and percentile bootstrap intervals (20,000 donor resamples).

Differential-expression *p* values were adjusted by the BH procedure over the genes tested in each contrast, gene set enrichment *p* values within each collection, and score-test *p* values within each family of dataset-by-contrast tests. Analyses were run in Python 3 and R 4.5.0 with DESeq2 1.48.2, fgsea 1.34.0, msigdbr 7.5.1, limma 3.64.0, edgeR 4.6.2, clusterProfiler 4.16.0, org.Hs.eg.db 3.21.0, singscore 1.28.0 and metafor 4.8.0.

#### Data and code availability

The gene count matrices, normalized expression and differential-expression results of the RNA sequencing libraries, processed experimental readouts, analysis code, microscopy images (lossless OME-TIFF), and plate-reader, qPCR and Western blot instrument exports are deposited at Zenodo (doi:10.5281/zenodo.23616198). Native qPCR and plate-reader project files are held under restricted access at Zenodo (doi:10.5281/zenodo.23617766) and are released to researchers on request. Raw sequencing reads from patient-derived cells are not publicly available because the ethical approval for the study does not permit their public release; they can be requested from the corresponding author and will be considered by the authors subject to institutional and ethical approval. Source data for all figures and Supplementary Data 1–4 are provided with this paper.

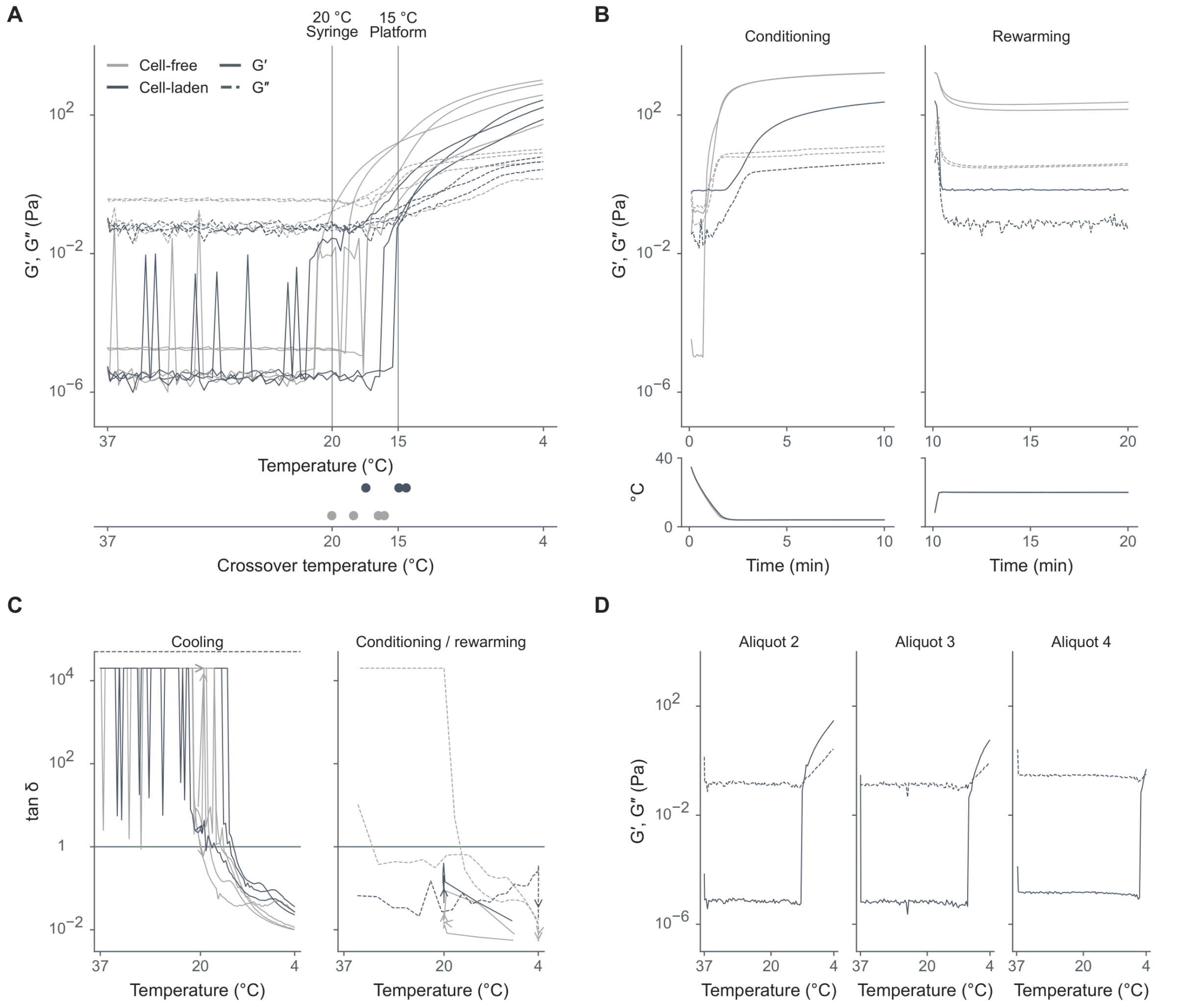


**Fig. S1. Rheology, compression and scanning electron microscopy of gelatin methacryloyl ink and constructs.** (A) Storage modulus G′ and loss modulus G″ (Pa) during cooling from 37 to 4 °C in seven independent traces, comprising four cell-free and three cell-laden recordings. Aligned dots mark each trace's last G′ = G″ crossing. Vertical references mark the syringe and platform temperatures of 20 and 15 °C. (B) G′ and G″ during separately recorded conditioning and rewarming stages, with measured temperature beneath the same time axis, in three traces (two cell-free and one cell-laden). (C) Loss tangent, tan δ = G″/G′, during the seven cooling and three conditioning/rewarming traces. Horizontal references mark tan δ = 1, and arrows indicate acquisition direction. Dashed curves denote conditioning. The fine dashed line marks the cooling-panel display limit of 50,000, with no curves truncated. (D) G′ and G″ during cooling for three later aliquots (2–4) of the same cell-laden preparation. Colors in (B)–(D) and line styles in (B) and (D) follow the key in (A). Modulus axes in (A), (B) and (D) and tan δ axes in (C) are logarithmic. (E) Force–displacement records (N versus mm) and compressive moduli (kPa) of six cast cylinders, three cell-free and three cell-laden. Solid, dashed and dash-dotted lines identify specimens 1, 2 and 3, respectively. Horizontal bars show individual moduli from stress–strain fits over strain 0.10–0.20. Negative force and displacement denote compression. (F) Failure force (N) and area-normalized compressive strength (MPa) of 12 printed specimens, three each for Control and transforming growth factor-β1, β2 and β3 (TGF-β1, TGF-β2 and TGF-β3). Each bar represents one specimen, without error bars. The same physical specimen occupies the same column in both outcome plots. (G) Low-magnification scanning electron microscopy (SEM) images of day-7 constructs at 2,000×. Rows identify two numbered samples, and columns identify four fields per sample (eight fields). Scale bars, 20 µm. (H) High-magnification SEM images at 30,000×, arranged in the same sample and field order as in (G), with four fields per sample (eight fields). Scale bars, 1 µm. Images in (G) and (H) use a linear 0–255 grayscale. The Sample 1, Field 0 images in (G) and (H) show the same fields as the corresponding images in Fig. 1C.

Fig. S1 (continued; 2 / 2)

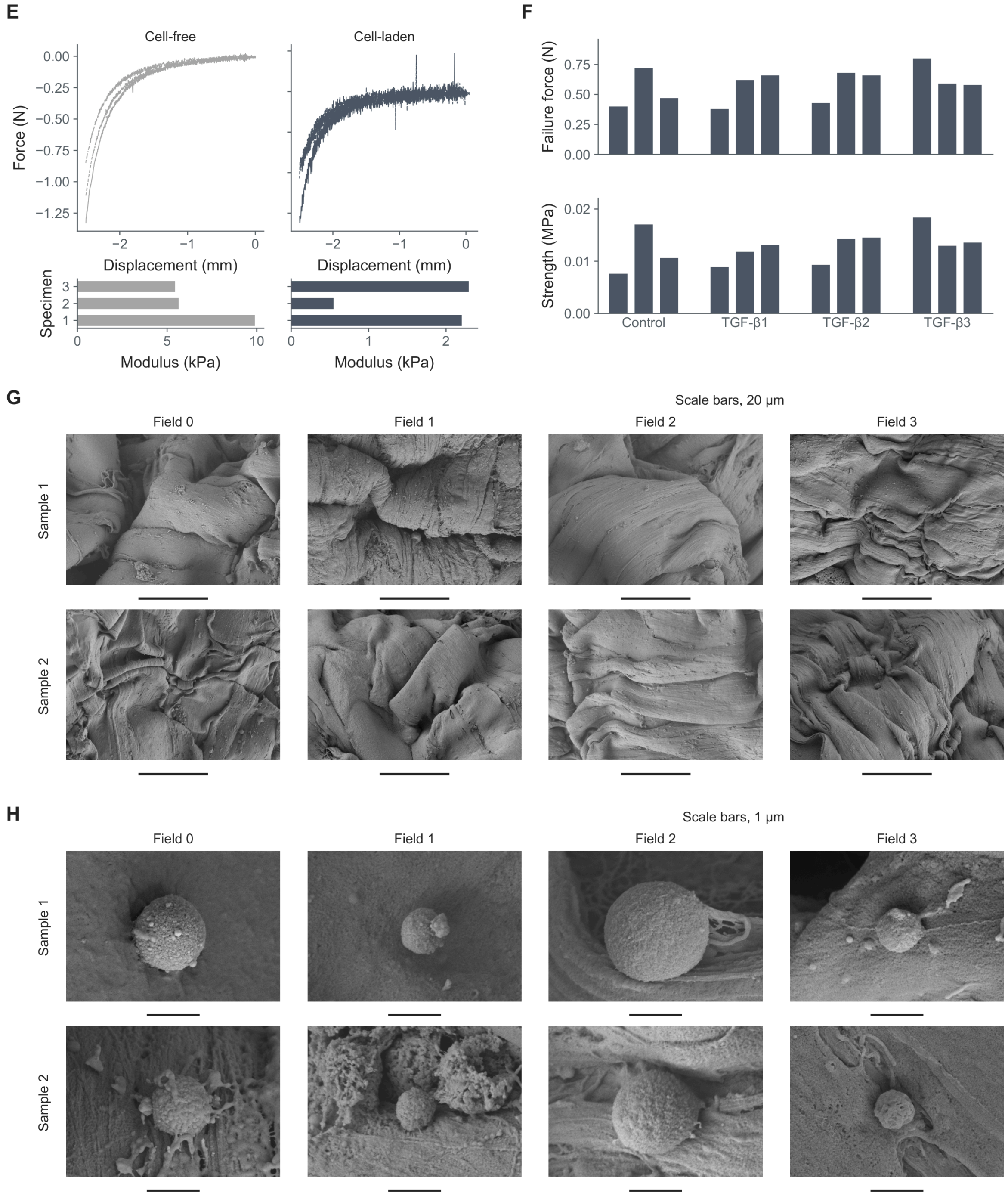

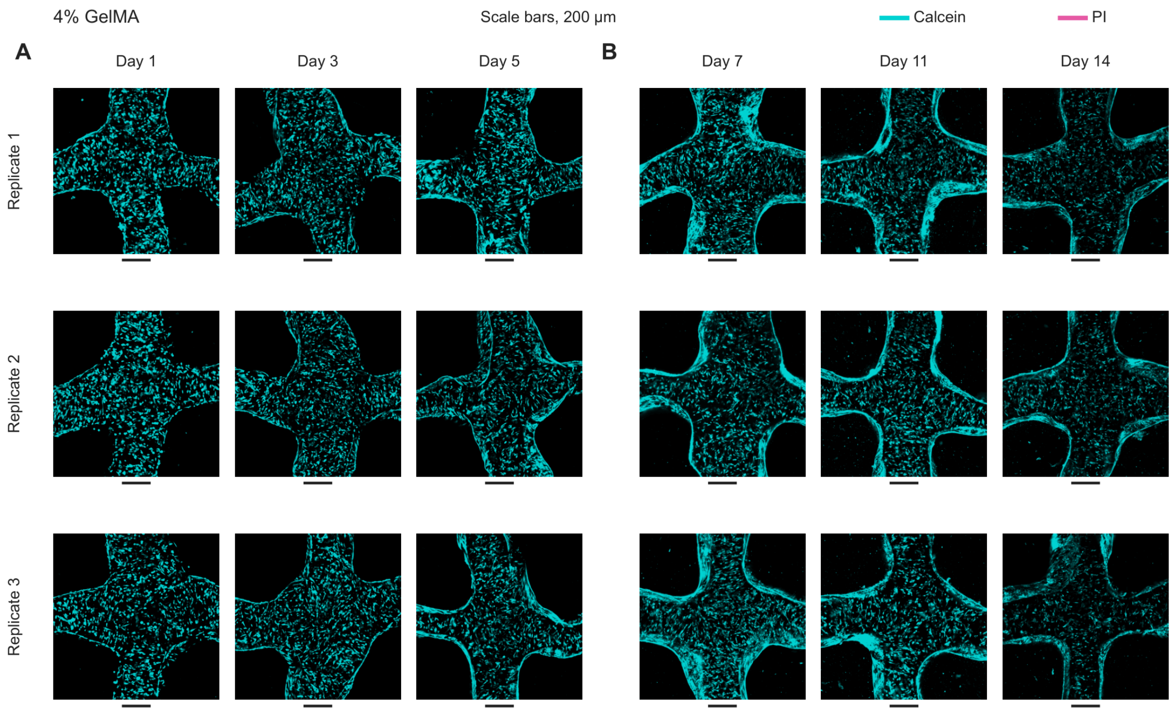


**Fig. S2. Live/dead fluorescence images across the GelMA concentration screen.** (A) Full-field live/dead fluorescence images in 4% gelatin methacryloyl (GelMA) on days 1, 3 and 5. (B) 4% GelMA on days 7, 11 and 14. (C) 5% GelMA on days 1, 3 and 5. (D) 5% GelMA on days 7, 11 and 14. (E) 6% GelMA on days 1, 3 and 5. (F) 6% GelMA on days 7, 11 and 14. (G) 8% GelMA on days 1, 3 and 5. (H) 8% GelMA on days 7, 11 and 14. (I) 10% GelMA on days 1, 3 and 5. (J) 10% GelMA on days 7, 11 and 14. (K) 12% GelMA on days 1, 3 and 5. (L) 12% GelMA on days 7, 11 and 14. Each image panel contains nine fields, with days as columns and three independent constructs per concentration and time point as rows, from one normal-donor cell isolate. Calcein and propidium iodide (PI) channels share a linear 0–255 display range (gamma 1). Scale bars, 200 µm. (M) Calcein-positive area as a fraction of the filament region of interest (ROI). Diamonds and lines show means, and circles show constructs. Mean symbols in M are offset horizontally for visibility. (N) PI objects per 100,000 ROI pixels, with bars showing means and circles showing constructs. M and N each show 108 constructs and 36 concentration–day means. Error bars are 95% t confidence intervals (mean ± t(0.975, 2) × standard deviation/√3), without truncation at zero. (O) Calcein-positive (left) and PI-positive (right) fractions of sampled filament volume in three independent constructs fabricated under identical conditions from the same cell batch, measured immediately after printing. Depth-bin boundaries are 0, 100, 200, 300, 400 and 500 µm, with A and B spanning 0–400 µm and C spanning 0–500 µm. All plotted axes are linear.

Fig. S2 (continued; 2 / 7)

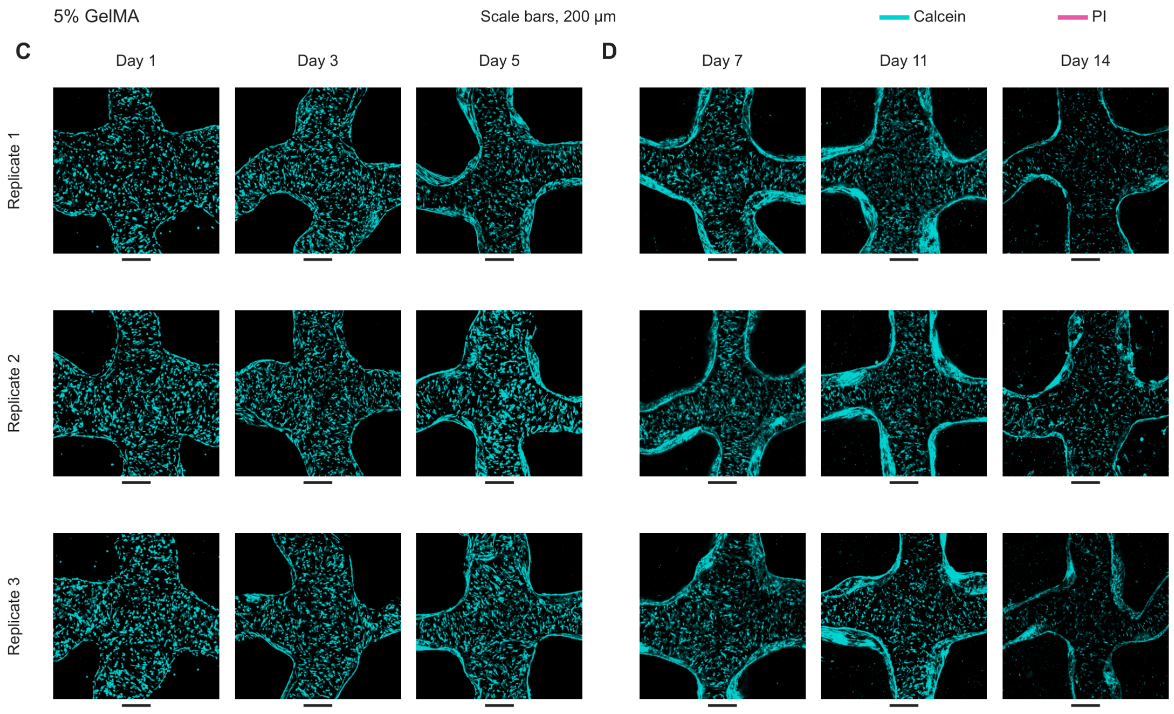

Fig. S2 (continued; 3 / 7)

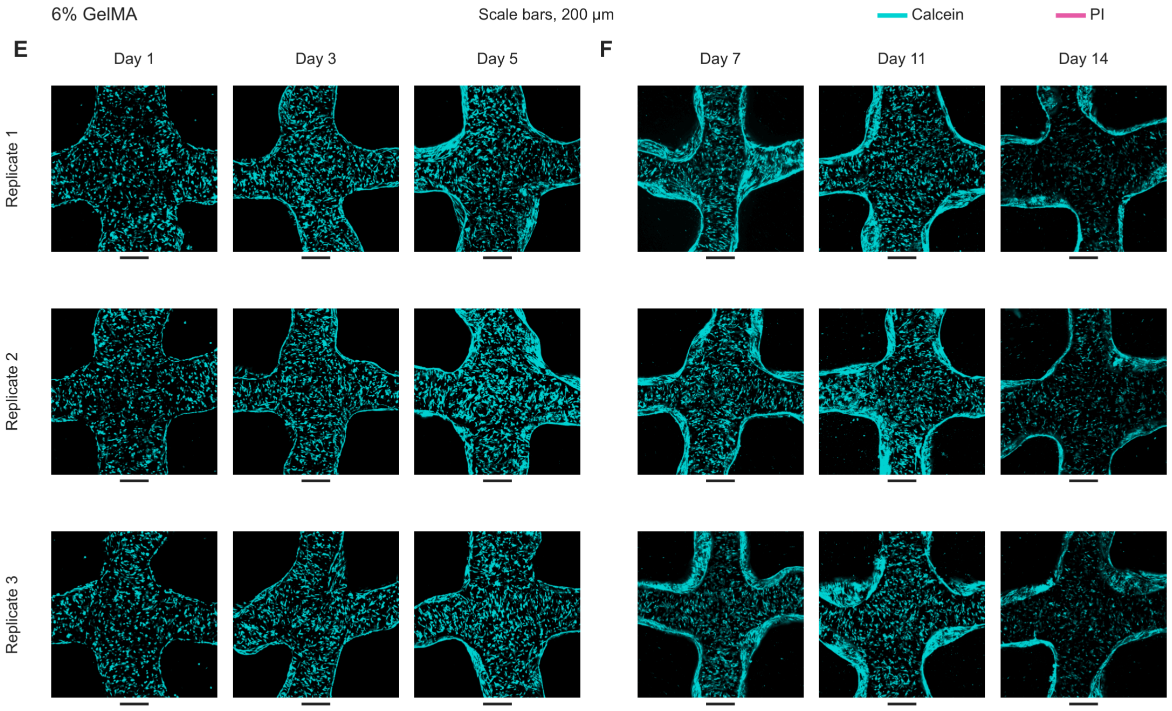

Fig. S2 (continued; 4 / 7)

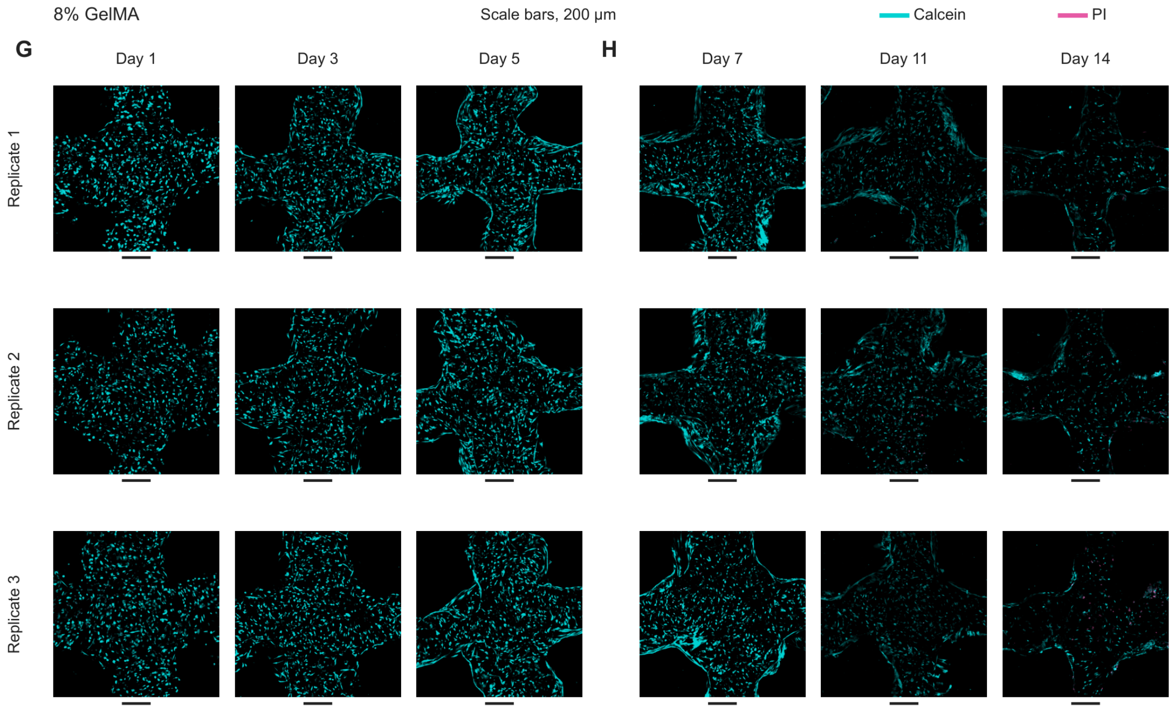

Fig. S2 (continued; 5 / 7)

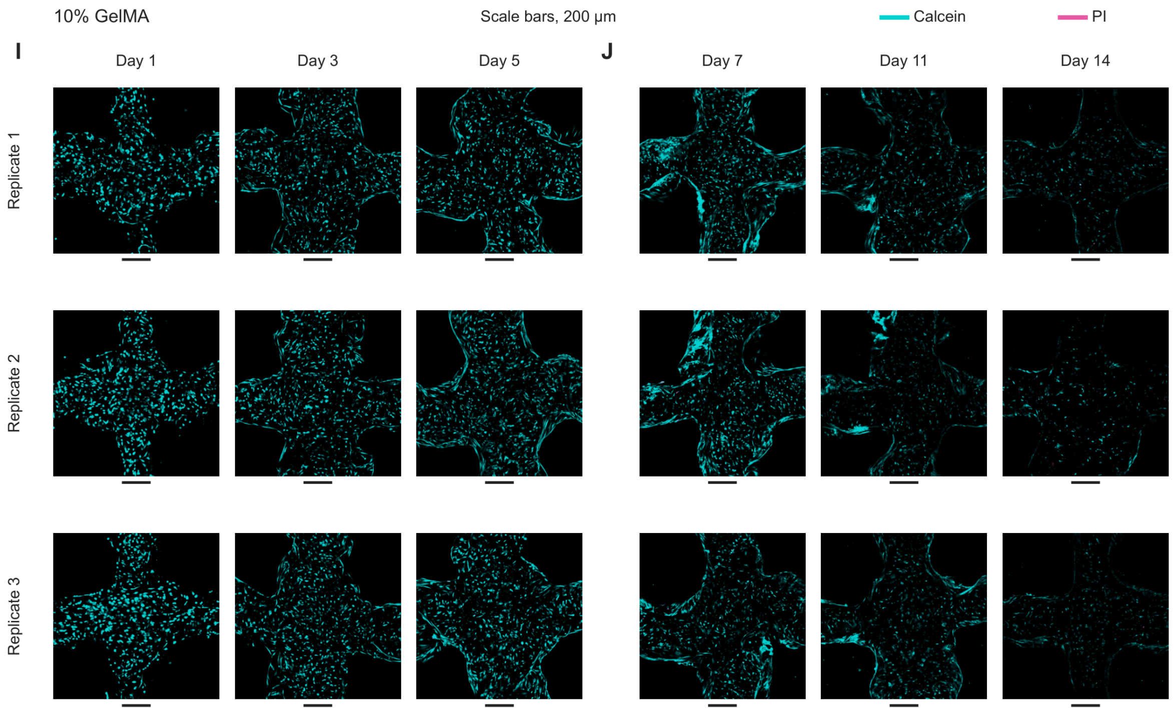

Fig. S2 (continued; 6 / 7)

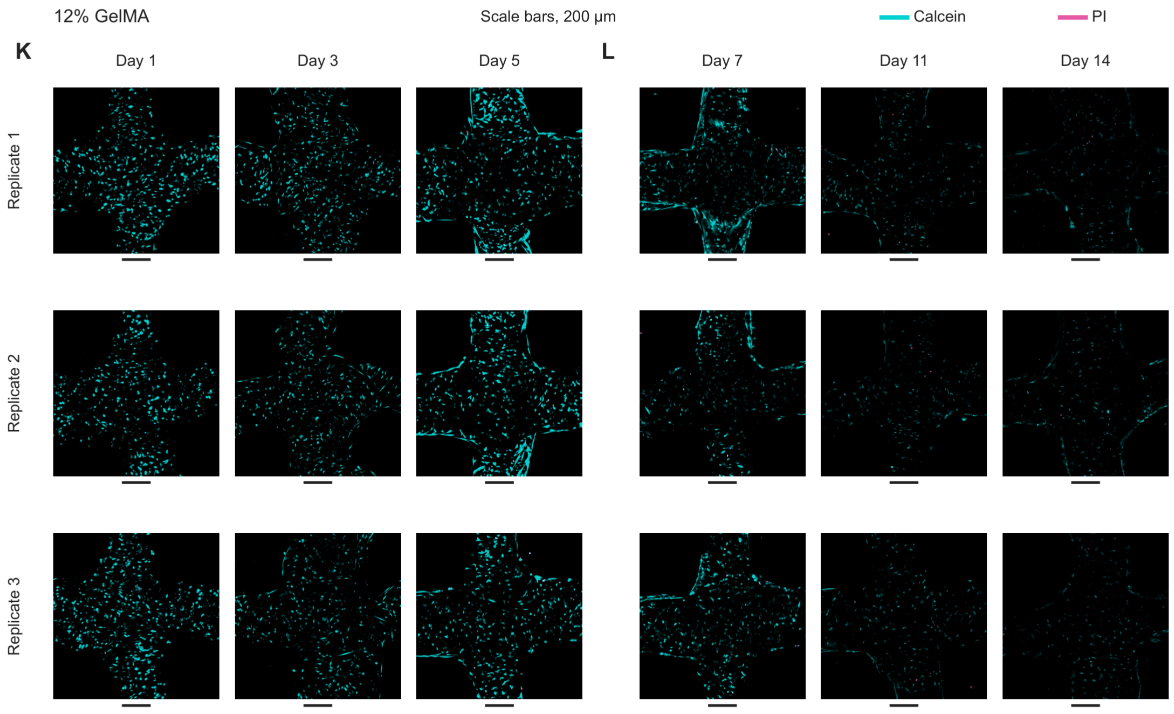

Fig. S2 (continued; 7 / 7)

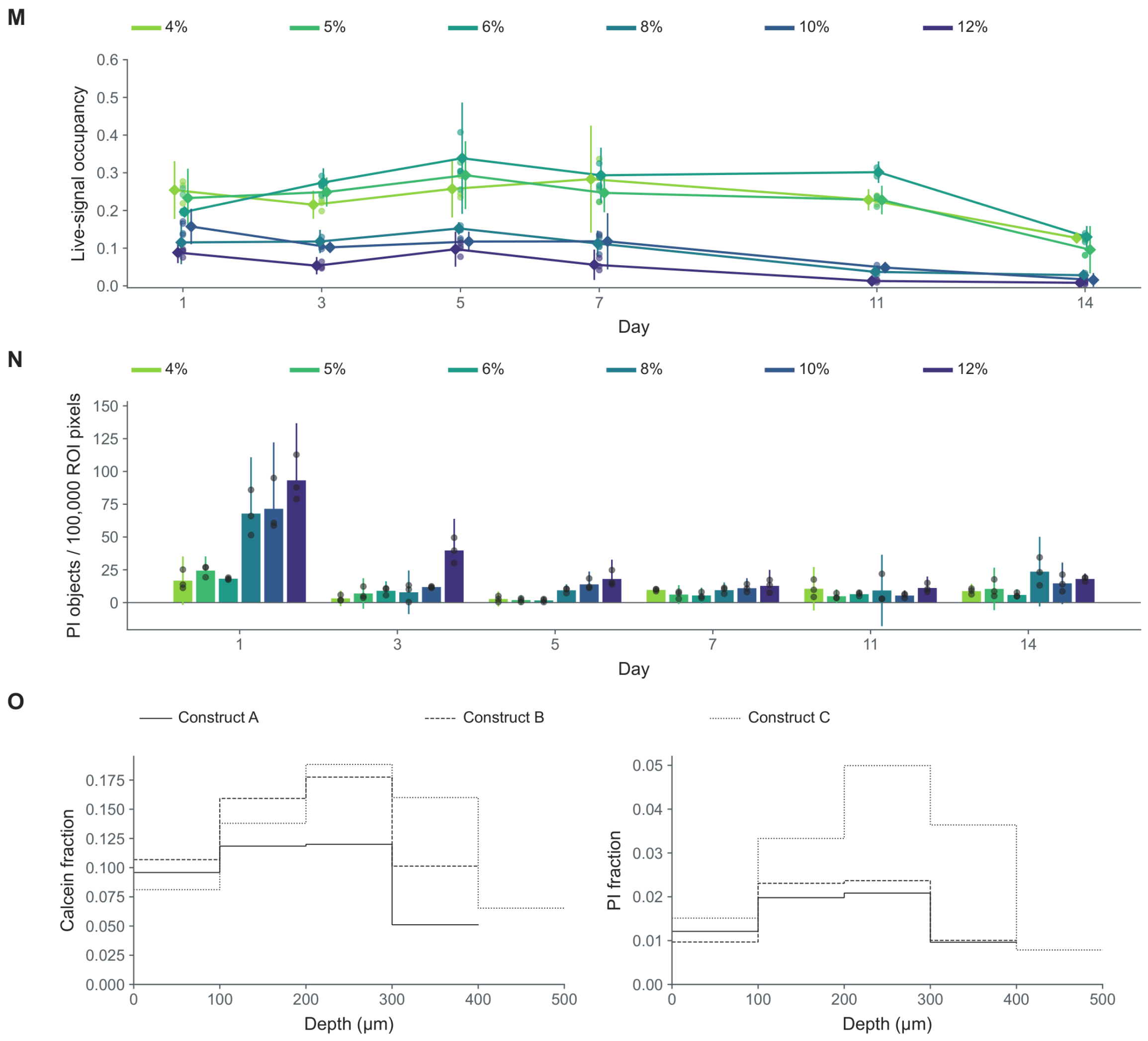

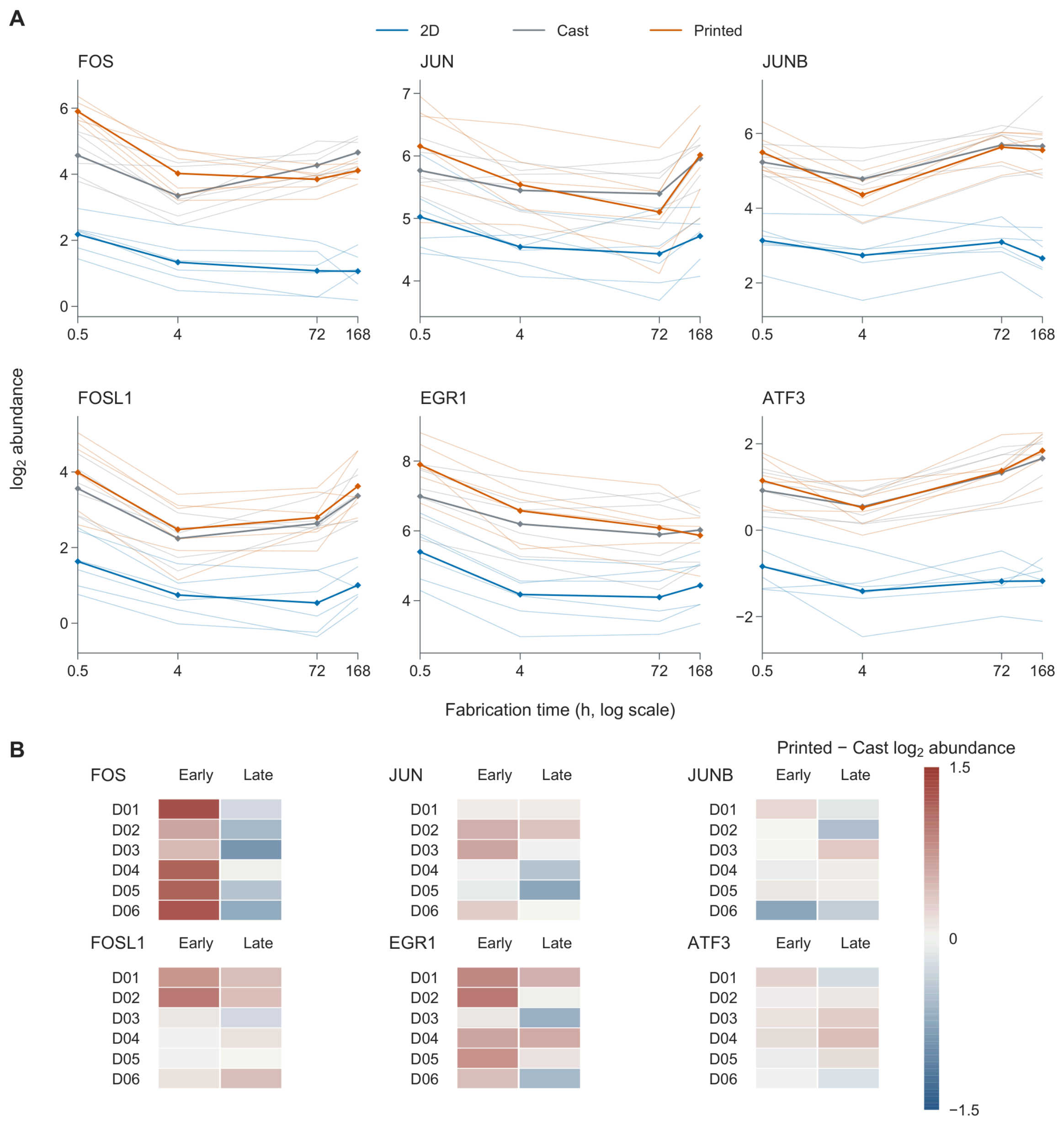


**Fig. S3. Immediate-early gene expression during construct fabrication.** (A) Quantitative polymerase chain reaction time courses of FOS, JUN, JUNB, FOSL1, EGR1 and ATF3 in unstimulated two-dimensional (2D), cast and printed cultures from six keratoconus (KC) donors. Expression is calibrated $\log_2$ abundance normalized to TBP, HPRT1 and RPLP0 after technical-well aggregation. Thin lines connect donor values, and thick lines with diamonds connect equal-donor arithmetic means. Each gene plot contains six donor trajectories per format at 0.5, 4, 72 and 168 h on a logarithmic time axis, giving 432 donor–gene–format–time observations and 72 means. (B) Heatmaps of printed-minus-cast $\log_2$ abundance differences for the same six genes and six KC donors. Each gene has six donor rows and two period columns, giving 72 cells. Each Early cell is the arithmetic mean of the matched 0.5 and 4 h differences and each Late cell of the matched 72 and 168 h differences. Both time points are required for each period mean. The shared color scale is centered on zero. No error bars or significance marks are shown.

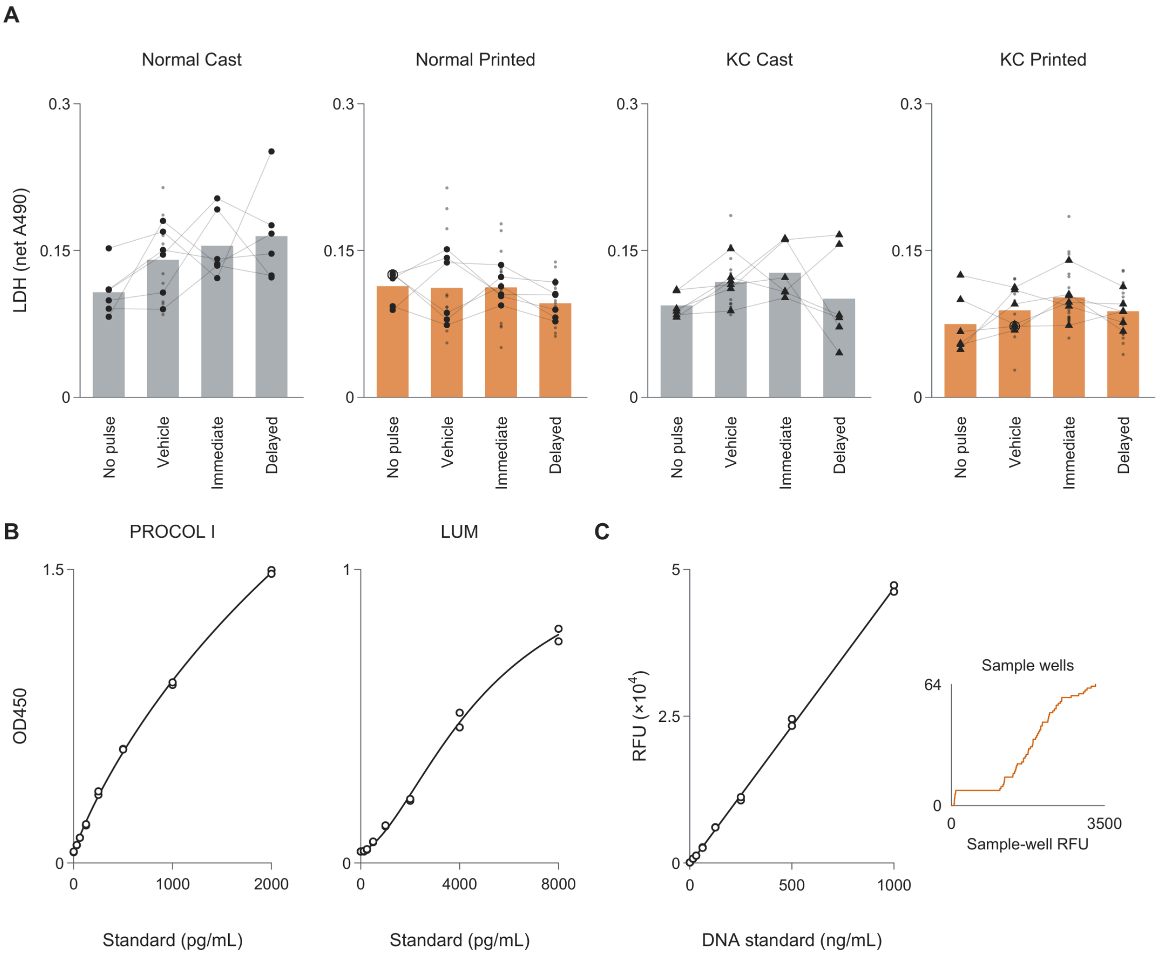


**Fig. S4. Lactate dehydrogenase release and assay calibrations in the functional batch.** (A) Lactate dehydrogenase (LDH) release in the 48–72 h collection window, expressed as background-corrected absorbance at 490 nm, in cast and printed constructs from six normal and six keratoconus (KC) donors. The four conditions are no pulse, vehicle, immediate SB431542 (4–72 h), and delayed SB431542 (24–72 h). Vehicle and both SB431542 arms received a 4 h transforming growth factor-β1 (TGF-β1) pulse. Small points show 168 constructs, large filled symbols show 96 donor–condition means, and lines connect the same donor. Each of the 16 bars shows the equal-donor arithmetic mean of six donor–condition means. Outer rings indicate coincident marks. (B) Procollagen I (PROCOL I) and lumican (LUM) enzyme-linked immunosorbent assay (ELISA) standard curves for the 48–72 h supernatant plates. Each plot shows 16 technical standard wells and a four-parameter logistic fit. Standard concentrations are in pg/mL, and optical density at 450 nm (OD450) is on the vertical axes. (C) Deoxyribonucleic acid (DNA) calibration from 16 technical standard wells, with a linear fit. Standard DNA concentration is shown in ng/mL and fluorescence in relative fluorescence units (RFU), displayed in units of $10^4$. The adjacent empirical cumulative distribution shows RFU horizontally and cumulative counts of 64 sample wells vertically over 0–3,500 RFU. The axes in (B) and (C) are linear. No error bars or significance marks are shown.

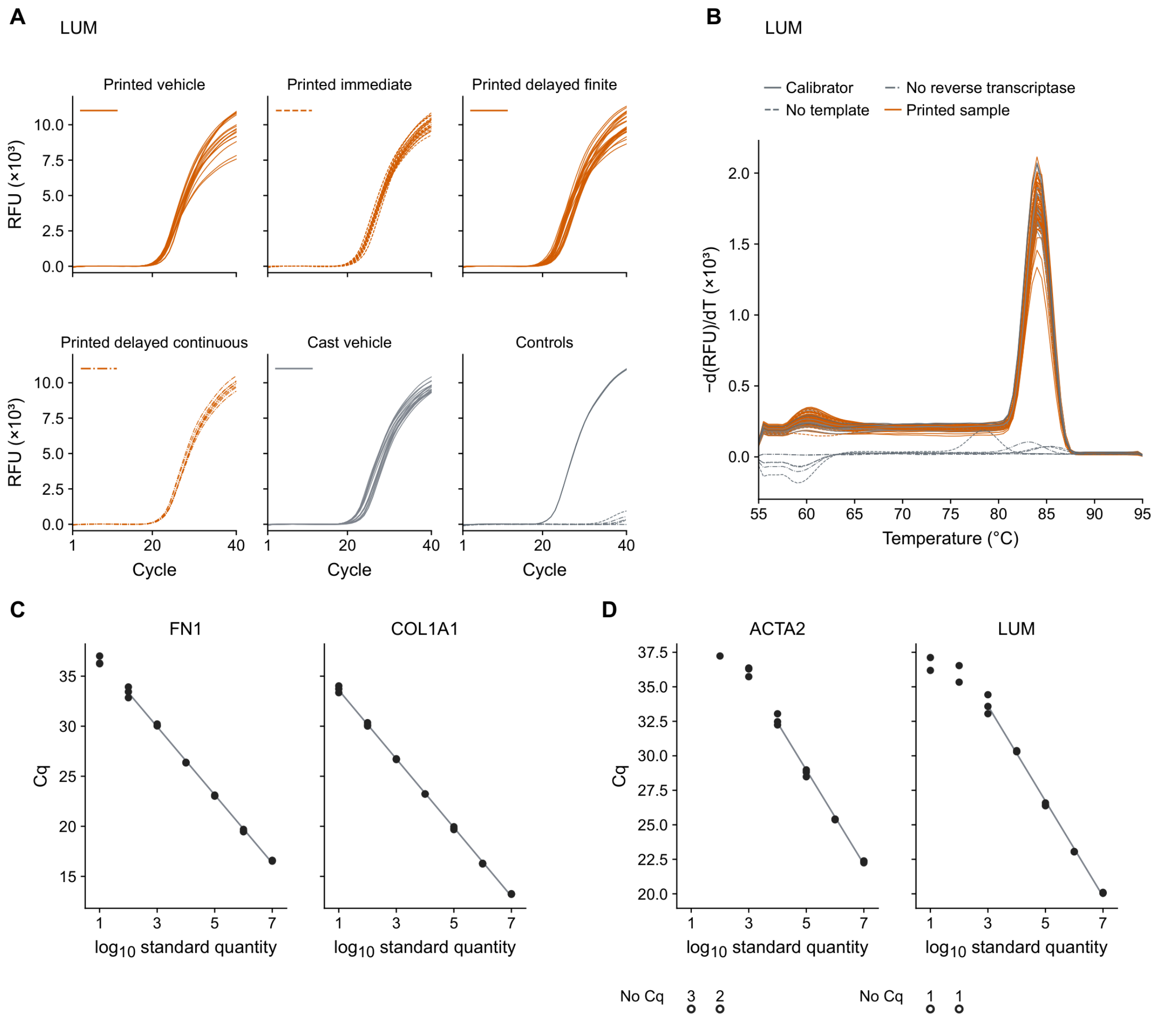


**Fig. S5. Amplification, melt curves, extract concentrations and assay calibrations.** (A) Lumican (LUM) amplification from 96 quantitative polymerase chain reaction (qPCR) wells over 40 cycles, in $10^3$ relative fluorescence units (RFU). Each curve represents one technical well: 88 sample wells from 44 constructs nested within four donors, plus two calibrator, two no-template and four no-reverse-transcriptase wells. Facets contain 16, 24, 24, 8, 16 and 8 wells, respectively, in reading order. Control line styles follow the key in (B). (B) Melt-curve derivatives of the same 96 wells from 55 to 95 °C in 0.5 °C steps, in $10^3$ RFU/°C. (C) Quantification cycle (Cq) against $\log_{10}$ standard quantity for fibronectin 1 (FN1) and collagen type I α1 (COL1A1), with three technical wells at each of seven quantities per target. Lines show linear fits over $\log_{10}$ quantities 2–7 and 1–7, respectively. (D) Cq against $\log_{10}$ standard quantity for actin α2 (ACTA2) and LUM, with three wells at each of seven quantities and linear fits over $\log_{10}$ quantities 4–7 and 3–7, respectively. In (C, D), filled circles show measurable Cq values. In (D), five ACTA2 and two LUM wells without Cq appear as open circles, with adjacent numbers giving the count at each dilution. In the separate No Cq lane, horizontal positions indicate standard quantity, while vertical positions do not represent Cq. (E) Type I collagen (COL I, ng/mL) and LUM (pg/mL) concentrations in construct extracts at 72 h, day 7 and day 14 (72, 168 and 336 h after pulse onset). Each analyte has nine boxes with individual extracts: 12 per box at 72 h and day 7 and 24 at day 14, nested within six normal and six keratoconus (KC) donors (168 cellular extracts per analyte). Boxes show medians and quartiles, and whiskers extend to the most extreme observations within 1.5 times the interquartile range beyond each box edge. COL I is shown on a linear axis and LUM on an inverse hyperbolic sine (asinh) axis with a linear width of 1,000 pg/mL. All cellular arms received a 4 h transforming growth factor-β1 (TGF-β1) pulse. Immediate and delayed SB431542 blockade were applied at 4–72 and 24–72 h, respectively, followed by withdrawal (stop), whereas delayed continued blockade lasted to 336 h. Vehicle arms received solvent after the pulse. Open squares show recorded concentrations of 14 COL I and 10 LUM cell-free extracts, all below the detection limit. At day 7, eight cell-free extracts (four per analyte) had no recorded concentration and are omitted without zero substitution. (F) COL I and LUM enzyme-linked immunosorbent assay (ELISA) calibration curves for day 14, each with 16 technical standard wells and a four-parameter logistic fit, with standards in ng/mL and pg/mL, respectively. Optical density (OD) was measured at 490 and 450 nm, respectively. Outer rings mark coincident wells, including two LUM wells at 2,000 pg/mL. (G) Ribonuclease P/MRP subunit p30 (RPP30) genomic DNA calibration, with 14 technical standard wells and a linear fit of Cq against $\log_{10}$ DNA mass (ng). Other axes are linear.

Fig. S5 (continued; 2 / 2)

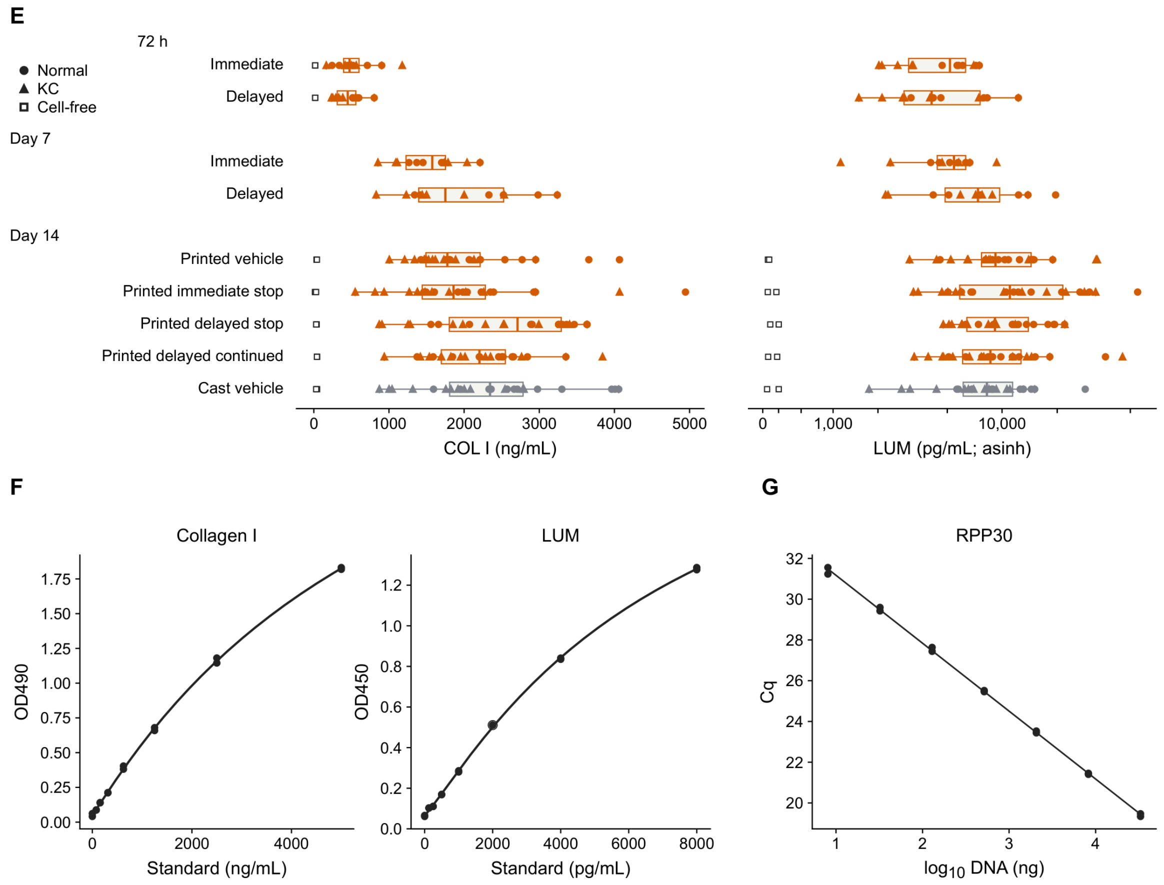

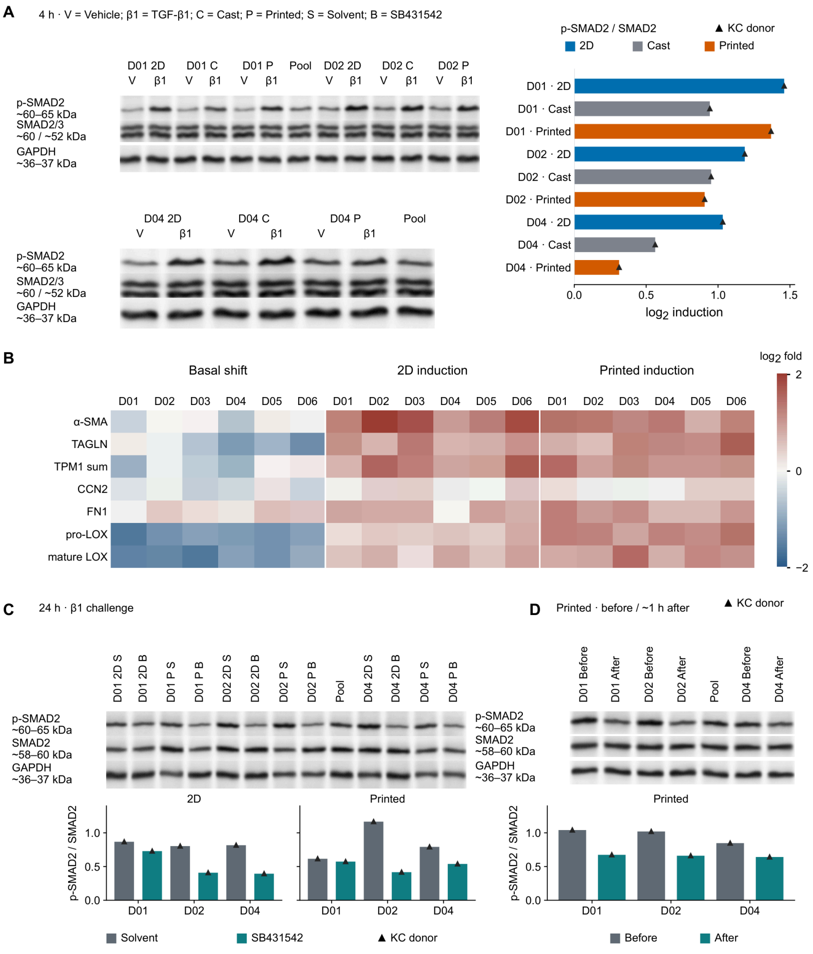


**Fig. S6. Protein components, SMAD2 signaling and isoform-resolved transcriptional responses across culture formats.** (A) Western blots of phosphorylated SMAD2 (p-SMAD2), total SMAD2/3 and glyceraldehyde-3-phosphate dehydrogenase (GAPDH) at 4 h. Three keratoconus (KC) donors contribute nine bars, each showing donor-level $\log_2$ induction of p-SMAD2/total SMAD2 by transforming growth factor-β1 (TGF-β1) relative to vehicle across two-dimensional (2D), cast and printed cultures. Total SMAD2 was quantified from the upper band of the SMAD2/3 doublet. The two sets of membrane strips have 13 and seven lanes, respectively, including Pool. (B) Donor-level $\log_2$ basal shifts (printed/2D, vehicle) and TGF-β1/vehicle induction within each format at 72 h. The seven rows show α-smooth muscle actin (α-SMA), transgelin (TAGLN), summed tropomyosin 1 (TPM1), cellular communication network factor 2 (CCN2), fibronectin (FN1), pro-lysyl oxidase (pro-LOX) and mature lysyl oxidase (LOX). Each of three contrasts has columns for six KC donors (D01–D06), totaling 126 cells. Values were normalized to DNA equivalents and the pooled calibrator. Each cell is one donor contrast on a zero-centered −2 to 2 scale, with endpoint colors for values outside this range. (C) Blots and raw p-SMAD2/total SMAD2 ratios at 24 h during a TGF-β1 challenge with solvent or SB431542 in 2D and printed cultures. Three KC donors contribute 12 bars, one per donor, format and condition. The membrane strips have 13 lanes including Pool. (D) Blots and raw p-SMAD2/total SMAD2 ratios in printed cultures with TGF-β1 history, before delayed SB431542 dosing and about 1 h afterward. Three KC donors contribute six bars, one per donor and condition. The membrane strips have seven lanes including Pool. In (A), (C) and (D), continuous native membrane crops are shown on a linear 0–255 grayscale without nonlinear adjustment. Bars represent individual donor values on linear axes without error bars. Pool denotes the pooled calibrator. Labeled molecular masses are expected values from antibody records. Total SMAD2 is the denominator throughout, using the single SMAD2 band in (C, D). Full membranes and lane assignments are in Supplementary Data 3. (E) Gene-level format-by-stimulus interactions for TGF-β1 versus TGF-β2 (upper) and TGF-β3 (lower), estimated from three KC donors. The interaction is the stimulus-induced $\log_2$ fold change in printed minus that in 2D culture. The axes are linear. Each plot contains the same 15,989 genes, one point per gene at its continuous coordinates, colored by occupancy in 0.2 × 0.2 bins on a shared logarithmic scale (1–683 genes per bin). Dashed y = x lines denote equal isoform interactions and pale horizontal and vertical lines mark zero. A likelihood ratio test of equal format-by-stimulus interactions among the three isoforms (2 degrees of freedom) and tests of the three pairwise differences in these interactions each gave no gene with Benjamini–Hochberg-adjusted $p < 0.05$ among the 15,989 genes. (F) Empirical cumulative distributions of printed/2D ratios of TGF-β-induced increases for 116 matrix and 110 contractile genes under each of TGF-β2 and TGF-β3 (452 gene–ligand records). Each ratio compares stimulus-minus-vehicle normalized counts, with the positive 2D increase as denominator. Each rug marks one gene–ligand ratio. The horizontal axis is symmetric logarithmic, linear from −1 to 1, with untransformed ticks. The vertical reference at 1 denotes equal induced increases between formats. The linear vertical axis gives the fraction of genes at or below each ratio.

Fig. S6 (continued; 2 / 2)

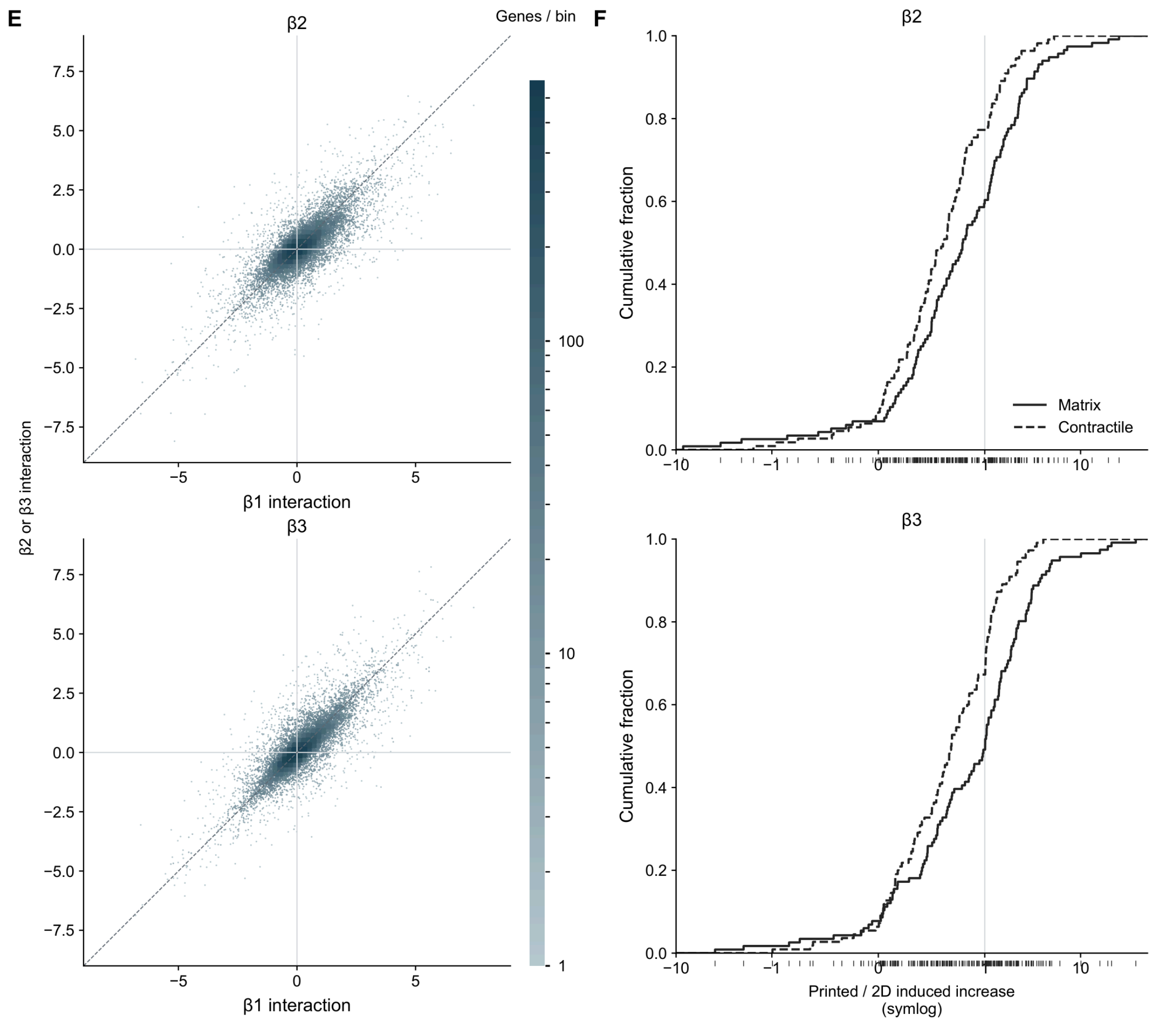

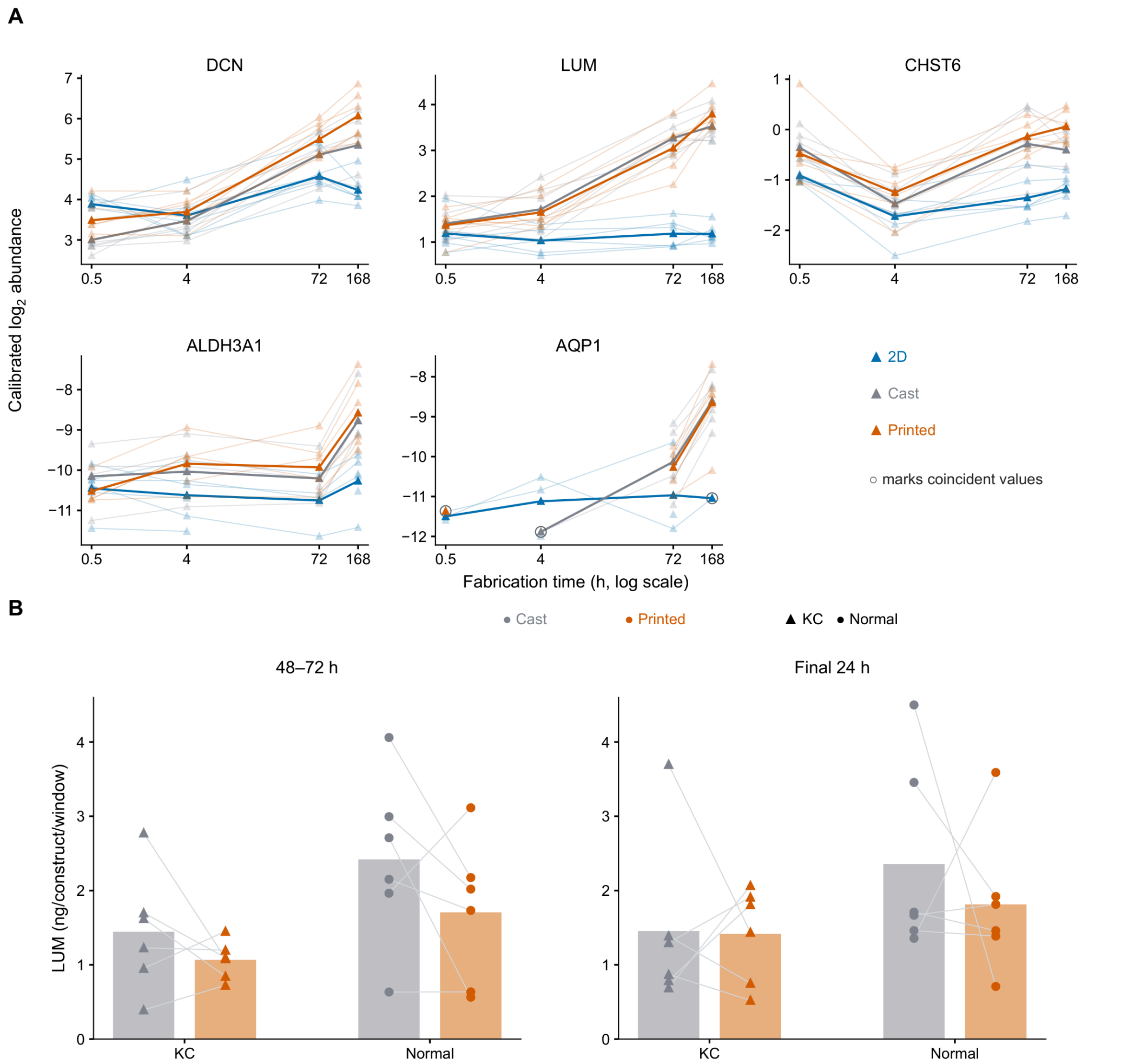


**Fig. S7. Stromal identity expression and lumican in medium and retained matrix.** (A) Fabrication time courses of decorin (DCN), lumican (LUM), carbohydrate sulfotransferase 6 (CHST6), aldehyde dehydrogenase 3 family member A1 (ALDH3A1) and aquaporin 1 (AQP1) in unstimulated two-dimensional (2D), cast and printed cultures from six keratoconus (KC) donors at 0.5, 4, 72 and 168 h. The five genes have separate vertical ranges and share the calibrated $\log_2$ abundance axis title. Fabrication time (h) is plotted on a logarithmic horizontal axis. Small symbols and thin paths show 313 donor–gene–format–time measurements, and larger symbols and thick paths show 58 arithmetic means of one to six available donors. Technical wells are aggregated before donor-level display, and missing time points are left unconnected. (B) Medium LUM in cast and printed vehicle-treated constructs after a transforming growth factor-β1 pulse, in ng/construct/window, during 48–72 h and the final 24 h ending at 336 h after pulse onset. Points show 48 measurements from six KC and six normal donors, with one construct per donor–format–window. The eight bars are arithmetic means of six donors each. Lines pair formats within donors. (C) Retained LUM (ng/construct) in printed sister constructs at 0, 4 and 11 days after SB431542 withdrawal at 72 h. Immediate and Delayed denote SB431542 at 4–72 h and 24–72 h after pulse onset, respectively. Each of the 12 facets represents one donor (D01–D06, KC; N01–N06, normal). Small pale symbols show 82 constructs, and larger symbols and lines show 67 donor–regimen–harvest means. Missing harvests are left unconnected. (D) Retained LUM and native type I collagen in cast and printed constructs receiving vehicle after the pulse and harvested at 336 h, in ng/construct. Small symbols show 86 construct measurements (42 LUM and 44 collagen), and larger symbols show 44 donor means. Each of the eight bars is the equally weighted mean of five KC or six normal donor means, with constructs averaged within donors. D05 is absent. Lines pair formats within donors. Paired donor-level comparisons of these constructs are in Fig. 2G. (E) Within-donor $\log_2$ delayed/immediate ratios of 48–72 h procollagen I secretion per µg DNA in cast and printed constructs from 12 donors. The two bars are arithmetic means of donor $\log_2$ ratios, and their labels are geometric mean ratios. Points show 24 donor–format values, paired by lines. The dashed line is the $\log_2$ of their equally weighted pooled geometric mean. The zero line denotes equal secretion. The printed $\log_2$ ratios were tested against zero using a two-sided paired exact sign-flip test ($p$ = 0.0029, $**p < 0.01$). Triangles denote KC and circles denote normal donors, including the small construct symbols in C. Outer circles mark coincident values.

Fig. S7 (continued; 2 / 2)

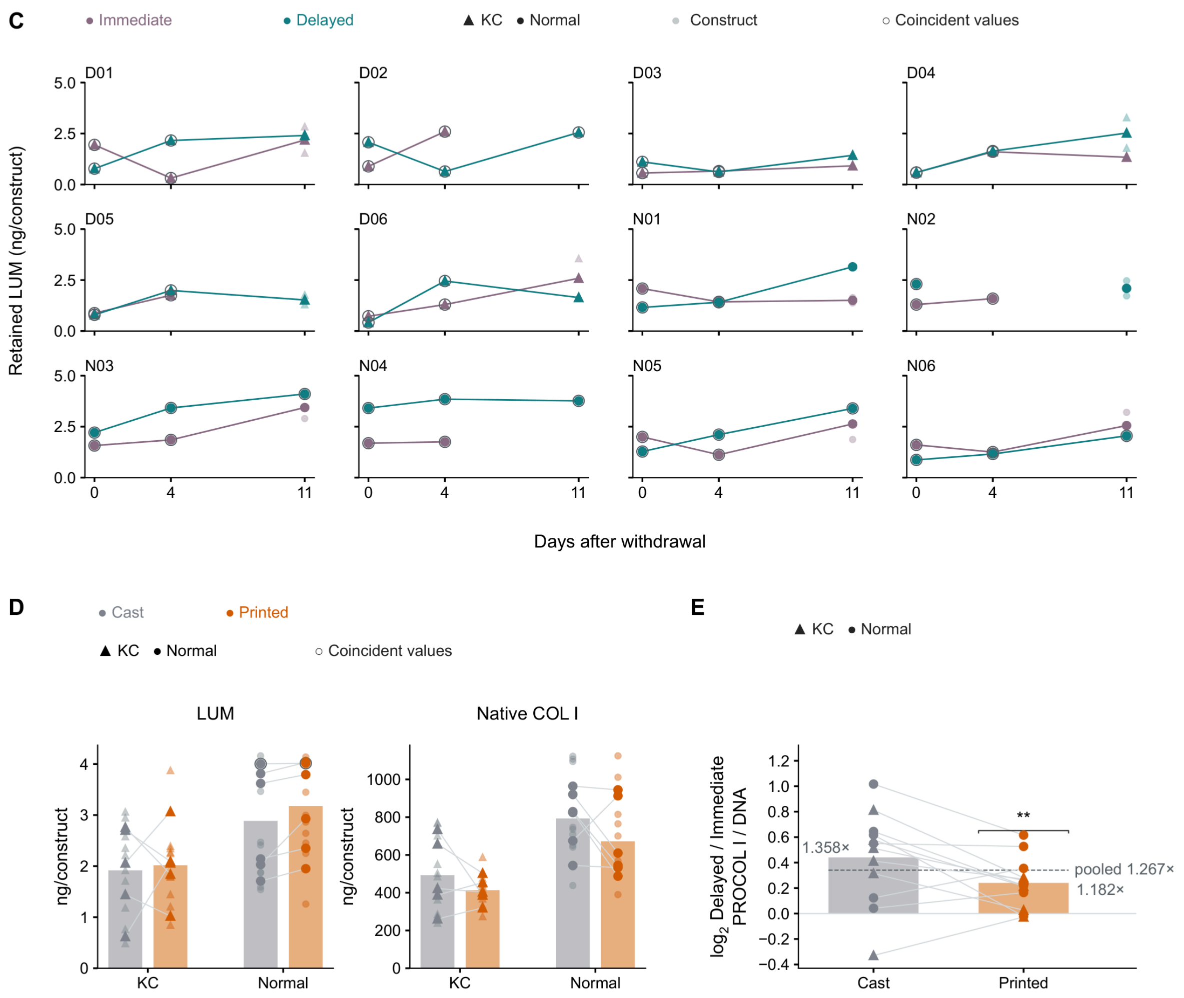

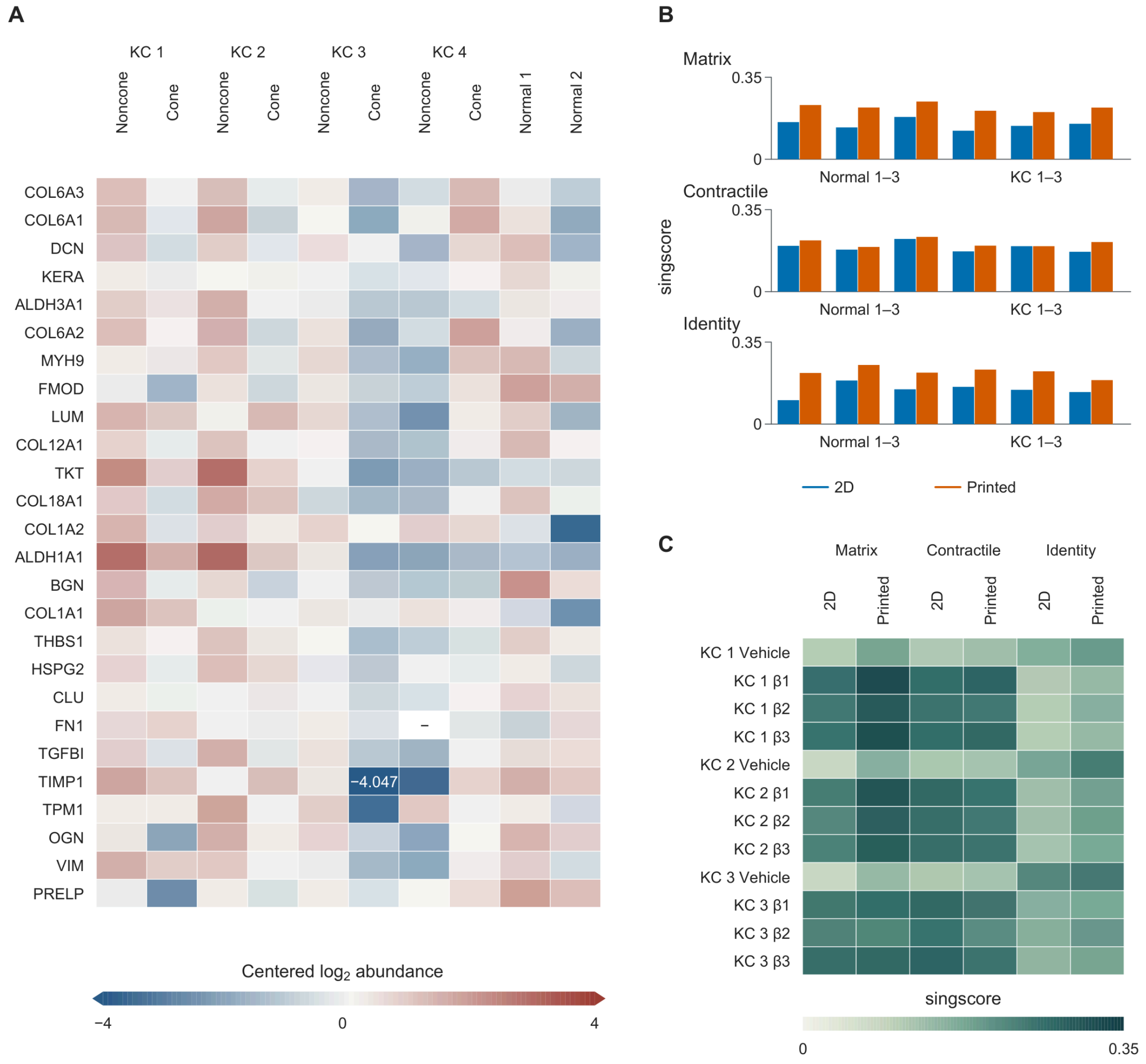


**Fig. S8. Patient stromal proteins and construct RNA program scores.** (A) Matrix, stromal identity and contractile proteins in the published stromal proteome of Yam et al., comprising cone and non-cone regions from four keratoconus (KC) corneas and two normal corneas. Rows show 26 proteins and columns show ten tissue samples. Cell colors show $\log_2$ abundance centered on each protein's mean across available samples, with the color scale capped at −4 and 4. The out-of-range TIMP1 value is printed. A dash marks the missing fibronectin (FN1) value in KC4 non-cone tissue. (B) Matrix, contractile and stromal identity gene-program scores calculated with singscore for 12 unstimulated RNA sequencing samples from six donors, comprising three normal and three KC donors, each in two-dimensional (2D) and printed culture. The 36 bars show individual sample scores on linear vertical axes. (C) The same program scores for 24 samples from three KC donors, each in both formats under vehicle or transforming growth factor-β1, β2 or β3. The 12 rows identify donor–stimulus combinations and the six columns pair formats within each program, giving 72 scores on a shared absolute color scale from 0 to 0.35.

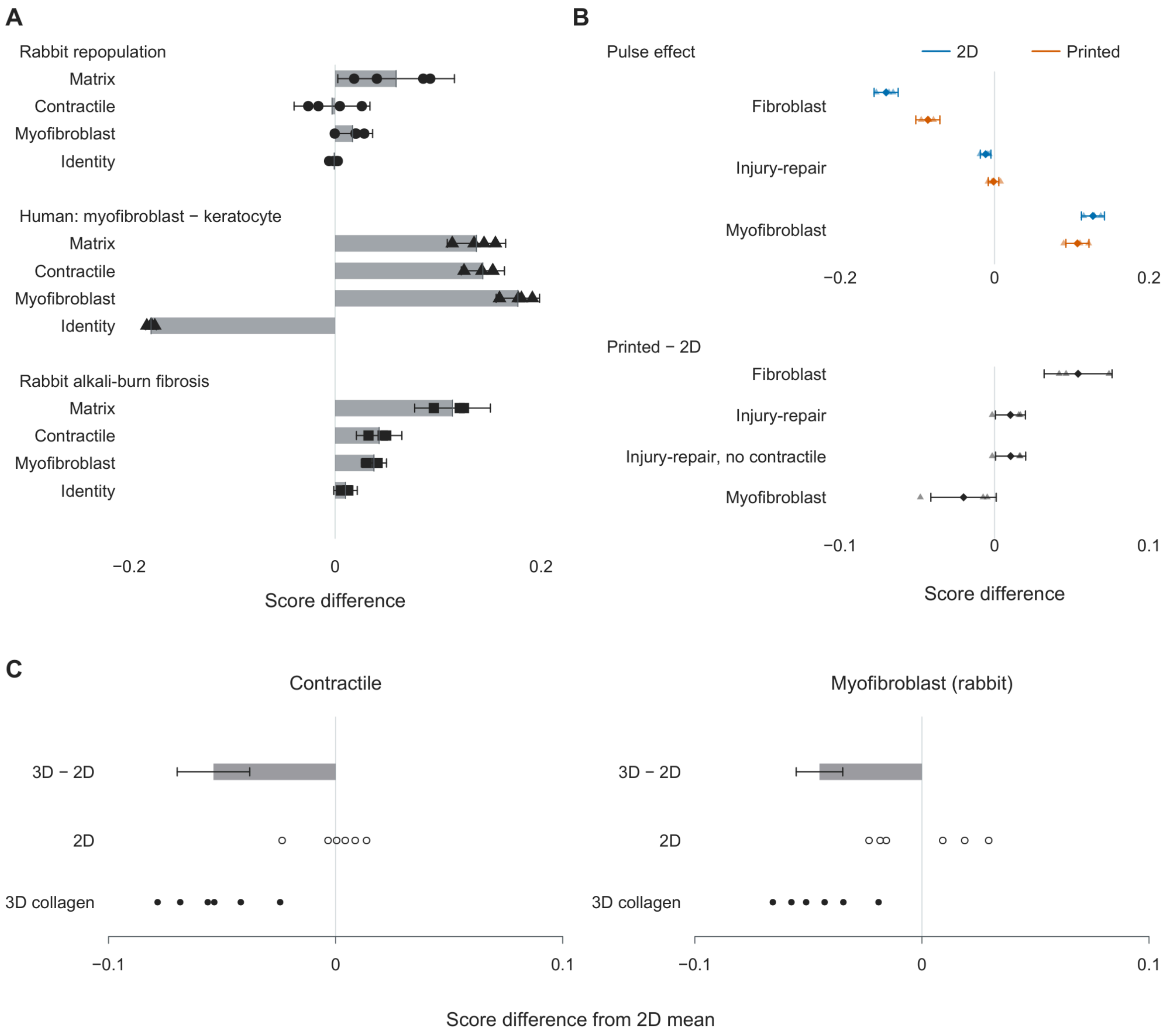


**Fig. S9. Matrix, contractile and stromal identity programs in repopulating stroma, fibrotic stroma and independent three-dimensional culture.** (A) Twelve horizontal bars show mean differences in centered singscore for four programs in three tissue references. Circles show paired differences between fibroblasts repopulating freeze-injured rabbit stroma and resting keratocytes from the same animals' uninjured eyes (GSE328779, four rabbits). Triangles show human myofibroblasts minus keratocytes (GSE308374, four curvature blocks). Squares show extracellular matrix-remodeling cells and activated keratocytes 14 days after rabbit corneal alkali burn minus uninjured keratocytes (GSE302936, three injured and three uninjured libraries), with each injured library centered on the uninjured-library mean. Horizontal error bars are 95% confidence intervals from the t distribution for paired rabbit and human block means (3 degrees of freedom) and Welch t intervals for the unpaired alkali-burn comparison. (B) Effects of the transforming growth factor-β1 (TGF-β1) pulse on the human fibroblast-state, injury-repair and human myofibroblast-state programs in cultures from three keratoconus (KC) donors. In the upper plot, six estimates show TGF-β1 minus control from the same donor and format in two-dimensional (2D) and printed cultures, with three donor differences per estimate (18 points). In the lower plot, four estimates show printed minus 2D pulse effects, including the injury-repair program without contractile genes, with three donor interactions per row (12 points). Each interaction is the printed pulse effect minus the 2D pulse effect, calculated from four matched sample scores per donor. Diamonds and horizontal bars show donor-adjusted linear-model estimates and 95% confidence intervals (14 degrees of freedom), and triangles show donors. (C) Contractile and rabbit myofibroblast-state scores of rabbit keratocytes in three-dimensional (3D) fibrillar collagen and 2D collagen-coated or fibrillar culture in GSE337743 (serum-free, no ligand). Each plot contains six open circles for 2D and six filled circles for 3D collagen, each centered on the six-library 2D mean. Bars show 3D minus 2D estimates with 95% confidence intervals from a linear model on culture group (10 degrees of freedom). All horizontal axes show score differences on linear scales. Vertical reference lines mark no difference. Gene-set definitions are given in Methods and all plotted values are in Table S3.

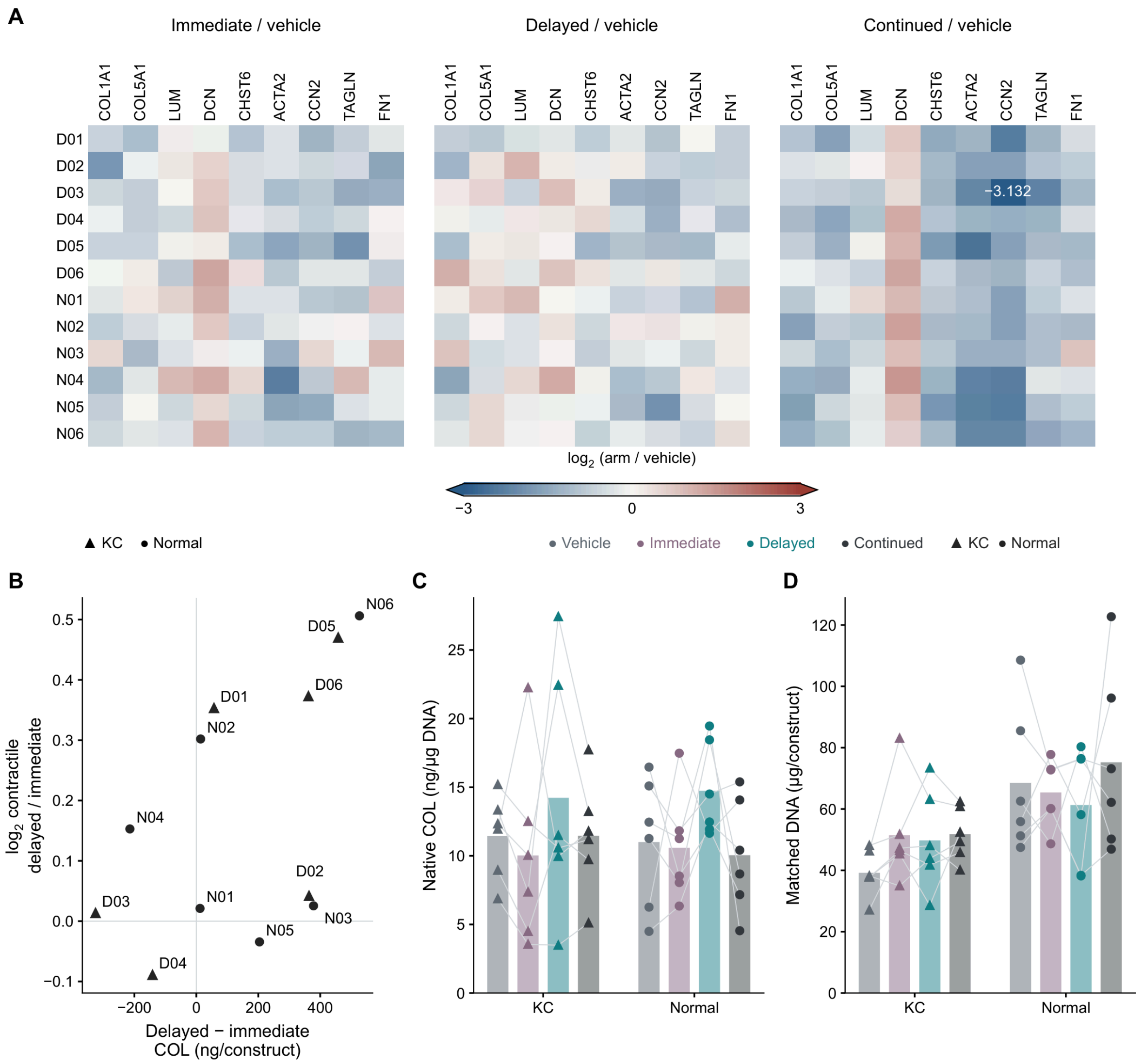


**Fig. S10. Day-14 transcription, retained collagen, and matched DNA across the treatment arms.** (A) Donor-specific $\log_2$ expression differences from vehicle for nine genes at 14 days after pulse initiation. Each block contains 12 donor rows (D01–D06, keratoconus (KC); N01–N06, normal) and nine gene columns, giving 324 donor–gene–contrast values across three blocks. The shared color scale is capped at −3 and 3. The sole below-range cell, D03/Continued/CCN2, is shown in the endpoint color and labeled −3.132. All arms received a 4 h transforming growth factor-β1 (TGF-β1) pulse. Immediate and Delayed denote SB431542 at 4–72 h and 24–72 h, respectively, and Continued denotes SB431542 from 24 to 336 h. Vehicle received solvent after the pulse. (B) Retained native type I collagen difference (ng/construct), Delayed minus Immediate, against the $\log_2$ Delayed/Immediate ratio of contractile gene expression, comprising ACTA2, CCN2 and TAGLN. Each labeled point is one donor (n = 12), and zero lines mark equality between arms. (C) Retained native type I collagen normalized to deoxyribonucleic acid (DNA), in ng per µg DNA, across the four printed arms. (D) Mean DNA quantity matched to the collagen measurements, in µg per construct, for the same 48 donor-arm combinations. C and D each show 48 donor values and eight bars, each giving the arithmetic mean of six donors within disease and arm, without error bars. Lines connect values from the same donor. (E) Retained native type I collagen (ng per µg DNA) in the Immediate and Delayed arms at withdrawal (day 0, 72 h), day 4 (168 h) and day 11 (336 h) after SB431542 withdrawal, with one panel per donor. The panels contain 96 sister-construct measurements and 72 donor–arm–harvest means from 12 donors. Large filled symbols show donor means, pale small symbols show constructs, and lines connect donor means. Outer rings identify coincident values without adding observations. (F) Native collagen under secretion-selected and retention-selected schedules at the same times. Each donor's schedule is fixed by the Fig. 8B selection. Secretion selection used the 48–72 h secretion window and retention selection used 336 h collagen. Left, sums of 12 donor-level means (ng/construct), with retention-selected minus secretion-selected differences of −258, −44 and +1,632 ng at days 0, 4 and 11, respectively. Right, 12 donor differences at each time, with the six changed choices labeled at day 11. The six unchanged choices coincide at zero and are indicated by an outer ring. All axes are linear in the quantities shown.

Fig. S10 (continued; 2 / 2)

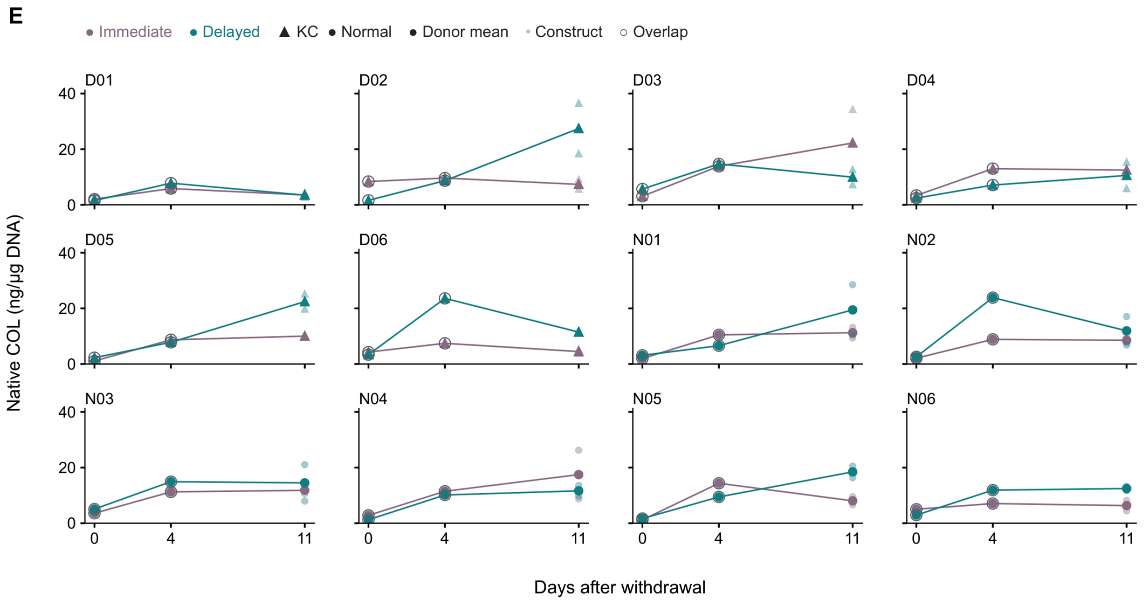


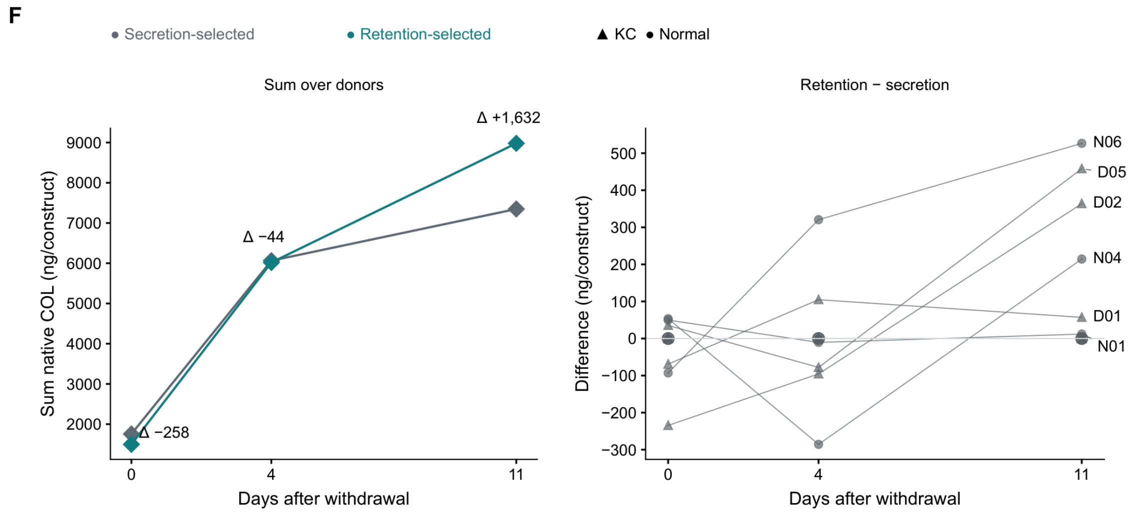

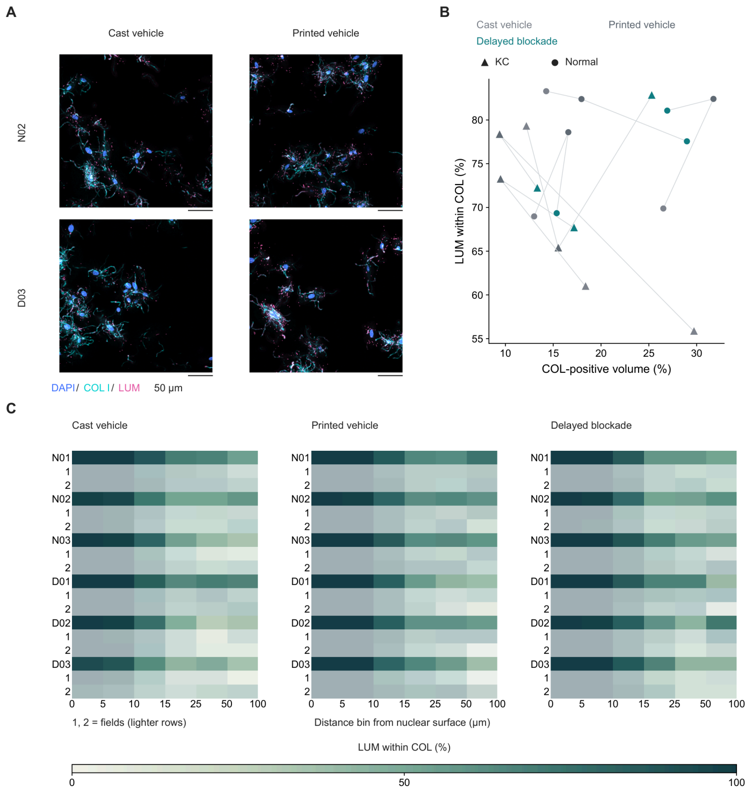


**Fig. S11. Lumican–collagen spatial relationships and α-smooth muscle actin readouts at day 14.** (A) Representative whole-mount confocal maximum-intensity projections at day 14, arranged with normal donor N02 above keratoconus (KC) donor D03 and cast vehicle beside printed vehicle. Each donor contributes one field from one construct per condition, giving four fields from four constructs. Images combine 4′,6-diamidino-2-phenylindole (DAPI), type I collagen (COL I, abbreviated COL in quantitative panels) and lumican (LUM). Display ranges are linear and shared between conditions. Scale bars, 50 µm. Vehicle denotes solvent after a 4 h transforming growth factor-β1 (TGF-β1) pulse, and Delayed blockade denotes SB431542 treatment at 24–72 h. (B) COL-positive gel volume (%) against LUM coverage within COL-positive volume (%) in 18 set B constructs from six donors (KC D01–D03 and normal N01–N03). Cast vehicle, printed vehicle and delayed blockade are distinct conditions, with one construct per donor per condition. Lines connect the three constructs from each donor. (C) Conditional LUM coverage within COL-positive volume across six distance intervals from the nearest nuclear surface, shown separately for the three conditions. Each heatmap has 18 rows comprising six construct rows with two field rows beneath each, and six columns representing 0–5, 5–10, 10–15, 15–25, 25–50 and 50–100 µm. The unequal physical intervals are displayed as equal-width columns, with ticks at interval boundaries. The shared color scale is 0–100%. (D) Individual grayscale maximum-intensity projections of four fields from four sister constructs from KC donor D03 under printed vehicle and delayed blockade, corresponding to Fig. 6A,B. Set A contains DAPI, COL I and α-smooth muscle actin (α-SMA), and set B contains DAPI, COL I and LUM. Linear display ranges, shared between conditions within each set, are 0–4095 detector counts for DAPI and COL I and 0–1997 for α-SMA in set A, and 0–4095 for DAPI and LUM and 0–3950 for COL I in set B. Scale bars, 50 µm. (E) From left to right, control-defined α-SMA-positive neighborhoods (%), α-SMA fiber length per nucleus (µm/nucleus), COL coherence and COL–filament angle (°). The first two plots each include no pulse, printed vehicle and delayed blockade, with six donors per condition and 18 construct values per plot. No pulse denotes printed constructs without a TGF-β1 pulse. COL coherence compares cast vehicle and printed vehicle, with six donors per condition and 12 construct values. COL–filament angle includes five printed vehicle donors and four delayed blockade donors, giving nine construct values. Construct values pool two fields. Bars are arithmetic means of donor values, points are construct values and lines connect values within donors. (F) Exact volume-weighted empirical cumulative distributions of COL connected-component volume (µm³, logarithmic axis) in printed vehicle and delayed blockade. Each condition includes 24 fields from 12 constructs and six donors, with 144,933 and 126,631 components, respectively. Each curve gives the cumulative fraction of total COL volume contained in components up to that volume. (G) COL fractions (%) under printed vehicle and delayed blockade, comprising collagen-positive fraction of sampled gel volume (Gel COL), median collagen coverage of extranuclear neighborhoods within 15 µm of the nearest nuclear surface (15 µm COL), and fraction of gel volume occupied by collagen-positive components spanning at least two nuclear neighborhoods (Spanning COL). Gel COL and 15 µm COL each comprise 48 field values, 24 construct values and 12 donor values, whereas Spanning COL comprises 24 construct values and 12 donor values. The six paired donors include three KC and three normal donors, with staining sets A and B equally weighted within each donor. (H) Median neighborhood COL fraction (0–1) at or below and above the α-SMA gate in set A. The 24 construct medians comprise six donors, two conditions and two α-SMA states. The α-SMA gate is the no-primary-control 95th percentile (0.4654). In (G) and (H), bars are equal-donor arithmetic means and lines connect paired donor values between conditions. Bars in (E), (G) and (H) have no error bars.

Fig. S11 (continued; 2 / 3)

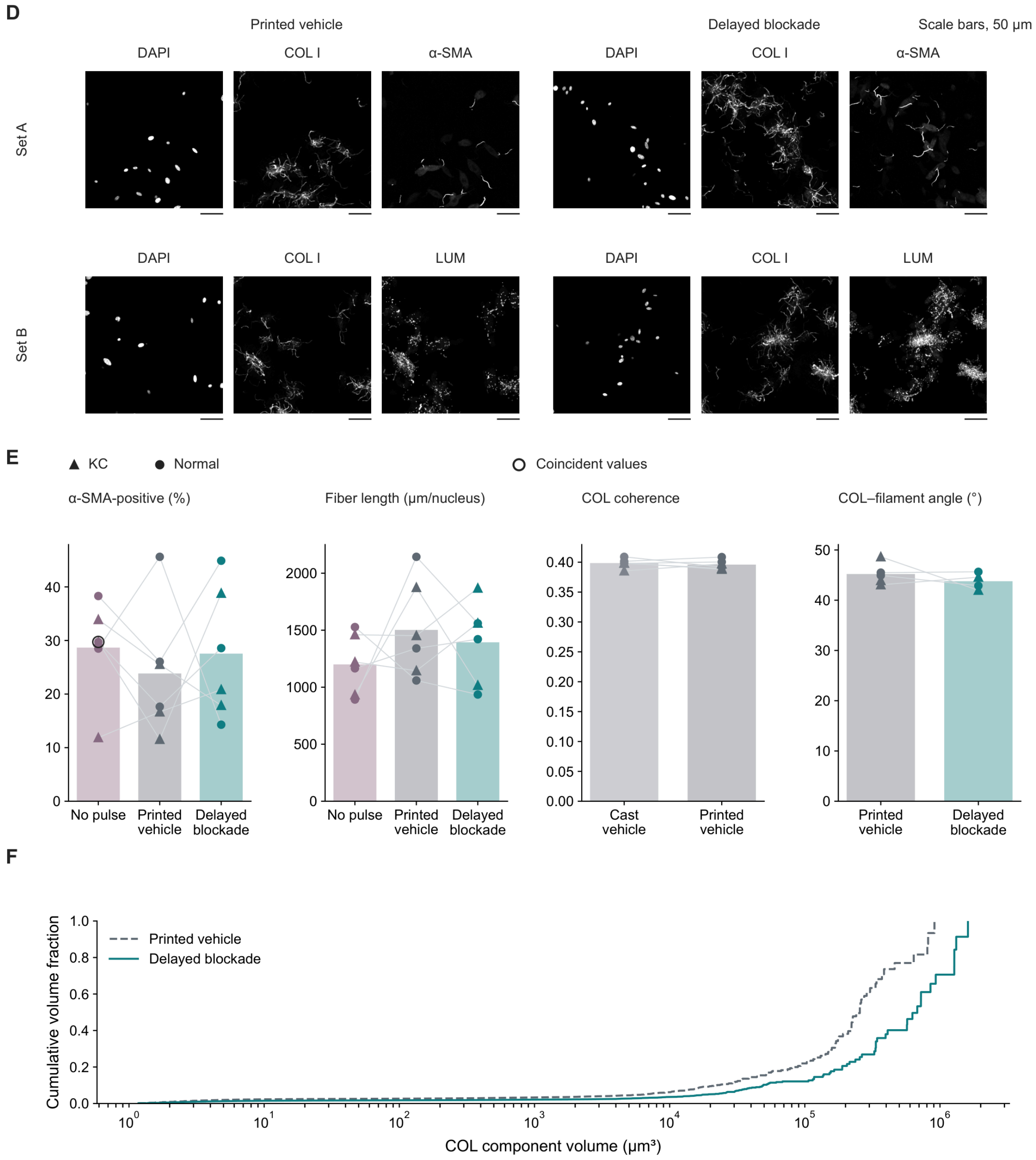

Fig. S11 (continued; 3 / 3)

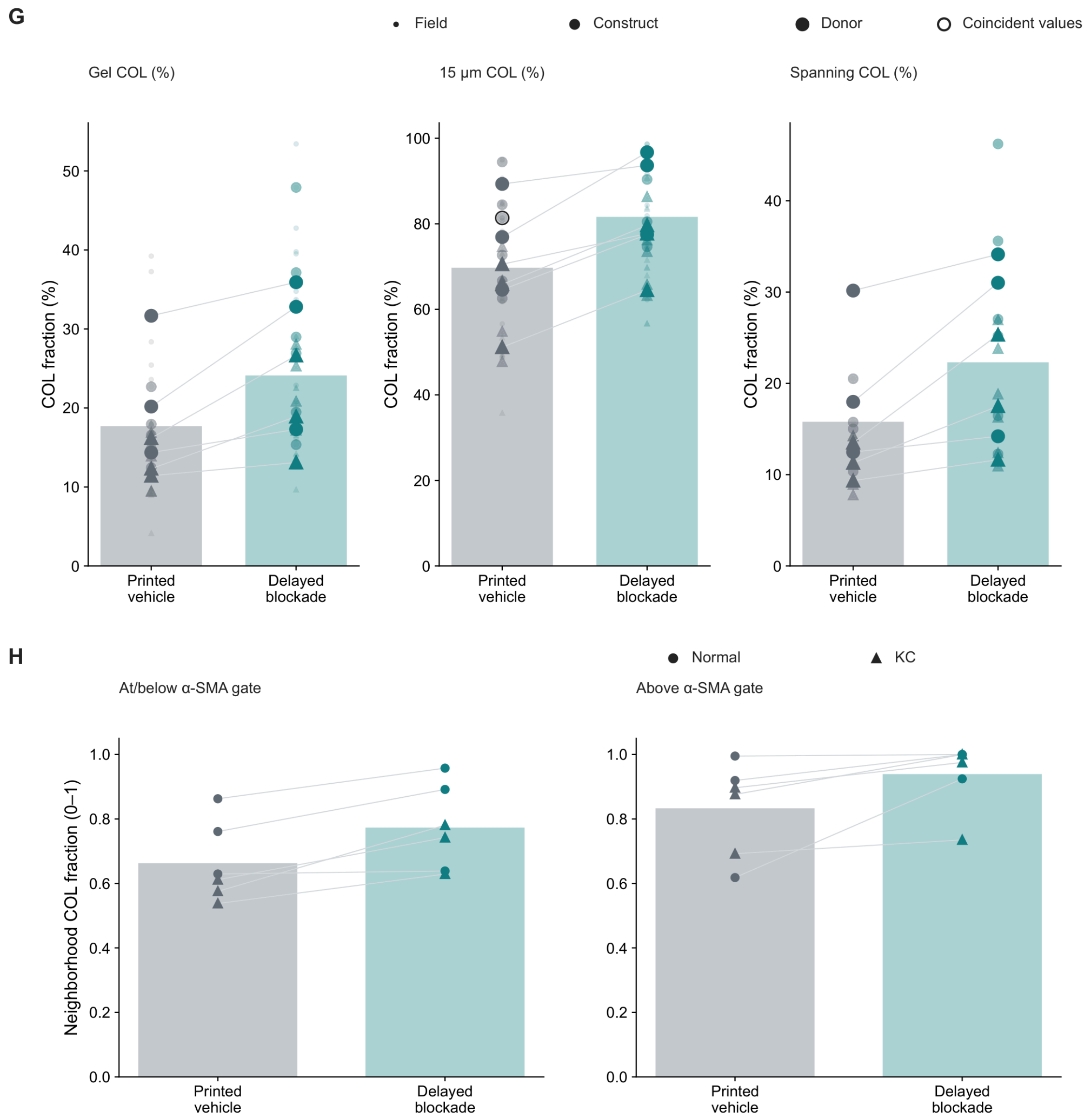

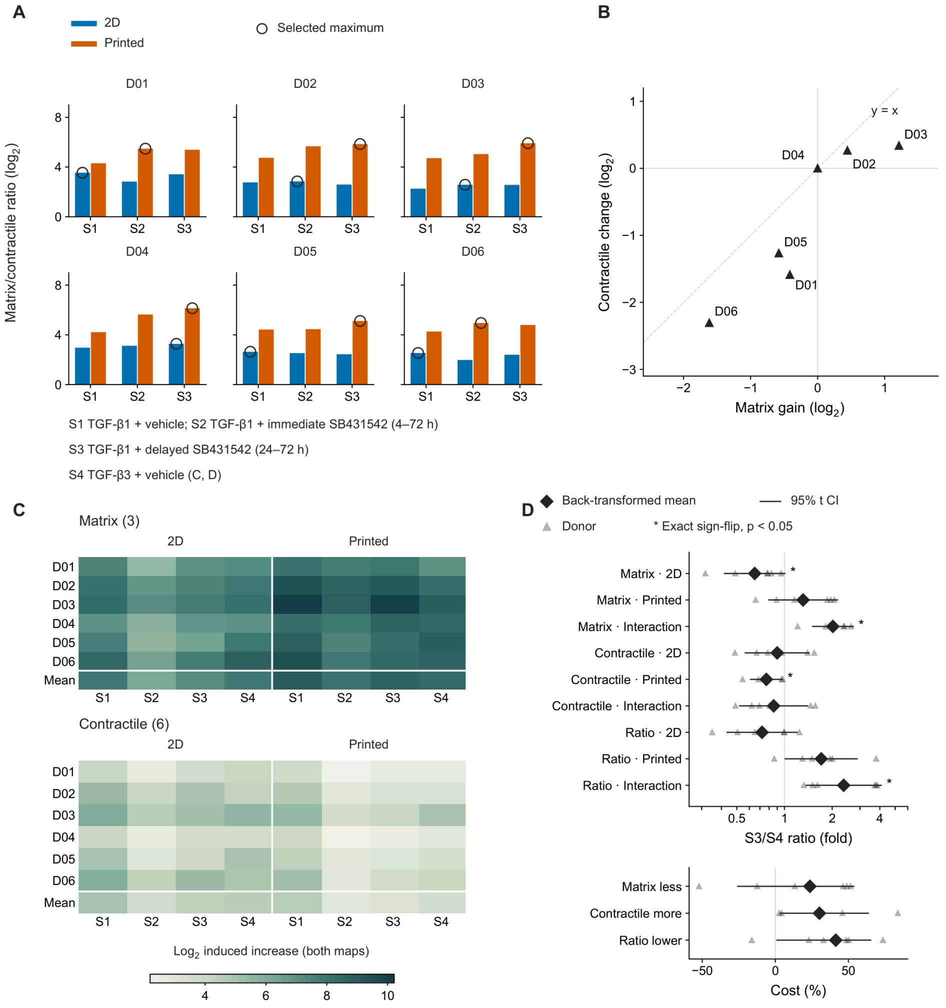


**Fig. S12. Regimen selection and matrix and contractile responses in keratoconus donor cultures.** (A) Matrix-to-contractile response ratios ($\log_2$) for three transforming growth factor-β1 (TGF-β1) regimens in six paired keratoconus (KC) donors, D01–D06. The 36 bars comprise one value per donor, regimen and culture format, with S1–S3 compared in two-dimensional (2D) and printed cultures. Regimens S1–S4 are those of Fig. 7A. The matrix-to-contractile response ratio ($\log_2$) is the difference between the $\log_2$ TGF-β1-induced increases in matrix genes (FN1, COL1A1 and COL5A1) and contractile genes (ACTA2, TAGLN, TPM1, CCN2, LOX and CCN1). TGF-β1-induced increases are pulse-minus-matched-vehicle abundances at 72 h. Open circles mark the selected maximum in each format. (B) Donor-specific $\log_2$ matrix gene gain and $\log_2$ contractile gene change when the regimen maximizing the matrix-to-contractile response ratio in printed culture replaces the regimen maximizing that ratio in 2D, with both regimens evaluated in printed constructs. Each of the six points is one donor. Both axes use the same physical scale per $\log_2$ unit, and the dashed diagonal, y = x, marks no change in the matrix-to-contractile response ratio. D04 selected delayed SB431542 in both formats and lies at the origin. (C) $\text{Log}_2$ induced increases in matrix and contractile genes under S1–S4 in 2D and printed cultures. Each heatmap has six donor rows and a Mean row across eight format–regimen columns, giving 96 donor cells and 16 group-mean cells across both maps. The matrix and contractile sets contain three and six genes, respectively, and share one color scale spanning 2.1896962326642253–10.217455011346456. Mean rows contain the group means of the $\log_2$ induced increases. (D) Paired-donor estimates comparing S3 with S4 in the six KC donors. The upper block gives nine S3/S4 ratios of matrix-gene induction, contractile-gene induction and their response ratio (u), each in 2D, in printed culture and for the regimen-by-format interaction, defined as printed S3/S4 divided by 2D S3/S4. The ratio axis is logarithmic with a vertical reference at 1. Triangles are individual donor ratios on the same axis, and diamonds are back-transformed mean $\log_2$ ratios with 95% t confidence intervals. The lower block gives, in printed culture, the percentage by which using the 2D-selected regimen (S4) instead of the printed-selected regimen (S3) lowers matrix-gene induction, raises contractile-gene induction and lowers the response ratio, on an axis with reference 0. Triangles are individual donor costs, and diamonds and horizontal intervals are group estimates and 95% t confidence intervals obtained by transforming the paired-donor $\log_2$ estimates and their intervals. The four asterisks denote the matrix S3/S4 comparison in 2D, the matrix regimen-by-format interaction, the contractile S3/S4 comparison in printed culture and the u regimen-by-format interaction, each from a two-sided paired exact sign-flip test across six donors (exact $p = 0.03125$, $p < 0.05$). (E) Changes in matrix and contractile responses when the regimen selected in printed culture replaces the regimen selected in 2D, with both choices evaluated in printed cultures. Each of the six donor plots contains matrix and contractile $\log_2$ changes, with the heading giving the regimen selected in 2D followed by that selected in printed culture among S1–S4. D05 selects S4 in both formats and has zero change in both responses.

Fig. S12 (continued; 2 / 2)

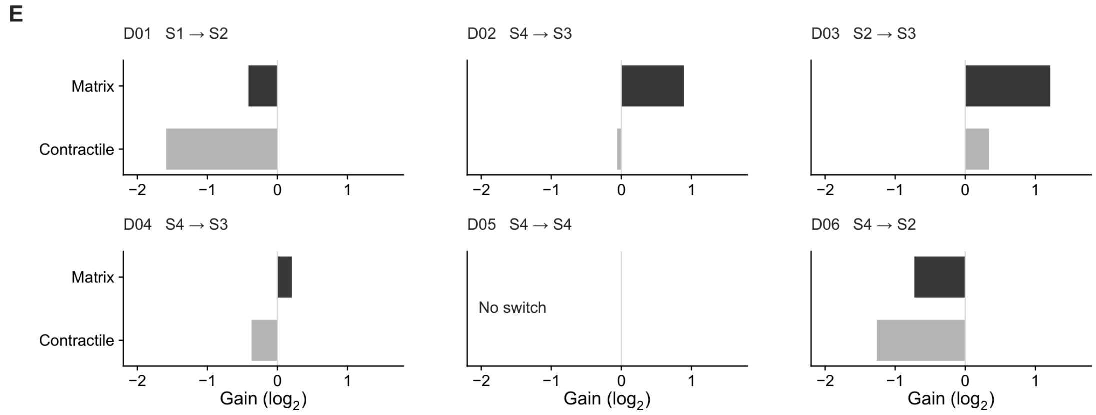

### Separate supplementary files

**Supplementary Data 1 — Supplementary_Data_1.pdf**
Lactate dehydrogenase release and assay calibrations in the functional batch.

**Supplementary Data 2 — Supplementary_Data_2.pdf**
Lumican amplification, melt curves and gene-specific standard curves.

**Supplementary Data 3 — Supplementary_Data_3.pdf**
Full membranes and lane assignments.

**Supplementary Data 4 — Supplementary_Data_4.pdf**
Whole-mount confocal images at 336 h after pulse onset (T336).

**Table S1 — Supplementary_Table_1.xlsx**
Transcriptome gene and gene-set effects and fixed disease-state programme weights for reuse of the RNA-sequencing comparisons.

**Table S2 — Supplementary_Table_2.xlsx**
Structural genetic signals, allele-oriented variant effects and gene expression responses in printed constructs and 2D cultures.

**Table S3 — Supplementary_Table_3.xlsx**
In vivo repair-state, in vivo fibrosis and independent three-dimensional reference scores.

**Table S4 — Supplementary_Table_4.xlsx**
Donor-level intervention and endpoint choices and their matrix, contractile and native collagen content consequences, and the coded outcomes of published corneal pathway-inhibition studies.

**Table S4 companion files — Supplementary_Table_4_companion_files.zip**
Study and endpoint coding of the KC-treatment evaluation (191 screened records, 2,128 endpoint records), deterministic summary, source records and codebook.

**Table S5 — Supplementary_Table_5.xlsx**
qPCR primer sequences and RefSeq accessions identify the recommended and backup assays for functional and post-withdrawal experiments.

**Table S6 — Supplementary_Table_6.xlsx**
Antibodies, reagents and kits document the protocol materials for protein assays, imaging and post-withdrawal experiments.

**Table S7 — Supplementary_Table_7.xlsx**
Experimental designs and donor, construct and sample identities for the functional, post-withdrawal, imaging and RNA-seq experiments.

**Table S8 — Supplementary_Table_8.xlsx**
Western blot filenames, lane assignments and exposure frames, plate maps and OIR acquisition metadata.

**Source Data — Source_Data.xlsx**
Source observations and associated summaries for the figure panels (Fig. 1A–Fig. S12E), with 116 panel data worksheets and README, Dictionary and Codes worksheets (119 worksheets in total).
