## Supplementary Data 1 for "Material context and TGF-β blockade timing determine matrix retained after withdrawal in patient-derived GelMA-bioprinted keratoconus stroma"

**A**

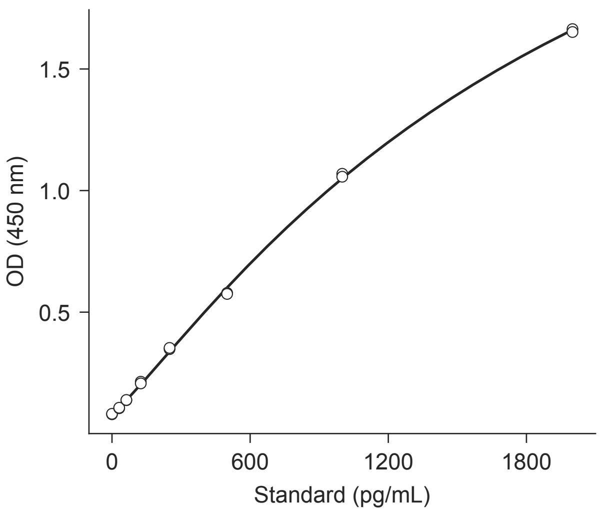


**B**

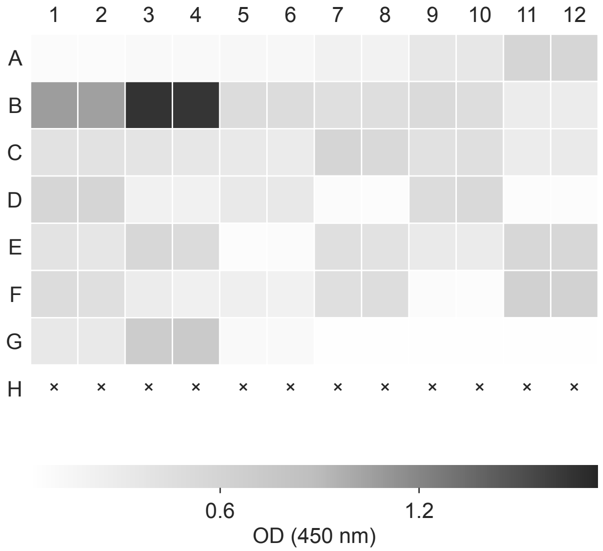


**C**

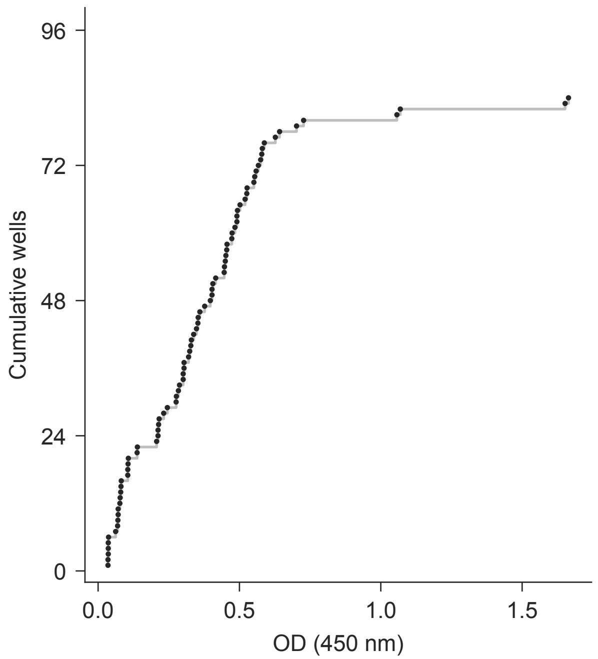


**D** elisa/plate_map.csv

| Well | Map rows | Dil. | Well | Map rows | Dil. |
|---|---|---|---|---|---|
| A1-2 | 1154-1155 | — | D7-8 | 1196-1197 | 1 |
| A3-4 | 1156-1157 | — | D9-10 | 1198-1199 | 1 |
| A5-6 | 1158-1159 | — | D11-12 | 1200-1201 | 1 |
| A7-8 | 1160-1161 | — | E1-2 | 1202-1203 | 1 |
| A9-10 | 1162-1163 | — | E3-4 | 1204-1205 | 1 |
| A11-12 | 1164-1165 | — | E5-6 | 1206-1207 | 1 |
| B1-2 | 1166-1167 | — | E7-8 | 1208-1209 | 1 |
| B3-4 | 1168-1169 | — | E9-10 | 1210-1211 | 1 |
| B5-8 | 1170-1173 | — | E11-12 | 1212-1213 | 1 |
| B9-10 | 1174-1175 | 1 | F1-2 | 1214-1215 | 1 |
| B11-12 | 1176-1177 | 1 | F3-4 | 1216-1217 | 1 |
| C1-2 | 1178-1179 | 1 | F5-6 | 1218-1219 | 1 |
| C3-4 | 1180-1181 | 1 | F7-8 | 1220-1221 | 1 |
| C5-6 | 1182-1183 | 1 | F9-10 | 1222-1223 | 1 |
| C7-8 | 1184-1185 | 1 | F11-12 | 1224-1225 | 1 |
| C9-10 | 1186-1187 | 1 | G1-2 | 1226-1227 | 1 |
| C11-12 | 1188-1189 | 1 | G3-4 | 1228-1229 | 1 |
| D1-2 | 1190-1191 | 1 | G5-12 | 1230-1237 | — |
| D3-4 | 1192-1193 | 1 | H1-12 | 1238-1249 | — |
| D5-6 | 1194-1195 | 1 | | | |

**A**

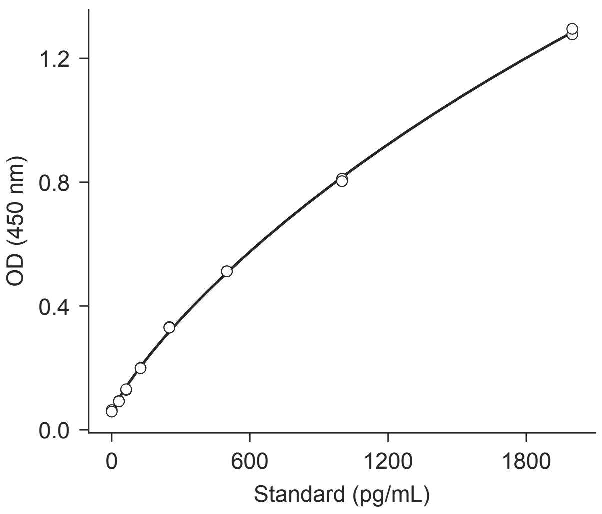


**B**

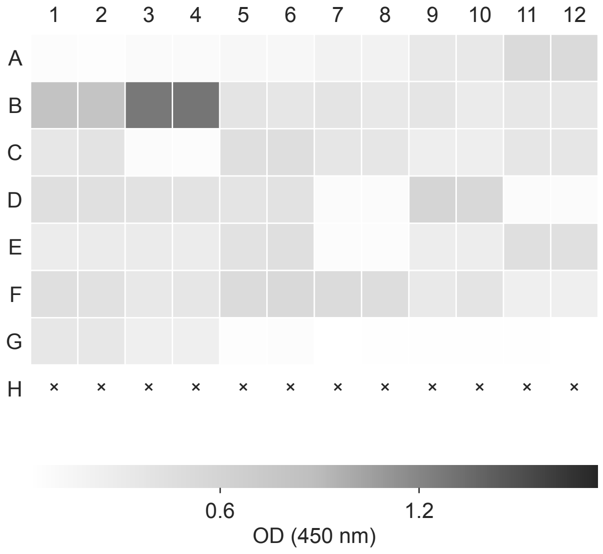


**C**

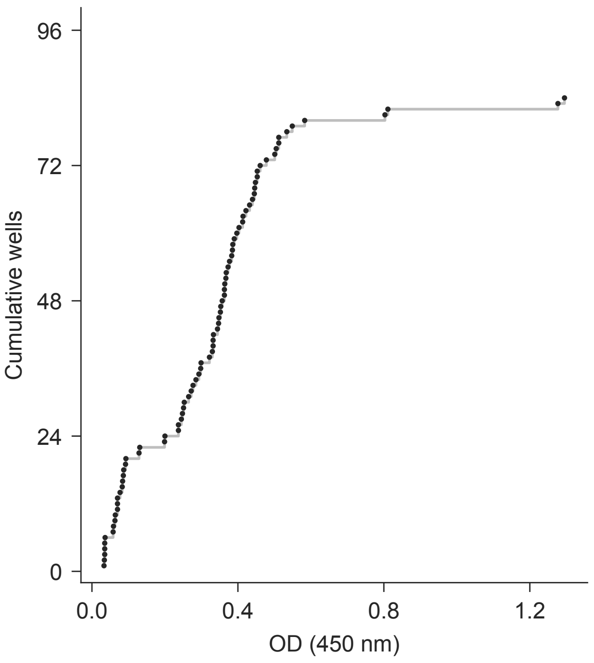


**D** elisa/plate_map.csv

| Well | Map rows | Dil. | Well | Map rows | Dil. |
|---|---|---|---|---|---|
| A1-2 | 1250-1251 | — | D7-8 | 1292-1293 | 1 |
| A3-4 | 1252-1253 | — | D9-10 | 1294-1295 | 1 |
| A5-6 | 1254-1255 | — | D11-12 | 1296-1297 | 1 |
| A7-8 | 1256-1257 | — | E1-2 | 1298-1299 | 1 |
| A9-10 | 1258-1259 | — | E3-4 | 1300-1301 | 1 |
| A11-12 | 1260-1261 | — | E5-6 | 1302-1303 | 1 |
| B1-2 | 1262-1263 | — | E7-8 | 1304-1305 | 1 |
| B3-4 | 1264-1265 | — | E9-10 | 1306-1307 | 1 |
| B5-8 | 1266-1269 | — | E11-12 | 1308-1309 | 1 |
| B9-10 | 1270-1271 | 1 | F1-2 | 1310-1311 | 1 |
| B11-12 | 1272-1273 | 1 | F3-4 | 1312-1313 | 1 |
| C1-2 | 1274-1275 | 1 | F5-6 | 1314-1315 | 1 |
| C3-4 | 1276-1277 | 1 | F7-8 | 1316-1317 | 1 |
| C5-6 | 1278-1279 | 1 | F9-10 | 1318-1319 | 1 |
| C7-8 | 1280-1281 | 1 | F11-12 | 1320-1321 | 1 |
| C9-10 | 1282-1283 | 1 | G1-2 | 1322-1323 | 1 |
| C11-12 | 1284-1285 | 1 | G3-4 | 1324-1325 | 1 |
| D1-2 | 1286-1287 | 1 | G5-12 | 1326-1333 | — |
| D3-4 | 1288-1289 | 1 | H1-12 | 1334-1345 | — |
| D5-6 | 1290-1291 | 1 | | | |

**A**

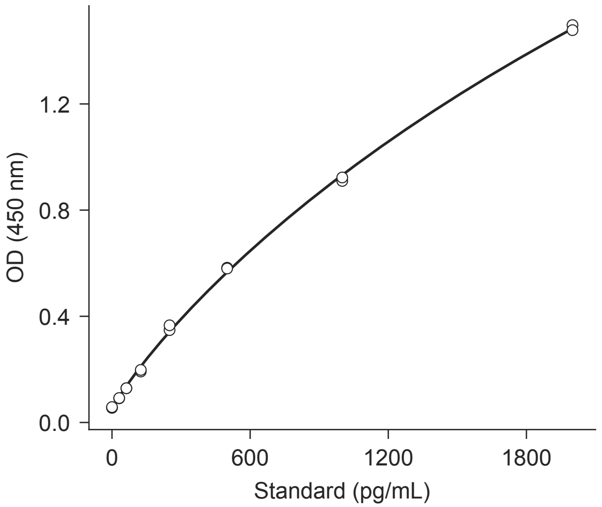


**B**

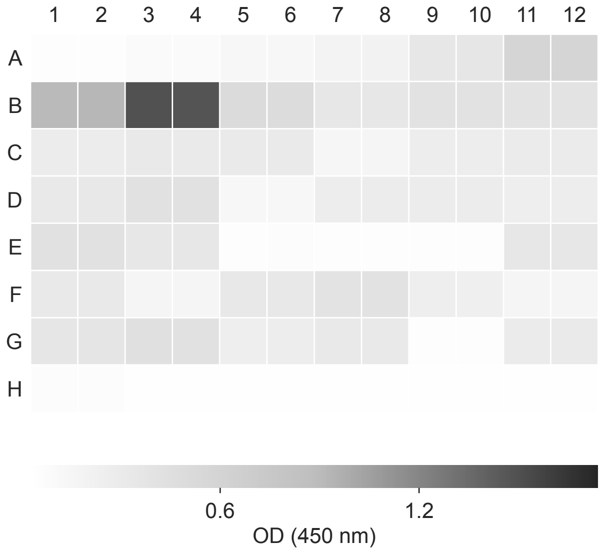


**C**

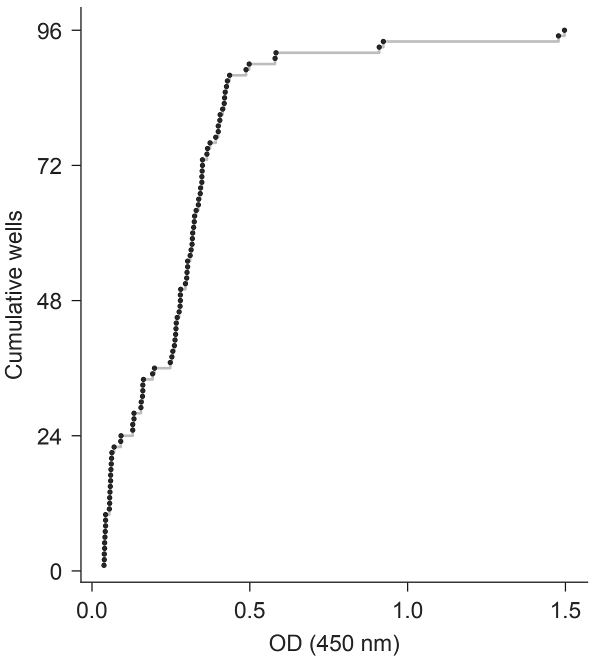


**D** elisa/plate_map.csv

| Well | Map rows | Dil. | Well | Map rows | Dil. |
|---|---|---|---|---|---|
| A1-2 | 2-3 | — | D9-10 | 46-47 | 1 |
| A3-4 | 4-5 | — | D11-12 | 48-49 | 1 |
| A5-6 | 6-7 | — | E1-2 | 50-51 | 1 |
| A7-8 | 8-9 | — | E3-4 | 52-53 | 1 |
| A9-10 | 10-11 | — | E5-6 | 54-55 | 1 |
| A11-12 | 12-13 | — | E7-8 | 56-57 | 1 |
| B1-2 | 14-15 | — | E9-10 | 58-59 | 1 |
| B3-4 | 16-17 | — | E11-12 | 60-61 | 1 |
| B5-6 | 18-19 | 1 | F1-2 | 62-63 | 1 |
| B7-8 | 20-21 | — | F3-4 | 64-65 | 1 |
| B9-10 | 22-23 | 1 | F5-6 | 66-67 | 1 |
| B11-12 | 24-25 | 1 | F7-8 | 68-69 | 1 |
| C1-2 | 26-27 | 1 | F9-10 | 70-71 | 1 |
| C3-4 | 28-29 | 1 | F11-12 | 72-73 | 1 |
| C5-6 | 30-31 | 1 | G1-2 | 74-75 | 1 |
| C7-8 | 32-33 | 1 | G3-4 | 76-77 | 1 |
| C9-10 | 34-35 | 1 | G5-6 | 78-79 | 1 |
| C11-12 | 36-37 | 1 | G7-8 | 80-81 | 1 |
| D1-2 | 38-39 | 1 | G9-10 | 82-83 | 1 |
| D3-4 | 40-41 | 1 | G11-12 | 84-85 | 1 |
| D5-6 | 42-43 | 1 | H1-12 | 86-97 | — |
| D7-8 | 44-45 | 1 | | | |

A

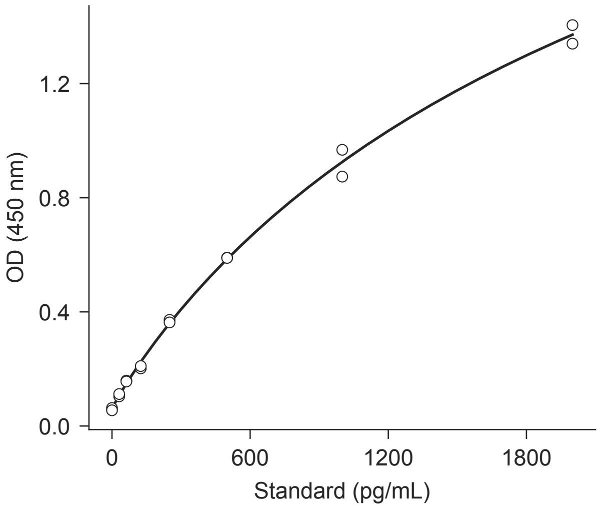


B

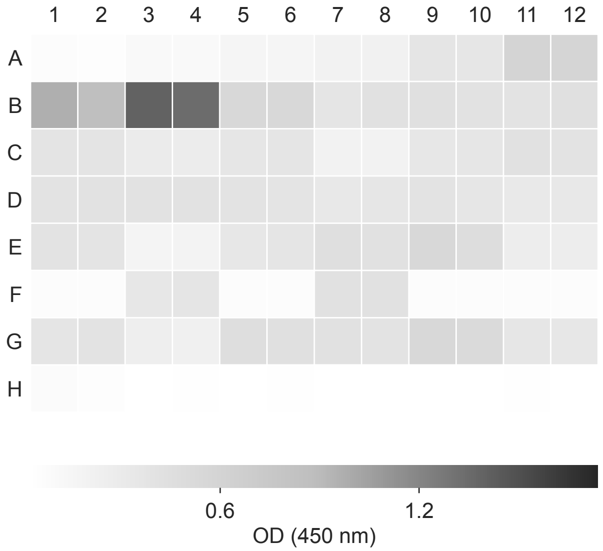


C

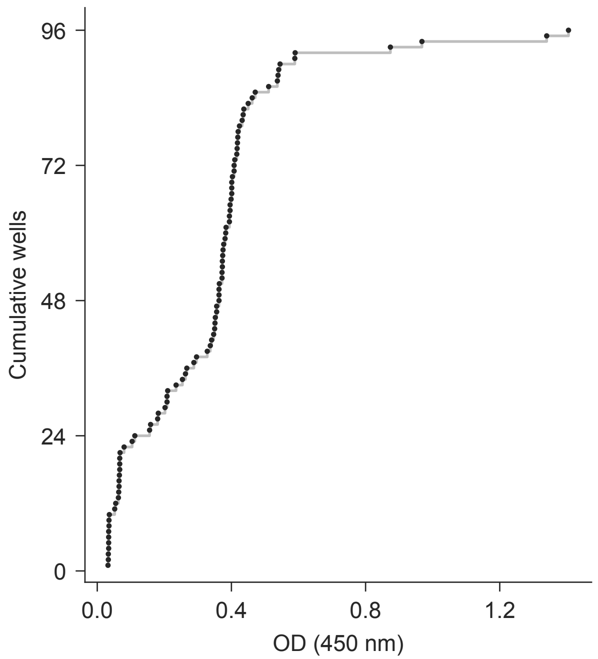


D

elisa/plate_map.csv

| Well | Map rows | Dil. | Well | Map rows | Dil. |
|---|---|---|---|---|---|
| A1-2 | 98-99 | — | D9-10 | 142-143 | 1 |
| A3-4 | 100-101 | — | D11-12 | 144-145 | 1 |
| A5-6 | 102-103 | — | E1-2 | 146-147 | 1 |
| A7-8 | 104-105 | — | E3-4 | 148-149 | 1 |
| A9-10 | 106-107 | — | E5-6 | 150-151 | 1 |
| A11-12 | 108-109 | — | E7-8 | 152-153 | 1 |
| B1-2 | 110-111 | — | E9-10 | 154-155 | 1 |
| B3-4 | 112-113 | — | E11-12 | 156-157 | 1 |
| B5-6 | 114-115 | 1 | F1-2 | 158-159 | 1 |
| B7-8 | 116-117 | — | F3-4 | 160-161 | 1 |
| B9-10 | 118-119 | 1 | F5-6 | 162-163 | 1 |
| B11-12 | 120-121 | 1 | F7-8 | 164-165 | 1 |
| C1-2 | 122-123 | 1 | F9-10 | 166-167 | 1 |
| C3-4 | 124-125 | 1 | F11-12 | 168-169 | 1 |
| C5-6 | 126-127 | 1 | G1-2 | 170-171 | 1 |
| C7-8 | 128-129 | 1 | G3-4 | 172-173 | 1 |
| C9-10 | 130-131 | 1 | G5-6 | 174-175 | 1 |
| C11-12 | 132-133 | 1 | G7-8 | 176-177 | 1 |
| D1-2 | 134-135 | 1 | G9-10 | 178-179 | 1 |
| D3-4 | 136-137 | 1 | G11-12 | 180-181 | 1 |
| D5-6 | 138-139 | 1 | H1-12 | 182-193 | — |
| D7-8 | 140-141 | 1 | | | |

**A**

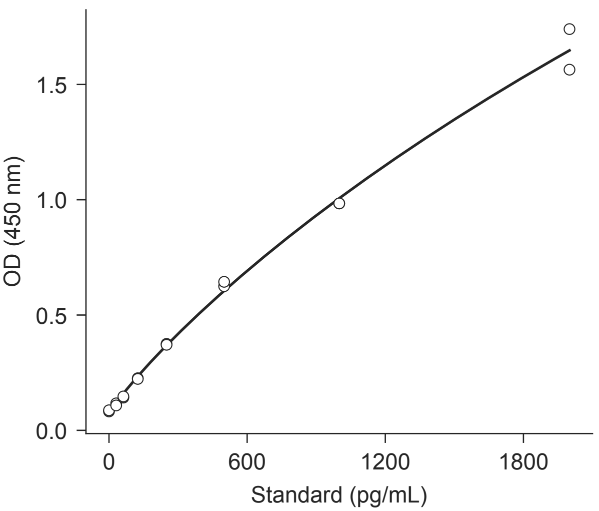


**B**

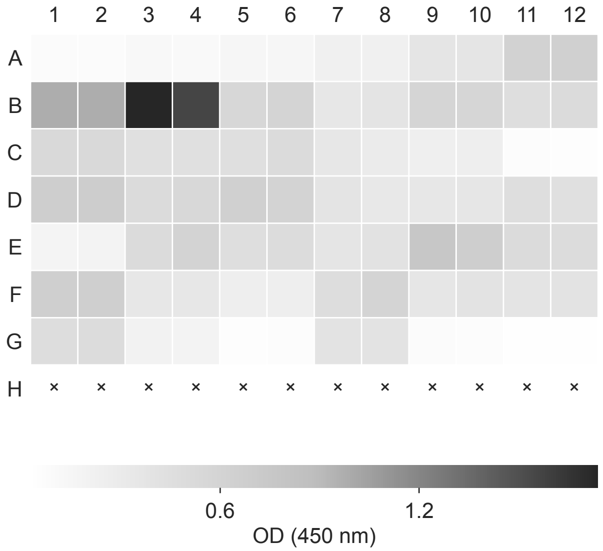


**C**

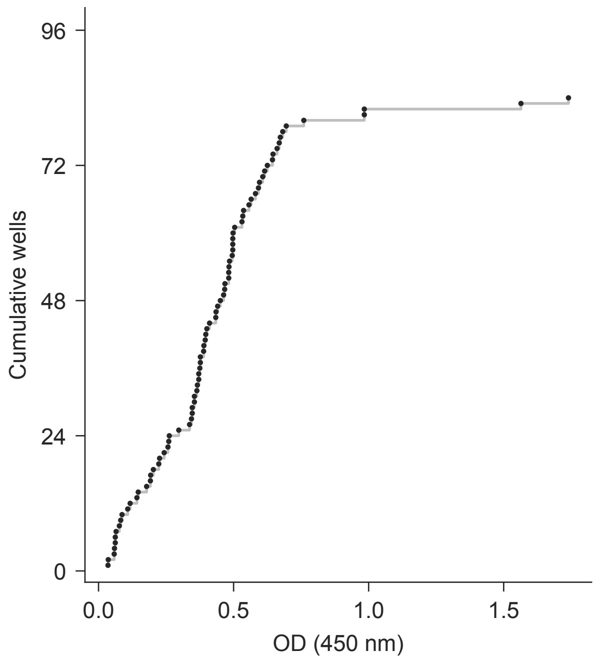


**D** elisa/plate_map.csv

| Well | Map rows | Dil. | Well | Map rows | Dil. |
|---|---|---|---|---|---|
| A1-2 | 194-195 | — | D7-8 | 236-237 | 1 |
| A3-4 | 196-197 | — | D9-10 | 238-239 | 1 |
| A5-6 | 198-199 | — | D11-12 | 240-241 | 1 |
| A7-8 | 200-201 | — | E1-2 | 242-243 | 1 |
| A9-10 | 202-203 | — | E3-4 | 244-245 | 1 |
| A11-12 | 204-205 | — | E5-6 | 246-247 | 1 |
| B1-2 | 206-207 | — | E7-8 | 248-249 | 1 |
| B3-4 | 208-209 | — | E9-10 | 250-251 | 1 |
| B5-6 | 210-211 | 1 | E11-12 | 252-253 | 1 |
| B7-8 | 212-213 | — | F1-2 | 254-255 | 1 |
| B9-10 | 214-215 | 1 | F3-4 | 256-257 | 1 |
| B11-12 | 216-217 | 1 | F5-6 | 258-259 | 1 |
| C1-2 | 218-219 | 1 | F7-8 | 260-261 | 1 |
| C3-4 | 220-221 | 1 | F9-10 | 262-263 | 1 |
| C5-6 | 222-223 | 1 | F11-12 | 264-265 | 1 |
| C7-8 | 224-225 | 1 | G1-2 | 266-267 | 1 |
| C9-10 | 226-227 | 1 | G3-4 | 268-269 | 1 |
| C11-12 | 228-229 | 1 | G5-6 | 270-271 | 1 |
| D1-2 | 230-231 | 1 | G7-8 | 272-273 | 1 |
| D3-4 | 232-233 | 1 | G9-12 | 274-277 | — |
| D5-6 | 234-235 | 1 | H1-12 | 278-289 | — |

**A**

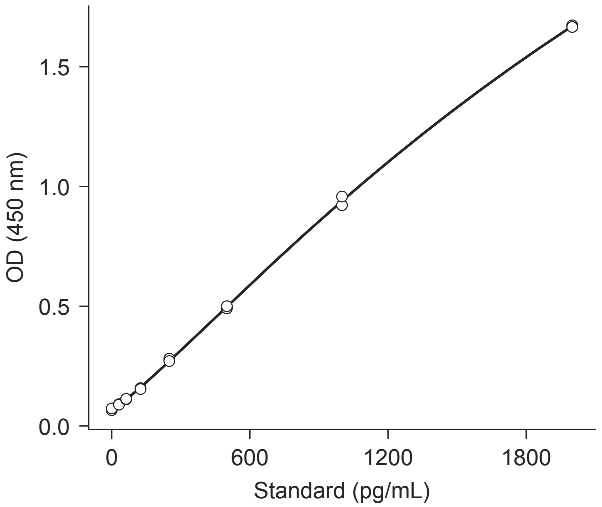


**B**

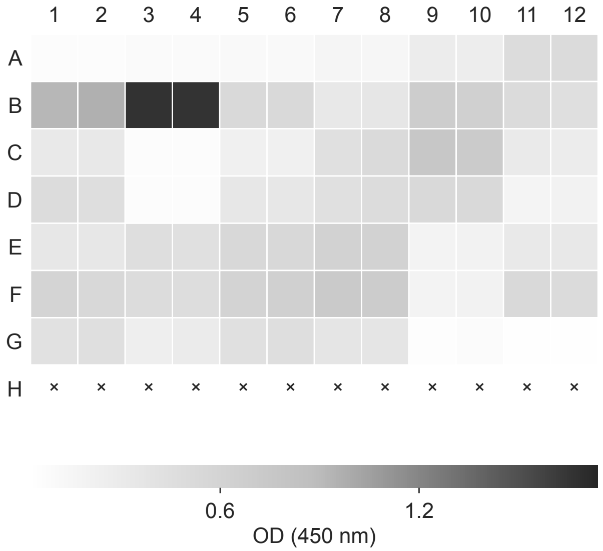


**C**

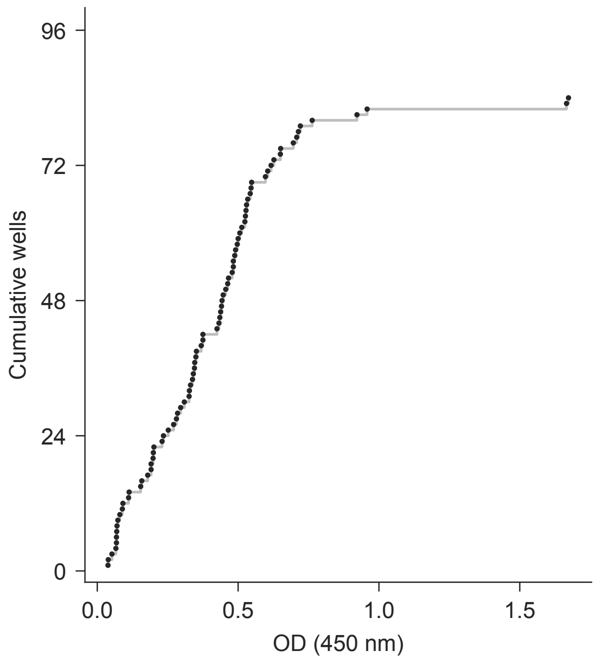


**D** elisa/plate_map.csv

| Well | Map rows | Dil. | Well | Map rows | Dil. |
|---|---|---|---|---|---|
| A1-2 | 290-291 | — | D7-8 | 332-333 | 1 |
| A3-4 | 292-293 | — | D9-10 | 334-335 | 1 |
| A5-6 | 294-295 | — | D11-12 | 336-337 | 1 |
| A7-8 | 296-297 | — | E1-2 | 338-339 | 1 |
| A9-10 | 298-299 | — | E3-4 | 340-341 | 1 |
| A11-12 | 300-301 | — | E5-6 | 342-343 | 1 |
| B1-2 | 302-303 | — | E7-8 | 344-345 | 1 |
| B3-4 | 304-305 | — | E9-10 | 346-347 | 1 |
| B5-6 | 306-307 | 1 | E11-12 | 348-349 | 1 |
| B7-8 | 308-309 | — | F1-2 | 350-351 | 1 |
| B9-10 | 310-311 | 1 | F3-4 | 352-353 | 1 |
| B11-12 | 312-313 | 1 | F5-6 | 354-355 | 1 |
| C1-2 | 314-315 | 1 | F7-8 | 356-357 | 1 |
| C3-4 | 316-317 | 1 | F9-10 | 358-359 | 1 |
| C5-6 | 318-319 | 1 | F11-12 | 360-361 | 1 |
| C7-8 | 320-321 | 1 | G1-2 | 362-363 | 1 |
| C9-10 | 322-323 | 1 | G3-4 | 364-365 | 1 |
| C11-12 | 324-325 | 1 | G5-6 | 366-367 | 1 |
| D1-2 | 326-327 | 1 | G7-8 | 368-369 | 1 |
| D3-4 | 328-329 | 1 | G9-12 | 370-373 | — |
| D5-6 | 330-331 | 1 | H1-12 | 374-385 | — |

A

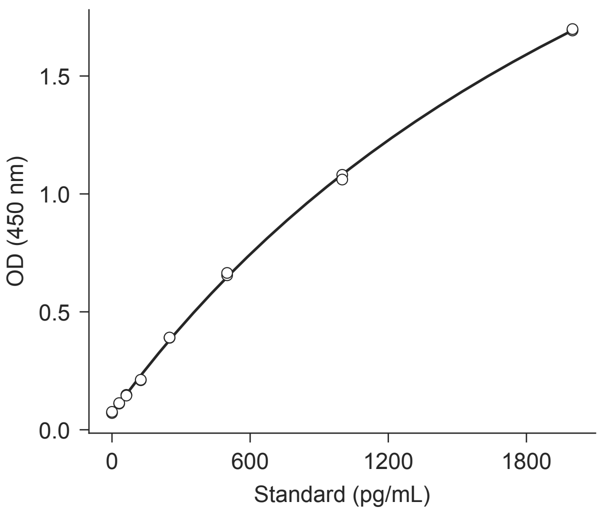


B

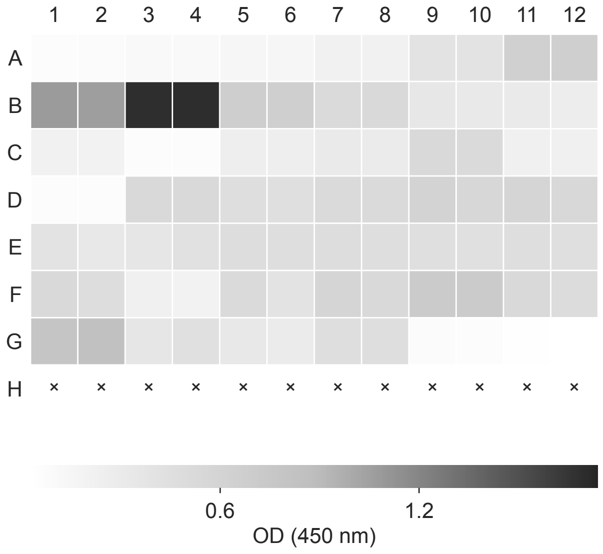


C

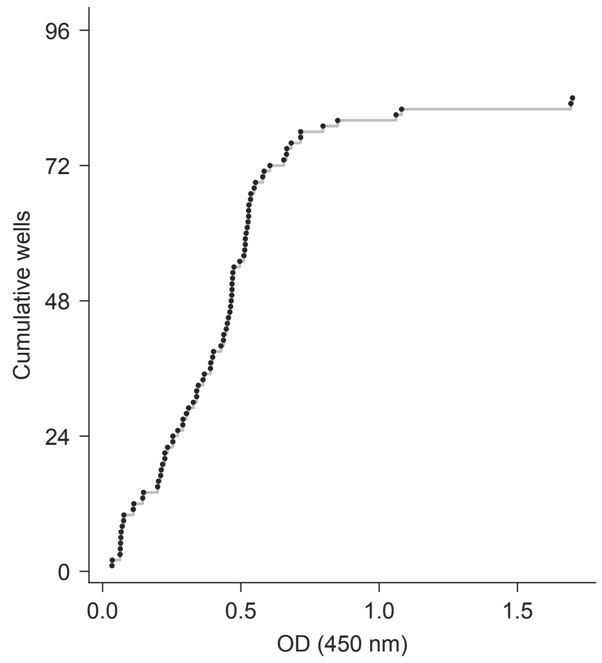


D elisa/plate_map.csv

| Well | Map rows | Dil. | Well | Map rows | Dil. |
|---|---|---|---|---|---|
| A1-2 | 386-387 | — | D7-8 | 428-429 | 1 |
| A3-4 | 388-389 | — | D9-10 | 430-431 | 1 |
| A5-6 | 390-391 | — | D11-12 | 432-433 | 1 |
| A7-8 | 392-393 | — | E1-2 | 434-435 | 1 |
| A9-10 | 394-395 | — | E3-4 | 436-437 | 1 |
| A11-12 | 396-397 | — | E5-6 | 438-439 | 1 |
| B1-2 | 398-399 | — | E7-8 | 440-441 | 1 |
| B3-4 | 400-401 | — | E9-10 | 442-443 | 1 |
| B5-6 | 402-403 | 1 | E11-12 | 444-445 | 1 |
| B7-8 | 404-405 | — | F1-2 | 446-447 | 1 |
| B9-10 | 406-407 | 1 | F3-4 | 448-449 | 1 |
| B11-12 | 408-409 | 1 | F5-6 | 450-451 | 1 |
| C1-2 | 410-411 | 1 | F7-8 | 452-453 | 1 |
| C3-4 | 412-413 | 1 | F9-10 | 454-455 | 1 |
| C5-6 | 414-415 | 1 | F11-12 | 456-457 | 1 |
| C7-8 | 416-417 | 1 | G1-2 | 458-459 | 1 |
| C9-10 | 418-419 | 1 | G3-4 | 460-461 | 1 |
| C11-12 | 420-421 | 1 | G5-6 | 462-463 | 1 |
| D1-2 | 422-423 | 1 | G7-8 | 464-465 | 1 |
| D3-4 | 424-425 | 1 | G9-12 | 466-469 | — |
| D5-6 | 426-427 | 1 | H1-12 | 470-481 | — |

**A**

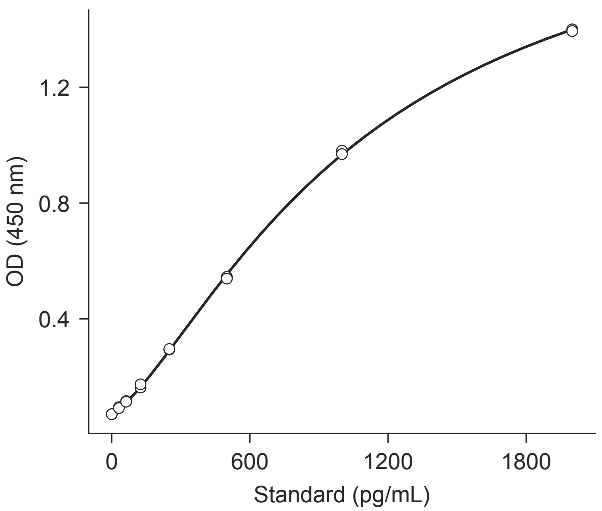


**B**

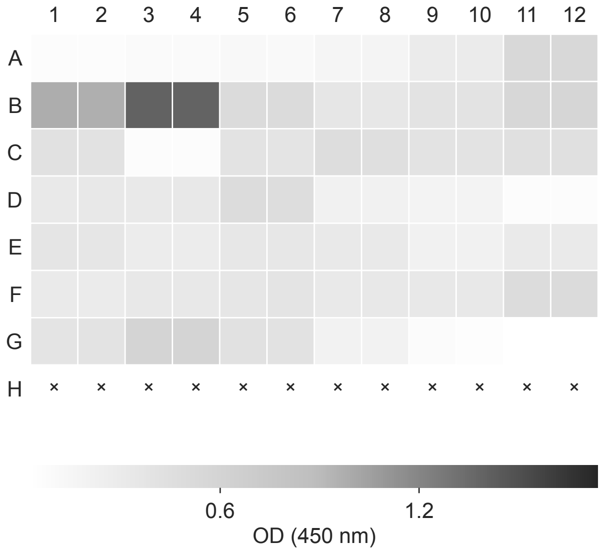


**C**

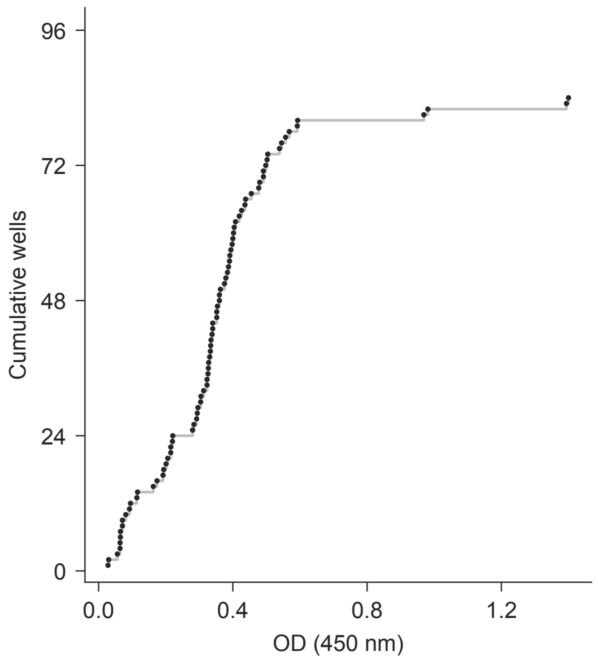


**D** elisa/plate_map.csv

| Well | Map rows | Dil. | Well | Map rows | Dil. |
|---|---|---|---|---|---|
| A1-2 | 482-483 | — | D7-8 | 524-525 | 1 |
| A3-4 | 484-485 | — | D9-10 | 526-527 | 1 |
| A5-6 | 486-487 | — | D11-12 | 528-529 | 1 |
| A7-8 | 488-489 | — | E1-2 | 530-531 | 1 |
| A9-10 | 490-491 | — | E3-4 | 532-533 | 1 |
| A11-12 | 492-493 | — | E5-6 | 534-535 | 1 |
| B1-2 | 494-495 | — | E7-8 | 536-537 | 1 |
| B3-4 | 496-497 | — | E9-10 | 538-539 | 1 |
| B5-6 | 498-499 | 1 | E11-12 | 540-541 | 1 |
| B7-8 | 500-501 | — | F1-2 | 542-543 | 1 |
| B9-10 | 502-503 | 1 | F3-4 | 544-545 | 1 |
| B11-12 | 504-505 | 1 | F5-6 | 546-547 | 1 |
| C1-2 | 506-507 | 1 | F7-8 | 548-549 | 1 |
| C3-4 | 508-509 | 1 | F9-10 | 550-551 | 1 |
| C5-6 | 510-511 | 1 | F11-12 | 552-553 | 1 |
| C7-8 | 512-513 | 1 | G1-2 | 554-555 | 1 |
| C9-10 | 514-515 | 1 | G3-4 | 556-557 | 1 |
| C11-12 | 516-517 | 1 | G5-6 | 558-559 | 1 |
| D1-2 | 518-519 | 1 | G7-8 | 560-561 | 1 |
| D3-4 | 520-521 | 1 | G9-12 | 562-565 | — |
| D5-6 | 522-523 | 1 | H1-12 | 566-577 | — |

**B**

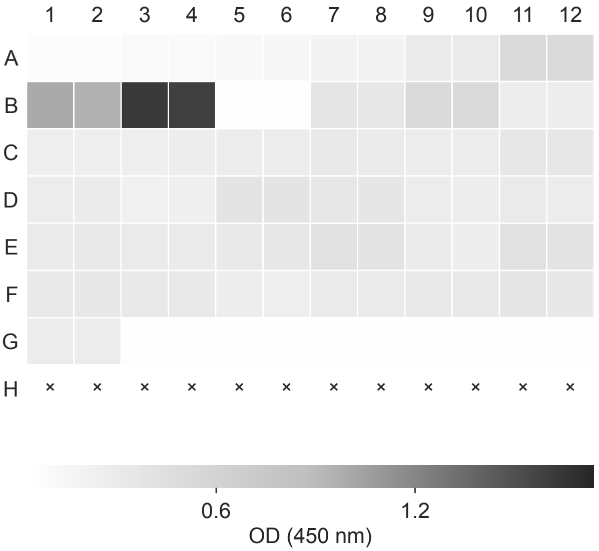


**C**

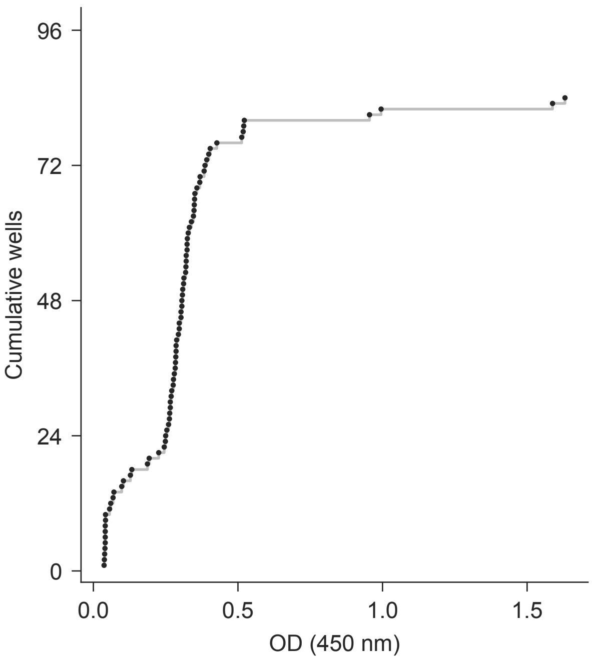


**D** OD (450 nm)

| Well | Value | Well | Value | Well | Value | Well | Value |
|---|---|---|---|---|---|---|---|
| A1 | 0.071 | C1 | 0.254 | E1 | 0.309 | G1 | 0.280 |
| A2 | 0.068 | C2 | 0.249 | E2 | 0.320 | G2 | 0.276 |
| A3 | 0.104 | C3 | 0.250 | E3 | 0.303 | G3 | 0.042 |
| A4 | 0.098 | C4 | 0.263 | E4 | 0.313 | G4 | 0.038 |
| A5 | 0.128 | C5 | 0.288 | E5 | 0.324 | G5 | 0.041 |
| A6 | 0.133 | C6 | 0.277 | E6 | 0.346 | G6 | 0.041 |
| A7 | 0.187 | C7 | 0.325 | E7 | 0.404 | G7 | 0.040 |
| A8 | 0.193 | C8 | 0.312 | E8 | 0.392 | G8 | 0.042 |
| A9 | 0.297 | C9 | 0.286 | E9 | 0.297 | G9 | 0.041 |
| A10 | 0.306 | C10 | 0.285 | E10 | 0.266 | G10 | 0.037 |
| A11 | 0.522 | C11 | 0.350 | E11 | 0.427 | G11 | 0.041 |
| A12 | 0.520 | C12 | 0.349 | E12 | 0.399 | G12 | 0.039 |
| B1 | 0.995 | D1 | 0.286 | F1 | 0.321 | H1 | — |
| B2 | 0.955 | D2 | 0.294 | F2 | 0.332 | H2 | — |
| B3 | 1.631 | D3 | 0.226 | F3 | 0.321 | H3 | — |
| B4 | 1.588 | D4 | 0.245 | F4 | 0.328 | H4 | — |
| B5 | 0.060 | D5 | 0.383 | F5 | 0.266 | H5 | — |
| B6 | 0.056 | D6 | 0.386 | F6 | 0.260 | H6 | — |
| B7 | 0.369 | D7 | 0.348 | F7 | 0.303 | H7 | — |
| B8 | 0.351 | D8 | 0.358 | F8 | 0.308 | H8 | — |
| B9 | 0.513 | D9 | 0.284 | F9 | 0.324 | H9 | — |
| B10 | 0.518 | D10 | 0.271 | F10 | 0.319 | H10 | — |
| B11 | 0.264 | D11 | 0.306 | F11 | 0.368 | H11 | — |
| B12 | 0.269 | D12 | 0.283 | F12 | 0.339 | H12 | — |

**A**

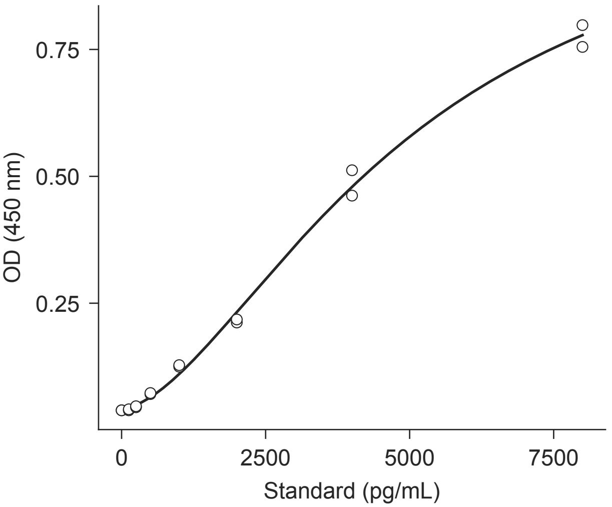


**B**

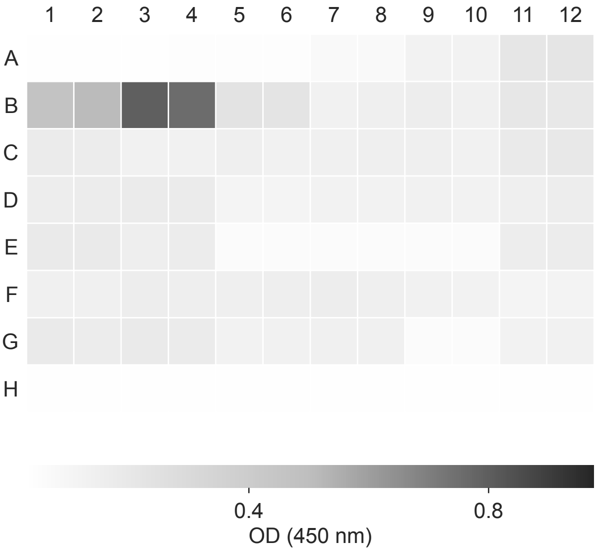


**C**

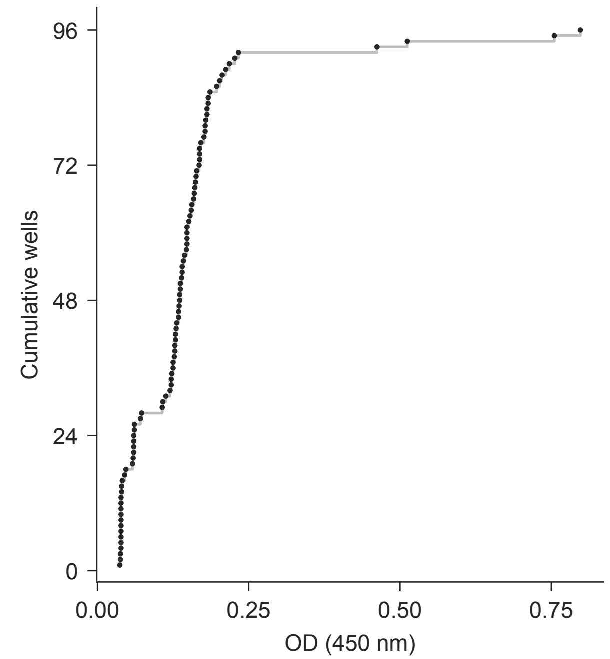


**D** elisa/plate_map.csv

| Well | Map rows | Dil. | Well | Map rows | Dil. |
|---|---|---|---|---|---|
| A1-2 | 578-579 | — | D9-10 | 622-623 | 1 |
| A3-4 | 580-581 | — | D11-12 | 624-625 | 1 |
| A5-6 | 582-583 | — | E1-2 | 626-627 | 1 |
| A7-8 | 584-585 | — | E3-4 | 628-629 | 1 |
| A9-10 | 586-587 | — | E5-6 | 630-631 | 1 |
| A11-12 | 588-589 | — | E7-8 | 632-633 | 1 |
| B1-2 | 590-591 | — | E9-10 | 634-635 | 1 |
| B3-4 | 592-593 | — | E11-12 | 636-637 | 1 |
| B5-6 | 594-595 | 1 | F1-2 | 638-639 | 1 |
| B7-8 | 596-597 | — | F3-4 | 640-641 | 1 |
| B9-10 | 598-599 | 1 | F5-6 | 642-643 | 1 |
| B11-12 | 600-601 | 1 | F7-8 | 644-645 | 1 |
| C1-2 | 602-603 | 1 | F9-10 | 646-647 | 1 |
| C3-4 | 604-605 | 1 | F11-12 | 648-649 | 1 |
| C5-6 | 606-607 | 1 | G1-2 | 650-651 | 1 |
| C7-8 | 608-609 | 1 | G3-4 | 652-653 | 1 |
| C9-10 | 610-611 | 1 | G5-6 | 654-655 | 1 |
| C11-12 | 612-613 | 1 | G7-8 | 656-657 | 1 |
| D1-2 | 614-615 | 1 | G9-10 | 658-659 | 1 |
| D3-4 | 616-617 | 1 | G11-12 | 660-661 | 1 |
| D5-6 | 618-619 | 1 | H1-12 | 662-673 | — |
| D7-8 | 620-621 | 1 | | | |

**A**

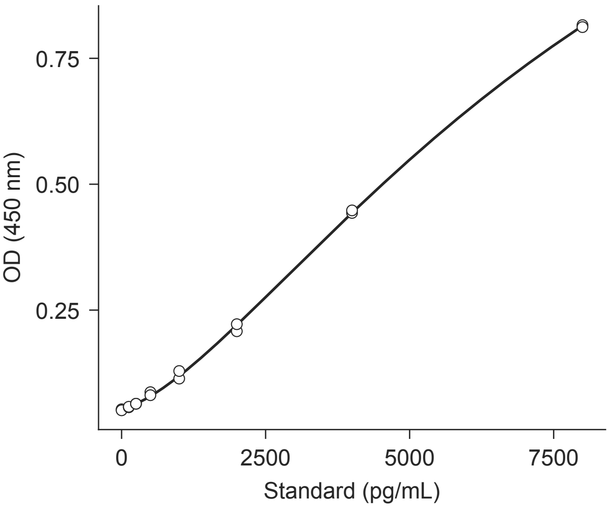


**B**

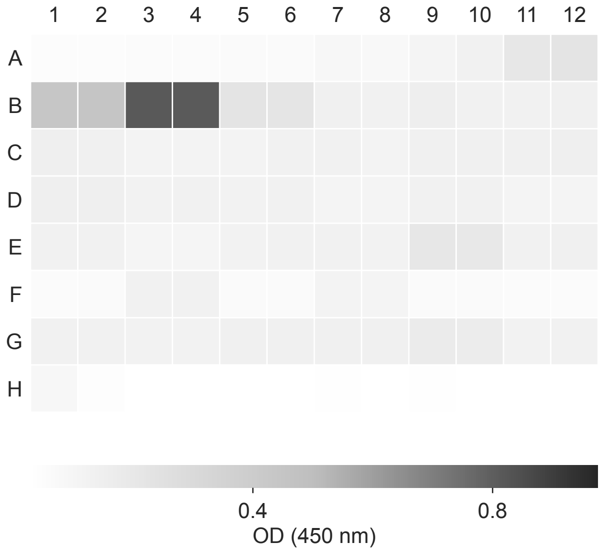


**C**

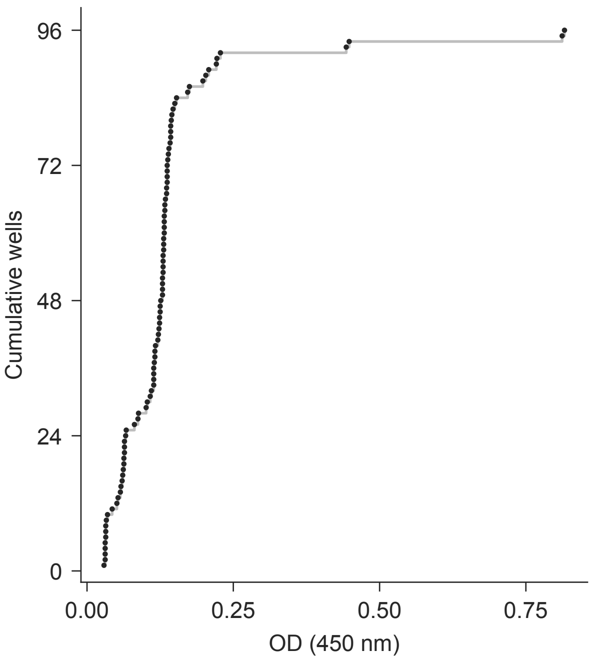


**D** elisa/plate_map.csv

| Well | Map rows | Dil. | Well | Map rows | Dil. |
|---|---|---|---|---|---|
| A1-2 | 674-675 | — | D9-10 | 718-719 | 1 |
| A3-4 | 676-677 | — | D11-12 | 720-721 | 1 |
| A5-6 | 678-679 | — | E1-2 | 722-723 | 1 |
| A7-8 | 680-681 | — | E3-4 | 724-725 | 1 |
| A9-10 | 682-683 | — | E5-6 | 726-727 | 1 |
| A11-12 | 684-685 | — | E7-8 | 728-729 | 1 |
| B1-2 | 686-687 | — | E9-10 | 730-731 | 1 |
| B3-4 | 688-689 | — | E11-12 | 732-733 | 1 |
| B5-6 | 690-691 | 1 | F1-2 | 734-735 | 1 |
| B7-8 | 692-693 | — | F3-4 | 736-737 | 1 |
| B9-10 | 694-695 | 1 | F5-6 | 738-739 | 1 |
| B11-12 | 696-697 | 1 | F7-8 | 740-741 | 1 |
| C1-2 | 698-699 | 1 | F9-10 | 742-743 | 1 |
| C3-4 | 700-701 | 1 | F11-12 | 744-745 | 1 |
| C5-6 | 702-703 | 1 | G1-2 | 746-747 | 1 |
| C7-8 | 704-705 | 1 | G3-4 | 748-749 | 1 |
| C9-10 | 706-707 | 1 | G5-6 | 750-751 | 1 |
| C11-12 | 708-709 | 1 | G7-8 | 752-753 | 1 |
| D1-2 | 710-711 | 1 | G9-10 | 754-755 | 1 |
| D3-4 | 712-713 | 1 | G11-12 | 756-757 | 1 |
| D5-6 | 714-715 | 1 | H1-12 | 758-769 | — |
| D7-8 | 716-717 | 1 | | | |

20260529_LUM_03.txt

Synergy H1 | 5/29/2026 | functional | LUM-03

**A**

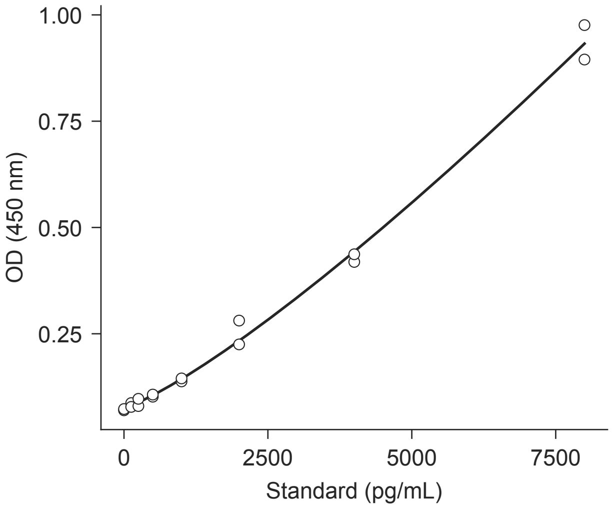


**B**

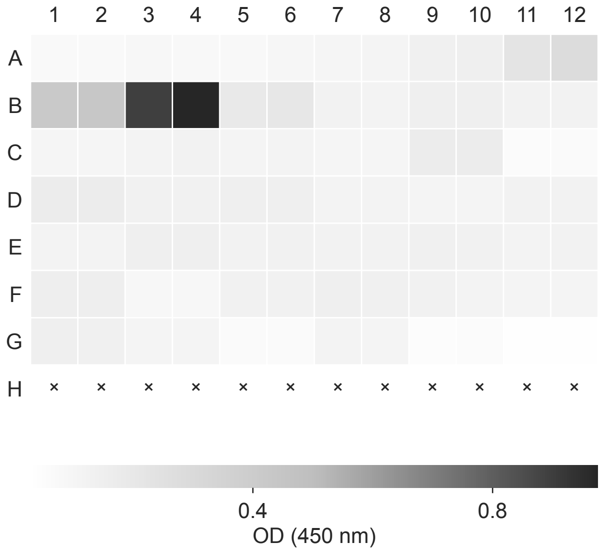


**C**

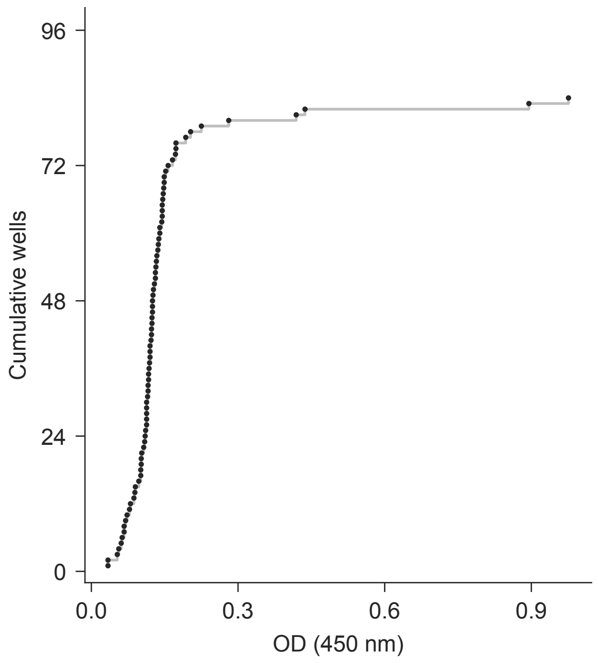


**D** elisa/plate_map.csv

| Well | Map rows | Dil. | Well | Map rows | Dil. |
|---|---|---|---|---|---|
| A1-2 | 770-771 | — | D7-8 | 812-813 | 1 |
| A3-4 | 772-773 | — | D9-10 | 814-815 | 1 |
| A5-6 | 774-775 | — | D11-12 | 816-817 | 1 |
| A7-8 | 776-777 | — | E1-2 | 818-819 | 1 |
| A9-10 | 778-779 | — | E3-4 | 820-821 | 1 |
| A11-12 | 780-781 | — | E5-6 | 822-823 | 1 |
| B1-2 | 782-783 | — | E7-8 | 824-825 | 1 |
| B3-4 | 784-785 | — | E9-10 | 826-827 | 1 |
| B5-6 | 786-787 | 1 | E11-12 | 828-829 | 1 |
| B7-8 | 788-789 | — | F1-2 | 830-831 | 1 |
| B9-10 | 790-791 | 1 | F3-4 | 832-833 | 1 |
| B11-12 | 792-793 | 1 | F5-6 | 834-835 | 1 |
| C1-2 | 794-795 | 1 | F7-8 | 836-837 | 1 |
| C3-4 | 796-797 | 1 | F9-10 | 838-839 | 1 |
| C5-6 | 798-799 | 1 | F11-12 | 840-841 | 1 |
| C7-8 | 800-801 | 1 | G1-2 | 842-843 | 1 |
| C9-10 | 802-803 | 1 | G3-4 | 844-845 | 1 |
| C11-12 | 804-805 | 1 | G5-6 | 846-847 | 1 |
| D1-2 | 806-807 | 1 | G7-8 | 848-849 | 1 |
| D3-4 | 808-809 | 1 | G9-12 | 850-853 | — |
| D5-6 | 810-811 | 1 | H1-12 | 854-865 | — |

**A**

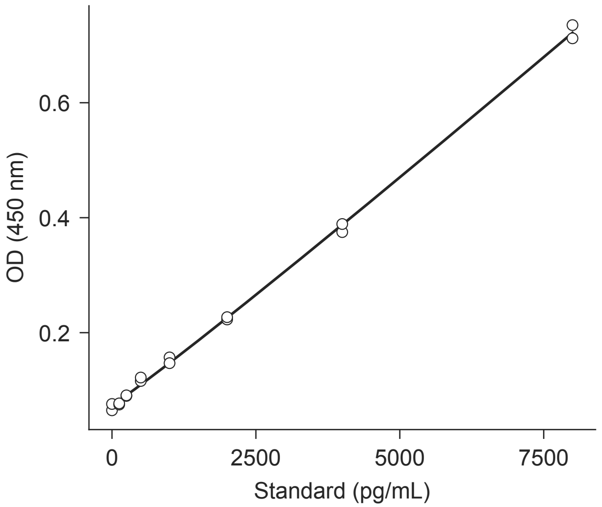


**B**

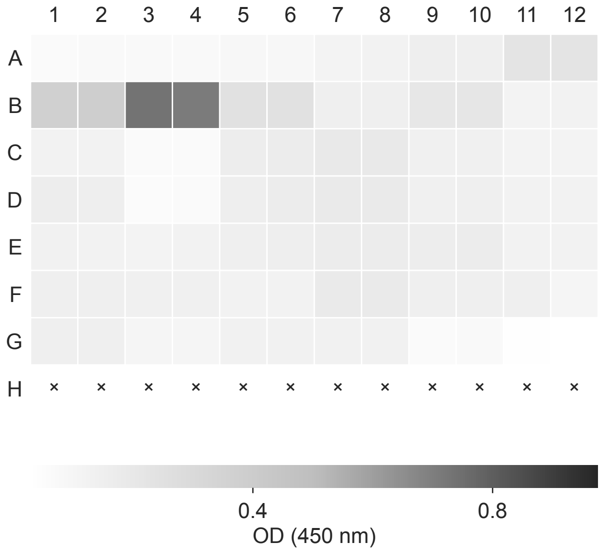


**C**

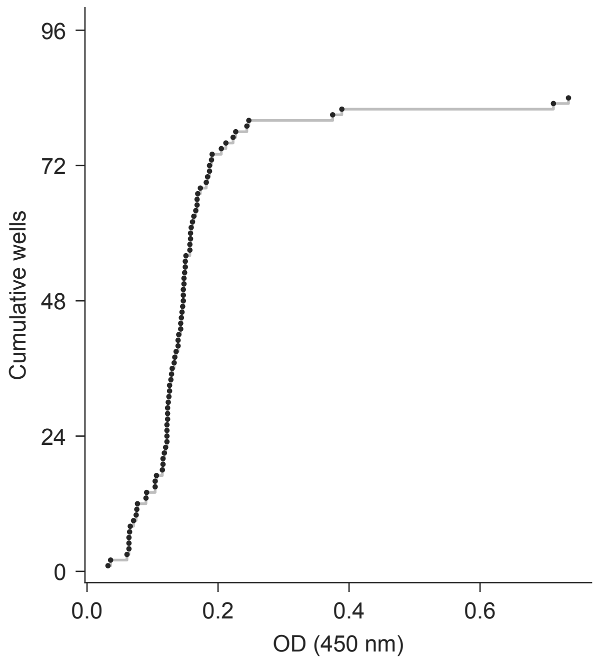


**D** elisa/plate_map.csv

| Well | Map rows | Dil. | Well | Map rows | Dil. |
|---|---|---|---|---|---|
| A1-2 | 866-867 | — | D7-8 | 908-909 | 1 |
| A3-4 | 868-869 | — | D9-10 | 910-911 | 1 |
| A5-6 | 870-871 | — | D11-12 | 912-913 | 1 |
| A7-8 | 872-873 | — | E1-2 | 914-915 | 1 |
| A9-10 | 874-875 | — | E3-4 | 916-917 | 1 |
| A11-12 | 876-877 | — | E5-6 | 918-919 | 1 |
| B1-2 | 878-879 | — | E7-8 | 920-921 | 1 |
| B3-4 | 880-881 | — | E9-10 | 922-923 | 1 |
| B5-6 | 882-883 | 1 | E11-12 | 924-925 | 1 |
| B7-8 | 884-885 | — | F1-2 | 926-927 | 1 |
| B9-10 | 886-887 | 1 | F3-4 | 928-929 | 1 |
| B11-12 | 888-889 | 1 | F5-6 | 930-931 | 1 |
| C1-2 | 890-891 | 1 | F7-8 | 932-933 | 1 |
| C3-4 | 892-893 | 1 | F9-10 | 934-935 | 1 |
| C5-6 | 894-895 | 1 | F11-12 | 936-937 | 1 |
| C7-8 | 896-897 | 1 | G1-2 | 938-939 | 1 |
| C9-10 | 898-899 | 1 | G3-4 | 940-941 | 1 |
| C11-12 | 900-901 | 1 | G5-6 | 942-943 | 1 |
| D1-2 | 902-903 | 1 | G7-8 | 944-945 | 1 |
| D3-4 | 904-905 | 1 | G9-12 | 946-949 | — |
| D5-6 | 906-907 | 1 | H1-12 | 950-961 | — |

**A**

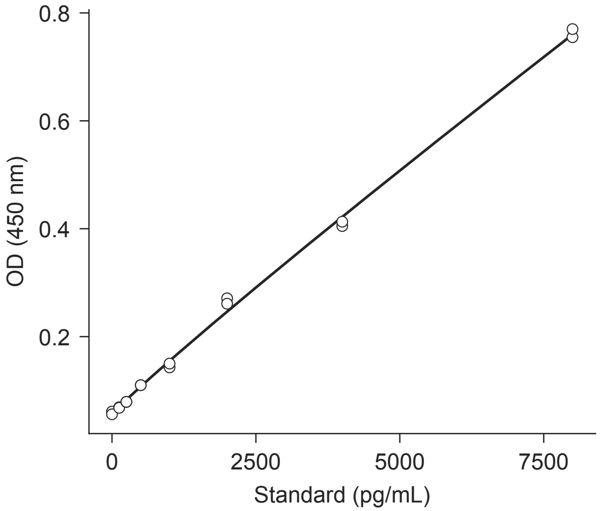


**B**

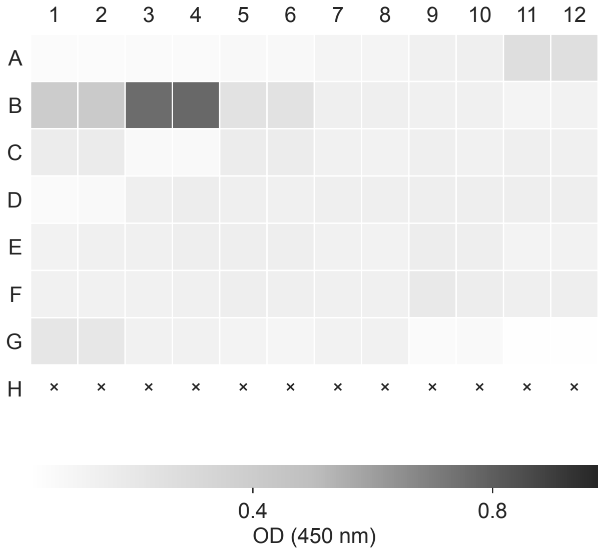


**C**

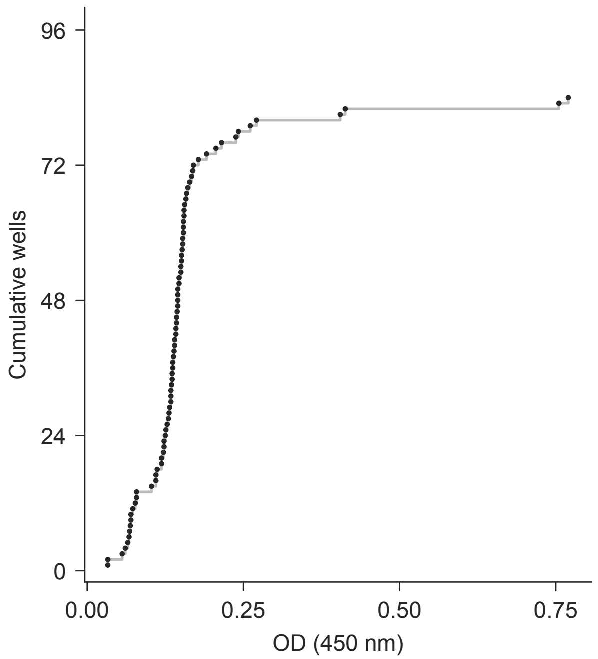


**D** elisa/plate_map.csv

| Well | Map rows | Dil. | Well | Map rows | Dil. |
|---|---|---|---|---|---|
| A1-2 | 962-963 | — | D7-8 | 1004-1005 | 1 |
| A3-4 | 964-965 | — | D9-10 | 1006-1007 | 1 |
| A5-6 | 966-967 | — | D11-12 | 1008-1009 | 1 |
| A7-8 | 968-969 | — | E1-2 | 1010-1011 | 1 |
| A9-10 | 970-971 | — | E3-4 | 1012-1013 | 1 |
| A11-12 | 972-973 | — | E5-6 | 1014-1015 | 1 |
| B1-2 | 974-975 | — | E7-8 | 1016-1017 | 1 |
| B3-4 | 976-977 | — | E9-10 | 1018-1019 | 1 |
| B5-6 | 978-979 | 1 | E11-12 | 1020-1021 | 1 |
| B7-8 | 980-981 | — | F1-2 | 1022-1023 | 1 |
| B9-10 | 982-983 | 1 | F3-4 | 1024-1025 | 1 |
| B11-12 | 984-985 | 1 | F5-6 | 1026-1027 | 1 |
| C1-2 | 986-987 | 1 | F7-8 | 1028-1029 | 1 |
| C3-4 | 988-989 | 1 | F9-10 | 1030-1031 | 1 |
| C5-6 | 990-991 | 1 | F11-12 | 1032-1033 | 1 |
| C7-8 | 992-993 | 1 | G1-2 | 1034-1035 | 1 |
| C9-10 | 994-995 | 1 | G3-4 | 1036-1037 | 1 |
| C11-12 | 996-997 | 1 | G5-6 | 1038-1039 | 1 |
| D1-2 | 998-999 | 1 | G7-8 | 1040-1041 | 1 |
| D3-4 | 1000-1001 | 1 | G9-12 | 1042-1045 | — |
| D5-6 | 1002-1003 | 1 | H1-12 | 1046-1057 | — |

**A**

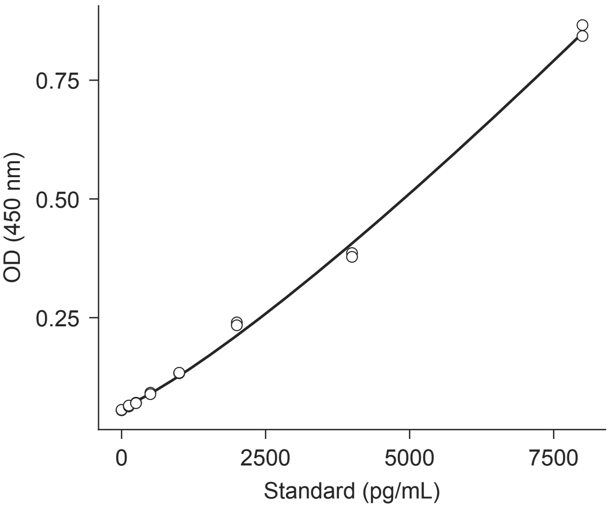


**B**

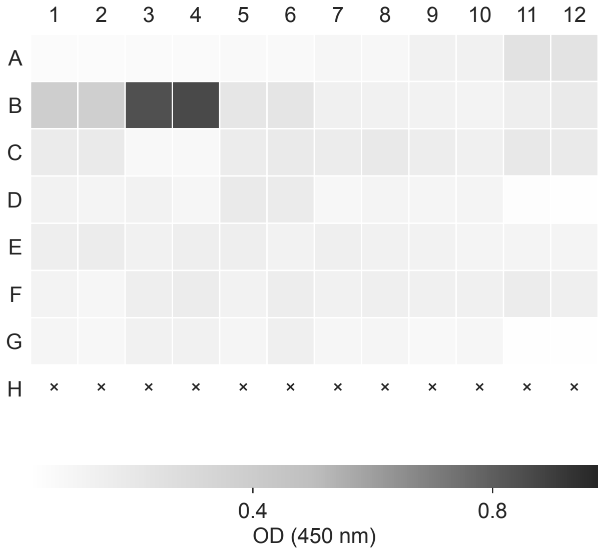


**C**

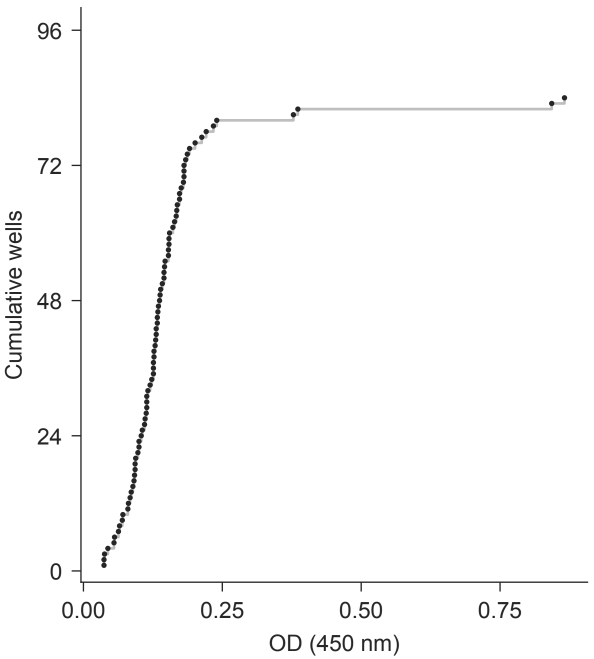


**D** elisa/plate_map.csv

| Well | Map rows | Dil. | Well | Map rows | Dil. |
|---|---|---|---|---|---|
| A1-2 | 1058-1059 | — | D7-8 | 1100-1101 | 1 |
| A3-4 | 1060-1061 | — | D9-10 | 1102-1103 | 1 |
| A5-6 | 1062-1063 | — | D11-12 | 1104-1105 | 1 |
| A7-8 | 1064-1065 | — | E1-2 | 1106-1107 | 1 |
| A9-10 | 1066-1067 | — | E3-4 | 1108-1109 | 1 |
| A11-12 | 1068-1069 | — | E5-6 | 1110-1111 | 1 |
| B1-2 | 1070-1071 | — | E7-8 | 1112-1113 | 1 |
| B3-4 | 1072-1073 | — | E9-10 | 1114-1115 | 1 |
| B5-6 | 1074-1075 | 1 | E11-12 | 1116-1117 | 1 |
| B7-8 | 1076-1077 | — | F1-2 | 1118-1119 | 1 |
| B9-10 | 1078-1079 | 1 | F3-4 | 1120-1121 | 1 |
| B11-12 | 1080-1081 | 1 | F5-6 | 1122-1123 | 1 |
| C1-2 | 1082-1083 | 1 | F7-8 | 1124-1125 | 1 |
| C3-4 | 1084-1085 | 1 | F9-10 | 1126-1127 | 1 |
| C5-6 | 1086-1087 | 1 | F11-12 | 1128-1129 | 1 |
| C7-8 | 1088-1089 | 1 | G1-2 | 1130-1131 | 1 |
| C9-10 | 1090-1091 | 1 | G3-4 | 1132-1133 | 1 |
| C11-12 | 1092-1093 | 1 | G5-6 | 1134-1135 | 1 |
| D1-2 | 1094-1095 | 1 | G7-8 | 1136-1137 | 1 |
| D3-4 | 1096-1097 | 1 | G9-12 | 1138-1141 | — |
| D5-6 | 1098-1099 | 1 | H1-12 | 1142-1153 | — |

**A**

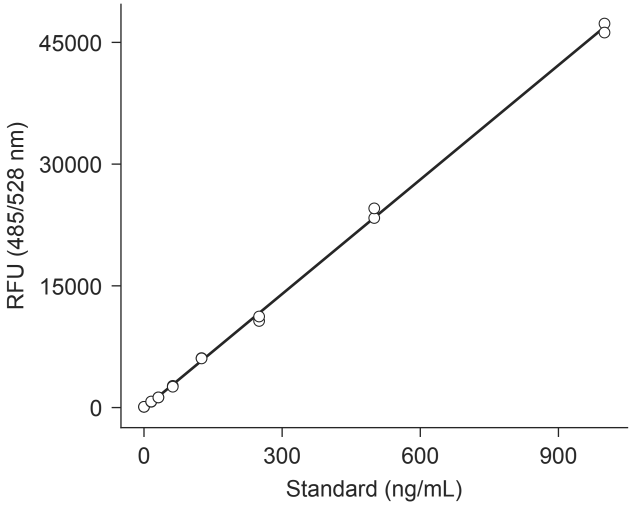


**B**

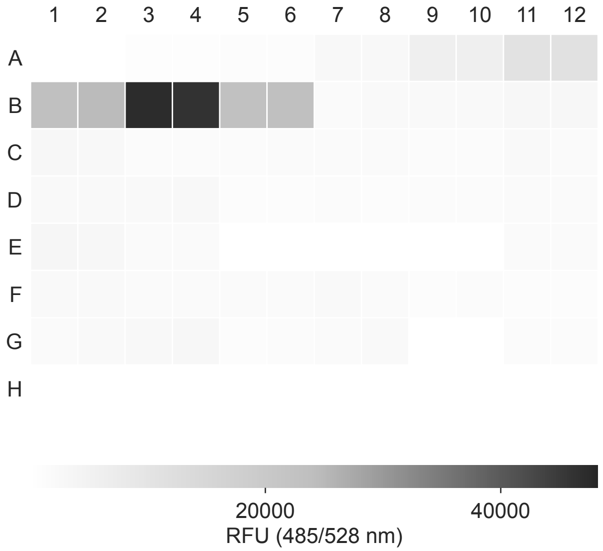


**C**

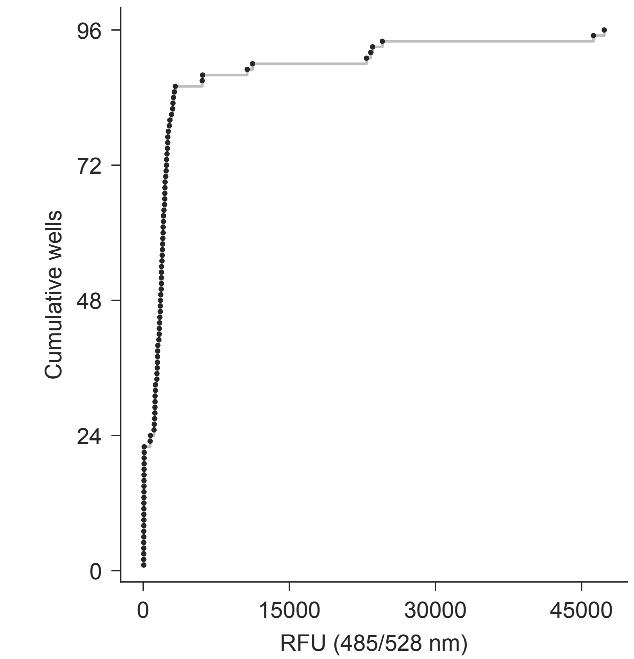


**D** dna/plate_map.csv

| Well | Map rows | Dil. | Well | Map rows | Dil. |
|---|---|---|---|---|---|
| A1-2 | 2-3 | — | D9-10 | 46-47 | 100 |
| A3-4 | 4-5 | — | D11-12 | 48-49 | 100 |
| A5-6 | 6-7 | — | E1-2 | 50-51 | 100 |
| A7-8 | 8-9 | — | E3-4 | 52-53 | 100 |
| A9-10 | 10-11 | — | E5-6 | 54-55 | 100 |
| A11-12 | 12-13 | — | E7-8 | 56-57 | 100 |
| B1-2 | 14-15 | — | E9-10 | 58-59 | 100 |
| B3-4 | 16-17 | — | E11-12 | 60-61 | 100 |
| B5-6 | 18-19 | 1 | F1-2 | 62-63 | 100 |
| B7-8 | 20-21 | — | F3-4 | 64-65 | 100 |
| B9-10 | 22-23 | 100 | F5-6 | 66-67 | 100 |
| B11-12 | 24-25 | 100 | F7-8 | 68-69 | 100 |
| C1-2 | 26-27 | 100 | F9-10 | 70-71 | 100 |
| C3-4 | 28-29 | 100 | F11-12 | 72-73 | 100 |
| C5-6 | 30-31 | 100 | G1-2 | 74-75 | 100 |
| C7-8 | 32-33 | 100 | G3-4 | 76-77 | 100 |
| C9-10 | 34-35 | 100 | G5-6 | 78-79 | 100 |
| C11-12 | 36-37 | 100 | G7-8 | 80-81 | 100 |
| D1-2 | 38-39 | 100 | G9-10 | 82-83 | 100 |
| D3-4 | 40-41 | 100 | G11-12 | 84-85 | 100 |
| D5-6 | 42-43 | 100 | H1-12 | 86-97 | — |
| D7-8 | 44-45 | 100 | | | |

**A**

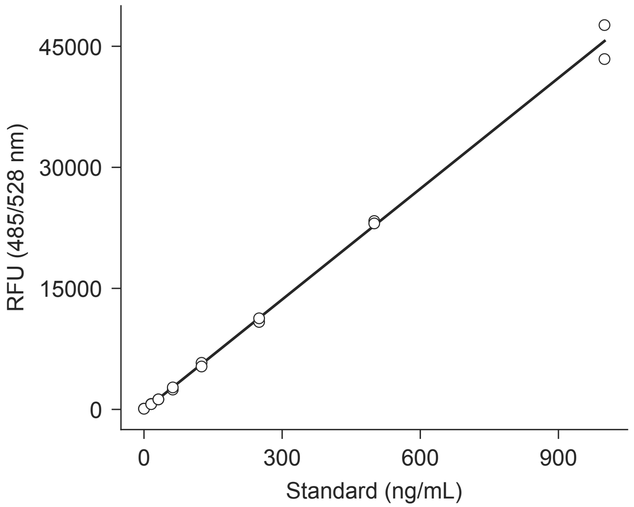


**B**

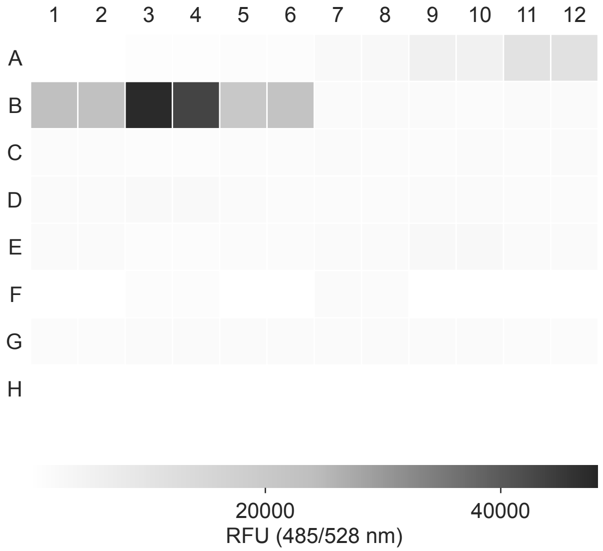


**C**

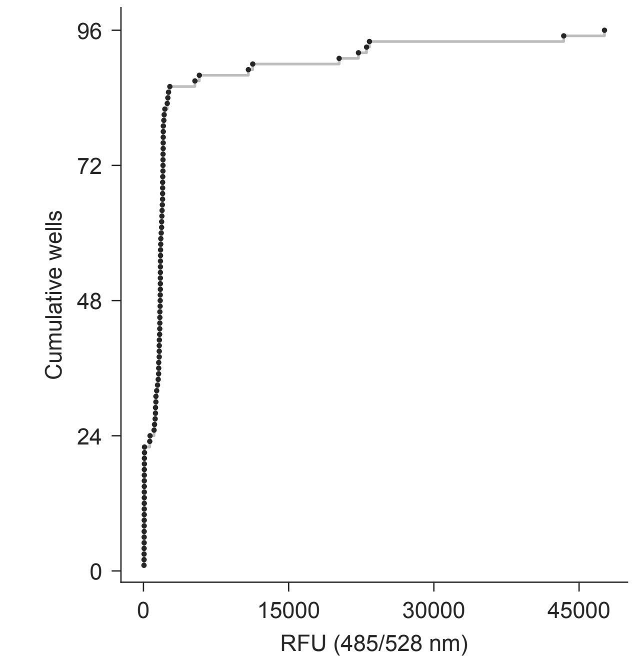


**D** dna/plate_map.csv

| Well | Map rows | Dil. | Well | Map rows | Dil. |
|---|---|---|---|---|---|
| A1-2 | 98-99 | — | D9-10 | 142-143 | 100 |
| A3-4 | 100-101 | — | D11-12 | 144-145 | 100 |
| A5-6 | 102-103 | — | E1-2 | 146-147 | 100 |
| A7-8 | 104-105 | — | E3-4 | 148-149 | 100 |
| A9-10 | 106-107 | — | E5-6 | 150-151 | 100 |
| A11-12 | 108-109 | — | E7-8 | 152-153 | 100 |
| B1-2 | 110-111 | — | E9-10 | 154-155 | 100 |
| B3-4 | 112-113 | — | E11-12 | 156-157 | 100 |
| B5-6 | 114-115 | 1 | F1-2 | 158-159 | 100 |
| B7-8 | 116-117 | — | F3-4 | 160-161 | 100 |
| B9-10 | 118-119 | 100 | F5-6 | 162-163 | 100 |
| B11-12 | 120-121 | 100 | F7-8 | 164-165 | 100 |
| C1-2 | 122-123 | 100 | F9-10 | 166-167 | 100 |
| C3-4 | 124-125 | 100 | F11-12 | 168-169 | 100 |
| C5-6 | 126-127 | 100 | G1-2 | 170-171 | 100 |
| C7-8 | 128-129 | 100 | G3-4 | 172-173 | 100 |
| C9-10 | 130-131 | 100 | G5-6 | 174-175 | 100 |
| C11-12 | 132-133 | 100 | G7-8 | 176-177 | 100 |
| D1-2 | 134-135 | 100 | G9-10 | 178-179 | 100 |
| D3-4 | 136-137 | 100 | G11-12 | 180-181 | 100 |
| D5-6 | 138-139 | 100 | H1-12 | 182-193 | — |
| D7-8 | 140-141 | 100 | | | |

**A**

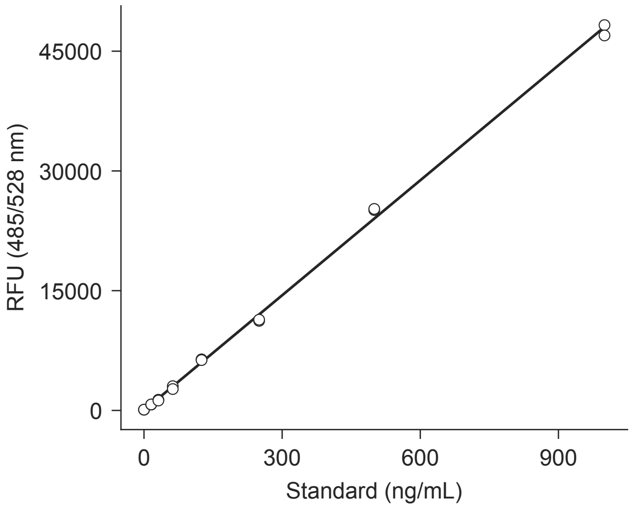


**B**

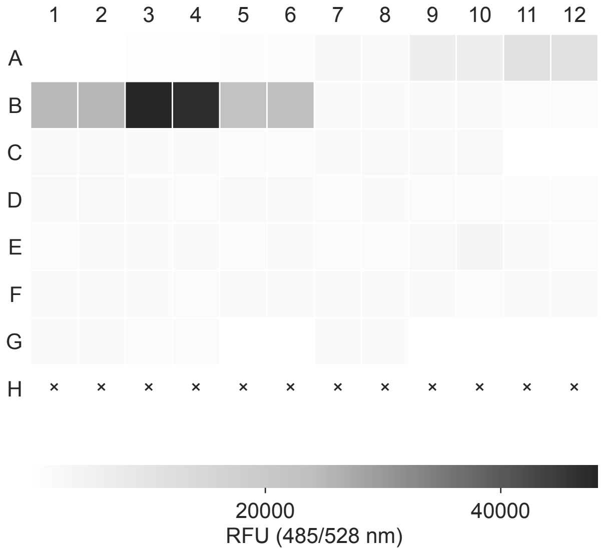


**C**

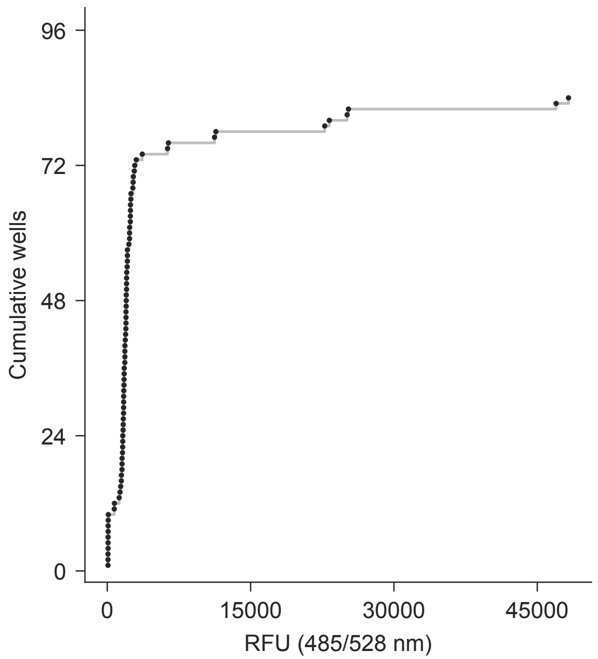


**D** dna/plate_map.csv

| Well | Map rows | Dil. | Well | Map rows | Dil. |
|---|---|---|---|---|---|
| A1-2 | 194-195 | — | D7-8 | 236-237 | 100 |
| A3-4 | 196-197 | — | D9-10 | 238-239 | 100 |
| A5-6 | 198-199 | — | D11-12 | 240-241 | 100 |
| A7-8 | 200-201 | — | E1-2 | 242-243 | 100 |
| A9-10 | 202-203 | — | E3-4 | 244-245 | 100 |
| A11-12 | 204-205 | — | E5-6 | 246-247 | 100 |
| B1-2 | 206-207 | — | E7-8 | 248-249 | 100 |
| B3-4 | 208-209 | — | E9-10 | 250-251 | 100 |
| B5-6 | 210-211 | 1 | E11-12 | 252-253 | 100 |
| B7-8 | 212-213 | — | F1-2 | 254-255 | 100 |
| B9-10 | 214-215 | 100 | F3-4 | 256-257 | 100 |
| B11-12 | 216-217 | 100 | F5-6 | 258-259 | 100 |
| C1-2 | 218-219 | 100 | F7-8 | 260-261 | 100 |
| C3-4 | 220-221 | 100 | F9-10 | 262-263 | 100 |
| C5-6 | 222-223 | 100 | F11-12 | 264-265 | 100 |
| C7-8 | 224-225 | 100 | G1-2 | 266-267 | 100 |
| C9-10 | 226-227 | 100 | G3-4 | 268-269 | 100 |
| C11-12 | 228-229 | 100 | G5-6 | 270-271 | 100 |
| D1-2 | 230-231 | 100 | G7-8 | 272-273 | 100 |
| D3-4 | 232-233 | 100 | G9-12 | 274-277 | — |
| D5-6 | 234-235 | 100 | H1-12 | 278-289 | — |

**A**

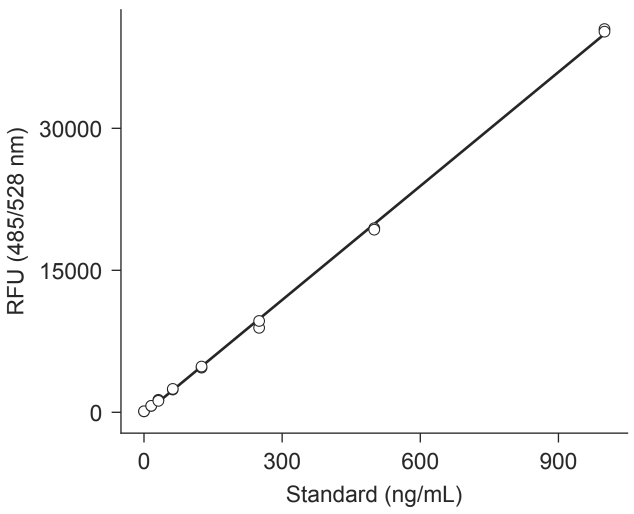


**B**

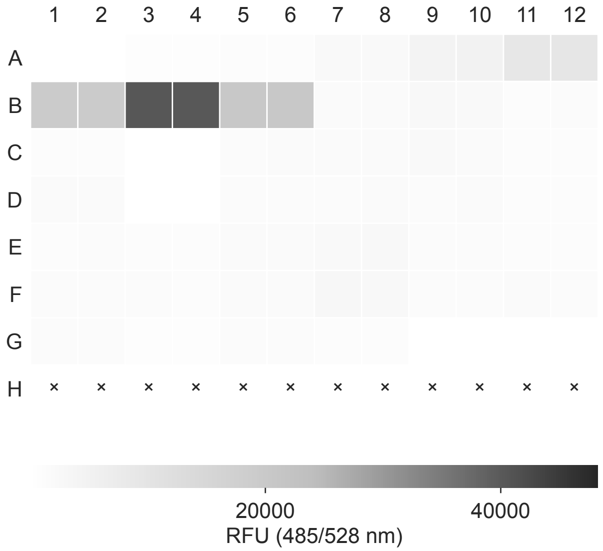


**C**

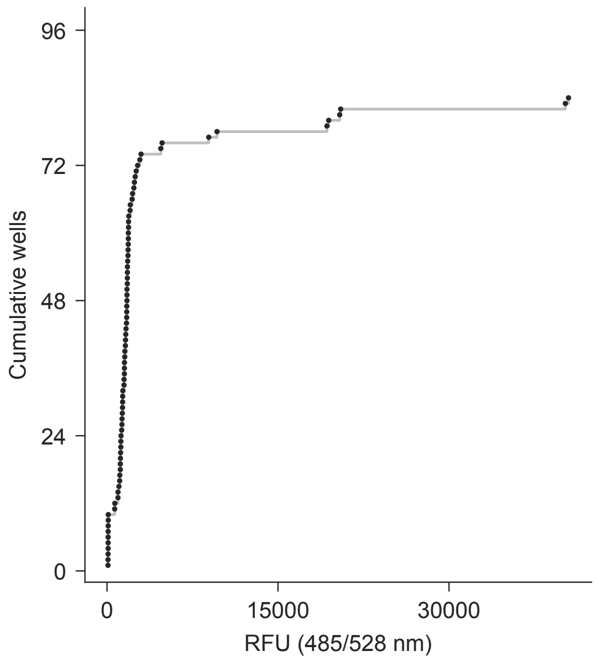


**D** dna/plate_map.csv

| Well | Map rows | Dil. | Well | Map rows | Dil. |
|---|---|---|---|---|---|
| A1-2 | 290-291 | — | D7-8 | 332-333 | 100 |
| A3-4 | 292-293 | — | D9-10 | 334-335 | 100 |
| A5-6 | 294-295 | — | D11-12 | 336-337 | 100 |
| A7-8 | 296-297 | — | E1-2 | 338-339 | 100 |
| A9-10 | 298-299 | — | E3-4 | 340-341 | 100 |
| A11-12 | 300-301 | — | E5-6 | 342-343 | 100 |
| B1-2 | 302-303 | — | E7-8 | 344-345 | 100 |
| B3-4 | 304-305 | — | E9-10 | 346-347 | 100 |
| B5-6 | 306-307 | 1 | E11-12 | 348-349 | 100 |
| B7-8 | 308-309 | — | F1-2 | 350-351 | 100 |
| B9-10 | 310-311 | 100 | F3-4 | 352-353 | 100 |
| B11-12 | 312-313 | 100 | F5-6 | 354-355 | 100 |
| C1-2 | 314-315 | 100 | F7-8 | 356-357 | 100 |
| C3-4 | 316-317 | 100 | F9-10 | 358-359 | 100 |
| C5-6 | 318-319 | 100 | F11-12 | 360-361 | 100 |
| C7-8 | 320-321 | 100 | G1-2 | 362-363 | 100 |
| C9-10 | 322-323 | 100 | G3-4 | 364-365 | 100 |
| C11-12 | 324-325 | 100 | G5-6 | 366-367 | 100 |
| D1-2 | 326-327 | 100 | G7-8 | 368-369 | 100 |
| D3-4 | 328-329 | 100 | G9-12 | 370-373 | — |
| D5-6 | 330-331 | 100 | H1-12 | 374-385 | — |

**A**

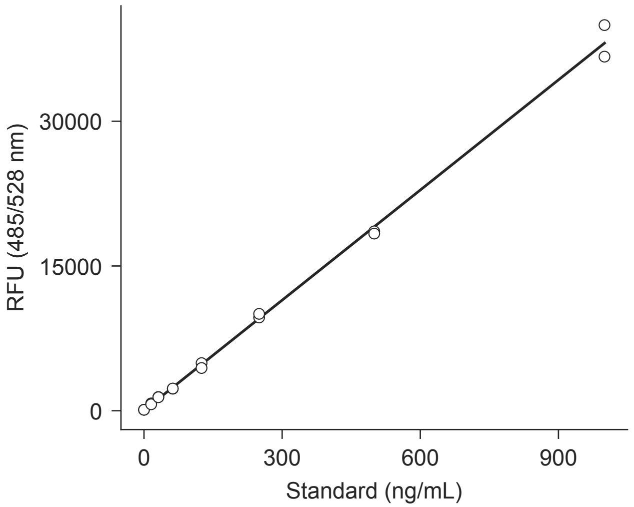


**B**

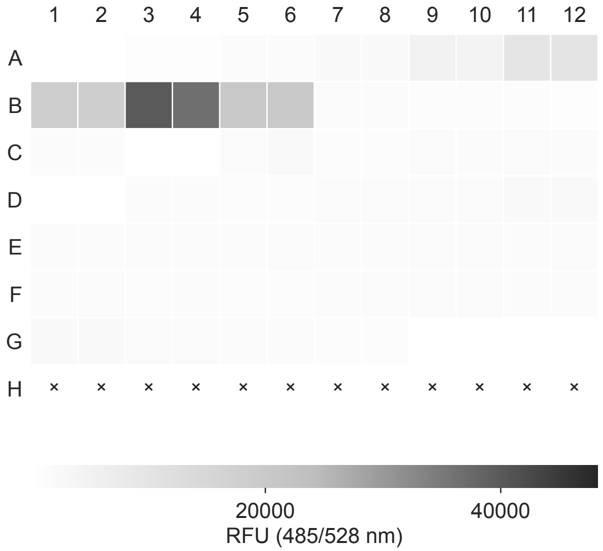


**C**

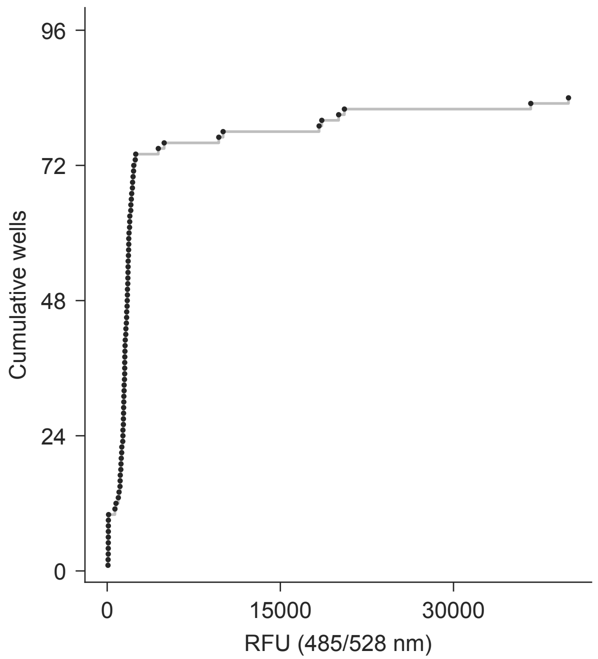


**D** dna/plate_map.csv

| Well | Map rows | Dil. | Well | Map rows | Dil. |
|---|---|---|---|---|---|
| A1-2 | 386-387 | — | D7-8 | 428-429 | 100 |
| A3-4 | 388-389 | — | D9-10 | 430-431 | 100 |
| A5-6 | 390-391 | — | D11-12 | 432-433 | 100 |
| A7-8 | 392-393 | — | E1-2 | 434-435 | 100 |
| A9-10 | 394-395 | — | E3-4 | 436-437 | 100 |
| A11-12 | 396-397 | — | E5-6 | 438-439 | 100 |
| B1-2 | 398-399 | — | E7-8 | 440-441 | 100 |
| B3-4 | 400-401 | — | E9-10 | 442-443 | 100 |
| B5-6 | 402-403 | 1 | E11-12 | 444-445 | 100 |
| B7-8 | 404-405 | — | F1-2 | 446-447 | 100 |
| B9-10 | 406-407 | 100 | F3-4 | 448-449 | 100 |
| B11-12 | 408-409 | 100 | F5-6 | 450-451 | 100 |
| C1-2 | 410-411 | 100 | F7-8 | 452-453 | 100 |
| C3-4 | 412-413 | 100 | F9-10 | 454-455 | 100 |
| C5-6 | 414-415 | 100 | F11-12 | 456-457 | 100 |
| C7-8 | 416-417 | 100 | G1-2 | 458-459 | 100 |
| C9-10 | 418-419 | 100 | G3-4 | 460-461 | 100 |
| C11-12 | 420-421 | 100 | G5-6 | 462-463 | 100 |
| D1-2 | 422-423 | 100 | G7-8 | 464-465 | 100 |
| D3-4 | 424-425 | 100 | G9-12 | 466-469 | — |
| D5-6 | 426-427 | 100 | H1-12 | 470-481 | — |

Synergy H1 | 6/27/2026 | functional | DNA-06

**A**

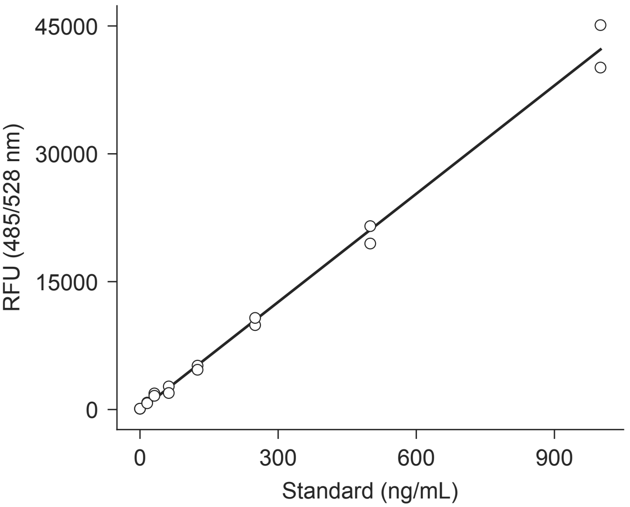


**B**

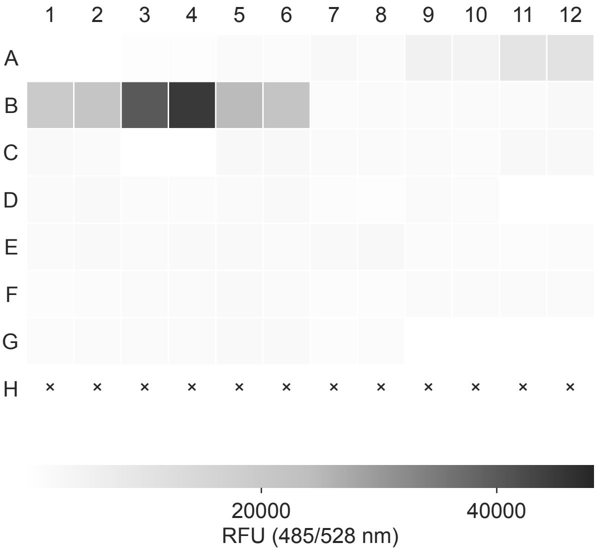


**C**

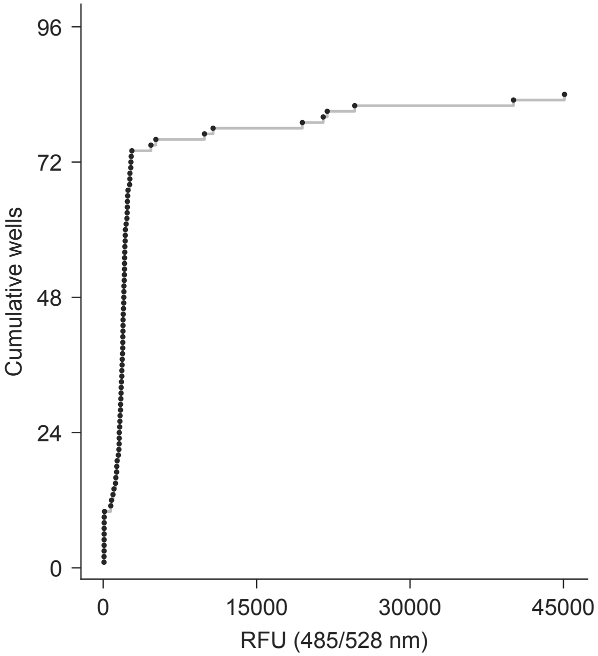


**D** dna/plate_map.csv

| Well | Map rows | Dil. | Well | Map rows | Dil. |
|---|---|---|---|---|---|
| A1-2 | 482-483 | — | D7-8 | 524-525 | 100 |
| A3-4 | 484-485 | — | D9-10 | 526-527 | 100 |
| A5-6 | 486-487 | — | D11-12 | 528-529 | 100 |
| A7-8 | 488-489 | — | E1-2 | 530-531 | 100 |
| A9-10 | 490-491 | — | E3-4 | 532-533 | 100 |
| A11-12 | 492-493 | — | E5-6 | 534-535 | 100 |
| B1-2 | 494-495 | — | E7-8 | 536-537 | 100 |
| B3-4 | 496-497 | — | E9-10 | 538-539 | 100 |
| B5-6 | 498-499 | 1 | E11-12 | 540-541 | 100 |
| B7-8 | 500-501 | — | F1-2 | 542-543 | 100 |
| B9-10 | 502-503 | 100 | F3-4 | 544-545 | 100 |
| B11-12 | 504-505 | 100 | F5-6 | 546-547 | 100 |
| C1-2 | 506-507 | 100 | F7-8 | 548-549 | 100 |
| C3-4 | 508-509 | 100 | F9-10 | 550-551 | 100 |
| C5-6 | 510-511 | 100 | F11-12 | 552-553 | 100 |
| C7-8 | 512-513 | 100 | G1-2 | 554-555 | 100 |
| C9-10 | 514-515 | 100 | G3-4 | 556-557 | 100 |
| C11-12 | 516-517 | 100 | G5-6 | 558-559 | 100 |
| D1-2 | 518-519 | 100 | G7-8 | 560-561 | 100 |
| D3-4 | 520-521 | 100 | G9-12 | 562-565 | — |
| D5-6 | 522-523 | 100 | H1-12 | 566-577 | — |

**B**

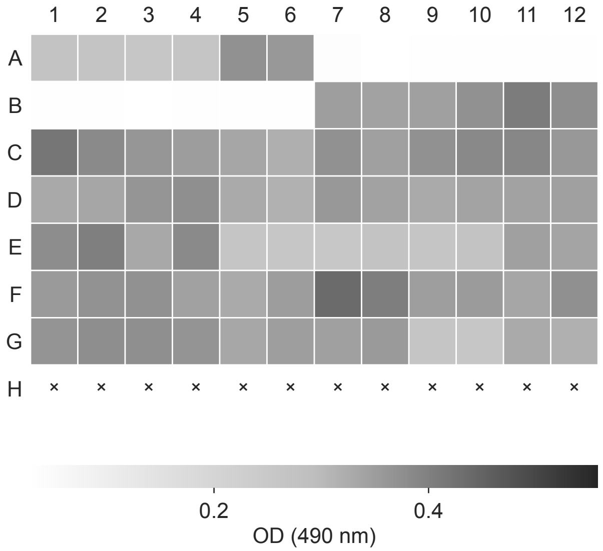


**C**

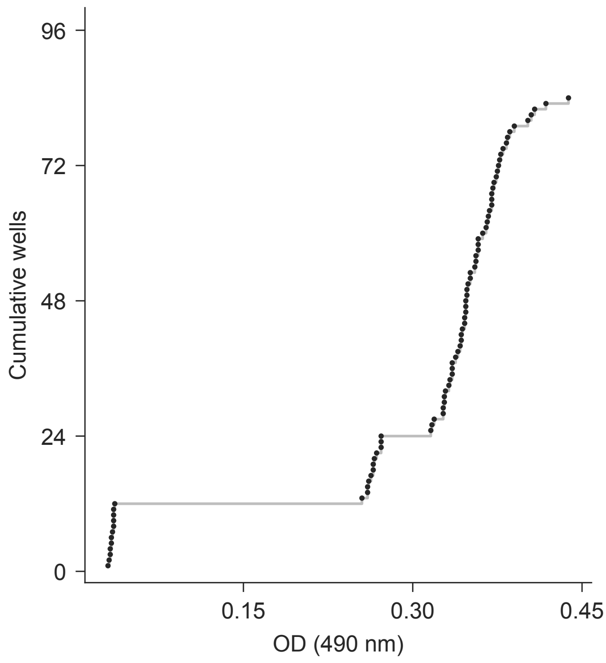


**D** ldh/plate_map.csv

| Well | Map rows | Dil. | Well | Map rows | Dil. |
|---|---|---|---|---|---|
| A1-4 | 2-5 | — | E1-2 | 50-51 | 1 |
| A5-6 | 6-7 | 1 | E3-4 | 52-53 | 1 |
| A7-12 | 8-13 | — | E5-6 | 54-55 | 1 |
| B1-6 | 14-19 | — | E7-8 | 56-57 | 1 |
| B7-8 | 20-21 | — | E9-10 | 58-59 | 1 |
| B9-10 | 22-23 | 1 | E11-12 | 60-61 | 1 |
| B11-12 | 24-25 | 1 | F1-2 | 62-63 | 1 |
| C1-2 | 26-27 | 1 | F3-4 | 64-65 | 1 |
| C3-4 | 28-29 | 1 | F5-6 | 66-67 | 1 |
| C5-6 | 30-31 | 1 | F7-8 | 68-69 | 1 |
| C7-8 | 32-33 | 1 | F9-10 | 70-71 | 1 |
| C9-10 | 34-35 | 1 | F11-12 | 72-73 | 1 |
| C11-12 | 36-37 | 1 | G1-2 | 74-75 | 1 |
| D1-2 | 38-39 | 1 | G3-4 | 76-77 | 1 |
| D3-4 | 40-41 | 1 | G5-6 | 78-79 | 1 |
| D5-6 | 42-43 | 1 | G7-8 | 80-81 | 1 |
| D7-8 | 44-45 | 1 | G9-10 | 82-83 | 1 |
| D9-10 | 46-47 | 1 | G11-12 | 84-85 | 1 |
| D11-12 | 48-49 | 1 | H1-12 | 86-97 | — |

**B**

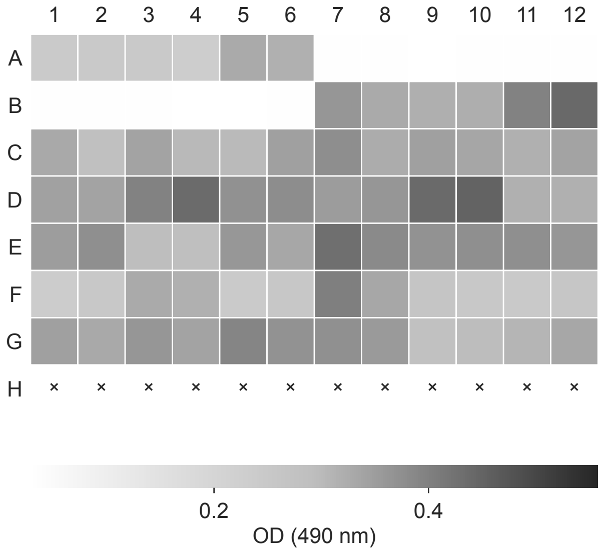


**C**

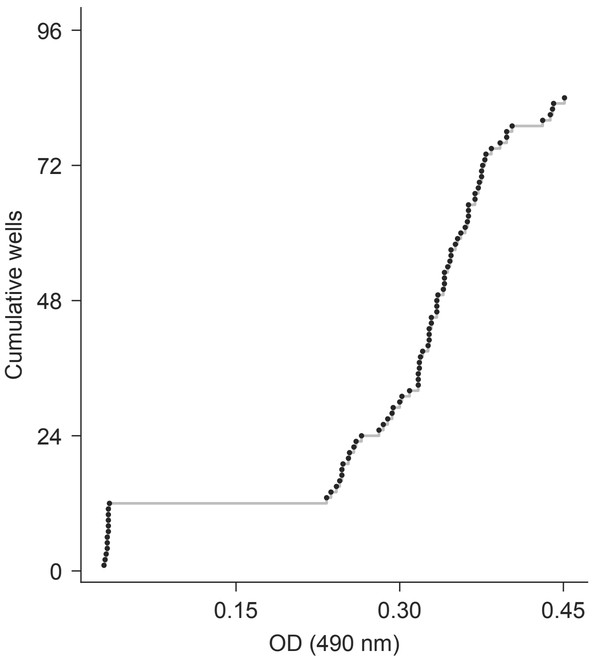


**D** ldh/plate_map.csv

| Well | Map rows | Dil. | Well | Map rows | Dil. |
|---|---|---|---|---|---|
| A1-4 | 98-101 | — | E1-2 | 146-147 | 1 |
| A5-6 | 102-103 | 1 | E3-4 | 148-149 | 1 |
| A7-12 | 104-109 | — | E5-6 | 150-151 | 1 |
| B1-6 | 110-115 | — | E7-8 | 152-153 | 1 |
| B7-8 | 116-117 | — | E9-10 | 154-155 | 1 |
| B9-10 | 118-119 | 1 | E11-12 | 156-157 | 1 |
| B11-12 | 120-121 | 1 | F1-2 | 158-159 | 1 |
| C1-2 | 122-123 | 1 | F3-4 | 160-161 | 1 |
| C3-4 | 124-125 | 1 | F5-6 | 162-163 | 1 |
| C5-6 | 126-127 | 1 | F7-8 | 164-165 | 1 |
| C7-8 | 128-129 | 1 | F9-10 | 166-167 | 1 |
| C9-10 | 130-131 | 1 | F11-12 | 168-169 | 1 |
| C11-12 | 132-133 | 1 | G1-2 | 170-171 | 1 |
| D1-2 | 134-135 | 1 | G3-4 | 172-173 | 1 |
| D3-4 | 136-137 | 1 | G5-6 | 174-175 | 1 |
| D5-6 | 138-139 | 1 | G7-8 | 176-177 | 1 |
| D7-8 | 140-141 | 1 | G9-10 | 178-179 | 1 |
| D9-10 | 142-143 | 1 | G11-12 | 180-181 | 1 |
| D11-12 | 144-145 | 1 | H1-12 | 182-193 | — |

LDH_Plate03_20260408_Results.txt

Synergy H1 | 4/8/2026 | functional | LDH-03

**B**

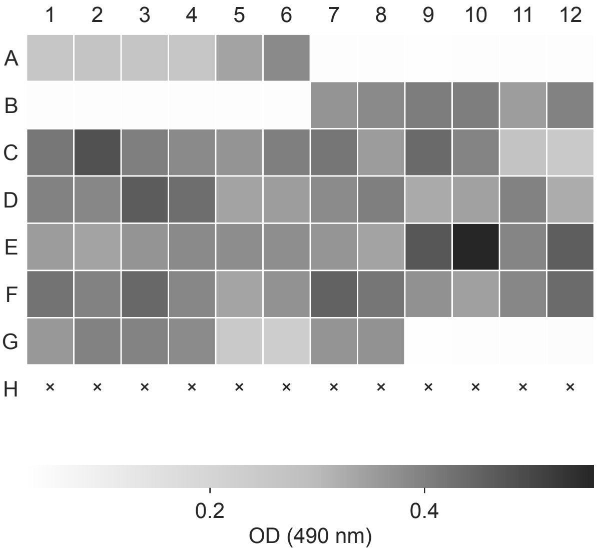


**C**

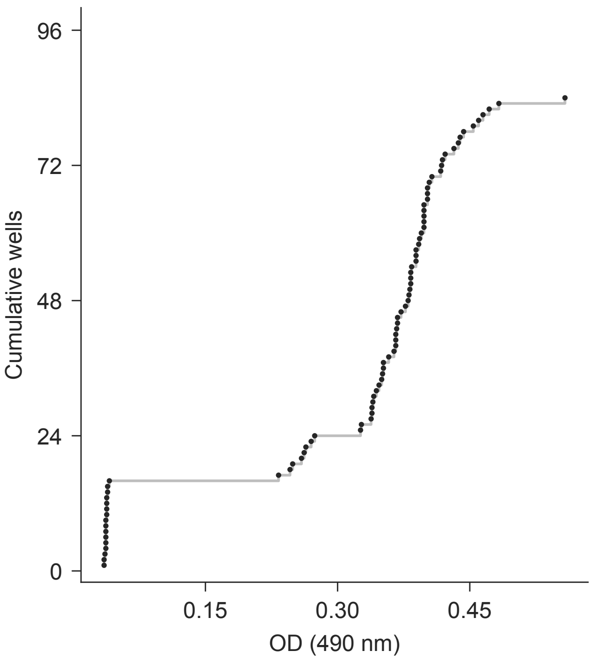


**D** ldh/plate_map.csv

| Well | Map rows | Dil. | Well | Map rows | Dil. |
|---|---|---|---|---|---|
| A1-4 | 194-197 | — | E1-2 | 242-243 | 1 |
| A5-6 | 198-199 | 1 | E3-4 | 244-245 | 1 |
| A7-12 | 200-205 | — | E5-6 | 246-247 | 1 |
| B1-6 | 206-211 | — | E7-8 | 248-249 | 1 |
| B7-8 | 212-213 | — | E9-10 | 250-251 | 1 |
| B9-10 | 214-215 | 1 | E11-12 | 252-253 | 1 |
| B11-12 | 216-217 | 1 | F1-2 | 254-255 | 1 |
| C1-2 | 218-219 | 1 | F3-4 | 256-257 | 1 |
| C3-4 | 220-221 | 1 | F5-6 | 258-259 | 1 |
| C5-6 | 222-223 | 1 | F7-8 | 260-261 | 1 |
| C7-8 | 224-225 | 1 | F9-10 | 262-263 | 1 |
| C9-10 | 226-227 | 1 | F11-12 | 264-265 | 1 |
| C11-12 | 228-229 | 1 | G1-2 | 266-267 | 1 |
| D1-2 | 230-231 | 1 | G3-4 | 268-269 | 1 |
| D3-4 | 232-233 | 1 | G5-6 | 270-271 | 1 |
| D5-6 | 234-235 | 1 | G7-8 | 272-273 | 1 |
| D7-8 | 236-237 | 1 | G9-12 | 274-277 | — |
| D9-10 | 238-239 | 1 | H1-12 | 278-289 | — |
| D11-12 | 240-241 | 1 | | | |

LDH_Plate04_20260414_Results.txt

Synergy H1 | 4/14/2026 | functional | LDH-04

**B**

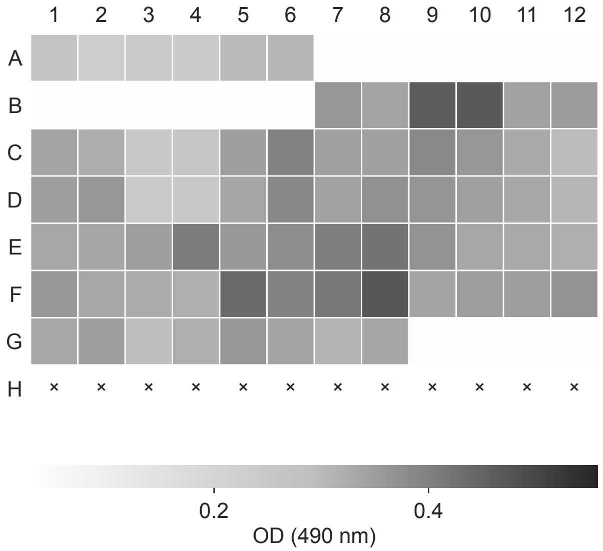


**C**

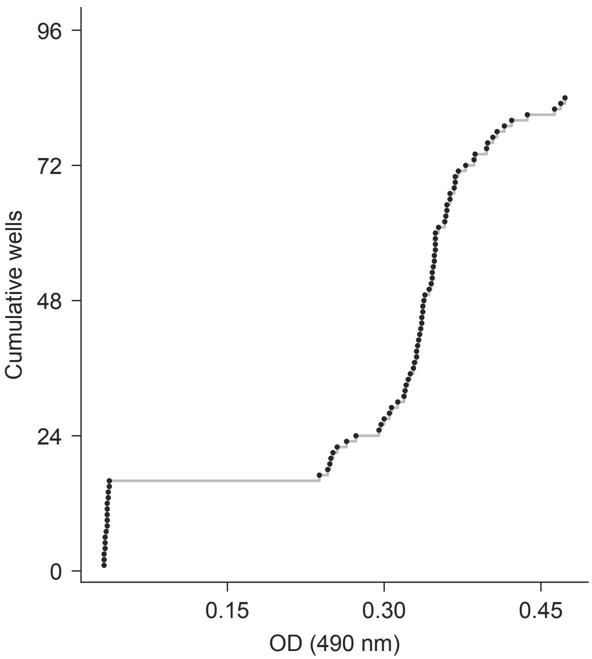


**D** ldh/plate_map.csv

| Well | Map rows | Dil. | Well | Map rows | Dil. |
|---|---|---|---|---|---|
| A1-4 | 290-293 | — | E1-2 | 338-339 | 1 |
| A5-6 | 294-295 | 1 | E3-4 | 340-341 | 1 |
| A7-12 | 296-301 | — | E5-6 | 342-343 | 1 |
| B1-6 | 302-307 | — | E7-8 | 344-345 | 1 |
| B7-8 | 308-309 | — | E9-10 | 346-347 | 1 |
| B9-10 | 310-311 | 1 | E11-12 | 348-349 | 1 |
| B11-12 | 312-313 | 1 | F1-2 | 350-351 | 1 |
| C1-2 | 314-315 | 1 | F3-4 | 352-353 | 1 |
| C3-4 | 316-317 | 1 | F5-6 | 354-355 | 1 |
| C5-6 | 318-319 | 1 | F7-8 | 356-357 | 1 |
| C7-8 | 320-321 | 1 | F9-10 | 358-359 | 1 |
| C9-10 | 322-323 | 1 | F11-12 | 360-361 | 1 |
| C11-12 | 324-325 | 1 | G1-2 | 362-363 | 1 |
| D1-2 | 326-327 | 1 | G3-4 | 364-365 | 1 |
| D3-4 | 328-329 | 1 | G5-6 | 366-367 | 1 |
| D5-6 | 330-331 | 1 | G7-8 | 368-369 | 1 |
| D7-8 | 332-333 | 1 | G9-12 | 370-373 | — |
| D9-10 | 334-335 | 1 | H1-12 | 374-385 | — |
| D11-12 | 336-337 | 1 | | | |

Synergy H1 | 4/14/2026 | functional | LDH-05

**B**

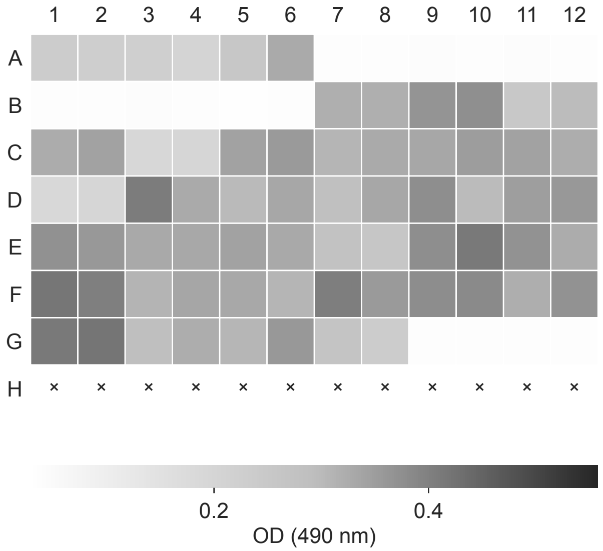


**C**

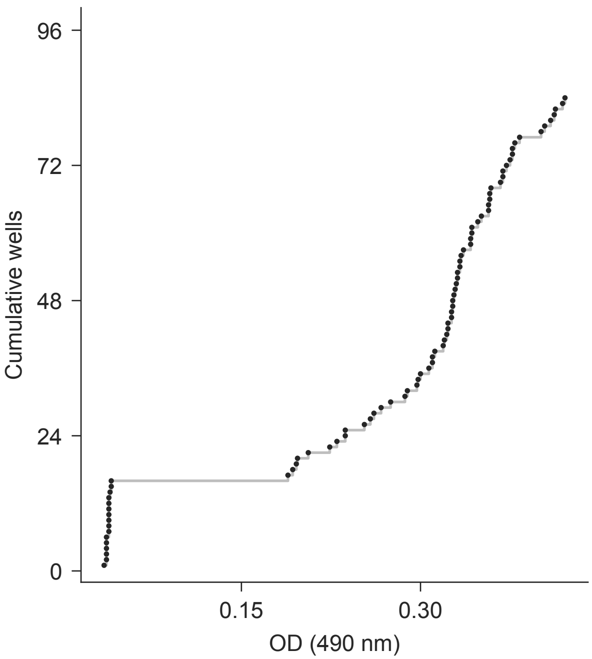


**D** ldh/plate_map.csv

| Well | Map rows | Dil. | Well | Map rows | Dil. |
|---|---|---|---|---|---|
| A1-4 | 386-389 | — | E1-2 | 434-435 | 1 |
| A5-6 | 390-391 | 1 | E3-4 | 436-437 | 1 |
| A7-12 | 392-397 | — | E5-6 | 438-439 | 1 |
| B1-6 | 398-403 | — | E7-8 | 440-441 | 1 |
| B7-8 | 404-405 | — | E9-10 | 442-443 | 1 |
| B9-10 | 406-407 | 1 | E11-12 | 444-445 | 1 |
| B11-12 | 408-409 | 1 | F1-2 | 446-447 | 1 |
| C1-2 | 410-411 | 1 | F3-4 | 448-449 | 1 |
| C3-4 | 412-413 | 1 | F5-6 | 450-451 | 1 |
| C5-6 | 414-415 | 1 | F7-8 | 452-453 | 1 |
| C7-8 | 416-417 | 1 | F9-10 | 454-455 | 1 |
| C9-10 | 418-419 | 1 | F11-12 | 456-457 | 1 |
| C11-12 | 420-421 | 1 | G1-2 | 458-459 | 1 |
| D1-2 | 422-423 | 1 | G3-4 | 460-461 | 1 |
| D3-4 | 424-425 | 1 | G5-6 | 462-463 | 1 |
| D5-6 | 426-427 | 1 | G7-8 | 464-465 | 1 |
| D7-8 | 428-429 | 1 | G9-12 | 466-469 | — |
| D9-10 | 430-431 | 1 | H1-12 | 470-481 | — |
| D11-12 | 432-433 | 1 | | | |

**B**

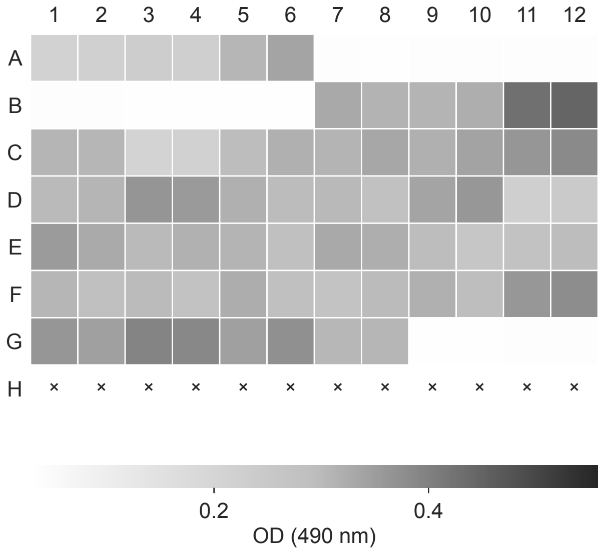


**C**

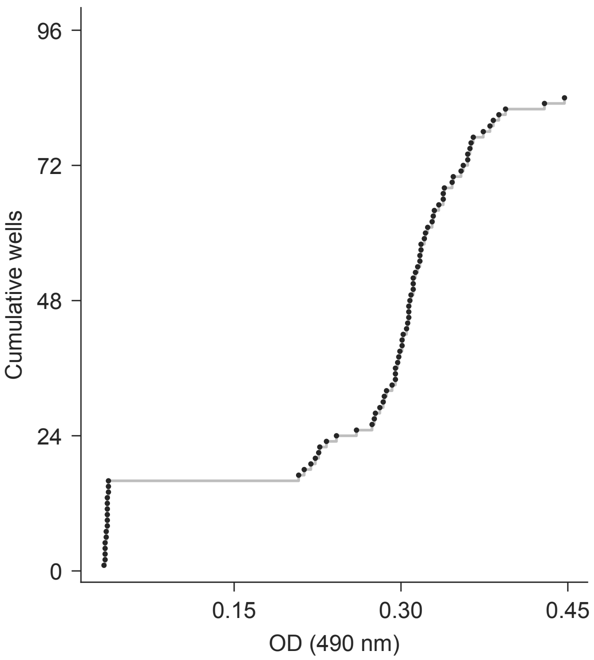


**D** ldh/plate_map.csv

| Well | Map rows | Dil. | Well | Map rows | Dil. |
|---|---|---|---|---|---|
| A1-4 | 482-485 | — | E1-2 | 530-531 | 1 |
| A5-6 | 486-487 | 1 | E3-4 | 532-533 | 1 |
| A7-12 | 488-493 | — | E5-6 | 534-535 | 1 |
| B1-6 | 494-499 | — | E7-8 | 536-537 | 1 |
| B7-8 | 500-501 | — | E9-10 | 538-539 | 1 |
| B9-10 | 502-503 | 1 | E11-12 | 540-541 | 1 |
| B11-12 | 504-505 | 1 | F1-2 | 542-543 | 1 |
| C1-2 | 506-507 | 1 | F3-4 | 544-545 | 1 |
| C3-4 | 508-509 | 1 | F5-6 | 546-547 | 1 |
| C5-6 | 510-511 | 1 | F7-8 | 548-549 | 1 |
| C7-8 | 512-513 | 1 | F9-10 | 550-551 | 1 |
| C9-10 | 514-515 | 1 | F11-12 | 552-553 | 1 |
| C11-12 | 516-517 | 1 | G1-2 | 554-555 | 1 |
| D1-2 | 518-519 | 1 | G3-4 | 556-557 | 1 |
| D3-4 | 520-521 | 1 | G5-6 | 558-559 | 1 |
| D5-6 | 522-523 | 1 | G7-8 | 560-561 | 1 |
| D7-8 | 524-525 | 1 | G9-12 | 562-565 | — |
| D9-10 | 526-527 | 1 | H1-12 | 566-577 | — |
| D11-12 | 528-529 | 1 | | | |

**A**

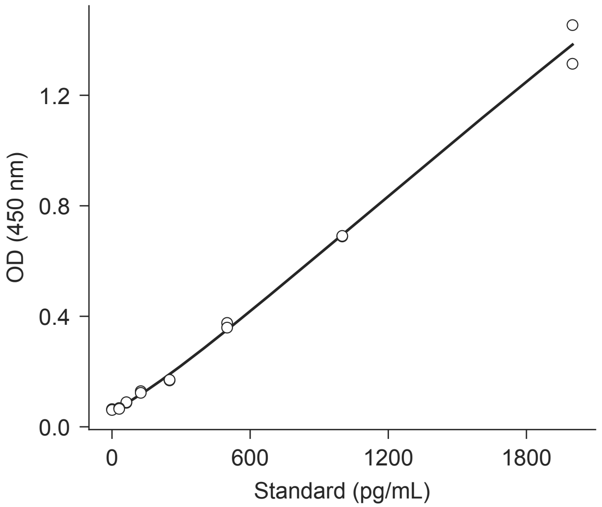


**B**

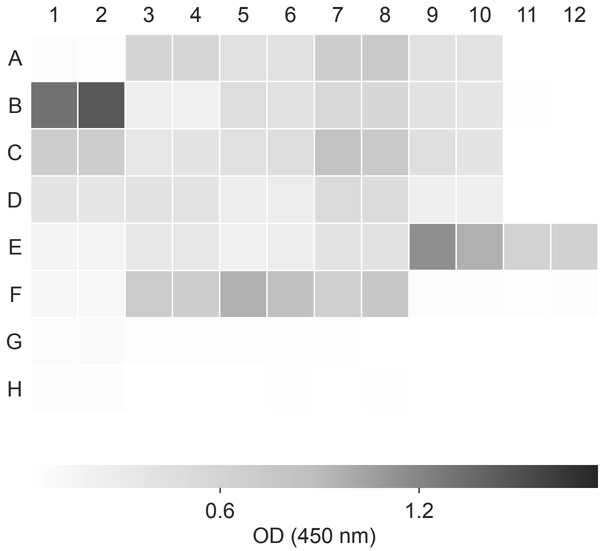


**C**

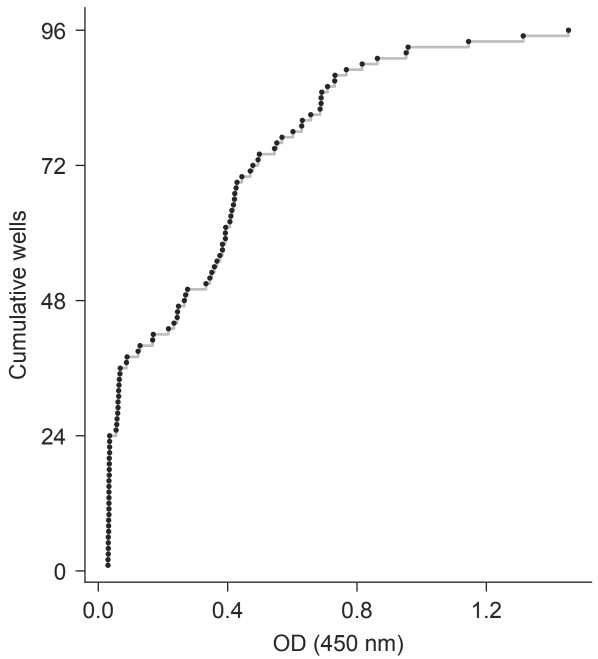


**D** elisa/plate_map.csv

| Well | Map rows | Dil. | Well | Map rows | Dil. |
|---|---|---|---|---|---|
| A1-2 | 2-3 | — | D7-8 | 44-45 | 1.0 |
| A3-4 | 4-5 | 1.0 | D9-10 | 46-47 | 1.0 |
| A5-6 | 6-7 | 1.0 | D11-12 | 48-49 | — |
| A7-8 | 8-9 | 1.0 | E1-2 | 50-51 | — |
| A9-10 | 10-11 | 1.0 | E3-4 | 52-53 | 1.0 |
| A11-12 | 12-13 | — | E5-6 | 54-55 | 1.0 |
| B1-2 | 14-15 | — | E7-8 | 56-57 | 1.0 |
| B3-4 | 16-17 | 1.0 | E9-10 | 58-59 | 1.0 |
| B5-6 | 18-19 | 1.0 | E11-12 | 60-61 | 1.0 |
| B7-8 | 20-21 | 1.0 | F1-2 | 62-63 | — |
| B9-10 | 22-23 | 1.0 | F3-4 | 64-65 | 1.0 |
| B11-12 | 24-25 | — | F5-6 | 66-67 | 1.0 |
| C1-2 | 26-27 | — | F7-8 | 68-69 | 1.0 |
| C3-4 | 28-29 | 1.0 | F9-10 | 70-71 | 1.0 |
| C5-6 | 30-31 | 1.0 | F11-12 | 72-73 | 1.0 |
| C7-8 | 32-33 | 1.0 | G1-2 | 74-75 | — |
| C9-10 | 34-35 | 1.0 | G3-4 | 76-77 | 1.0 |
| C11-12 | 36-37 | — | G5-6 | 78-79 | 1.0 |
| D1-2 | 38-39 | — | G7-12 | 80-85 | — |
| D3-4 | 40-41 | 1.0 | H1-2 | 86-87 | — |
| D5-6 | 42-43 | 1.0 | H3-12 | 88-97 | — |

**A**

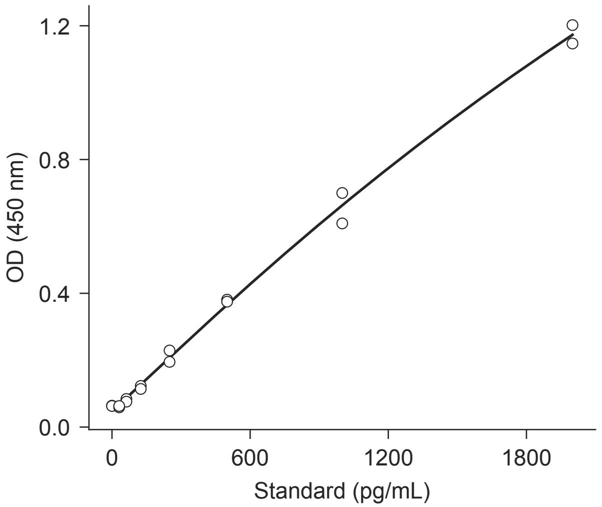


**B**

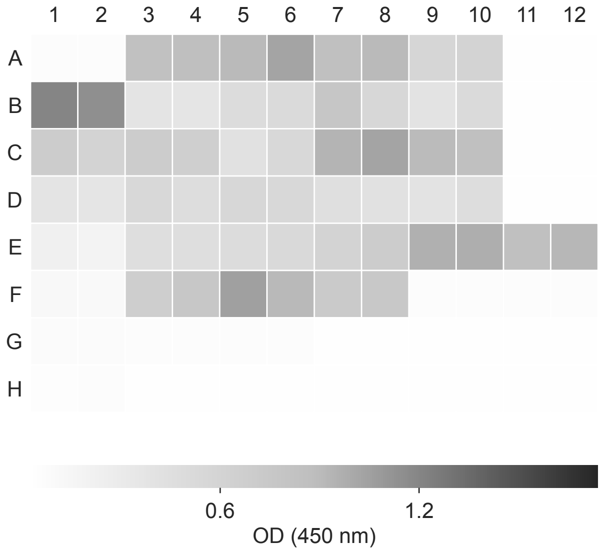


**C**

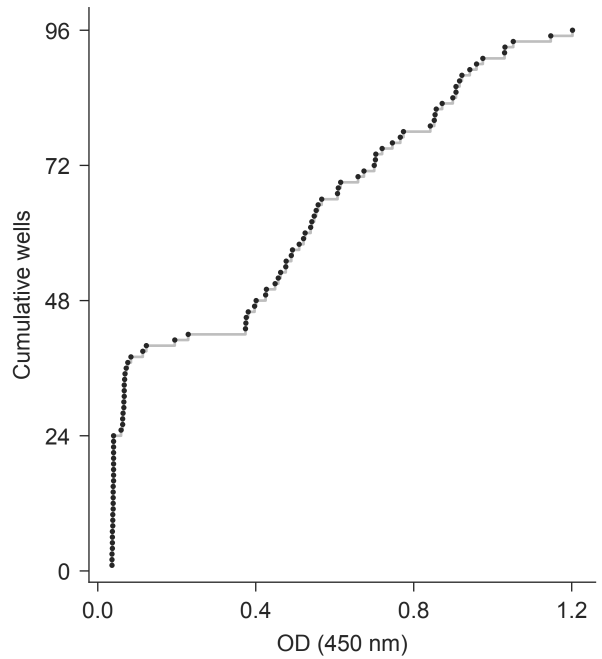


**D** elisa/plate_map.csv

| Well | Map rows | Dil. | Well | Map rows | Dil. |
|---|---|---|---|---|---|
| A1-2 | 98-99 | — | D7-8 | 140-141 | 1.0 |
| A3-4 | 100-101 | 1.0 | D9-10 | 142-143 | 1.0 |
| A5-6 | 102-103 | 1.0 | D11-12 | 144-145 | — |
| A7-8 | 104-105 | 1.0 | E1-2 | 146-147 | — |
| A9-10 | 106-107 | 1.0 | E3-4 | 148-149 | 1.0 |
| A11-12 | 108-109 | — | E5-6 | 150-151 | 1.0 |
| B1-2 | 110-111 | — | E7-8 | 152-153 | 1.0 |
| B3-4 | 112-113 | 1.0 | E9-10 | 154-155 | 1.0 |
| B5-6 | 114-115 | 1.0 | E11-12 | 156-157 | 1.0 |
| B7-8 | 116-117 | 1.0 | F1-2 | 158-159 | — |
| B9-10 | 118-119 | 1.0 | F3-4 | 160-161 | 1.0 |
| B11-12 | 120-121 | — | F5-6 | 162-163 | 1.0 |
| C1-2 | 122-123 | — | F7-8 | 164-165 | 1.0 |
| C3-4 | 124-125 | 1.0 | F9-10 | 166-167 | 1.0 |
| C5-6 | 126-127 | 1.0 | F11-12 | 168-169 | 1.0 |
| C7-8 | 128-129 | 1.0 | G1-2 | 170-171 | — |
| C9-10 | 130-131 | 1.0 | G3-4 | 172-173 | 1.0 |
| C11-12 | 132-133 | — | G5-6 | 174-175 | 1.0 |
| D1-2 | 134-135 | — | G7-12 | 176-181 | — |
| D3-4 | 136-137 | 1.0 | H1-2 | 182-183 | — |
| D5-6 | 138-139 | 1.0 | H3-12 | 184-193 | — |

**A**

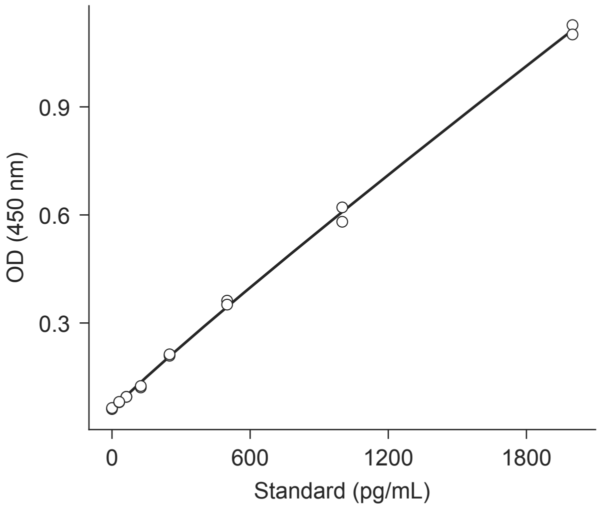


**B**

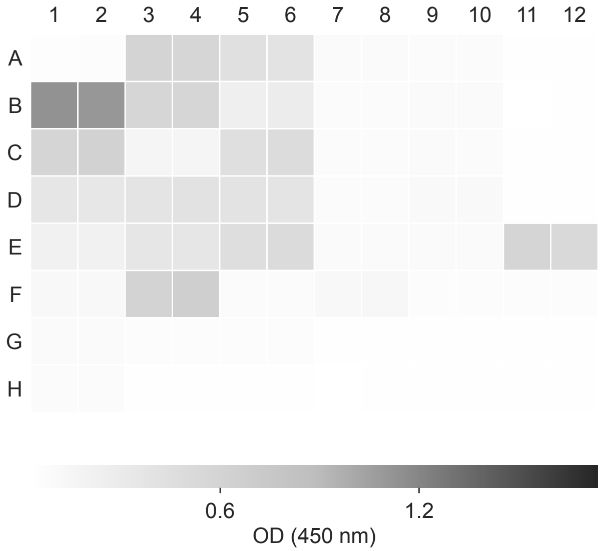


**C**

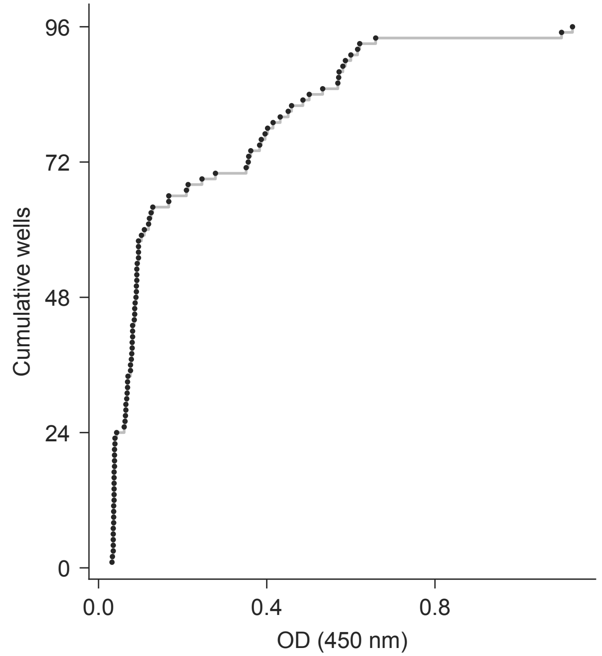


**D** elisa/plate_map.csv

| Well | Map rows | Dil. | Well | Map rows | Dil. |
|---|---|---|---|---|---|
| A1-2 | 194-195 | — | D7-8 | 236-237 | 1.0 |
| A3-4 | 196-197 | 1.0 | D9-10 | 238-239 | 1.0 |
| A5-6 | 198-199 | 1.0 | D11-12 | 240-241 | — |
| A7-8 | 200-201 | 1.0 | E1-2 | 242-243 | — |
| A9-10 | 202-203 | 1.0 | E3-4 | 244-245 | 1.0 |
| A11-12 | 204-205 | — | E5-6 | 246-247 | 1.0 |
| B1-2 | 206-207 | — | E7-8 | 248-249 | 1.0 |
| B3-4 | 208-209 | 1.0 | E9-10 | 250-251 | 1.0 |
| B5-6 | 210-211 | 1.0 | E11-12 | 252-253 | 1.0 |
| B7-8 | 212-213 | 1.0 | F1-2 | 254-255 | — |
| B9-10 | 214-215 | 1.0 | F3-4 | 256-257 | 1.0 |
| B11-12 | 216-217 | — | F5-6 | 258-259 | 1.0 |
| C1-2 | 218-219 | — | F7-8 | 260-261 | 1.0 |
| C3-4 | 220-221 | 1.0 | F9-10 | 262-263 | 1.0 |
| C5-6 | 222-223 | 1.0 | F11-12 | 264-265 | 1.0 |
| C7-8 | 224-225 | 1.0 | G1-2 | 266-267 | — |
| C9-10 | 226-227 | 1.0 | G3-4 | 268-269 | 1.0 |
| C11-12 | 228-229 | — | G5-6 | 270-271 | 1.0 |
| D1-2 | 230-231 | — | G7-12 | 272-277 | — |
| D3-4 | 232-233 | 1.0 | H1-2 | 278-279 | — |
| D5-6 | 234-235 | 1.0 | H3-12 | 280-289 | — |

**A**

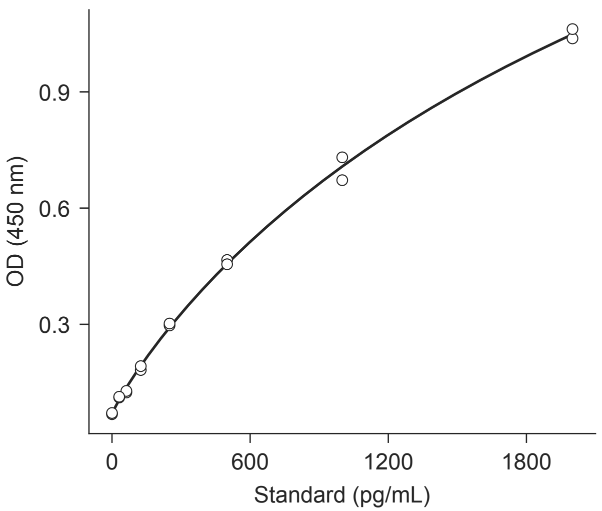


**B**

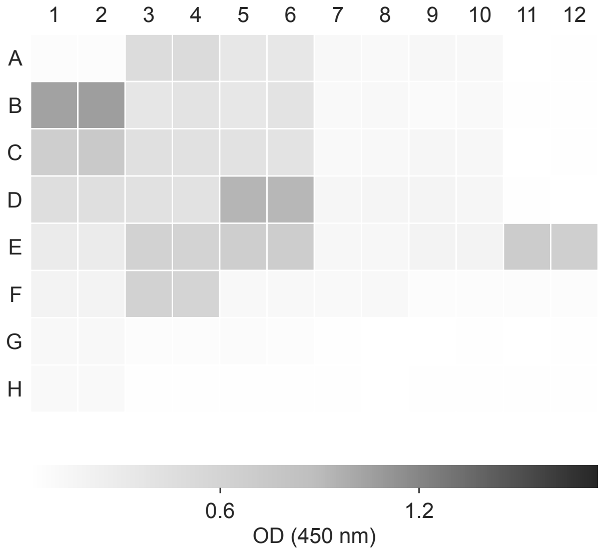


**C**

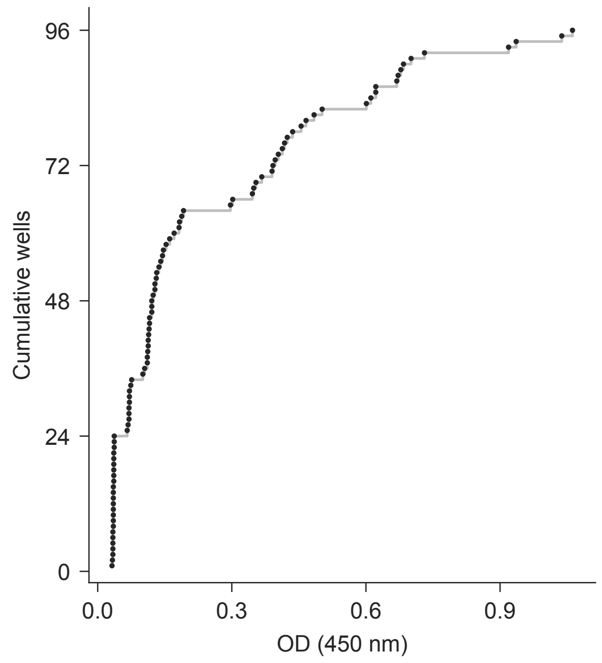


**D** elisa/plate_map.csv

| Well | Map rows | Dil. | Well | Map rows | Dil. |
|---|---|---|---|---|---|
| A1-2 | 290-291 | — | D7-8 | 332-333 | 1.0 |
| A3-4 | 292-293 | 1.0 | D9-10 | 334-335 | 1.0 |
| A5-6 | 294-295 | 1.0 | D11-12 | 336-337 | — |
| A7-8 | 296-297 | 1.0 | E1-2 | 338-339 | — |
| A9-10 | 298-299 | 1.0 | E3-4 | 340-341 | 1.0 |
| A11-12 | 300-301 | — | E5-6 | 342-343 | 1.0 |
| B1-2 | 302-303 | — | E7-8 | 344-345 | 1.0 |
| B3-4 | 304-305 | 1.0 | E9-10 | 346-347 | 1.0 |
| B5-6 | 306-307 | 1.0 | E11-12 | 348-349 | 1.0 |
| B7-8 | 308-309 | 1.0 | F1-2 | 350-351 | — |
| B9-10 | 310-311 | 1.0 | F3-4 | 352-353 | 1.0 |
| B11-12 | 312-313 | — | F5-6 | 354-355 | 1.0 |
| C1-2 | 314-315 | — | F7-8 | 356-357 | 1.0 |
| C3-4 | 316-317 | 1.0 | F9-10 | 358-359 | 1.0 |
| C5-6 | 318-319 | 1.0 | F11-12 | 360-361 | 1.0 |
| C7-8 | 320-321 | 1.0 | G1-2 | 362-363 | — |
| C9-10 | 322-323 | 1.0 | G3-4 | 364-365 | 1.0 |
| C11-12 | 324-325 | — | G5-6 | 366-367 | 1.0 |
| D1-2 | 326-327 | — | G7-12 | 368-373 | — |
| D3-4 | 328-329 | 1.0 | H1-2 | 374-375 | — |
| D5-6 | 330-331 | 1.0 | H3-12 | 376-385 | — |

**A**

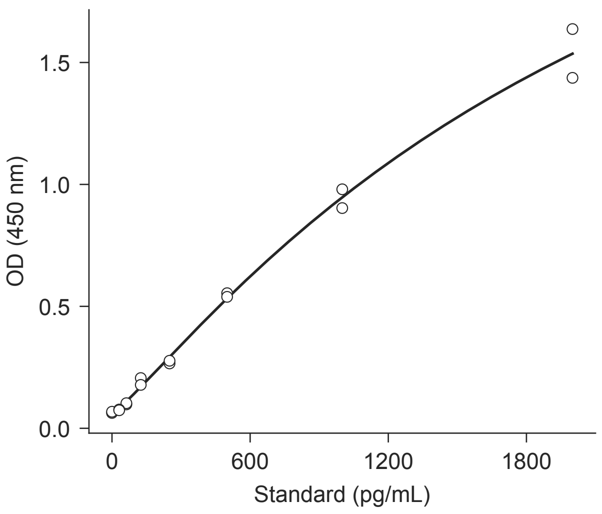


**B**

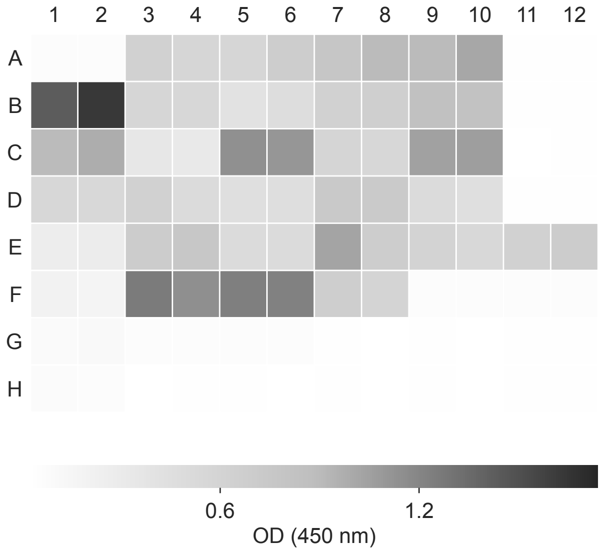


**C**

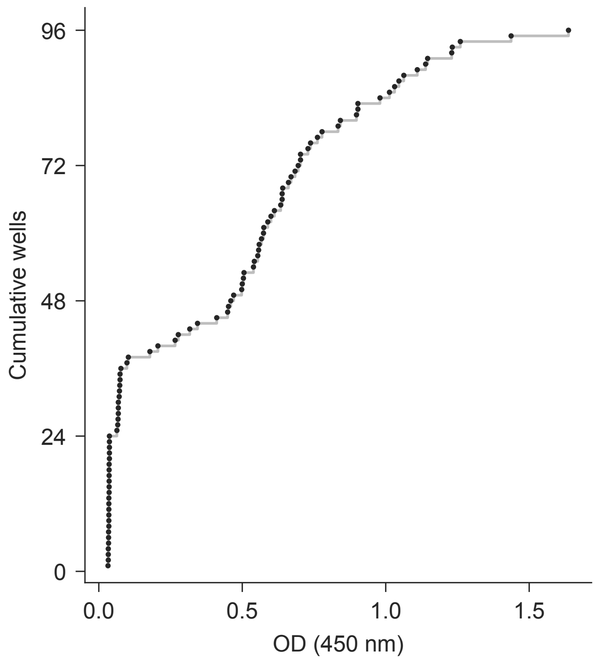


**D** elisa/plate_map.csv

| Well | Map rows | Dil. | Well | Map rows | Dil. |
|---|---|---|---|---|---|
| A1-2 | 386-387 | — | D7-8 | 428-429 | 1.0 |
| A3-4 | 388-389 | 1.0 | D9-10 | 430-431 | 1.0 |
| A5-6 | 390-391 | 1.0 | D11-12 | 432-433 | — |
| A7-8 | 392-393 | 1.0 | E1-2 | 434-435 | — |
| A9-10 | 394-395 | 1.0 | E3-4 | 436-437 | 1.0 |
| A11-12 | 396-397 | — | E5-6 | 438-439 | 1.0 |
| B1-2 | 398-399 | — | E7-8 | 440-441 | 1.0 |
| B3-4 | 400-401 | 1.0 | E9-10 | 442-443 | 1.0 |
| B5-6 | 402-403 | 1.0 | E11-12 | 444-445 | 1.0 |
| B7-8 | 404-405 | 1.0 | F1-2 | 446-447 | — |
| B9-10 | 406-407 | 1.0 | F3-4 | 448-449 | 1.0 |
| B11-12 | 408-409 | — | F5-6 | 450-451 | 1.0 |
| C1-2 | 410-411 | — | F7-8 | 452-453 | 1.0 |
| C3-4 | 412-413 | 1.0 | F9-10 | 454-455 | 1.0 |
| C5-6 | 414-415 | 1.0 | F11-12 | 456-457 | 1.0 |
| C7-8 | 416-417 | 1.0 | G1-2 | 458-459 | — |
| C9-10 | 418-419 | 1.0 | G3-4 | 460-461 | 1.0 |
| C11-12 | 420-421 | — | G5-6 | 462-463 | 1.0 |
| D1-2 | 422-423 | — | G7-12 | 464-469 | — |
| D3-4 | 424-425 | 1.0 | H1-2 | 470-471 | — |
| D5-6 | 426-427 | 1.0 | H3-12 | 472-481 | — |

**A**

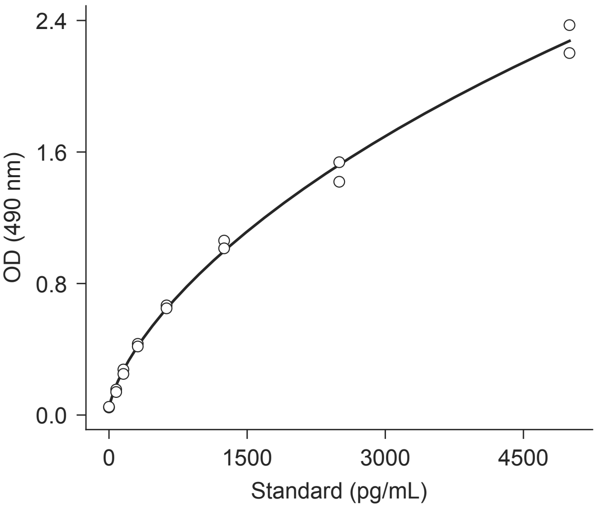


**B**

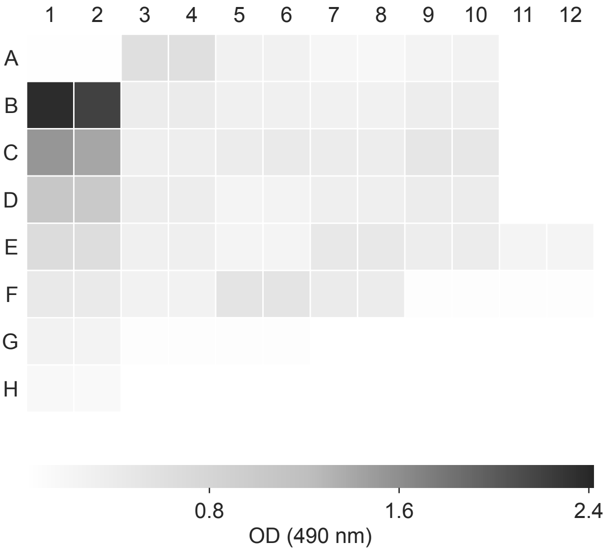


**C**

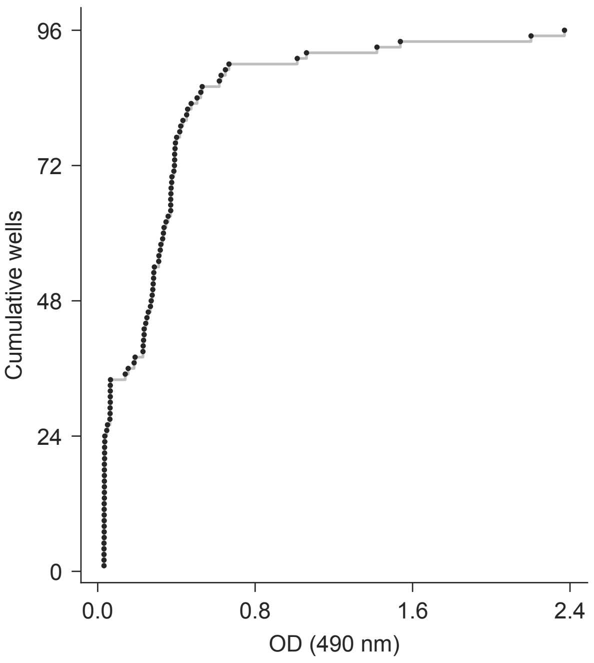


**D** elisa/plate_map.csv

| Well | Map rows | Dil. | Well | Map rows | Dil. |
|---|---|---|---|---|---|
| A1-2 | 482-483 | — | D7-8 | 524-525 | 2 |
| A3-4 | 484-485 | 2 | D9-10 | 526-527 | 2 |
| A5-6 | 486-487 | 2 | D11-12 | 528-529 | — |
| A7-8 | 488-489 | 2 | E1-2 | 530-531 | — |
| A9-10 | 490-491 | 2 | E3-4 | 532-533 | 2 |
| A11-12 | 492-493 | — | E5-6 | 534-535 | 2 |
| B1-2 | 494-495 | — | E7-8 | 536-537 | 2 |
| B3-4 | 496-497 | 2 | E9-10 | 538-539 | 2 |
| B5-6 | 498-499 | 2 | E11-12 | 540-541 | 2 |
| B7-8 | 500-501 | 2 | F1-2 | 542-543 | — |
| B9-10 | 502-503 | 2 | F3-4 | 544-545 | 2 |
| B11-12 | 504-505 | — | F5-6 | 546-547 | 2 |
| C1-2 | 506-507 | — | F7-8 | 548-549 | 2 |
| C3-4 | 508-509 | 2 | F9-10 | 550-551 | 2 |
| C5-6 | 510-511 | 2 | F11-12 | 552-553 | 2 |
| C7-8 | 512-513 | 2 | G1-2 | 554-555 | — |
| C9-10 | 514-515 | 2 | G3-4 | 556-557 | 2 |
| C11-12 | 516-517 | — | G5-6 | 558-559 | 2 |
| D1-2 | 518-519 | — | G7-12 | 560-565 | — |
| D3-4 | 520-521 | 2 | H1-2 | 566-567 | — |
| D5-6 | 522-523 | 2 | H3-12 | 568-577 | — |

**A**

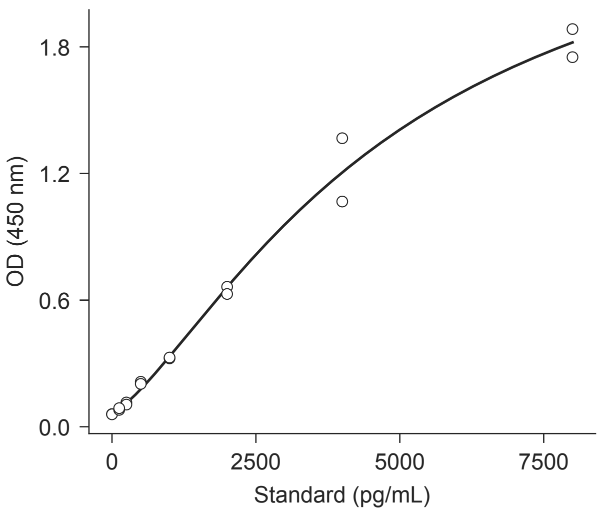


**B**

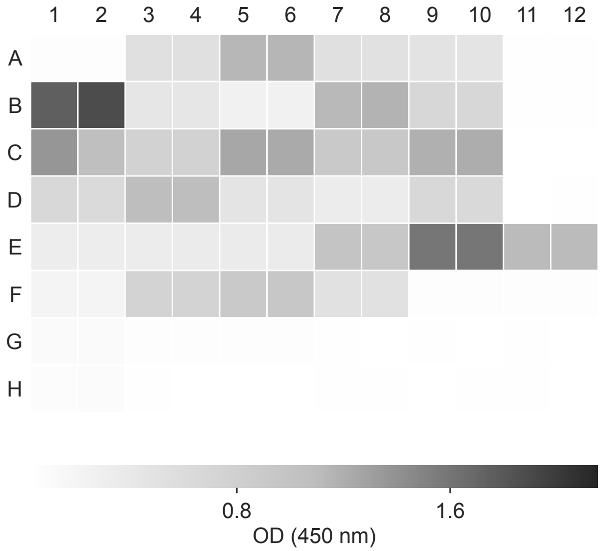


**C**

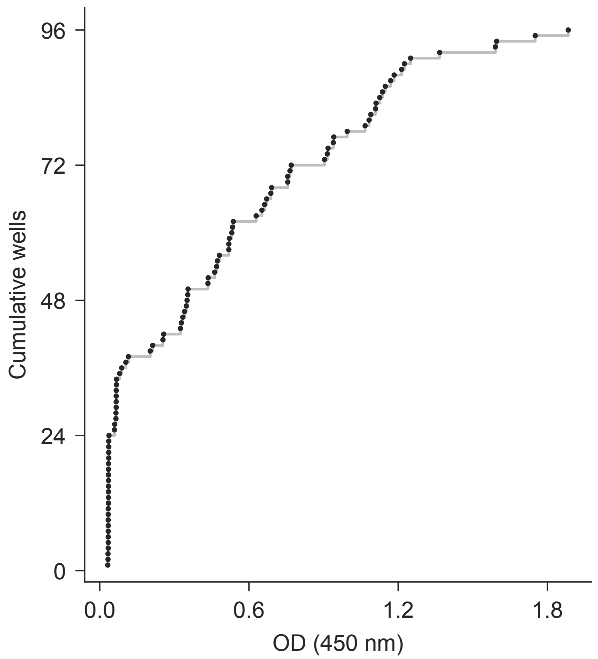


**D** elisa/plate_map.csv

| Well | Map rows | Dil. | Well | Map rows | Dil. |
|---|---|---|---|---|---|
| A1-2 | 578-579 | — | D7-8 | 620-621 | 2 |
| A3-4 | 580-581 | 2 | D9-10 | 622-623 | 2 |
| A5-6 | 582-583 | 2 | D11-12 | 624-625 | — |
| A7-8 | 584-585 | 2 | E1-2 | 626-627 | — |
| A9-10 | 586-587 | 2 | E3-4 | 628-629 | 2 |
| A11-12 | 588-589 | — | E5-6 | 630-631 | 2 |
| B1-2 | 590-591 | — | E7-8 | 632-633 | 2 |
| B3-4 | 592-593 | 2 | E9-10 | 634-635 | 2 |
| B5-6 | 594-595 | 2 | E11-12 | 636-637 | 2 |
| B7-8 | 596-597 | 2 | F1-2 | 638-639 | — |
| B9-10 | 598-599 | 2 | F3-4 | 640-641 | 2 |
| B11-12 | 600-601 | — | F5-6 | 642-643 | 2 |
| C1-2 | 602-603 | — | F7-8 | 644-645 | 2 |
| C3-4 | 604-605 | 2 | F9-10 | 646-647 | 2 |
| C5-6 | 606-607 | 2 | F11-12 | 648-649 | 2 |
| C7-8 | 608-609 | 2 | G1-2 | 650-651 | — |
| C9-10 | 610-611 | 2 | G3-4 | 652-653 | 2 |
| C11-12 | 612-613 | — | G5-6 | 654-655 | 2 |
| D1-2 | 614-615 | — | G7-12 | 656-661 | — |
| D3-4 | 616-617 | 2 | H1-2 | 662-663 | — |
| D5-6 | 618-619 | 2 | H3-12 | 664-673 | — |

**A**

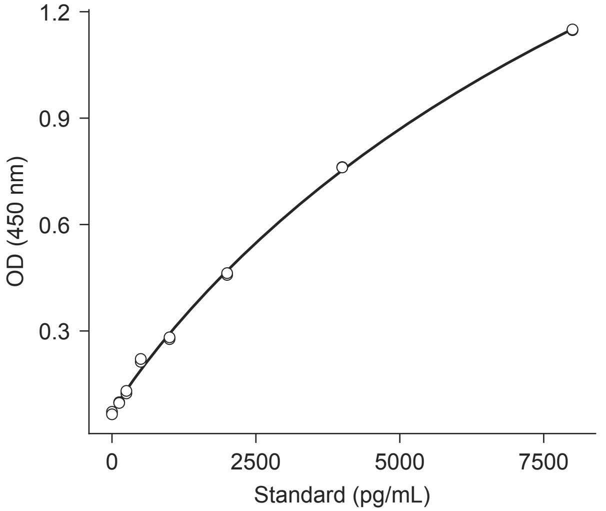


**B**

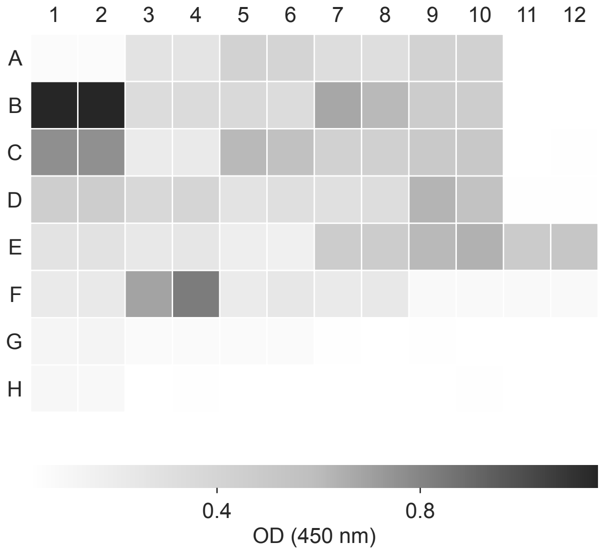


**C**

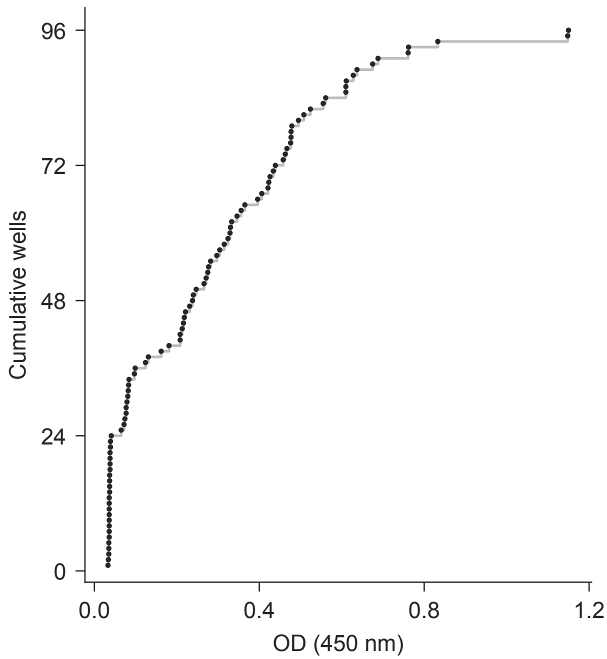


**D** elisa/plate_map.csv

| Well | Map rows | Dil. | Well | Map rows | Dil. |
|---|---|---|---|---|---|
| A1-2 | 674-675 | — | D7-8 | 716-717 | 1.0 |
| A3-4 | 676-677 | 1.0 | D9-10 | 718-719 | 1.0 |
| A5-6 | 678-679 | 1.0 | D11-12 | 720-721 | — |
| A7-8 | 680-681 | 1.0 | E1-2 | 722-723 | — |
| A9-10 | 682-683 | 1.0 | E3-4 | 724-725 | 1.0 |
| A11-12 | 684-685 | — | E5-6 | 726-727 | 1.0 |
| B1-2 | 686-687 | — | E7-8 | 728-729 | 1.0 |
| B3-4 | 688-689 | 1.0 | E9-10 | 730-731 | 1.0 |
| B5-6 | 690-691 | 1.0 | E11-12 | 732-733 | 1.0 |
| B7-8 | 692-693 | 1.0 | F1-2 | 734-735 | — |
| B9-10 | 694-695 | 1.0 | F3-4 | 736-737 | 1.0 |
| B11-12 | 696-697 | — | F5-6 | 738-739 | 1.0 |
| C1-2 | 698-699 | — | F7-8 | 740-741 | 1.0 |
| C3-4 | 700-701 | 1.0 | F9-10 | 742-743 | 1.0 |
| C5-6 | 702-703 | 1.0 | F11-12 | 744-745 | 1.0 |
| C7-8 | 704-705 | 1.0 | G1-2 | 746-747 | — |
| C9-10 | 706-707 | 1.0 | G3-4 | 748-749 | 1.0 |
| C11-12 | 708-709 | — | G5-6 | 750-751 | 1.0 |
| D1-2 | 710-711 | — | G7-12 | 752-757 | — |
| D3-4 | 712-713 | 1.0 | H1-2 | 758-759 | — |
| D5-6 | 714-715 | 1.0 | H3-12 | 760-769 | — |

**A**

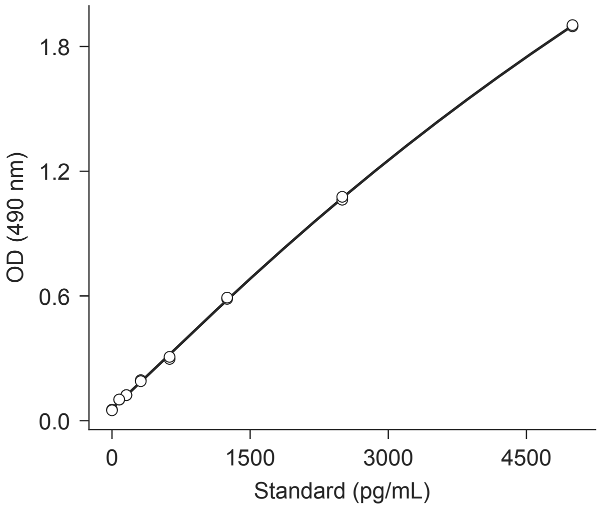


**B**

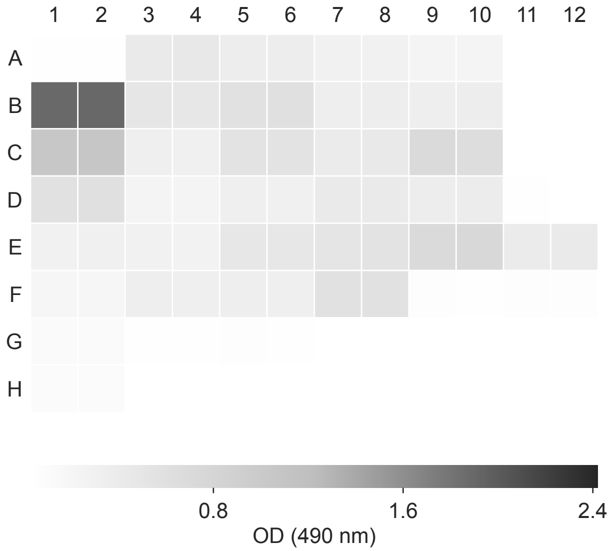


**C**

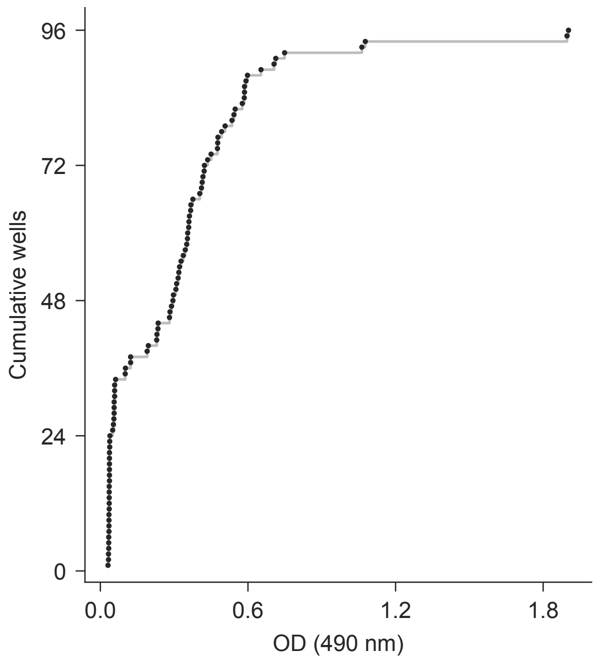


**D** elisa/plate_map.csv

| Well | Map rows | Dil. | Well | Map rows | Dil. |
|---|---|---|---|---|---|
| A1-2 | 770-771 | — | D7-8 | 812-813 | 2 |
| A3-4 | 772-773 | 2 | D9-10 | 814-815 | 2 |
| A5-6 | 774-775 | 2 | D11-12 | 816-817 | — |
| A7-8 | 776-777 | 2 | E1-2 | 818-819 | — |
| A9-10 | 778-779 | 2 | E3-4 | 820-821 | 2 |
| A11-12 | 780-781 | — | E5-6 | 822-823 | 2 |
| B1-2 | 782-783 | — | E7-8 | 824-825 | 2 |
| B3-4 | 784-785 | 2 | E9-10 | 826-827 | 2 |
| B5-6 | 786-787 | 2 | E11-12 | 828-829 | 2 |
| B7-8 | 788-789 | 2 | F1-2 | 830-831 | — |
| B9-10 | 790-791 | 2 | F3-4 | 832-833 | 2 |
| B11-12 | 792-793 | — | F5-6 | 834-835 | 2 |
| C1-2 | 794-795 | — | F7-8 | 836-837 | 2 |
| C3-4 | 796-797 | 2 | F9-10 | 838-839 | 2 |
| C5-6 | 798-799 | 2 | F11-12 | 840-841 | 2 |
| C7-8 | 800-801 | 2 | G1-2 | 842-843 | — |
| C9-10 | 802-803 | 2 | G3-4 | 844-845 | 2 |
| C11-12 | 804-805 | — | G5-6 | 846-847 | 2 |
| D1-2 | 806-807 | — | G7-12 | 848-853 | — |
| D3-4 | 808-809 | 2 | H1-2 | 854-855 | — |
| D5-6 | 810-811 | 2 | H3-12 | 856-865 | — |

**A**

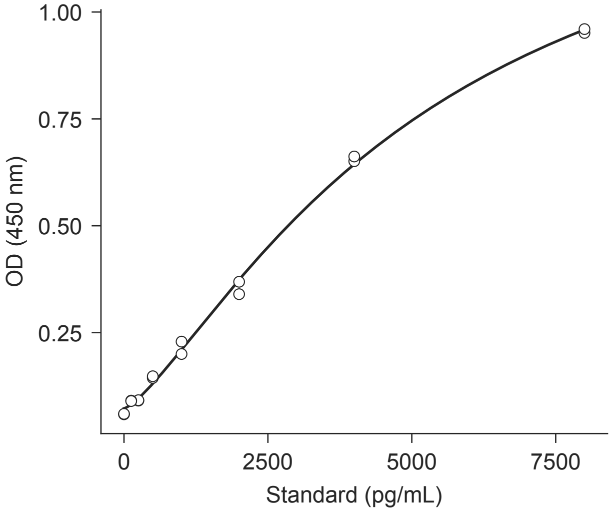


**B**

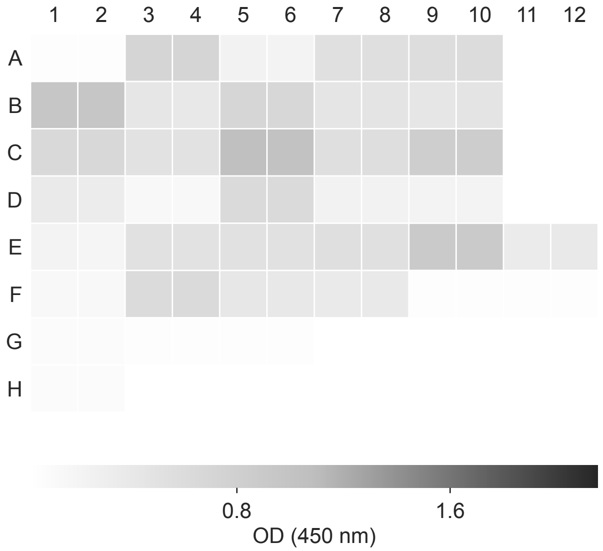


**C**

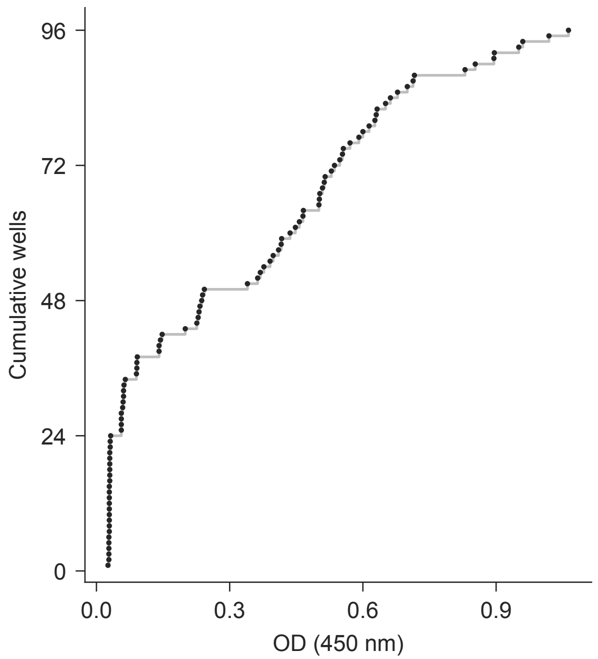


**D** elisa/plate_map.csv

| Well | Map rows | Dil. | Well | Map rows | Dil. |
|---|---|---|---|---|---|
| A1-2 | 866-867 | — | D7-8 | 908-909 | 2 |
| A3-4 | 868-869 | 2 | D9-10 | 910-911 | 2 |
| A5-6 | 870-871 | 2 | D11-12 | 912-913 | — |
| A7-8 | 872-873 | 2 | E1-2 | 914-915 | — |
| A9-10 | 874-875 | 2 | E3-4 | 916-917 | 2 |
| A11-12 | 876-877 | — | E5-6 | 918-919 | 2 |
| B1-2 | 878-879 | — | E7-8 | 920-921 | 2 |
| B3-4 | 880-881 | 2 | E9-10 | 922-923 | 2 |
| B5-6 | 882-883 | 2 | E11-12 | 924-925 | 2 |
| B7-8 | 884-885 | 2 | F1-2 | 926-927 | — |
| B9-10 | 886-887 | 2 | F3-4 | 928-929 | 2 |
| B11-12 | 888-889 | — | F5-6 | 930-931 | 2 |
| C1-2 | 890-891 | — | F7-8 | 932-933 | 2 |
| C3-4 | 892-893 | 2 | F9-10 | 934-935 | 2 |
| C5-6 | 894-895 | 2 | F11-12 | 936-937 | 2 |
| C7-8 | 896-897 | 2 | G1-2 | 938-939 | — |
| C9-10 | 898-899 | 2 | G3-4 | 940-941 | 2 |
| C11-12 | 900-901 | — | G5-6 | 942-943 | 2 |
| D1-2 | 902-903 | — | G7-12 | 944-949 | — |
| D3-4 | 904-905 | 2 | H1-2 | 950-951 | — |
| D5-6 | 906-907 | 2 | H3-12 | 952-961 | — |

**A**

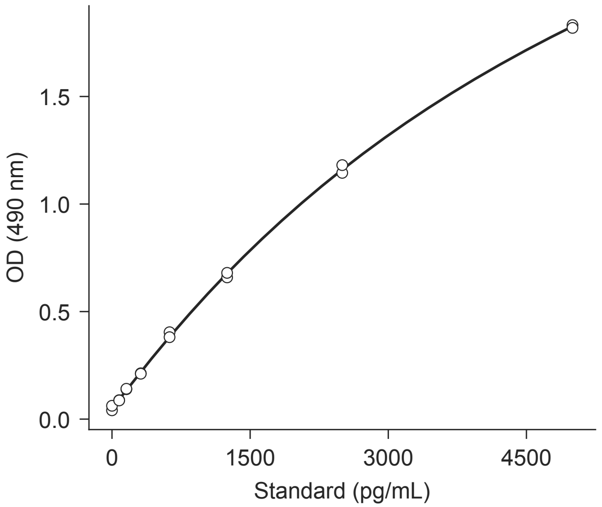


**B**

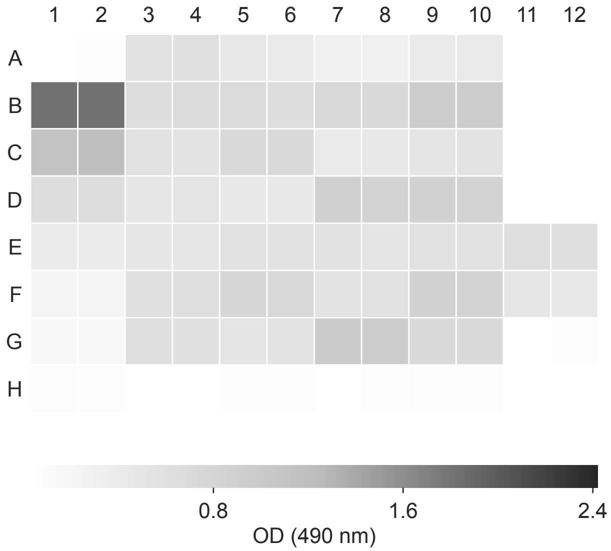


**C**

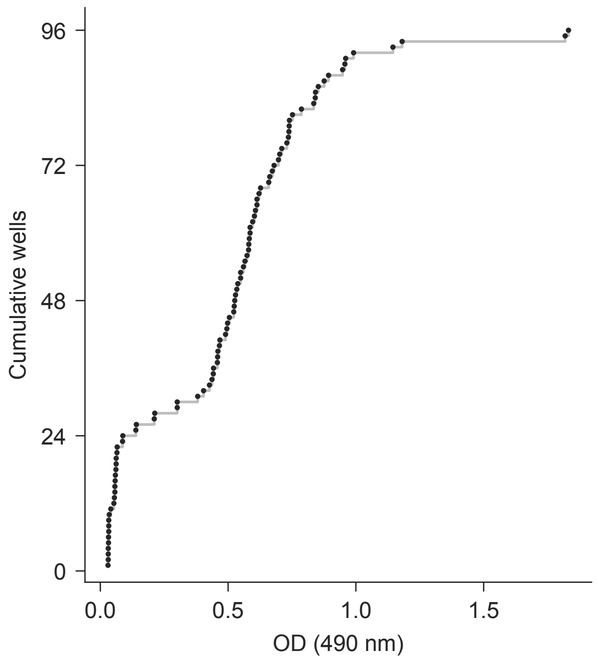


**D** elisa/plate_map.csv

| Well | Map rows | Dil. | Well | Map rows | Dil. |
|---|---|---|---|---|---|
| A1-2 | 962-963 | — | E1-2 | 1010-1011 | — |
| A3-4 | 964-965 | 2 | E3-4 | 1012-1013 | 2 |
| A5-6 | 966-967 | 2 | E5-6 | 1014-1015 | 2 |
| A7-8 | 968-969 | 2 | E7-8 | 1016-1017 | 2 |
| A9-10 | 970-971 | 2 | E9-10 | 1018-1019 | 2 |
| A11-12 | 972-973 | — | E11-12 | 1020-1021 | 2 |
| B1-2 | 974-975 | — | F1-2 | 1022-1023 | — |
| B3-4 | 976-977 | 2 | F3-4 | 1024-1025 | 2 |
| B5-6 | 978-979 | 2 | F5-6 | 1026-1027 | 2 |
| B7-8 | 980-981 | 2 | F7-8 | 1028-1029 | 2 |
| B9-10 | 982-983 | 2 | F9-10 | 1030-1031 | 2 |
| B11-12 | 984-985 | — | F11-12 | 1032-1033 | 2 |
| C1-2 | 986-987 | — | G1-2 | 1034-1035 | — |
| C3-4 | 988-989 | 2 | G3-4 | 1036-1037 | 2 |
| C5-6 | 990-991 | 2 | G5-6 | 1038-1039 | 2 |
| C7-8 | 992-993 | 2 | G7-8 | 1040-1041 | 2 |
| C9-10 | 994-995 | 2 | G9-10 | 1042-1043 | 2 |
| C11-12 | 996-997 | — | G11-12 | 1044-1045 | 2 |
| D1-2 | 998-999 | — | H1-2 | 1046-1047 | — |
| D3-4 | 1000-1001 | 2 | H3-4 | 1048-1049 | 2 |
| D5-6 | 1002-1003 | 2 | H5-6 | 1050-1051 | 2 |
| D7-8 | 1004-1005 | 2 | H7-8 | 1052-1053 | 2 |
| D9-10 | 1006-1007 | 2 | H9-10 | 1054-1055 | 2 |
| D11-12 | 1008-1009 | — | H11-12 | 1056-1057 | — |

**A**

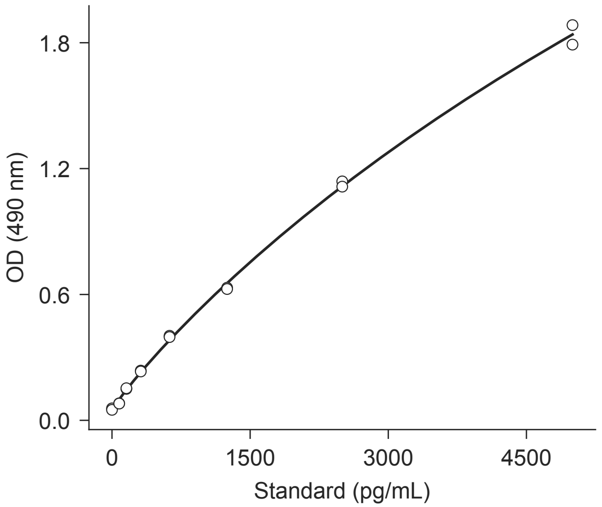


**B**

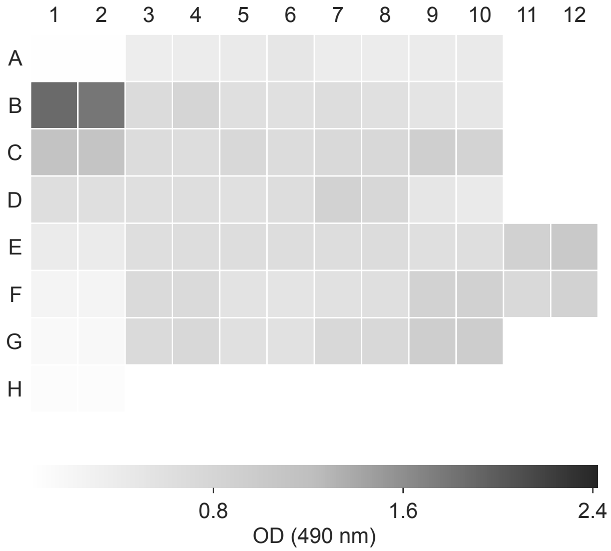


**C**

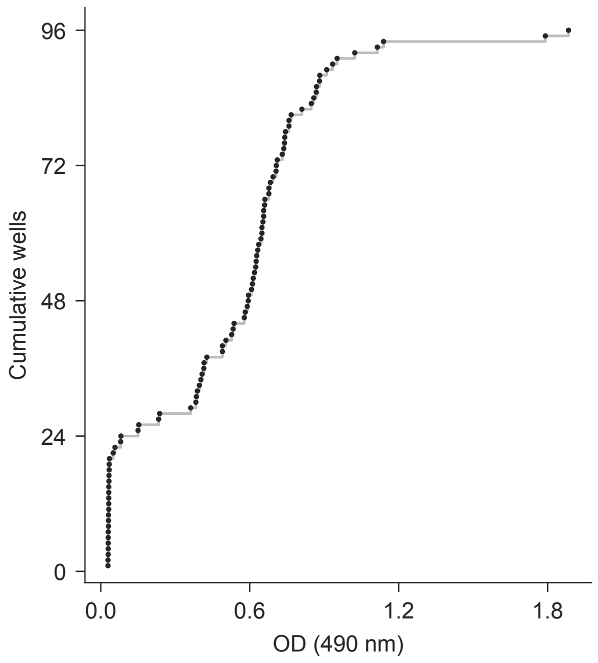


**D** elisa/plate_map.csv

| Well | Map rows | Dil. | Well | Map rows | Dil. |
|---|---|---|---|---|---|
| A1-2 | 1058-1059 | — | E1-2 | 1106-1107 | — |
| A3-4 | 1060-1061 | 2 | E3-4 | 1108-1109 | 2 |
| A5-6 | 1062-1063 | 2 | E5-6 | 1110-1111 | 2 |
| A7-8 | 1064-1065 | 2 | E7-8 | 1112-1113 | 2 |
| A9-10 | 1066-1067 | 2 | E9-10 | 1114-1115 | 2 |
| A11-12 | 1068-1069 | — | E11-12 | 1116-1117 | 2 |
| B1-2 | 1070-1071 | — | F1-2 | 1118-1119 | — |
| B3-4 | 1072-1073 | 2 | F3-4 | 1120-1121 | 2 |
| B5-6 | 1074-1075 | 2 | F5-6 | 1122-1123 | 2 |
| B7-8 | 1076-1077 | 2 | F7-8 | 1124-1125 | 2 |
| B9-10 | 1078-1079 | 2 | F9-10 | 1126-1127 | 2 |
| B11-12 | 1080-1081 | — | F11-12 | 1128-1129 | 2 |
| C1-2 | 1082-1083 | — | G1-2 | 1130-1131 | — |
| C3-4 | 1084-1085 | 2 | G3-4 | 1132-1133 | 2 |
| C5-6 | 1086-1087 | 2 | G5-6 | 1134-1135 | 2 |
| C7-8 | 1088-1089 | 2 | G7-8 | 1136-1137 | 2 |
| C9-10 | 1090-1091 | 2 | G9-10 | 1138-1139 | 2 |
| C11-12 | 1092-1093 | — | G11-12 | 1140-1141 | 2 |
| D1-2 | 1094-1095 | — | H1-2 | 1142-1143 | — |
| D3-4 | 1096-1097 | 2 | H3-4 | 1144-1145 | 2 |
| D5-6 | 1098-1099 | 2 | H5-6 | 1146-1147 | 2 |
| D7-8 | 1100-1101 | 2 | H7-8 | 1148-1149 | 2 |
| D9-10 | 1102-1103 | 2 | H9-10 | 1150-1151 | 2 |
| D11-12 | 1104-1105 | — | H11-12 | 1152-1153 | — |

**A**

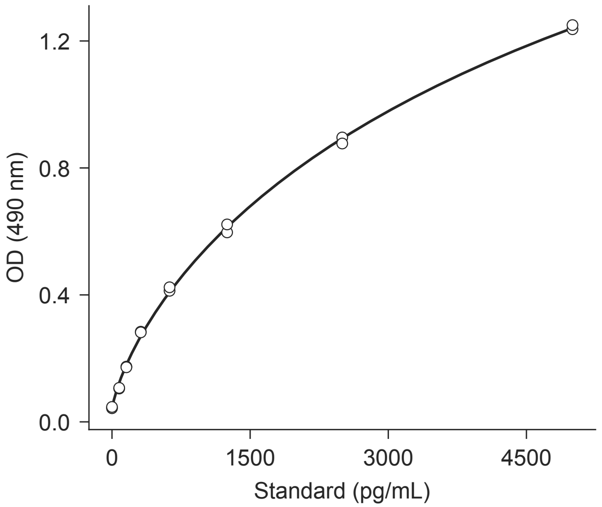


**B**

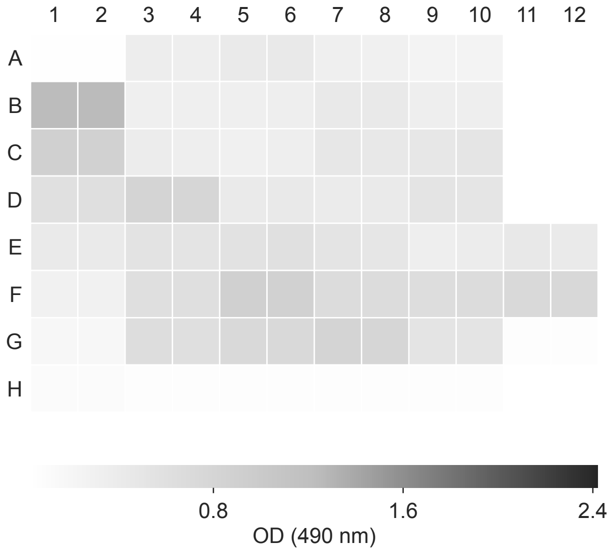


**C**

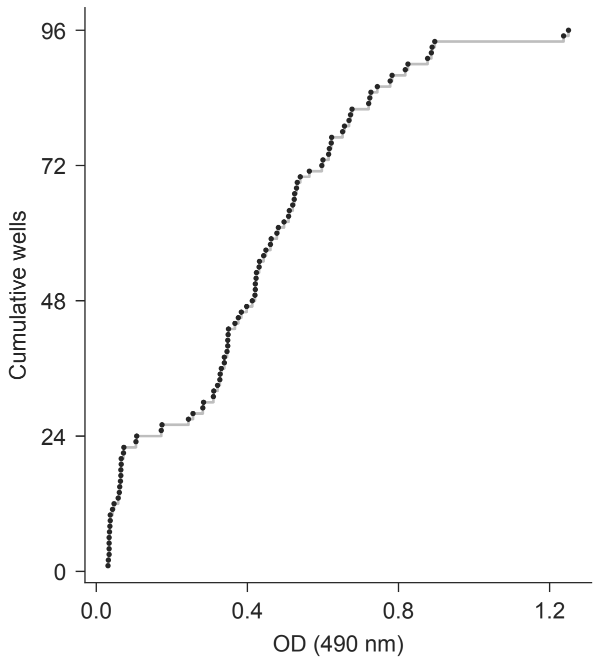


**D** elisa/plate_map.csv

| Well | Map rows | Dil. | Well | Map rows | Dil. |
|---|---|---|---|---|---|
| A1-2 | 1154-1155 | — | E1-2 | 1202-1203 | — |
| A3-4 | 1156-1157 | 2 | E3-4 | 1204-1205 | 2 |
| A5-6 | 1158-1159 | 2 | E5-6 | 1206-1207 | 2 |
| A7-8 | 1160-1161 | 2 | E7-8 | 1208-1209 | 2 |
| A9-10 | 1162-1163 | 2 | E9-10 | 1210-1211 | 2 |
| A11-12 | 1164-1165 | — | E11-12 | 1212-1213 | 2 |
| B1-2 | 1166-1167 | — | F1-2 | 1214-1215 | — |
| B3-4 | 1168-1169 | 2 | F3-4 | 1216-1217 | 2 |
| B5-6 | 1170-1171 | 2 | F5-6 | 1218-1219 | 2 |
| B7-8 | 1172-1173 | 2 | F7-8 | 1220-1221 | 2 |
| B9-10 | 1174-1175 | 2 | F9-10 | 1222-1223 | 2 |
| B11-12 | 1176-1177 | — | F11-12 | 1224-1225 | 2 |
| C1-2 | 1178-1179 | — | G1-2 | 1226-1227 | — |
| C3-4 | 1180-1181 | 2 | G3-4 | 1228-1229 | 2 |
| C5-6 | 1182-1183 | 2 | G5-6 | 1230-1231 | 2 |
| C7-8 | 1184-1185 | 2 | G7-8 | 1232-1233 | 2 |
| C9-10 | 1186-1187 | 2 | G9-10 | 1234-1235 | 2 |
| C11-12 | 1188-1189 | — | G11-12 | 1236-1237 | 2 |
| D1-2 | 1190-1191 | — | H1-2 | 1238-1239 | — |
| D3-4 | 1192-1193 | 2 | H3-4 | 1240-1241 | 2 |
| D5-6 | 1194-1195 | 2 | H5-6 | 1242-1243 | 2 |
| D7-8 | 1196-1197 | 2 | H7-8 | 1244-1245 | 2 |
| D9-10 | 1198-1199 | 2 | H9-10 | 1246-1247 | 2 |
| D11-12 | 1200-1201 | — | H11-12 | 1248-1249 | — |

20260803_COL_RET_T336_4.txt

Synergy H1 | 8/3/2026 | retention | COL_MAIN_T336_4

**A**

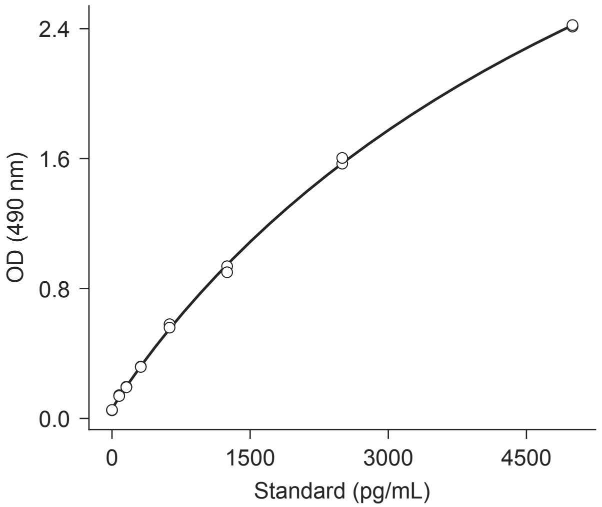


**B**

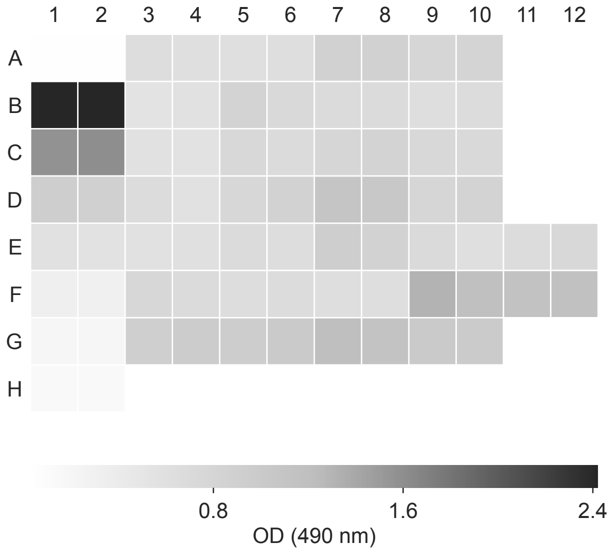


**C**

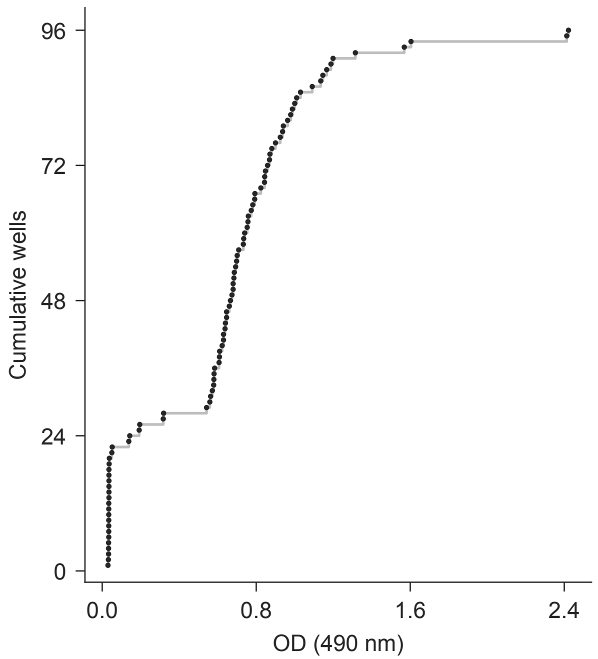


**D** elisa/plate_map.csv

| Well | Map rows | Dil. | Well | Map rows | Dil. |
|---|---|---|---|---|---|
| A1-2 | 1250-1251 | — | E1-2 | 1298-1299 | — |
| A3-4 | 1252-1253 | 2 | E3-4 | 1300-1301 | 2 |
| A5-6 | 1254-1255 | 2 | E5-6 | 1302-1303 | 2 |
| A7-8 | 1256-1257 | 2 | E7-8 | 1304-1305 | 2 |
| A9-10 | 1258-1259 | 2 | E9-10 | 1306-1307 | 2 |
| A11-12 | 1260-1261 | — | E11-12 | 1308-1309 | 2 |
| B1-2 | 1262-1263 | — | F1-2 | 1310-1311 | — |
| B3-4 | 1264-1265 | 2 | F3-4 | 1312-1313 | 2 |
| B5-6 | 1266-1267 | 2 | F5-6 | 1314-1315 | 2 |
| B7-8 | 1268-1269 | 2 | F7-8 | 1316-1317 | 2 |
| B9-10 | 1270-1271 | 2 | F9-10 | 1318-1319 | 2 |
| B11-12 | 1272-1273 | — | F11-12 | 1320-1321 | 2 |
| C1-2 | 1274-1275 | — | G1-2 | 1322-1323 | — |
| C3-4 | 1276-1277 | 2 | G3-4 | 1324-1325 | 2 |
| C5-6 | 1278-1279 | 2 | G5-6 | 1326-1327 | 2 |
| C7-8 | 1280-1281 | 2 | G7-8 | 1328-1329 | 2 |
| C9-10 | 1282-1283 | 2 | G9-10 | 1330-1331 | 2 |
| C11-12 | 1284-1285 | — | G11-12 | 1332-1333 | 2 |
| D1-2 | 1286-1287 | — | H1-2 | 1334-1335 | — |
| D3-4 | 1288-1289 | 2 | H3-4 | 1336-1337 | 2 |
| D5-6 | 1290-1291 | 2 | H5-6 | 1338-1339 | 2 |
| D7-8 | 1292-1293 | 2 | H7-8 | 1340-1341 | 2 |
| D9-10 | 1294-1295 | 2 | H9-10 | 1342-1343 | 2 |
| D11-12 | 1296-1297 | — | H11-12 | 1344-1345 | — |

20260804_LUM_RET_T336_1.txt

Synergy H1 | 8/4/2026 | retention | LUM_MAIN_T336_1

**A**

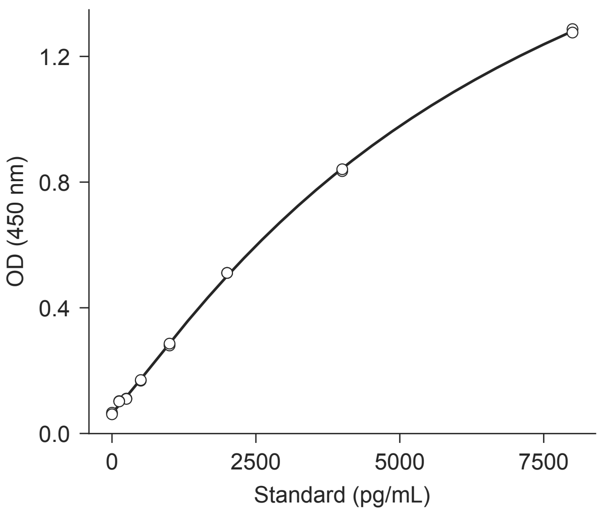


**B**

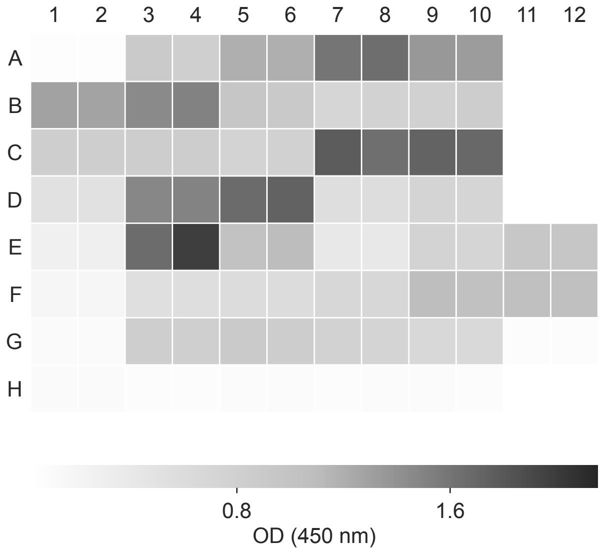


**C**

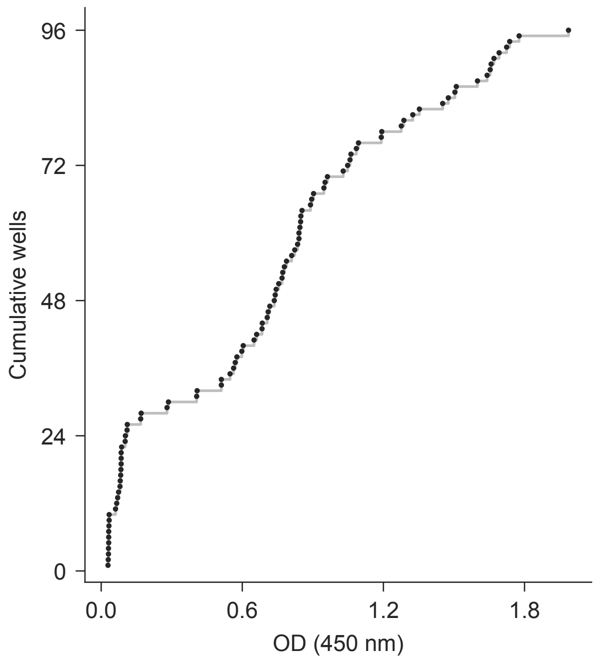


**D** elisa/plate_map.csv

| Well | Map rows | Dil. | Well | Map rows | Dil. |
|---|---|---|---|---|---|
| A1-2 | 1346-1347 | — | E1-2 | 1394-1395 | — |
| A3-4 | 1348-1349 | 2 | E3-4 | 1396-1397 | 2 |
| A5-6 | 1350-1351 | 2 | E5-6 | 1398-1399 | 2 |
| A7-8 | 1352-1353 | 2 | E7-8 | 1400-1401 | 2 |
| A9-10 | 1354-1355 | 2 | E9-10 | 1402-1403 | 2 |
| A11-12 | 1356-1357 | — | E11-12 | 1404-1405 | 2 |
| B1-2 | 1358-1359 | — | F1-2 | 1406-1407 | — |
| B3-4 | 1360-1361 | 2 | F3-4 | 1408-1409 | 2 |
| B5-6 | 1362-1363 | 2 | F5-6 | 1410-1411 | 2 |
| B7-8 | 1364-1365 | 2 | F7-8 | 1412-1413 | 2 |
| B9-10 | 1366-1367 | 2 | F9-10 | 1414-1415 | 2 |
| B11-12 | 1368-1369 | — | F11-12 | 1416-1417 | 2 |
| C1-2 | 1370-1371 | — | G1-2 | 1418-1419 | — |
| C3-4 | 1372-1373 | 2 | G3-4 | 1420-1421 | 2 |
| C5-6 | 1374-1375 | 2 | G5-6 | 1422-1423 | 2 |
| C7-8 | 1376-1377 | 2 | G7-8 | 1424-1425 | 2 |
| C9-10 | 1378-1379 | 2 | G9-10 | 1426-1427 | 2 |
| C11-12 | 1380-1381 | — | G11-12 | 1428-1429 | 2 |
| D1-2 | 1382-1383 | — | H1-2 | 1430-1431 | — |
| D3-4 | 1384-1385 | 2 | H3-4 | 1432-1433 | 2 |
| D5-6 | 1386-1387 | 2 | H5-6 | 1434-1435 | 2 |
| D7-8 | 1388-1389 | 2 | H7-8 | 1436-1437 | 2 |
| D9-10 | 1390-1391 | 2 | H9-10 | 1438-1439 | 2 |
| D11-12 | 1392-1393 | — | H11-12 | 1440-1441 | — |

**A**

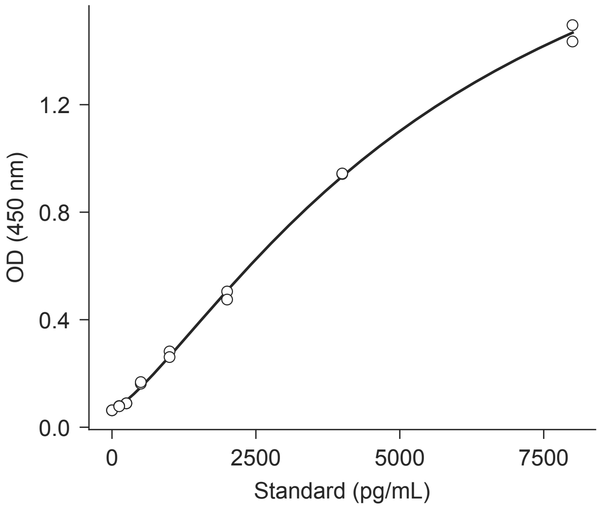


**B**

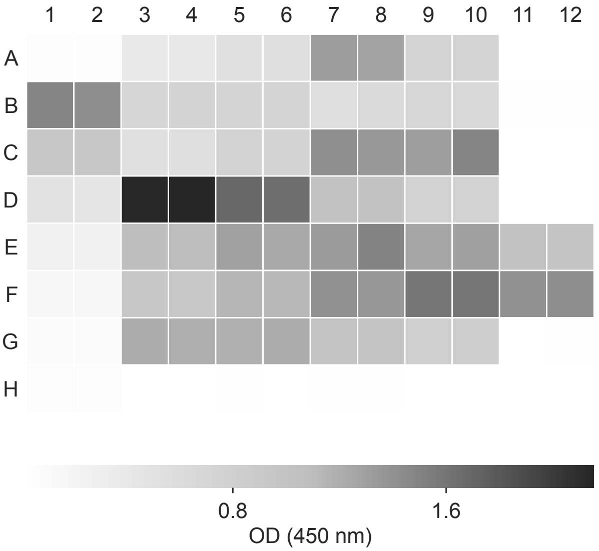


**C**

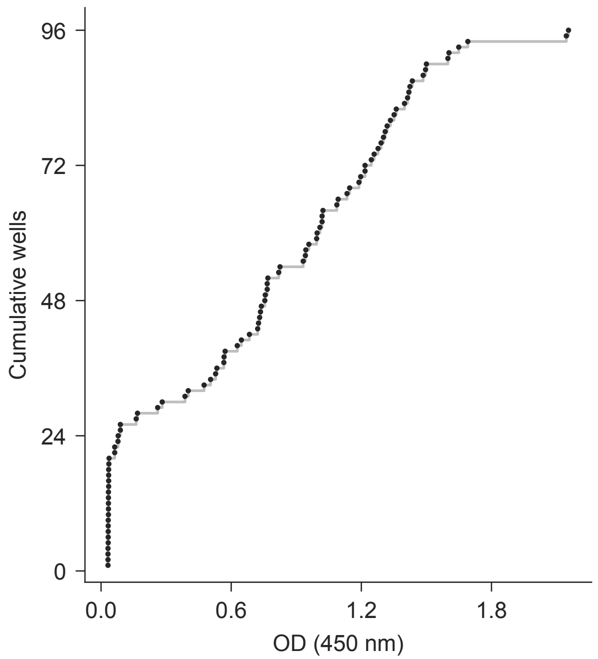


**D** elisa/plate_map.csv

| Well | Map rows | Dil. | Well | Map rows | Dil. |
|---|---|---|---|---|---|
| A1-2 | 1442-1443 | — | E1-2 | 1490-1491 | — |
| A3-4 | 1444-1445 | 2 | E3-4 | 1492-1493 | 2 |
| A5-6 | 1446-1447 | 2 | E5-6 | 1494-1495 | 2 |
| A7-8 | 1448-1449 | 2 | E7-8 | 1496-1497 | 2 |
| A9-10 | 1450-1451 | 2 | E9-10 | 1498-1499 | 2 |
| A11-12 | 1452-1453 | — | E11-12 | 1500-1501 | 2 |
| B1-2 | 1454-1455 | — | F1-2 | 1502-1503 | — |
| B3-4 | 1456-1457 | 2 | F3-4 | 1504-1505 | 2 |
| B5-6 | 1458-1459 | 2 | F5-6 | 1506-1507 | 2 |
| B7-8 | 1460-1461 | 2 | F7-8 | 1508-1509 | 2 |
| B9-10 | 1462-1463 | 2 | F9-10 | 1510-1511 | 2 |
| B11-12 | 1464-1465 | — | F11-12 | 1512-1513 | 2 |
| C1-2 | 1466-1467 | — | G1-2 | 1514-1515 | — |
| C3-4 | 1468-1469 | 2 | G3-4 | 1516-1517 | 2 |
| C5-6 | 1470-1471 | 2 | G5-6 | 1518-1519 | 2 |
| C7-8 | 1472-1473 | 2 | G7-8 | 1520-1521 | 2 |
| C9-10 | 1474-1475 | 2 | G9-10 | 1522-1523 | 2 |
| C11-12 | 1476-1477 | — | G11-12 | 1524-1525 | 2 |
| D1-2 | 1478-1479 | — | H1-2 | 1526-1527 | — |
| D3-4 | 1480-1481 | 2 | H3-4 | 1528-1529 | 2 |
| D5-6 | 1482-1483 | 2 | H5-6 | 1530-1531 | 2 |
| D7-8 | 1484-1485 | 2 | H7-8 | 1532-1533 | 2 |
| D9-10 | 1486-1487 | 2 | H9-10 | 1534-1535 | 2 |
| D11-12 | 1488-1489 | — | H11-12 | 1536-1537 | — |

**A**

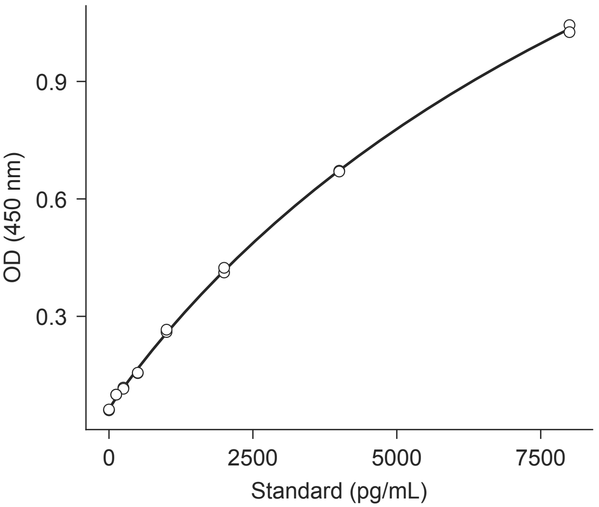


**B**

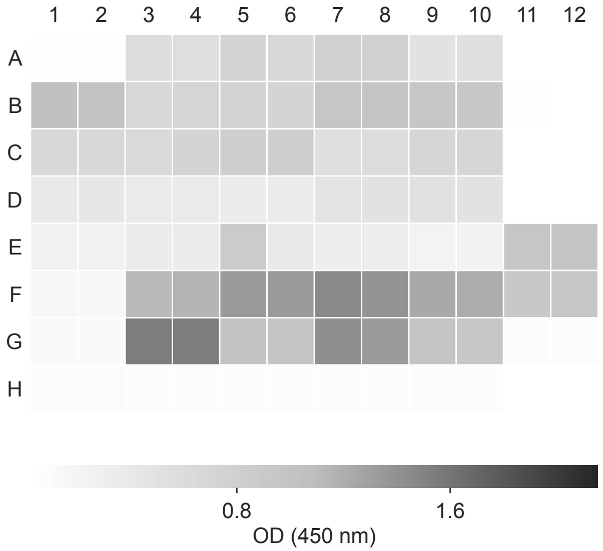


**C**

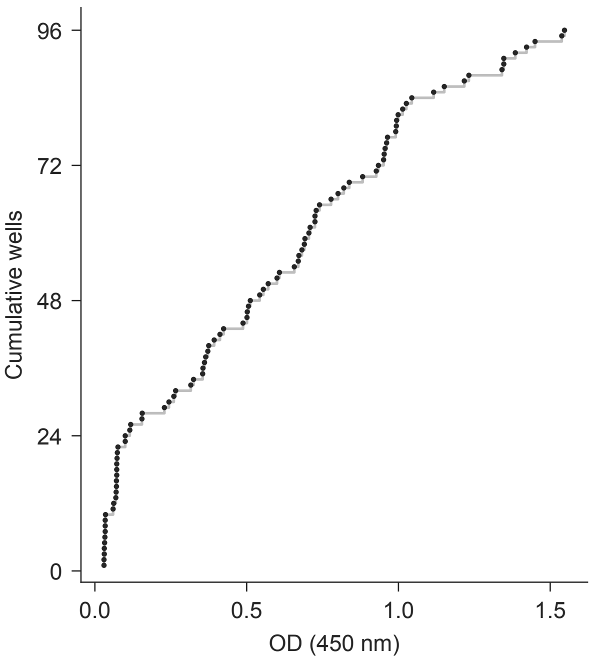


**D** elisa/plate_map.csv

| Well | Map rows | Dil. | Well | Map rows | Dil. |
|---|---|---|---|---|---|
| A1-2 | 1538-1539 | — | E1-2 | 1586-1587 | — |
| A3-4 | 1540-1541 | 2 | E3-4 | 1588-1589 | 2 |
| A5-6 | 1542-1543 | 2 | E5-6 | 1590-1591 | 2 |
| A7-8 | 1544-1545 | 2 | E7-8 | 1592-1593 | 2 |
| A9-10 | 1546-1547 | 2 | E9-10 | 1594-1595 | 2 |
| A11-12 | 1548-1549 | — | E11-12 | 1596-1597 | 2 |
| B1-2 | 1550-1551 | — | F1-2 | 1598-1599 | — |
| B3-4 | 1552-1553 | 2 | F3-4 | 1600-1601 | 2 |
| B5-6 | 1554-1555 | 2 | F5-6 | 1602-1603 | 2 |
| B7-8 | 1556-1557 | 2 | F7-8 | 1604-1605 | 2 |
| B9-10 | 1558-1559 | 2 | F9-10 | 1606-1607 | 2 |
| B11-12 | 1560-1561 | — | F11-12 | 1608-1609 | 2 |
| C1-2 | 1562-1563 | — | G1-2 | 1610-1611 | — |
| C3-4 | 1564-1565 | 2 | G3-4 | 1612-1613 | 2 |
| C5-6 | 1566-1567 | 2 | G5-6 | 1614-1615 | 2 |
| C7-8 | 1568-1569 | 2 | G7-8 | 1616-1617 | 2 |
| C9-10 | 1570-1571 | 2 | G9-10 | 1618-1619 | 2 |
| C11-12 | 1572-1573 | — | G11-12 | 1620-1621 | 2 |
| D1-2 | 1574-1575 | — | H1-2 | 1622-1623 | — |
| D3-4 | 1576-1577 | 2 | H3-4 | 1624-1625 | 2 |
| D5-6 | 1578-1579 | 2 | H5-6 | 1626-1627 | 2 |
| D7-8 | 1580-1581 | 2 | H7-8 | 1628-1629 | 2 |
| D9-10 | 1582-1583 | 2 | H9-10 | 1630-1631 | 2 |
| D11-12 | 1584-1585 | — | H11-12 | 1632-1633 | — |

**A**

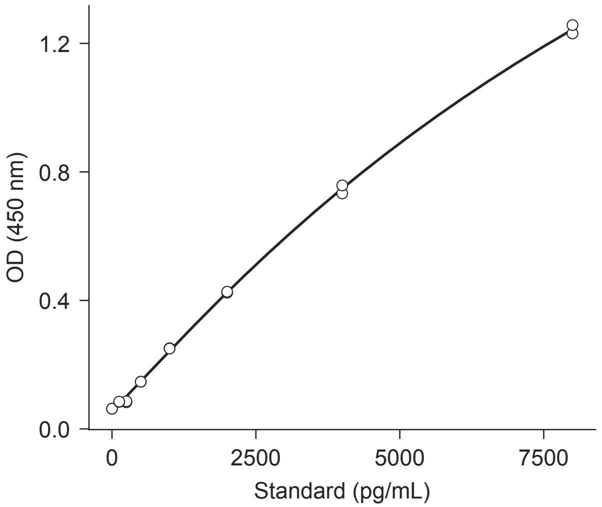


**B**

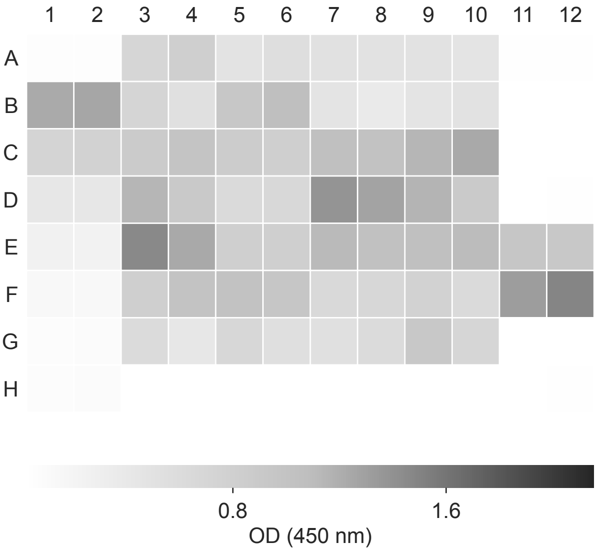


**C**

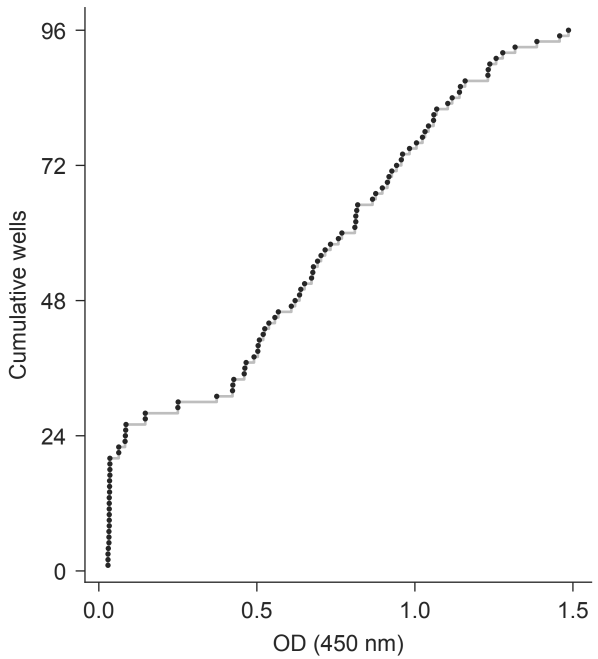


**D** elisa/plate_map.csv

| Well | Map rows | Dil. | Well | Map rows | Dil. |
|---|---|---|---|---|---|
| A1-2 | 1634-1635 | — | E1-2 | 1682-1683 | — |
| A3-4 | 1636-1637 | 2 | E3-4 | 1684-1685 | 2 |
| A5-6 | 1638-1639 | 2 | E5-6 | 1686-1687 | 2 |
| A7-8 | 1640-1641 | 2 | E7-8 | 1688-1689 | 2 |
| A9-10 | 1642-1643 | 2 | E9-10 | 1690-1691 | 2 |
| A11-12 | 1644-1645 | — | E11-12 | 1692-1693 | 2 |
| B1-2 | 1646-1647 | — | F1-2 | 1694-1695 | — |
| B3-4 | 1648-1649 | 2 | F3-4 | 1696-1697 | 2 |
| B5-6 | 1650-1651 | 2 | F5-6 | 1698-1699 | 2 |
| B7-8 | 1652-1653 | 2 | F7-8 | 1700-1701 | 2 |
| B9-10 | 1654-1655 | 2 | F9-10 | 1702-1703 | 2 |
| B11-12 | 1656-1657 | — | F11-12 | 1704-1705 | 2 |
| C1-2 | 1658-1659 | — | G1-2 | 1706-1707 | — |
| C3-4 | 1660-1661 | 2 | G3-4 | 1708-1709 | 2 |
| C5-6 | 1662-1663 | 2 | G5-6 | 1710-1711 | 2 |
| C7-8 | 1664-1665 | 2 | G7-8 | 1712-1713 | 2 |
| C9-10 | 1666-1667 | 2 | G9-10 | 1714-1715 | 2 |
| C11-12 | 1668-1669 | — | G11-12 | 1716-1717 | 2 |
| D1-2 | 1670-1671 | — | H1-2 | 1718-1719 | — |
| D3-4 | 1672-1673 | 2 | H3-4 | 1720-1721 | 2 |
| D5-6 | 1674-1675 | 2 | H5-6 | 1722-1723 | 2 |
| D7-8 | 1676-1677 | 2 | H7-8 | 1724-1725 | 2 |
| D9-10 | 1678-1679 | 2 | H9-10 | 1726-1727 | 2 |
| D11-12 | 1680-1681 | — | H11-12 | 1728-1729 | — |

**A**

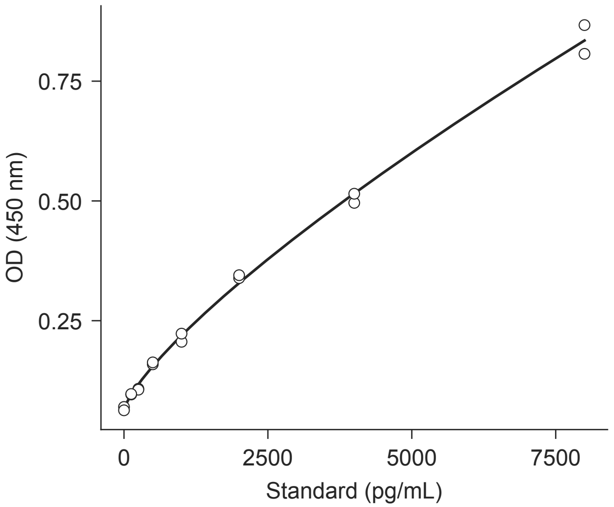


**B**

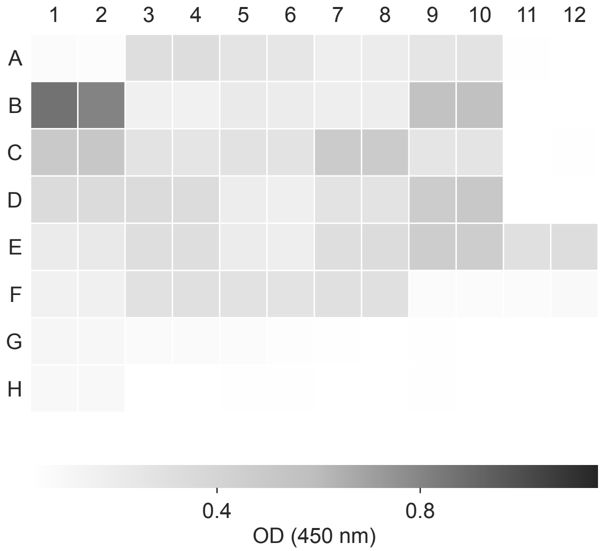


**C**

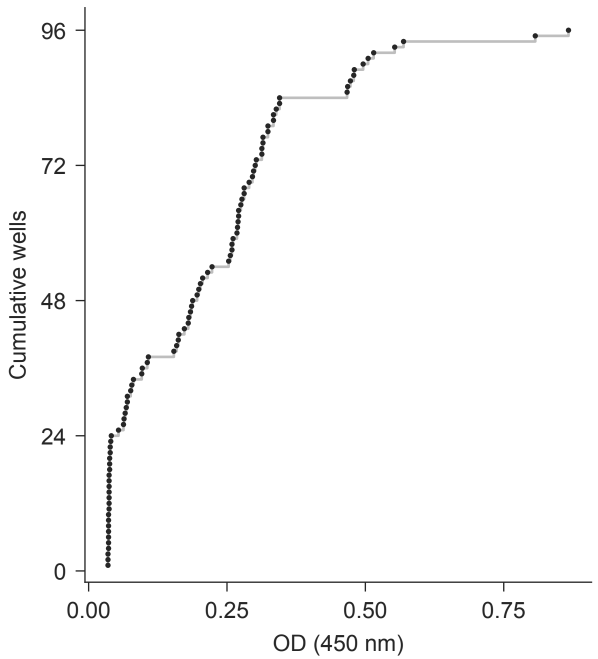


**D** elisa/plate_map.csv

| Well | Map rows | Dil. | Well | Map rows | Dil. |
|---|---|---|---|---|---|
| A1-2 | 1730-1731 | — | D7-8 | 1772-1773 | 1.0 |
| A3-4 | 1732-1733 | 1.0 | D9-10 | 1774-1775 | 1.0 |
| A5-6 | 1734-1735 | 1.0 | D11-12 | 1776-1777 | — |
| A7-8 | 1736-1737 | 1.0 | E1-2 | 1778-1779 | — |
| A9-10 | 1738-1739 | 1.0 | E3-4 | 1780-1781 | 1.0 |
| A11-12 | 1740-1741 | — | E5-6 | 1782-1783 | 1.0 |
| B1-2 | 1742-1743 | — | E7-8 | 1784-1785 | 1.0 |
| B3-4 | 1744-1745 | 1.0 | E9-10 | 1786-1787 | 1.0 |
| B5-6 | 1746-1747 | 1.0 | E11-12 | 1788-1789 | 1.0 |
| B7-8 | 1748-1749 | 1.0 | F1-2 | 1790-1791 | — |
| B9-10 | 1750-1751 | 1.0 | F3-4 | 1792-1793 | 1.0 |
| B11-12 | 1752-1753 | — | F5-6 | 1794-1795 | 1.0 |
| C1-2 | 1754-1755 | — | F7-8 | 1796-1797 | 1.0 |
| C3-4 | 1756-1757 | 1.0 | F9-10 | 1798-1799 | 1.0 |
| C5-6 | 1758-1759 | 1.0 | F11-12 | 1800-1801 | 1.0 |
| C7-8 | 1760-1761 | 1.0 | G1-2 | 1802-1803 | — |
| C9-10 | 1762-1763 | 1.0 | G3-4 | 1804-1805 | 1.0 |
| C11-12 | 1764-1765 | — | G5-6 | 1806-1807 | 1.0 |
| D1-2 | 1766-1767 | — | G7-12 | 1808-1813 | — |
| D3-4 | 1768-1769 | 1.0 | H1-2 | 1814-1815 | — |
| D5-6 | 1770-1771 | 1.0 | H3-12 | 1816-1825 | — |

**B**

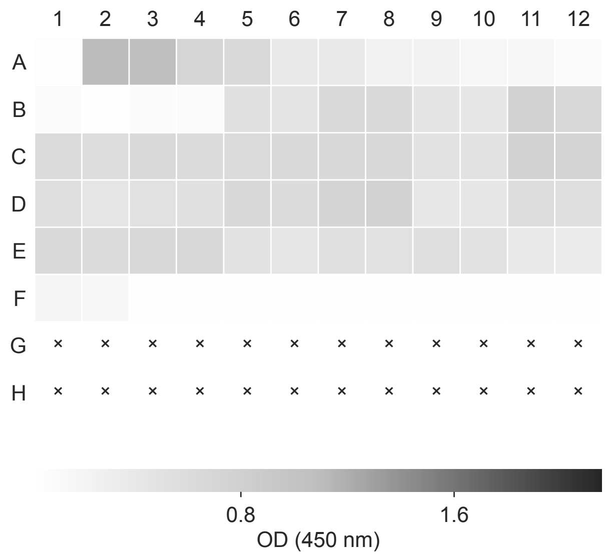


**C**

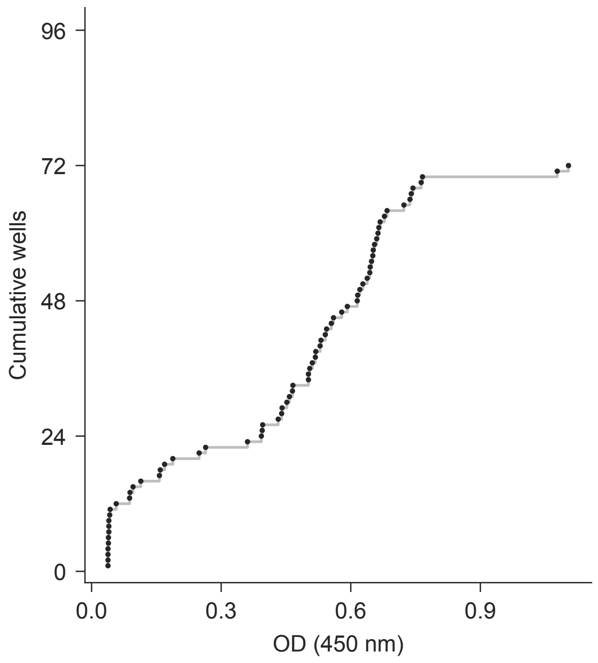


**D** OD (450 nm)

| Well | Value | Well | Value | Well | Value | Well | Value |
|---|---|---|---|---|---|---|---|
| A1 | 0.057 | C1 | 0.628 | E1 | 0.648 | G1 | — |
| A2 | 1.104 | C2 | 0.579 | E2 | 0.616 | G2 | — |
| A3 | 1.078 | C3 | 0.644 | E3 | 0.663 | G3 | — |
| A4 | 0.723 | C4 | 0.621 | E4 | 0.684 | G4 | — |
| A5 | 0.655 | C5 | 0.638 | E5 | 0.505 | G5 | — |
| A6 | 0.393 | C6 | 0.651 | E6 | 0.458 | G6 | — |
| A7 | 0.395 | C7 | 0.668 | E7 | 0.529 | G7 | — |
| A8 | 0.264 | C8 | 0.660 | E8 | 0.518 | G8 | — |
| A9 | 0.249 | C9 | 0.502 | E9 | 0.560 | G9 | — |
| A10 | 0.169 | C10 | 0.502 | E10 | 0.511 | G10 | — |
| A11 | 0.157 | C11 | 0.763 | E11 | 0.396 | G11 | — |
| A12 | 0.096 | C12 | 0.737 | E12 | 0.361 | G12 | — |
| B1 | 0.114 | D1 | 0.555 | F1 | 0.188 | H1 | — |
| B2 | 0.043 | D2 | 0.452 | F2 | 0.159 | H2 | — |
| B3 | 0.089 | D3 | 0.519 | F3 | 0.039 | H3 | — |
| B4 | 0.088 | D4 | 0.531 | F4 | 0.038 | H4 | — |
| B5 | 0.541 | D5 | 0.665 | F5 | 0.038 | H5 | — |
| B6 | 0.465 | D6 | 0.615 | F6 | 0.039 | H6 | — |
| B7 | 0.652 | D7 | 0.740 | F7 | 0.038 | H7 | — |
| B8 | 0.645 | D8 | 0.766 | F8 | 0.038 | H8 | — |
| B9 | 0.466 | D9 | 0.432 | F9 | 0.040 | H9 | — |
| B10 | 0.440 | D10 | 0.441 | F10 | 0.042 | H10 | — |
| B11 | 0.744 | D11 | 0.592 | F11 | 0.040 | H11 | — |
| B12 | 0.678 | D12 | 0.544 | F12 | 0.040 | H12 | — |
