## Supplementary Data 3 for "Material context and TGF-β blockade timing determine matrix retained after withdrawal in patient-derived GelMA-bioprinted keratoconus stroma"

2.tif *

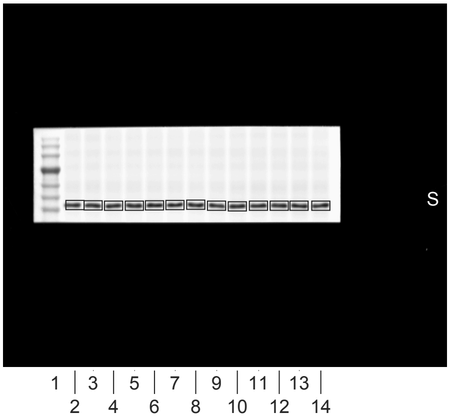


1.tif

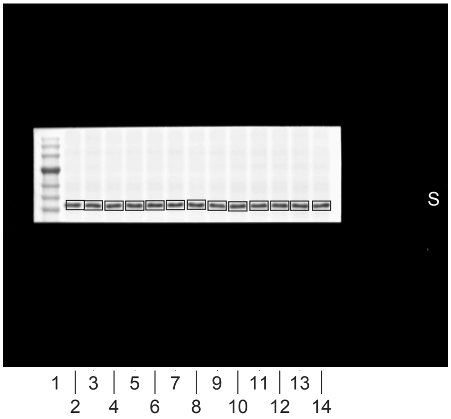


3.tif

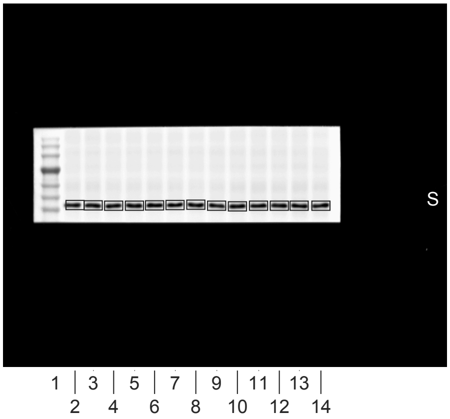


4.tif

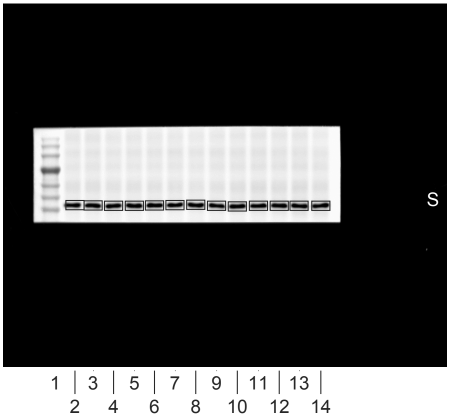


5.tif

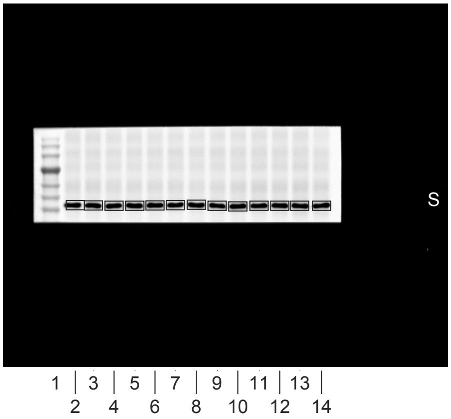


0.tif

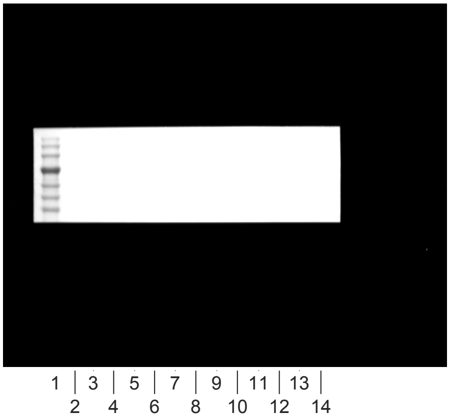


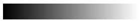
linear RGB

0 255

| Lane | Donor | Format | Treatment | Lane type |
|---|---|---|---|---|
| 1 | - | - | Marker | marker |
| 2 | D01 | 2D | Vehicle | sample |
| 3 | D01 | 2D | TGF-beta1 | sample |
| 4 | D01 | cast | Vehicle | sample |
| 5 | D01 | cast | TGF-beta1 | sample |
| 6 | D01 | printed | Vehicle | sample |
| 7 | D01 | printed | TGF-beta1 | sample |
| 8 | D01-D06 pool | 2D | TGF-beta1, T72 h | calibration pool |
| 9 | D04 | 2D | Vehicle | sample |
| 10 | D04 | 2D | TGF-beta1 | sample |
| 11 | D04 | cast | Vehicle | sample |
| 12 | D04 | cast | TGF-beta1 | sample |
| 13 | D04 | printed | Vehicle | sample |
| 14 | D04 | printed | TGF-beta1 | sample |
| 15 | - | - | loading buffer | empty |

1.tif *

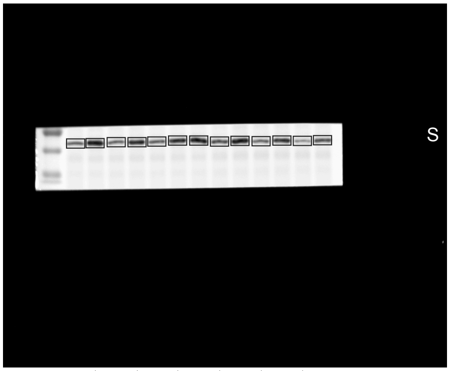


2.tif

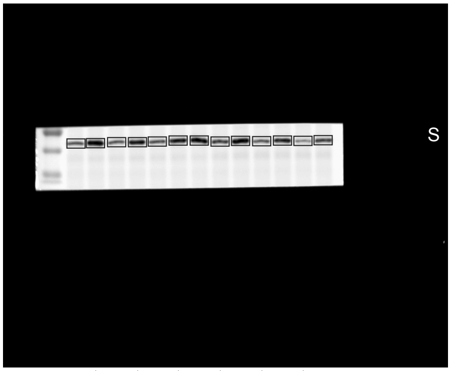


3.tif

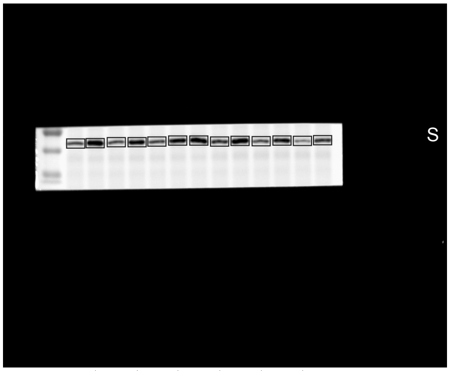


4.tif

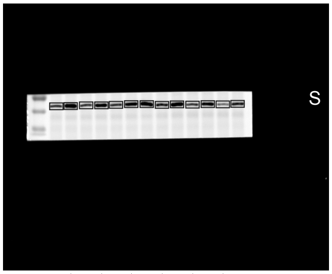


5.tif

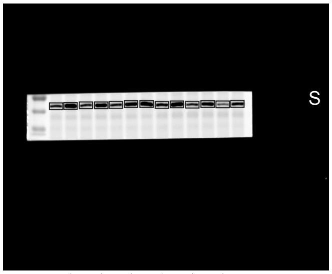


6.tif

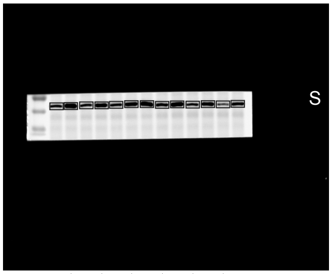


0.tif

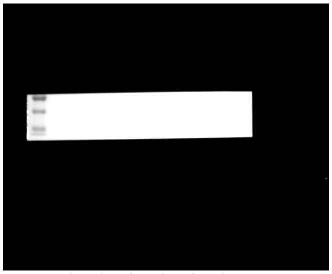


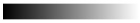


| Lane | Donor | Format | Treatment | Lane type |
|---|---|---|---|---|
| 1 | - | - | Marker | marker |
| 2 | D01 | 2D | Vehicle | sample |
| 3 | D01 | 2D | TGF-beta1 | sample |
| 4 | D01 | cast | Vehicle | sample |
| 5 | D01 | cast | TGF-beta1 | sample |
| 6 | D01 | printed | Vehicle | sample |
| 7 | D01 | printed | TGF-beta1 | sample |
| 8 | D01-D06 pool | 2D | TGF-beta1, T72 h | calibration pool |
| 9 | D04 | 2D | Vehicle | sample |
| 10 | D04 | 2D | TGF-beta1 | sample |
| 11 | D04 | cast | Vehicle | sample |
| 12 | D04 | cast | TGF-beta1 | sample |
| 13 | D04 | printed | Vehicle | sample |
| 14 | D04 | printed | TGF-beta1 | sample |
| 15 | - | - | loading buffer | empty |

1.tif *

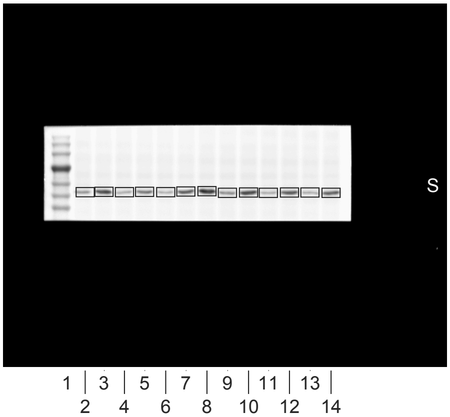


2.tif

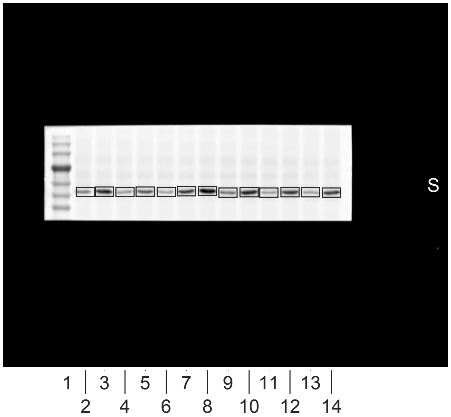


3.tif

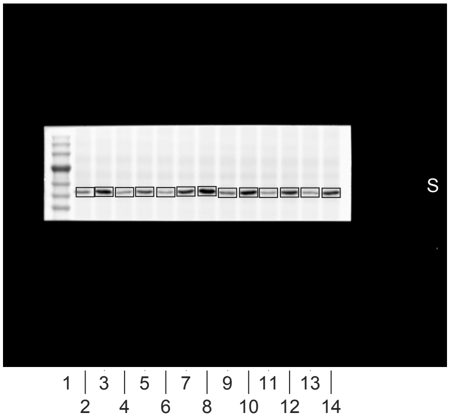


4.tif

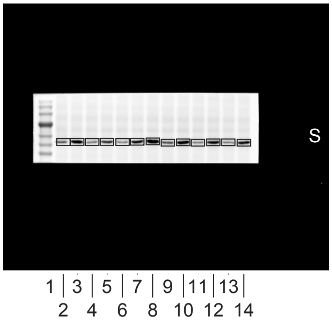


5.tif

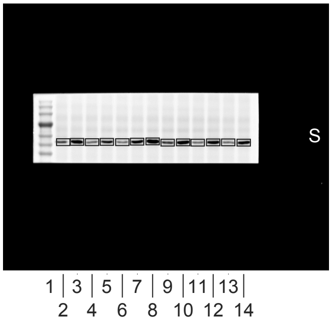


6.tif

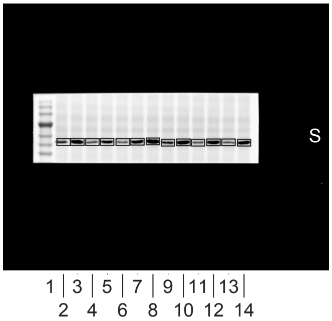


0.tif

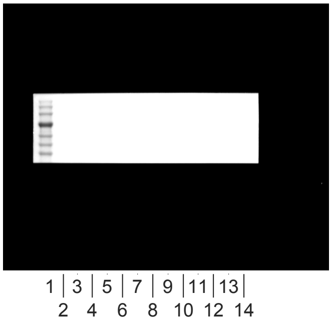


linear RGB
0 255

| Lane | Donor | Format | Treatment | Lane type |
|---|---|---|---|---|
| 1 | - | - | Marker | marker |
| 2 | D01 | 2D | Vehicle | sample |
| 3 | D01 | 2D | TGF-beta1 | sample |
| 4 | D01 | cast | Vehicle | sample |
| 5 | D01 | cast | TGF-beta1 | sample |
| 6 | D01 | printed | Vehicle | sample |
| 7 | D01 | printed | TGF-beta1 | sample |
| 8 | D01-D06 pool | 2D | TGF-beta1, T72 h | calibration pool |
| 9 | D04 | 2D | Vehicle | sample |
| 10 | D04 | 2D | TGF-beta1 | sample |
| 11 | D04 | cast | Vehicle | sample |
| 12 | D04 | cast | TGF-beta1 | sample |
| 13 | D04 | printed | Vehicle | sample |
| 14 | D04 | printed | TGF-beta1 | sample |
| 15 | - | - | loading buffer | empty |

1.tif *

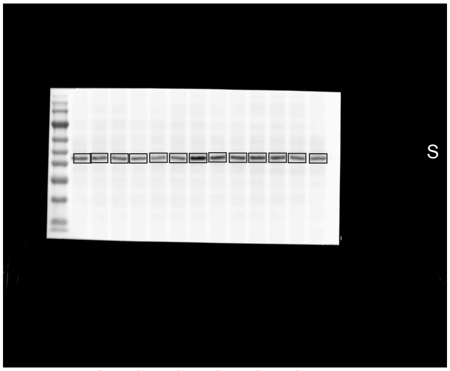


1 | 3 | 5 | 7 | 9 | 11 | 13 |
2 4 6 8 10 12 14

2.tif

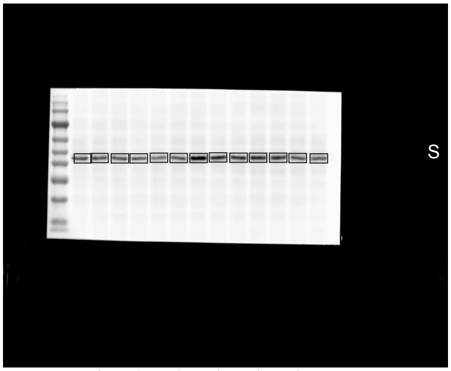


1 | 3 | 5 | 7 | 9 | 11 | 13 |
2 4 6 8 10 12 14

3.tif

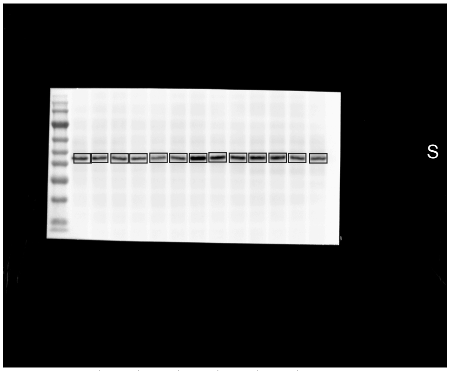


1 | 3 | 5 | 7 | 9 | 11 | 13 |
2 4 6 8 10 12 14

4.tif

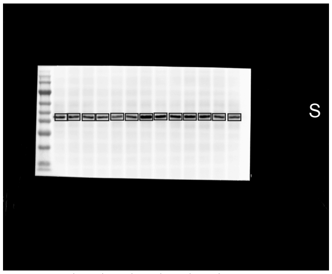


1 | 3 | 5 | 7 | 9 | 11 | 13 |
2 4 6 8 10 12 14

5.tif

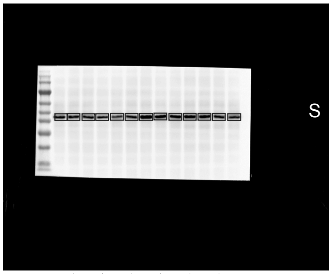


1 | 3 | 5 | 7 | 9 | 11 | 13 |
2 4 6 8 10 12 14

6.tif

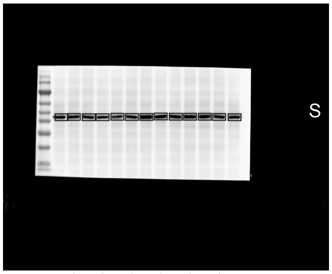


1 | 3 | 5 | 7 | 9 | 11 | 13 |
2 4 6 8 10 12 14

0.tif

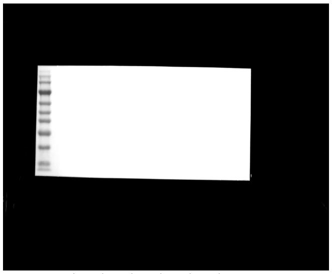

1 | 3 | 5 | 7 | 9 | 11 | 13 |
2 4 6 8 10 12 14

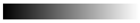

linear RGB

0 255

| Lane | Donor | Format | Treatment | Lane type |
|---|---|---|---|---|
| 1 | - | - | Marker | marker |
| 2 | D01 | 2D | Vehicle | sample |
| 3 | D01 | 2D | TGF-beta1 | sample |
| 4 | D01 | cast | Vehicle | sample |
| 5 | D01 | cast | TGF-beta1 | sample |
| 6 | D01 | printed | Vehicle | sample |
| 7 | D01 | printed | TGF-beta1 | sample |
| 8 | D01-D06 pool | 2D | TGF-beta1, T72 h | calibration pool |
| 9 | D04 | 2D | Vehicle | sample |
| 10 | D04 | 2D | TGF-beta1 | sample |
| 11 | D04 | cast | Vehicle | sample |
| 12 | D04 | cast | TGF-beta1 | sample |
| 13 | D04 | printed | Vehicle | sample |
| 14 | D04 | printed | TGF-beta1 | sample |
| 15 | - | - | loading buffer | empty |

2.tif *

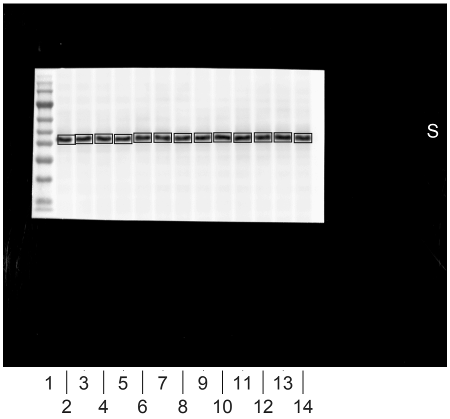


1.tif

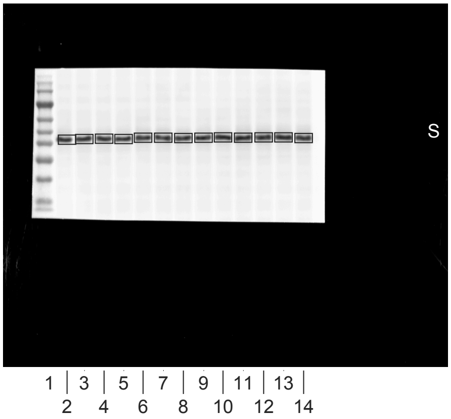


3.tif

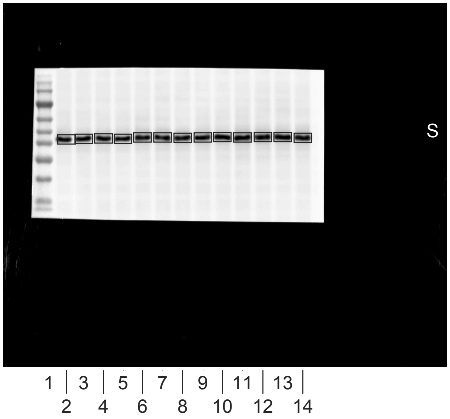


4.tif

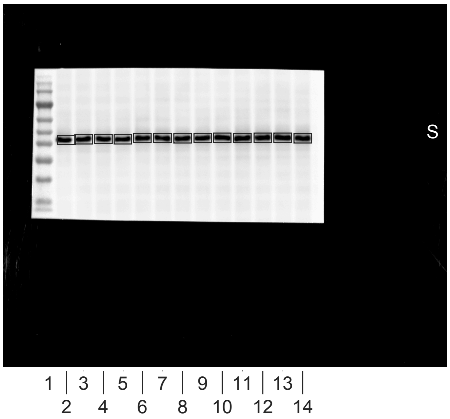


5.tif

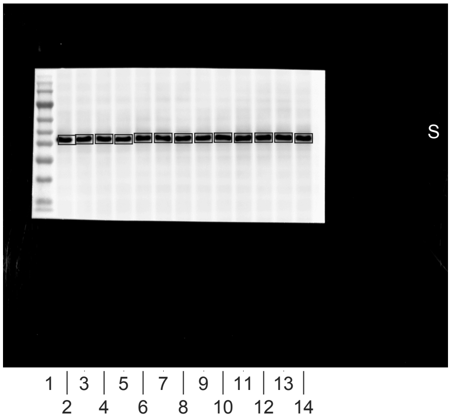


0.tif

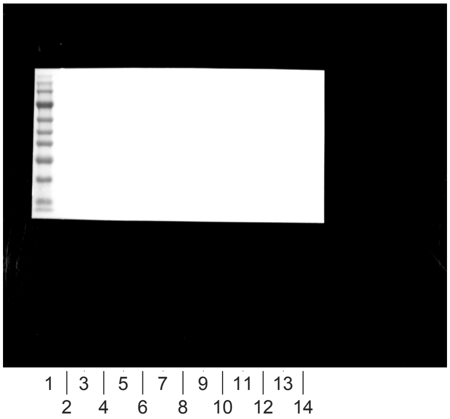


linear RGB
0 255

| Lane | Donor | Format | Treatment | Lane type |
|---|---|---|---|---|
| 1 | - | - | Marker | marker |
| 2 | D01 | 2D | Vehicle | sample |
| 3 | D01 | 2D | TGF-beta1 | sample |
| 4 | D01 | cast | Vehicle | sample |
| 5 | D01 | cast | TGF-beta1 | sample |
| 6 | D01 | printed | Vehicle | sample |
| 7 | D01 | printed | TGF-beta1 | sample |
| 8 | D01-D06 pool | 2D | TGF-beta1, T72 h | calibration pool |
| 9 | D04 | 2D | Vehicle | sample |
| 10 | D04 | 2D | TGF-beta1 | sample |
| 11 | D04 | cast | Vehicle | sample |
| 12 | D04 | cast | TGF-beta1 | sample |
| 13 | D04 | printed | Vehicle | sample |
| 14 | D04 | printed | TGF-beta1 | sample |
| 15 | - | - | loading buffer | empty |

3.tif *

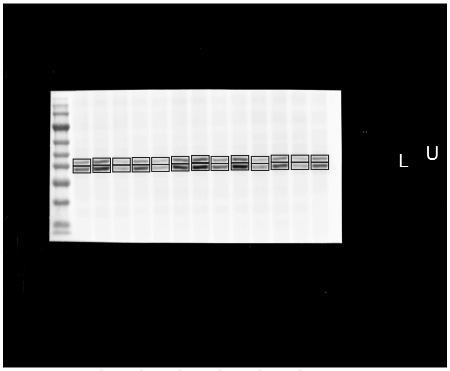


1 | 3 | 5 | 7 | 9 | 11 | 13 |
2 4 6 8 10 12 14

1.tif

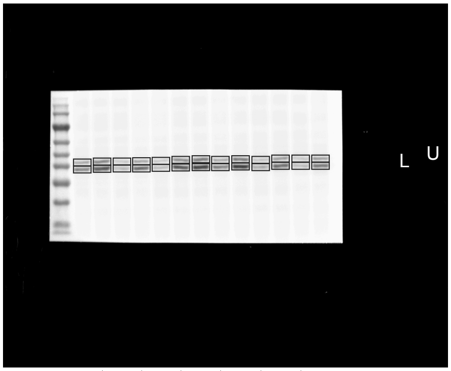


1 | 3 | 5 | 7 | 9 | 11 | 13 |
2 4 6 8 10 12 14

2.tif

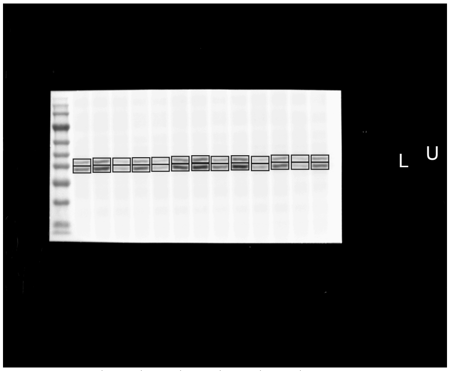


1 | 3 | 5 | 7 | 9 | 11 | 13 |
2 4 6 8 10 12 14

4.tif

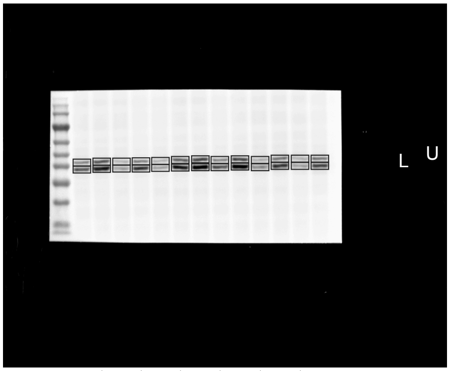


1 | 3 | 5 | 7 | 9 | 11 | 13 |
2 4 6 8 10 12 14

5.tif

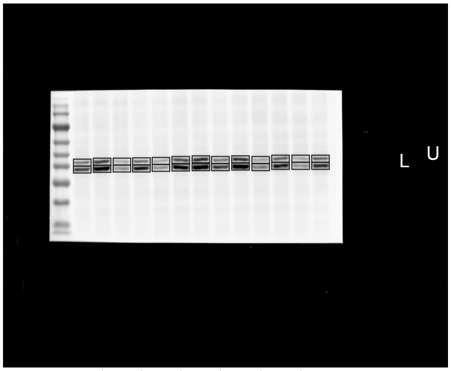


1 | 3 | 5 | 7 | 9 | 11 | 13 |
2 4 6 8 10 12 14

0.tif

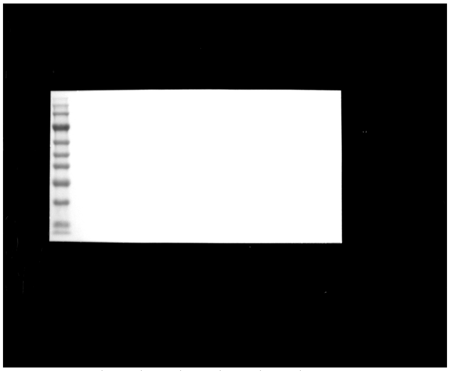

1 | 3 | 5 | 7 | 9 | 11 | 13 |
2 4 6 8 10 12 14

linear RGB
0 255

| Lane | Donor | Format | Treatment | Lane type |
|---|---|---|---|---|
| 1 | - | - | Marker | marker |
| 2 | D01 | 2D | Vehicle | sample |
| 3 | D01 | 2D | TGF-beta1 | sample |
| 4 | D01 | cast | Vehicle | sample |
| 5 | D01 | cast | TGF-beta1 | sample |
| 6 | D01 | printed | Vehicle | sample |
| 7 | D01 | printed | TGF-beta1 | sample |
| 8 | D01-D06 pool | 2D | TGF-beta1, T72 h | calibration pool |
| 9 | D04 | 2D | Vehicle | sample |
| 10 | D04 | 2D | TGF-beta1 | sample |
| 11 | D04 | cast | Vehicle | sample |
| 12 | D04 | cast | TGF-beta1 | sample |
| 13 | D04 | printed | Vehicle | sample |
| 14 | D04 | printed | TGF-beta1 | sample |
| 15 | - | - | loading buffer | empty |

1.tif *

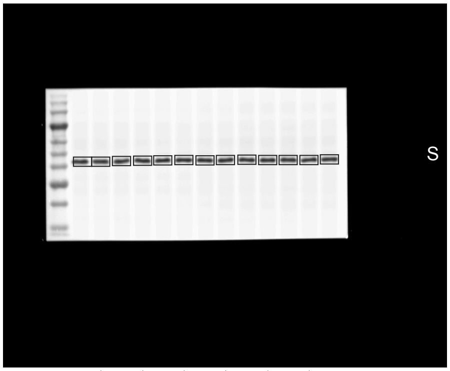


1 | 3 | 5 | 7 | 9 | 11 | 13 |
2 4 6 8 10 12 14

2.tif

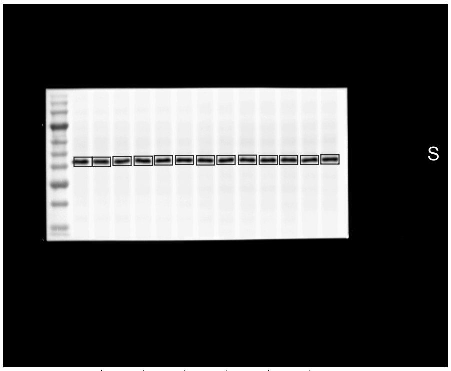


1 | 3 | 5 | 7 | 9 | 11 | 13 |
2 4 6 8 10 12 14

3.tif

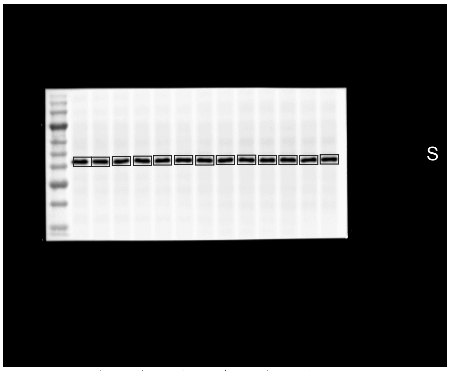


1 | 3 | 5 | 7 | 9 | 11 | 13 |
2 4 6 8 10 12 14

4.tif

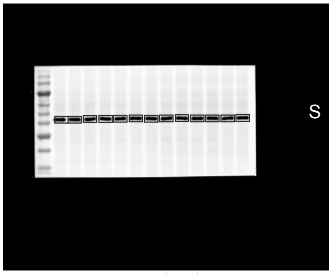


1 | 3 | 5 | 7 | 9 | 11 | 13 |
2 4 6 8 10 12 14

5.tif

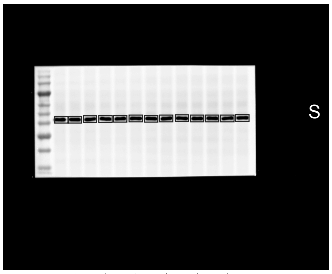


1 | 3 | 5 | 7 | 9 | 11 | 13 |
2 4 6 8 10 12 14

6.tif

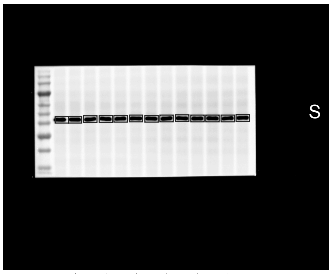


1 | 3 | 5 | 7 | 9 | 11 | 13 |
2 4 6 8 10 12 14

0.tif

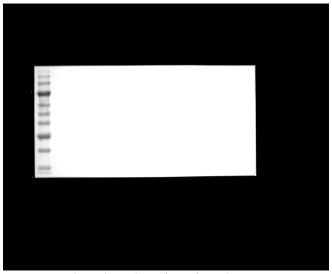

1 | 3 | 5 | 7 | 9 | 11 | 13 |
2 4 6 8 10 12 14

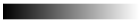

linear RGB
0 255

| Lane | Donor | Format | Treatment | Lane type |
|---|---|---|---|---|
| 1 | - | - | Marker | marker |
| 2 | D02 | 2D | Vehicle | sample |
| 3 | D02 | 2D | TGF-beta1 | sample |
| 4 | D02 | cast | Vehicle | sample |
| 5 | D02 | cast | TGF-beta1 | sample |
| 6 | D02 | printed | Vehicle | sample |
| 7 | D02 | printed | TGF-beta1 | sample |
| 8 | D01-D06 pool | 2D | TGF-beta1, T72 h | calibration pool |
| 9 | D05 | 2D | Vehicle | sample |
| 10 | D05 | 2D | TGF-beta1 | sample |
| 11 | D05 | cast | Vehicle | sample |
| 12 | D05 | cast | TGF-beta1 | sample |
| 13 | D05 | printed | Vehicle | sample |
| 14 | D05 | printed | TGF-beta1 | sample |
| 15 | - | - | loading buffer | empty |

1.tif

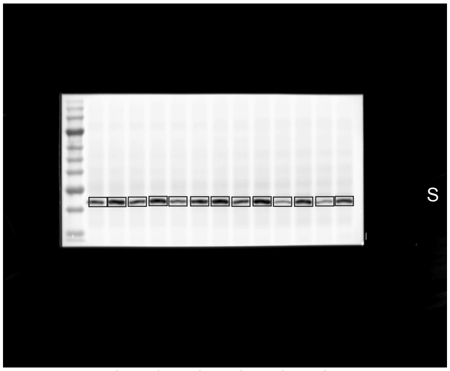


1 3 5 7 9 11 13
2 4 6 8 10 12 14

2.tif

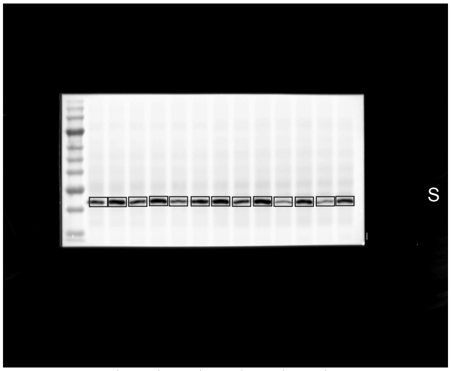


1 3 5 7 9 11 13
2 4 6 8 10 12 14

3.tif

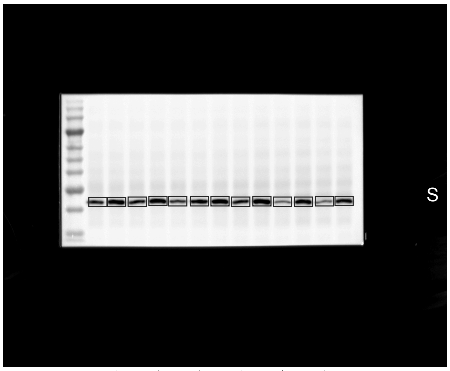


1 3 5 7 9 11 13
2 4 6 8 10 12 14

4.tif

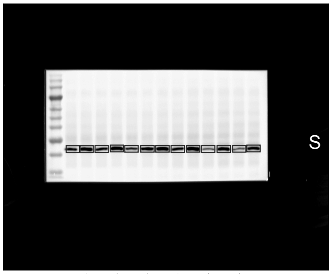


1 3 5 7 9 11 13
2 4 6 8 10 12 14

5.tif

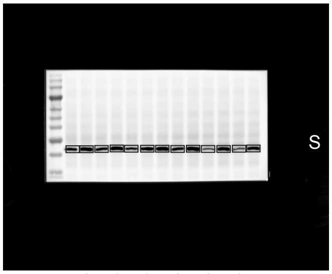


1 3 5 7 9 11 13
2 4 6 8 10 12 14

6.tif

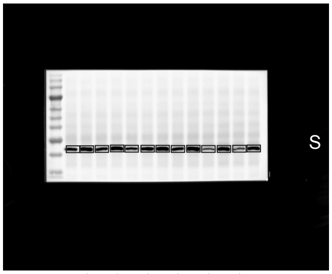


1 3 5 7 9 11 13
2 4 6 8 10 12 14

0.tif

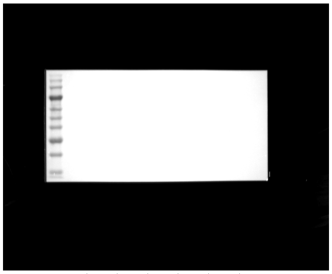

1 3 5 7 9 11 13
2 4 6 8 10 12 14

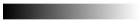

linear RGB

0 255

| Lane | Donor | Format | Treatment | Lane type |
|---|---|---|---|---|
| 1 | - | - | Marker | marker |
| 2 | D02 | 2D | Vehicle | sample |
| 3 | D02 | 2D | TGF-beta1 | sample |
| 4 | D02 | cast | Vehicle | sample |
| 5 | D02 | cast | TGF-beta1 | sample |
| 6 | D02 | printed | Vehicle | sample |
| 7 | D02 | printed | TGF-beta1 | sample |
| 8 | D01-D06 pool | 2D | TGF-beta1, T72 h | calibration pool |
| 9 | D05 | 2D | Vehicle | sample |
| 10 | D05 | 2D | TGF-beta1 | sample |
| 11 | D05 | cast | Vehicle | sample |
| 12 | D05 | cast | TGF-beta1 | sample |
| 13 | D05 | printed | Vehicle | sample |
| 14 | D05 | printed | TGF-beta1 | sample |
| 15 | - | - | loading buffer | empty |

3.tif *

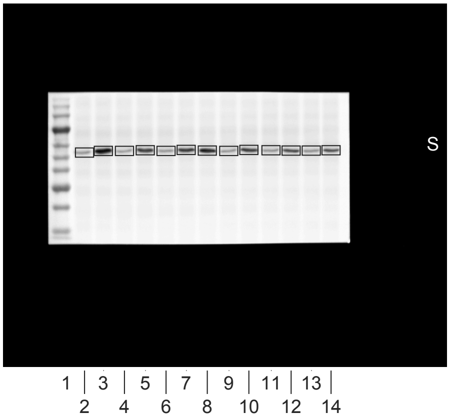


1.tif

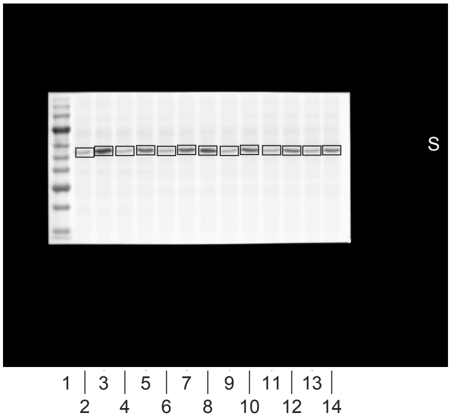


2.tif

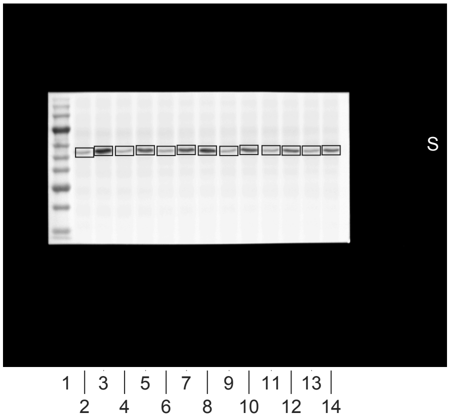


4.tif

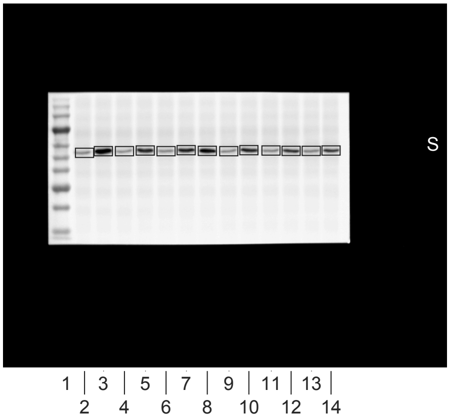


5.tif

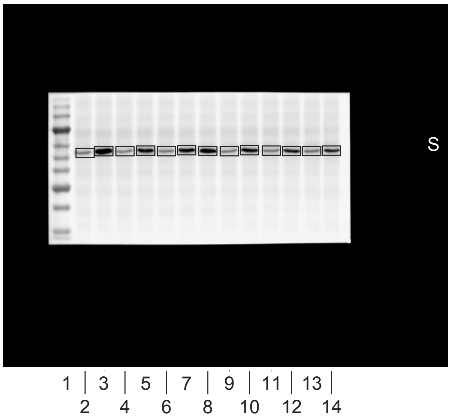


0.tif

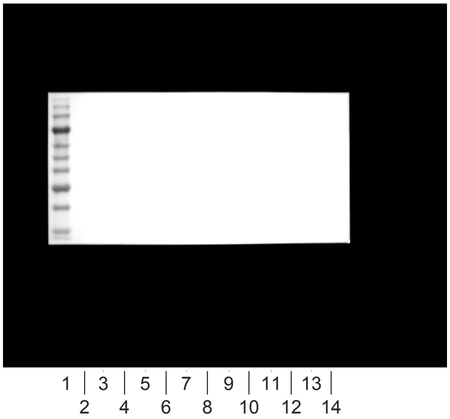


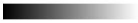
linear RGB
0 255

| Lane | Donor | Format | Treatment | Lane type |
|---|---|---|---|---|
| 1 | - | - | Marker | marker |
| 2 | D02 | 2D | Vehicle | sample |
| 3 | D02 | 2D | TGF-beta1 | sample |
| 4 | D02 | cast | Vehicle | sample |
| 5 | D02 | cast | TGF-beta1 | sample |
| 6 | D02 | printed | Vehicle | sample |
| 7 | D02 | printed | TGF-beta1 | sample |
| 8 | D01-D06 pool | 2D | TGF-beta1, T72 h | calibration pool |
| 9 | D05 | 2D | Vehicle | sample |
| 10 | D05 | 2D | TGF-beta1 | sample |
| 11 | D05 | cast | Vehicle | sample |
| 12 | D05 | cast | TGF-beta1 | sample |
| 13 | D05 | printed | Vehicle | sample |
| 14 | D05 | printed | TGF-beta1 | sample |
| 15 | - | - | loading buffer | empty |

1.tif *

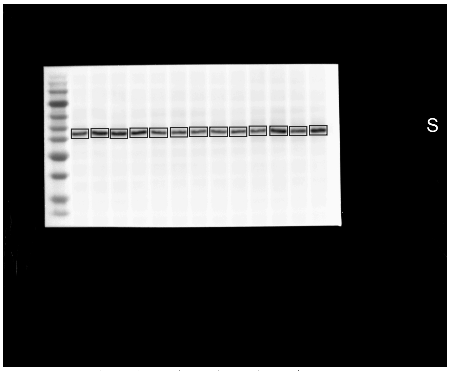


1 | 3 | 5 | 7 | 9 | 11 | 13 |
2 4 6 8 10 12 14

2.tif

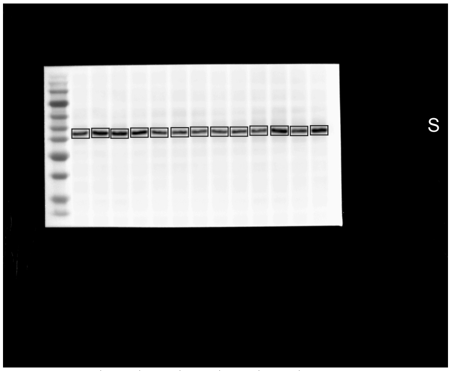


1 | 3 | 5 | 7 | 9 | 11 | 13 |
2 4 6 8 10 12 14

3.tif

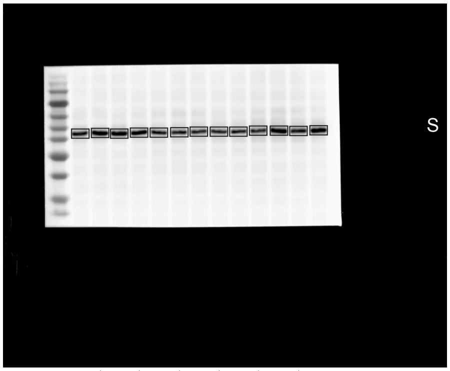


1 | 3 | 5 | 7 | 9 | 11 | 13 |
2 4 6 8 10 12 14

4.tif

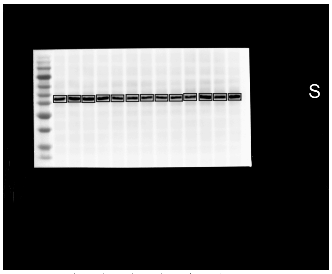


1 | 3 | 5 | 7 | 9 | 11 | 13 |
2 4 6 8 10 12 14

5.tif

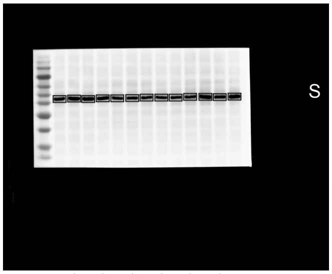


1 | 3 | 5 | 7 | 9 | 11 | 13 |
2 4 6 8 10 12 14

6.tif

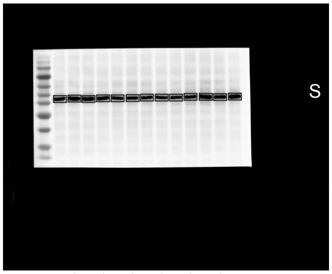


1 | 3 | 5 | 7 | 9 | 11 | 13 |
2 4 6 8 10 12 14

0.tif

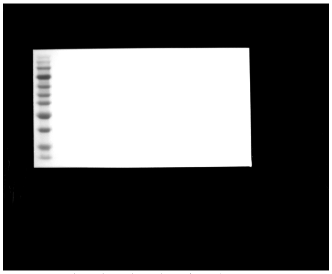

1 | 3 | 5 | 7 | 9 | 11 | 13 |
2 4 6 8 10 12 14

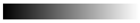
linear RGB
0 255

| Lane | Donor | Format | Treatment | Lane type |
|---|---|---|---|---|
| 1 | - | - | Marker | marker |
| 2 | D02 | 2D | Vehicle | sample |
| 3 | D02 | 2D | TGF-beta1 | sample |
| 4 | D02 | cast | Vehicle | sample |
| 5 | D02 | cast | TGF-beta1 | sample |
| 6 | D02 | printed | Vehicle | sample |
| 7 | D02 | printed | TGF-beta1 | sample |
| 8 | D01-D06 pool | 2D | TGF-beta1, T72 h | calibration pool |
| 9 | D05 | 2D | Vehicle | sample |
| 10 | D05 | 2D | TGF-beta1 | sample |
| 11 | D05 | cast | Vehicle | sample |
| 12 | D05 | cast | TGF-beta1 | sample |
| 13 | D05 | printed | Vehicle | sample |
| 14 | D05 | printed | TGF-beta1 | sample |
| 15 | - | - | loading buffer | empty |

3.tif *

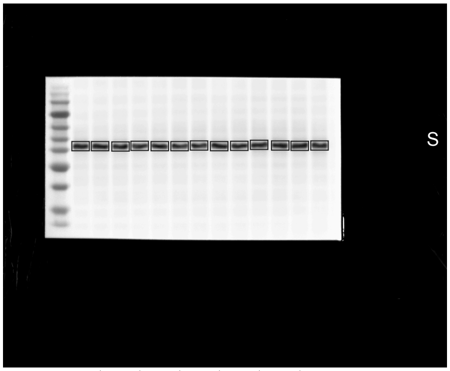


1.tif

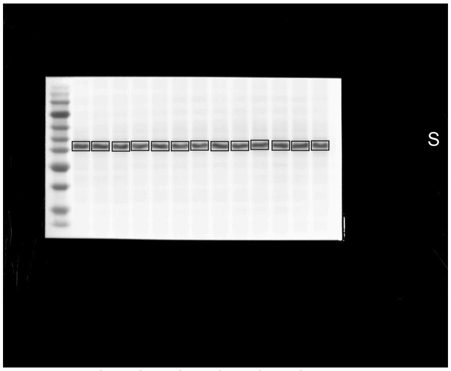


2.tif

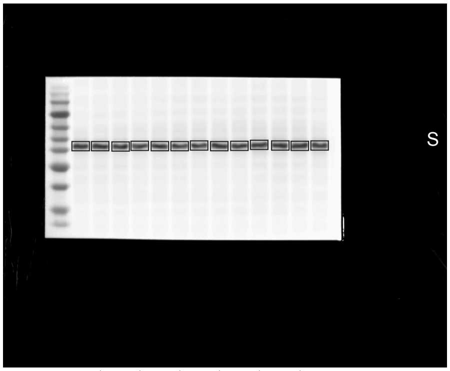


4.tif

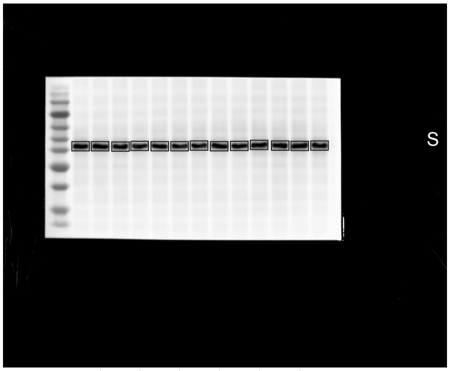


5.tif

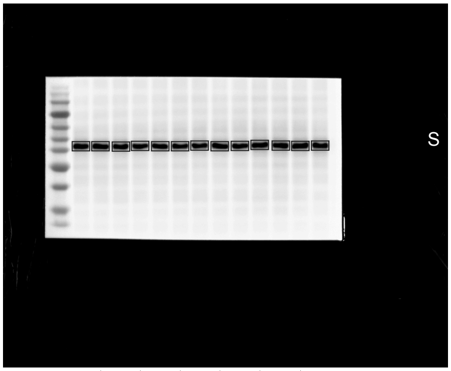


0.tif

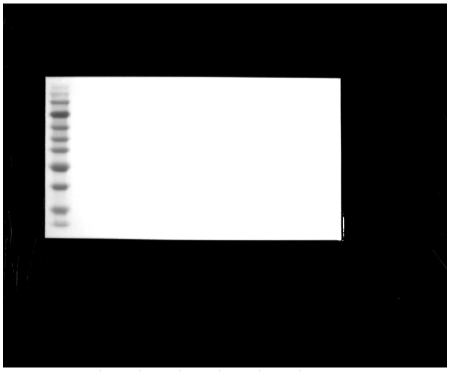


linear RGB
0 255

| Lane | Donor | Format | Treatment | Lane type |
|---|---|---|---|---|
| 1 | - | - | Marker | marker |
| 2 | D02 | 2D | Vehicle | sample |
| 3 | D02 | 2D | TGF-beta1 | sample |
| 4 | D02 | cast | Vehicle | sample |
| 5 | D02 | cast | TGF-beta1 | sample |
| 6 | D02 | printed | Vehicle | sample |
| 7 | D02 | printed | TGF-beta1 | sample |
| 8 | D01-D06 pool | 2D | TGF-beta1, T72 h | calibration pool |
| 9 | D05 | 2D | Vehicle | sample |
| 10 | D05 | 2D | TGF-beta1 | sample |
| 11 | D05 | cast | Vehicle | sample |
| 12 | D05 | cast | TGF-beta1 | sample |
| 13 | D05 | printed | Vehicle | sample |
| 14 | D05 | printed | TGF-beta1 | sample |
| 15 | - | - | loading buffer | empty |

1.tif

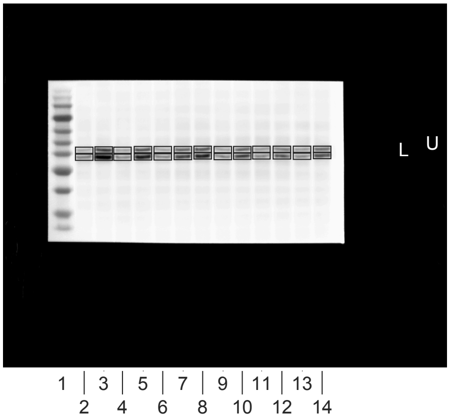


2.tif

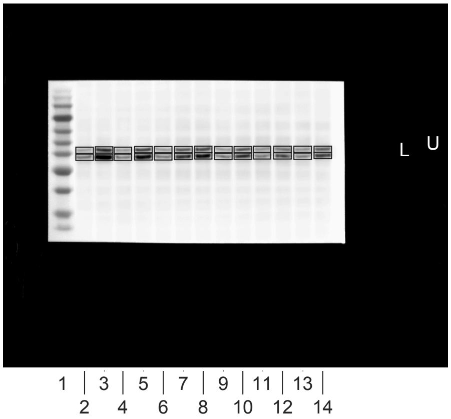


3.tif

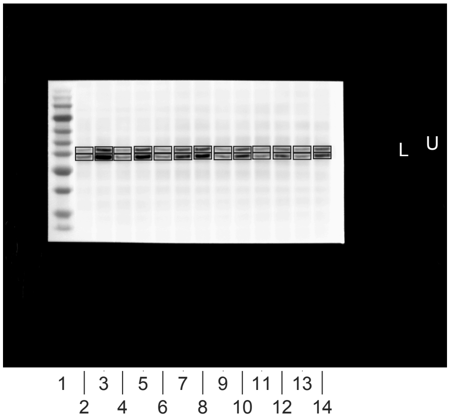


4.tif

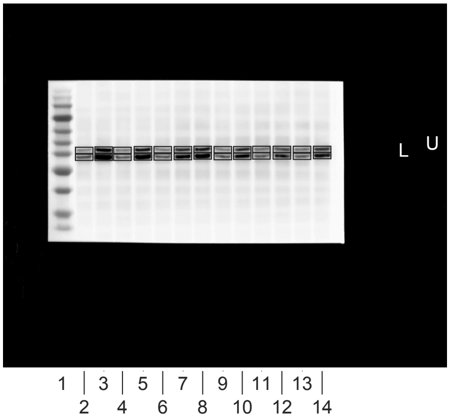


5.tif

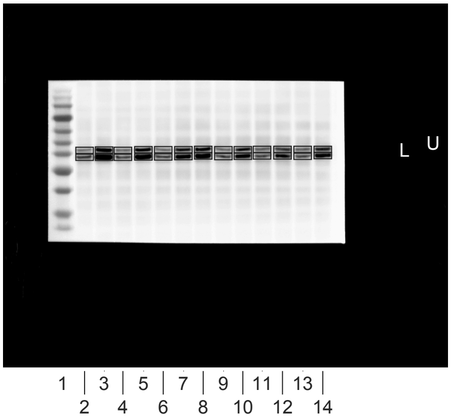


0.tif

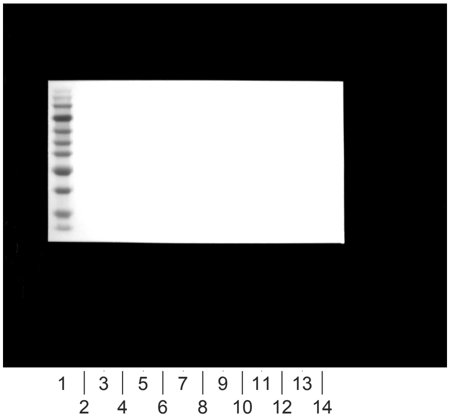


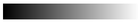
linear RGB

0 255

| Lane | Donor | Format | Treatment | Lane type |
|---|---|---|---|---|
| 1 | - | - | Marker | marker |
| 2 | D02 | 2D | Vehicle | sample |
| 3 | D02 | 2D | TGF-beta1 | sample |
| 4 | D02 | cast | Vehicle | sample |
| 5 | D02 | cast | TGF-beta1 | sample |
| 6 | D02 | printed | Vehicle | sample |
| 7 | D02 | printed | TGF-beta1 | sample |
| 8 | D01-D06 pool | 2D | TGF-beta1, T72 h | calibration pool |
| 9 | D05 | 2D | Vehicle | sample |
| 10 | D05 | 2D | TGF-beta1 | sample |
| 11 | D05 | cast | Vehicle | sample |
| 12 | D05 | cast | TGF-beta1 | sample |
| 13 | D05 | printed | Vehicle | sample |
| 14 | D05 | printed | TGF-beta1 | sample |
| 15 | - | - | loading buffer | empty |

1.tif *

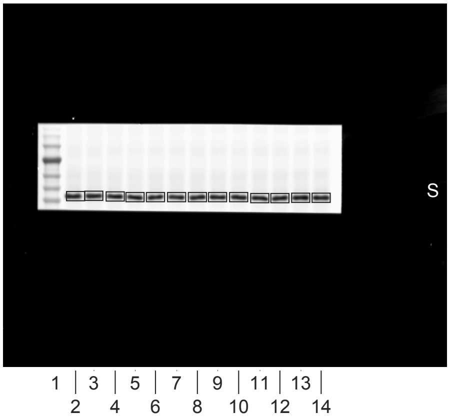


2.tif

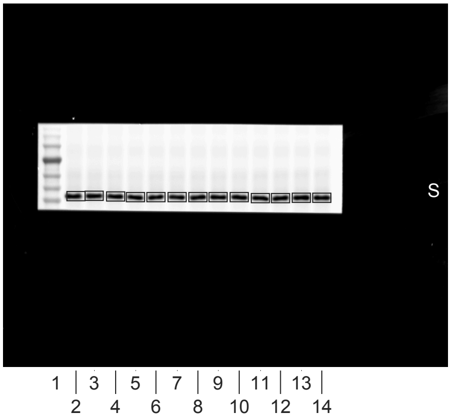


3.tif

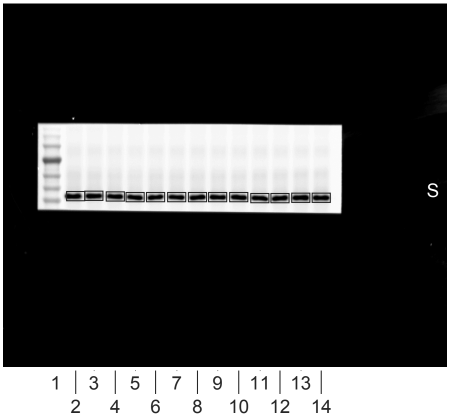


4.tif

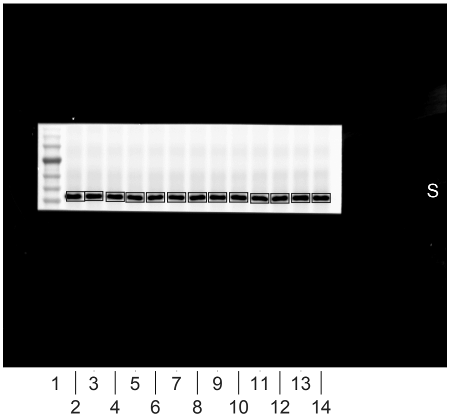


5.tif

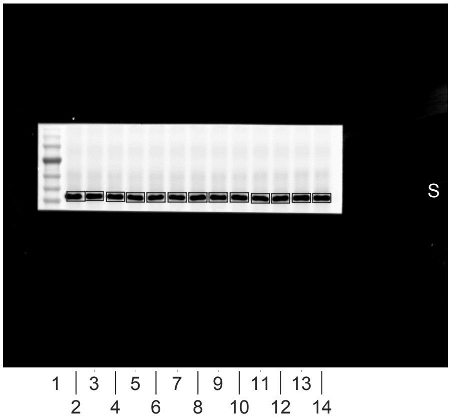


0.tif

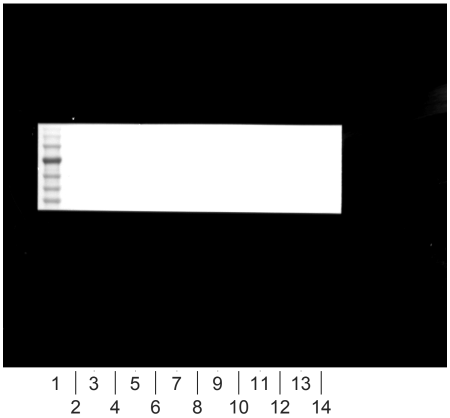


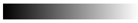

linear RGB

0 255

| Lane | Donor | Format | Treatment | Lane type |
|---|---|---|---|---|
| 1 | - | - | Marker | marker |
| 2 | D03 | 2D | Vehicle | sample |
| 3 | D03 | 2D | TGF-beta1 | sample |
| 4 | D03 | cast | Vehicle | sample |
| 5 | D03 | cast | TGF-beta1 | sample |
| 6 | D03 | printed | Vehicle | sample |
| 7 | D03 | printed | TGF-beta1 | sample |
| 8 | D01-D06 pool | 2D | TGF-beta1, T72 h | calibration pool |
| 9 | D06 | 2D | Vehicle | sample |
| 10 | D06 | 2D | TGF-beta1 | sample |
| 11 | D06 | cast | Vehicle | sample |
| 12 | D06 | cast | TGF-beta1 | sample |
| 13 | D06 | printed | Vehicle | sample |
| 14 | D06 | printed | TGF-beta1 | sample |
| 15 | - | - | loading buffer | empty |

1.tif

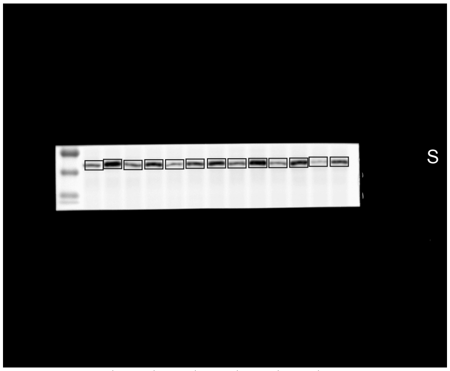


2.tif

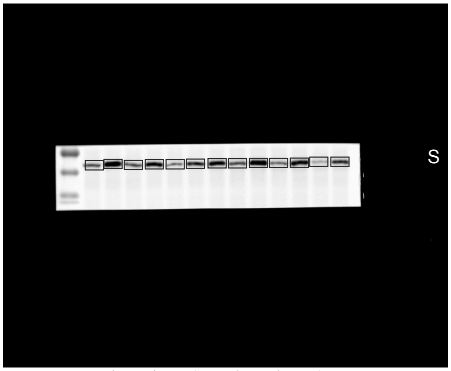


3.tif

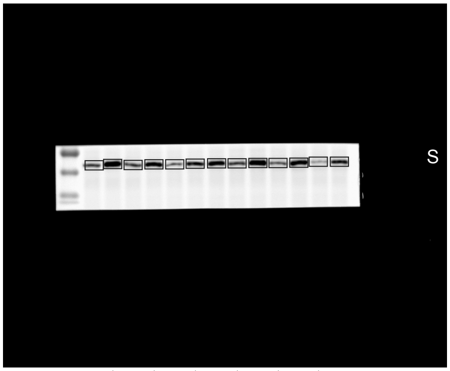


4.tif

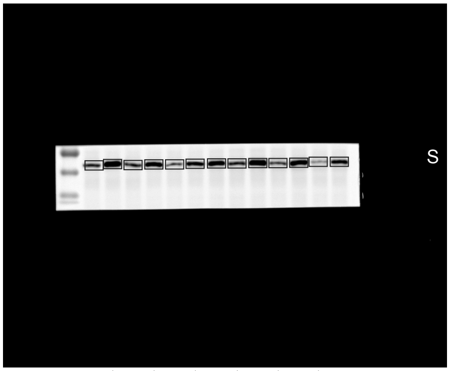


5.tif

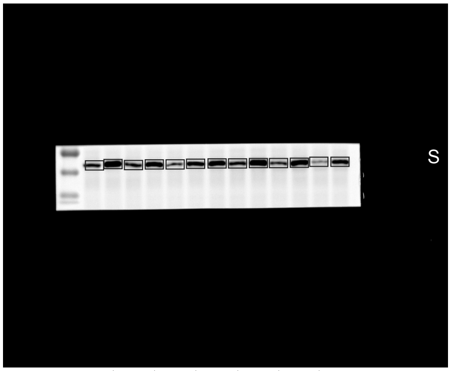


0.tif

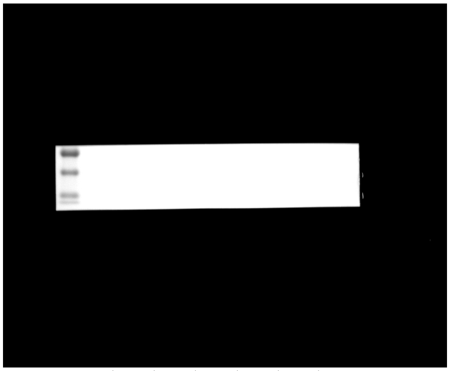


linear RGB

0 255

| Lane | Donor | Format | Treatment | Lane type |
|---|---|---|---|---|
| 1 | - | - | Marker | marker |
| 2 | D03 | 2D | Vehicle | sample |
| 3 | D03 | 2D | TGF-beta1 | sample |
| 4 | D03 | cast | Vehicle | sample |
| 5 | D03 | cast | TGF-beta1 | sample |
| 6 | D03 | printed | Vehicle | sample |
| 7 | D03 | printed | TGF-beta1 | sample |
| 8 | D01-D06 pool | 2D | TGF-beta1, T72 h | calibration pool |
| 9 | D06 | 2D | Vehicle | sample |
| 10 | D06 | 2D | TGF-beta1 | sample |
| 11 | D06 | cast | Vehicle | sample |
| 12 | D06 | cast | TGF-beta1 | sample |
| 13 | D06 | printed | Vehicle | sample |
| 14 | D06 | printed | TGF-beta1 | sample |
| 15 | - | - | loading buffer | empty |

1.tif *

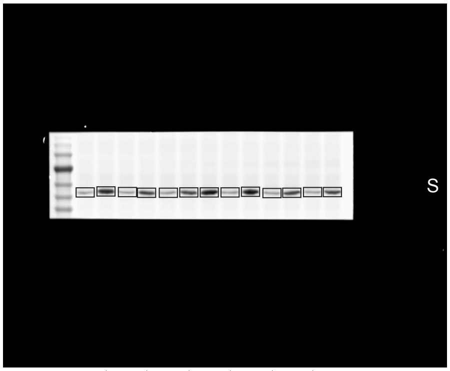


1 | 3 | 5 | 7 | 9 | 11 | 13 |
2 4 6 8 10 12 14

2.tif

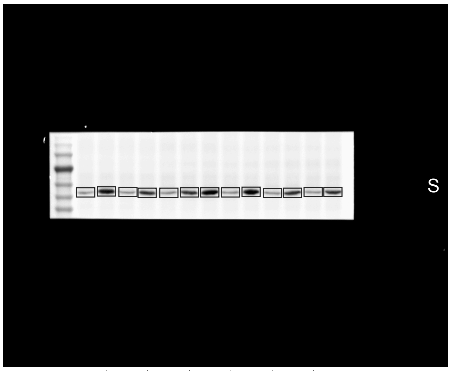


1 | 3 | 5 | 7 | 9 | 11 | 13 |
2 4 6 8 10 12 14

3.tif

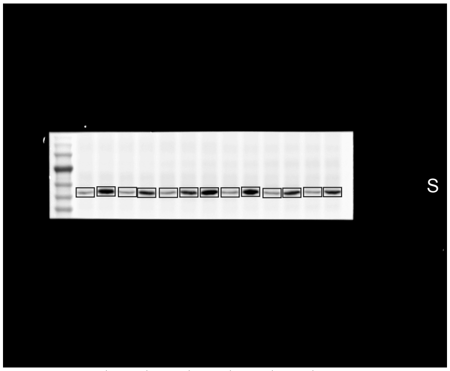


1 | 3 | 5 | 7 | 9 | 11 | 13 |
2 4 6 8 10 12 14

4.tif

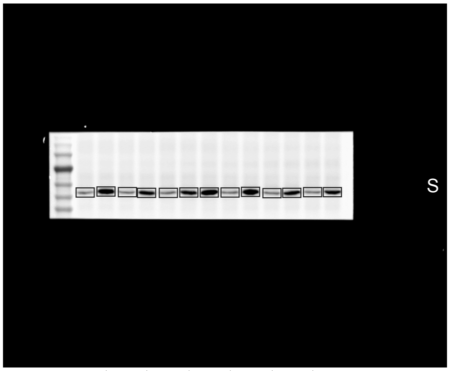


1 | 3 | 5 | 7 | 9 | 11 | 13 |
2 4 6 8 10 12 14

5.tif

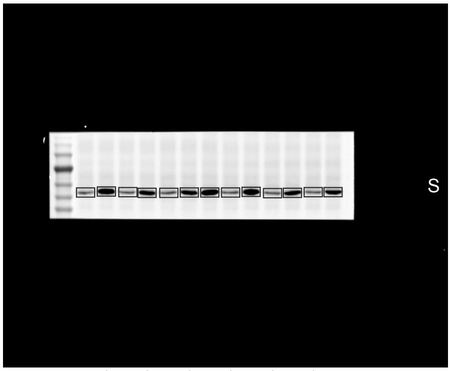


1 | 3 | 5 | 7 | 9 | 11 | 13 |
2 4 6 8 10 12 14

0.tif

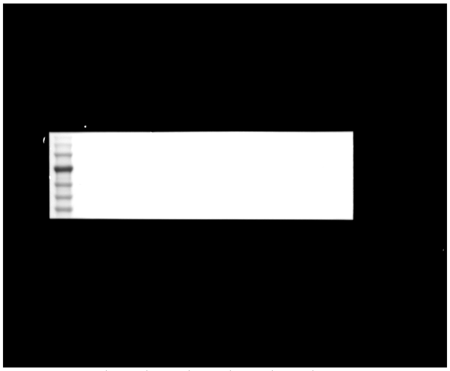

1 | 3 | 5 | 7 | 9 | 11 | 13 |
2 4 6 8 10 12 14

linear RGB

0 255

| Lane | Donor | Format | Treatment | Lane type |
|---|---|---|---|---|
| 1 | - | - | Marker | marker |
| 2 | D03 | 2D | Vehicle | sample |
| 3 | D03 | 2D | TGF-beta1 | sample |
| 4 | D03 | cast | Vehicle | sample |
| 5 | D03 | cast | TGF-beta1 | sample |
| 6 | D03 | printed | Vehicle | sample |
| 7 | D03 | printed | TGF-beta1 | sample |
| 8 | D01-D06 pool | 2D | TGF-beta1, T72 h | calibration pool |
| 9 | D06 | 2D | Vehicle | sample |
| 10 | D06 | 2D | TGF-beta1 | sample |
| 11 | D06 | cast | Vehicle | sample |
| 12 | D06 | cast | TGF-beta1 | sample |
| 13 | D06 | printed | Vehicle | sample |
| 14 | D06 | printed | TGF-beta1 | sample |
| 15 | - | - | loading buffer | empty |

1.tif *

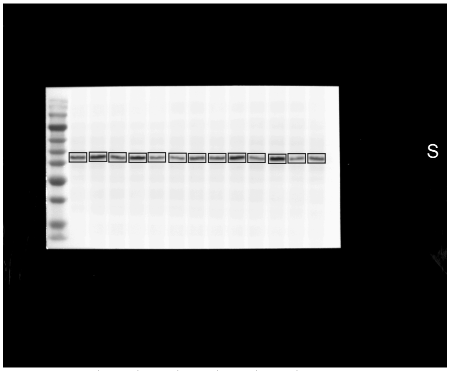


1 | 3 | 5 | 7 | 9 | 11 | 13 |
2 4 6 8 10 12 14

2.tif

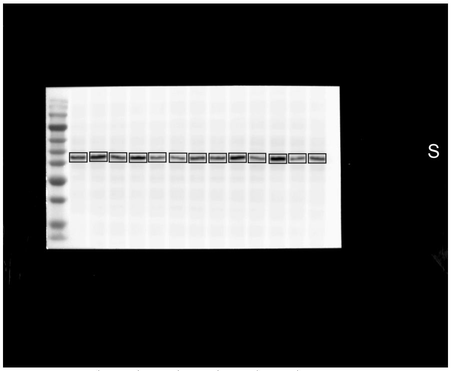


1 | 3 | 5 | 7 | 9 | 11 | 13 |
2 4 6 8 10 12 14

3.tif

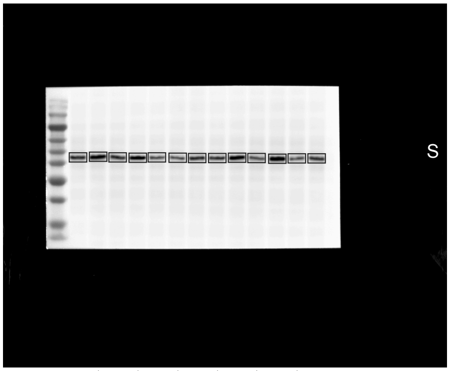


1 | 3 | 5 | 7 | 9 | 11 | 13 |
2 4 6 8 10 12 14

4.tif

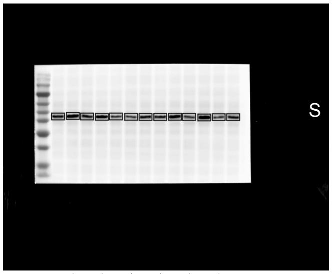


1 | 3 | 5 | 7 | 9 | 11 | 13 |
2 4 6 8 10 12 14

5.tif

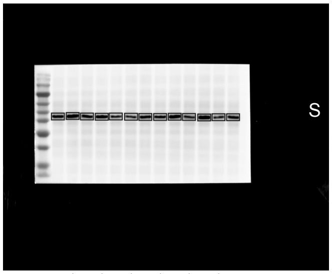


1 | 3 | 5 | 7 | 9 | 11 | 13 |
2 4 6 8 10 12 14

6.tif

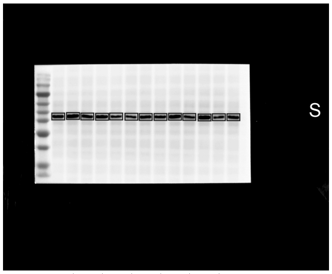


1 | 3 | 5 | 7 | 9 | 11 | 13 |
2 4 6 8 10 12 14

0.tif

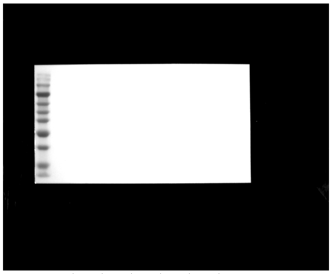

1 | 3 | 5 | 7 | 9 | 11 | 13 |
2 4 6 8 10 12 14

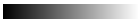

linear RGB

0 255

| Lane | Donor | Format | Treatment | Lane type |
|---|---|---|---|---|
| 1 | - | - | Marker | marker |
| 2 | D03 | 2D | Vehicle | sample |
| 3 | D03 | 2D | TGF-beta1 | sample |
| 4 | D03 | cast | Vehicle | sample |
| 5 | D03 | cast | TGF-beta1 | sample |
| 6 | D03 | printed | Vehicle | sample |
| 7 | D03 | printed | TGF-beta1 | sample |
| 8 | D01-D06 pool | 2D | TGF-beta1, T72 h | calibration pool |
| 9 | D06 | 2D | Vehicle | sample |
| 10 | D06 | 2D | TGF-beta1 | sample |
| 11 | D06 | cast | Vehicle | sample |
| 12 | D06 | cast | TGF-beta1 | sample |
| 13 | D06 | printed | Vehicle | sample |
| 14 | D06 | printed | TGF-beta1 | sample |
| 15 | - | - | loading buffer | empty |

2.tif *

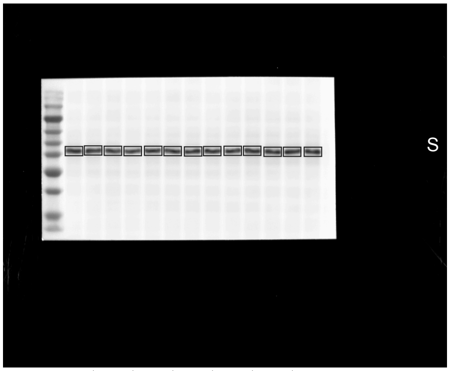


1 3 5 7 9 11 13
2 4 6 8 10 12 14

1.tif

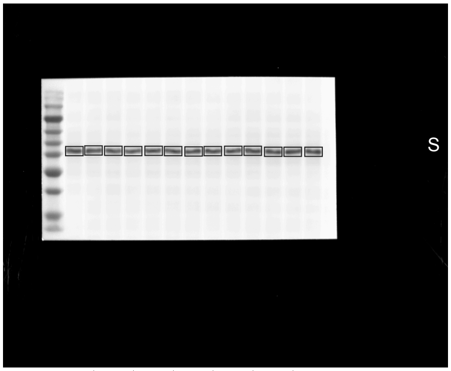


1 3 5 7 9 11 13
2 4 6 8 10 12 14

3.tif

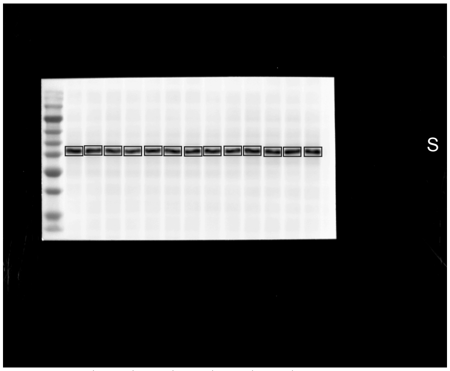


1 3 5 7 9 11 13
2 4 6 8 10 12 14

4.tif

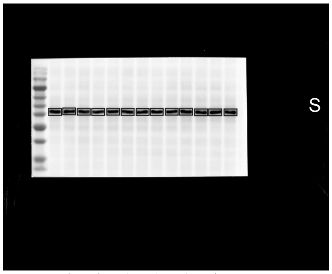


1 3 5 7 9 11 13
2 4 6 8 10 12 14

5.tif

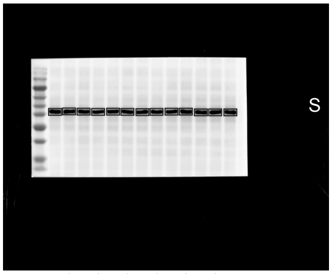


1 3 5 7 9 11 13
2 4 6 8 10 12 14

6.tif

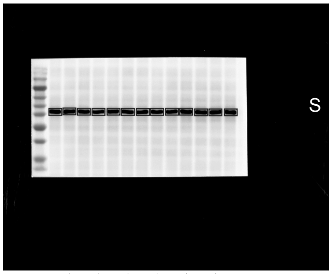


1 3 5 7 9 11 13
2 4 6 8 10 12 14

0.tif

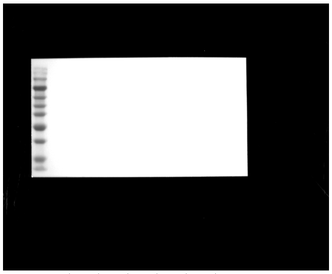

1 3 5 7 9 11 13
2 4 6 8 10 12 14

linear RGB
0 255

| Lane | Donor | Format | Treatment | Lane type |
|---|---|---|---|---|
| 1 | - | - | Marker | marker |
| 2 | D03 | 2D | Vehicle | sample |
| 3 | D03 | 2D | TGF-beta1 | sample |
| 4 | D03 | cast | Vehicle | sample |
| 5 | D03 | cast | TGF-beta1 | sample |
| 6 | D03 | printed | Vehicle | sample |
| 7 | D03 | printed | TGF-beta1 | sample |
| 8 | D01-D06 pool | 2D | TGF-beta1, T72 h | calibration pool |
| 9 | D06 | 2D | Vehicle | sample |
| 10 | D06 | 2D | TGF-beta1 | sample |
| 11 | D06 | cast | Vehicle | sample |
| 12 | D06 | cast | TGF-beta1 | sample |
| 13 | D06 | printed | Vehicle | sample |
| 14 | D06 | printed | TGF-beta1 | sample |
| 15 | - | - | loading buffer | empty |

1.tif

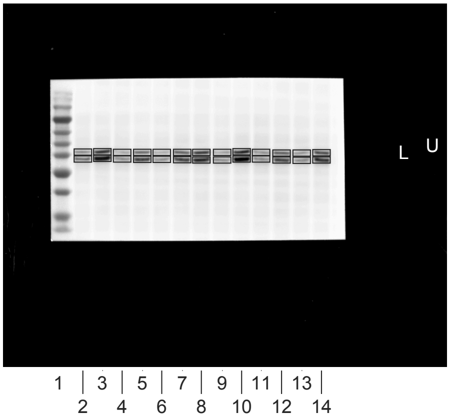


2.tif

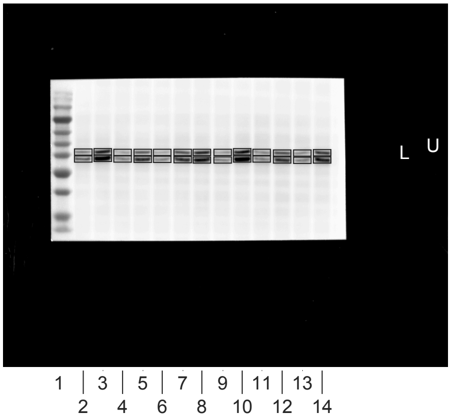


3.tif

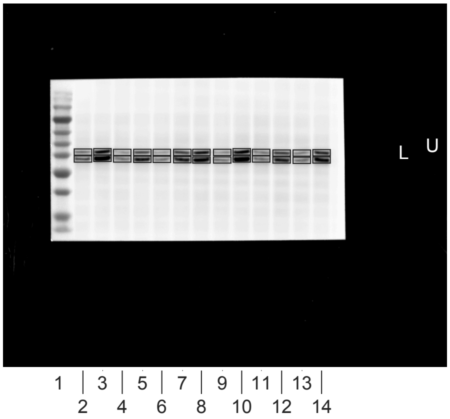


4.tif

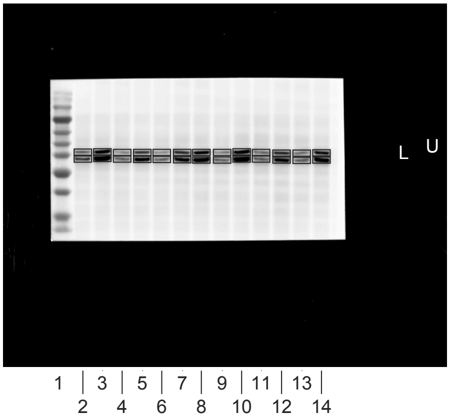


5.tif

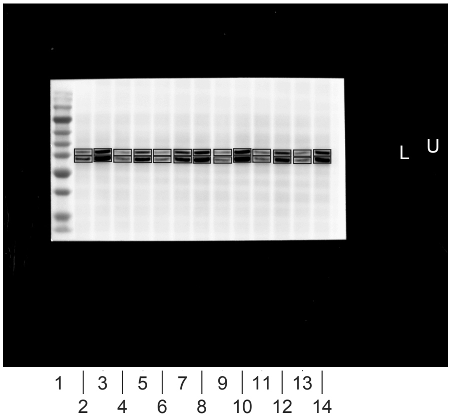


0.tif

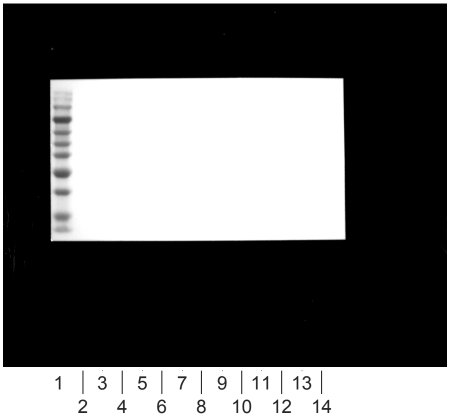


linear RGB
0 255

| Lane | Donor | Format | Treatment | Lane type |
|---|---|---|---|---|
| 1 | - | - | Marker | marker |
| 2 | D03 | 2D | Vehicle | sample |
| 3 | D03 | 2D | TGF-beta1 | sample |
| 4 | D03 | cast | Vehicle | sample |
| 5 | D03 | cast | TGF-beta1 | sample |
| 6 | D03 | printed | Vehicle | sample |
| 7 | D03 | printed | TGF-beta1 | sample |
| 8 | D01-D06 pool | 2D | TGF-beta1, T72 h | calibration pool |
| 9 | D06 | 2D | Vehicle | sample |
| 10 | D06 | 2D | TGF-beta1 | sample |
| 11 | D06 | cast | Vehicle | sample |
| 12 | D06 | cast | TGF-beta1 | sample |
| 13 | D06 | printed | Vehicle | sample |
| 14 | D06 | printed | TGF-beta1 | sample |
| 15 | - | - | loading buffer | empty |

2.tif *

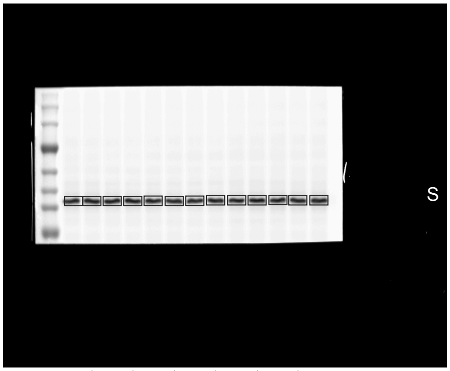


1.tif

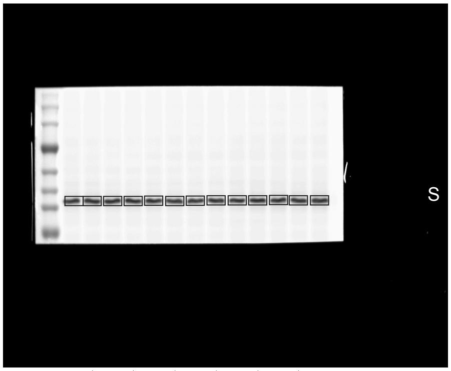


3.tif

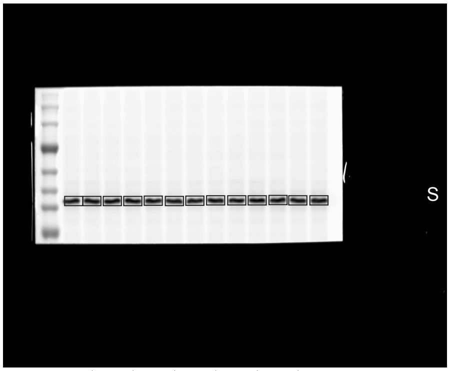


4.tif

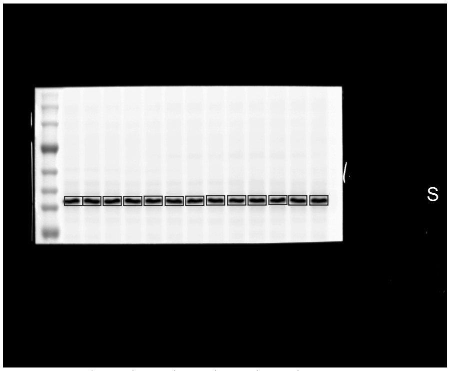


5.tif

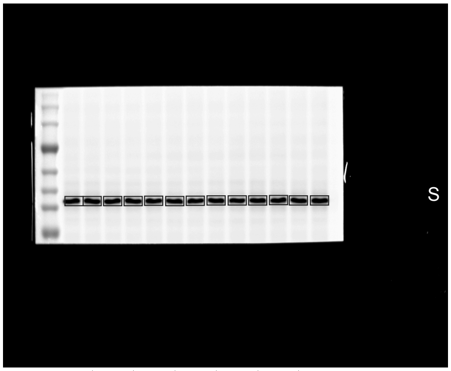


0.tif

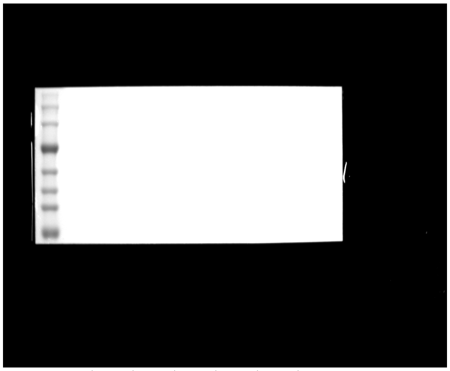


linear RGB
0 255

| Lane | Donor | Format | Treatment | Lane type |
|---|---|---|---|---|
| 1 | - | - | Marker | marker |
| 2 | D01 | 2D | Vehicle | sample |
| 3 | D01 | 2D | TGF-beta1 | sample |
| 4 | D01 | cast | Vehicle | sample |
| 5 | D01 | cast | TGF-beta1 | sample |
| 6 | D01 | printed | Vehicle | sample |
| 7 | D01 | printed | TGF-beta1 | sample |
| 8 | D01-D06 pool | 2D | TGF-beta1, T72 h | calibration pool |
| 9 | D02 | 2D | Vehicle | sample |
| 10 | D02 | 2D | TGF-beta1 | sample |
| 11 | D02 | cast | Vehicle | sample |
| 12 | D02 | cast | TGF-beta1 | sample |
| 13 | D02 | printed | Vehicle | sample |
| 14 | D02 | printed | TGF-beta1 | sample |
| 15 | - | - | loading buffer | empty |

1.tif *

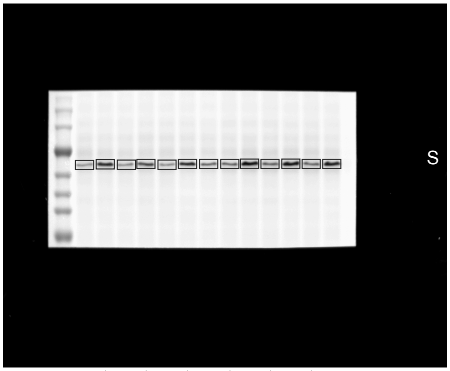


1 3 5 7 9 11 13
2 4 6 8 10 12 14

2.tif

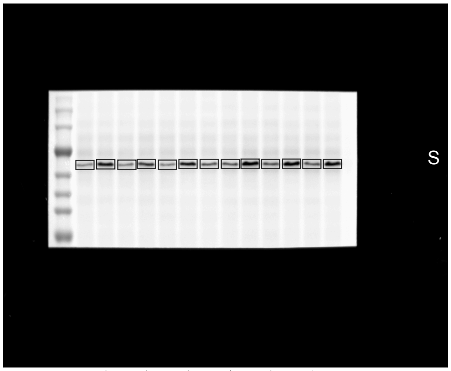


1 3 5 7 9 11 13
2 4 6 8 10 12 14

3.tif

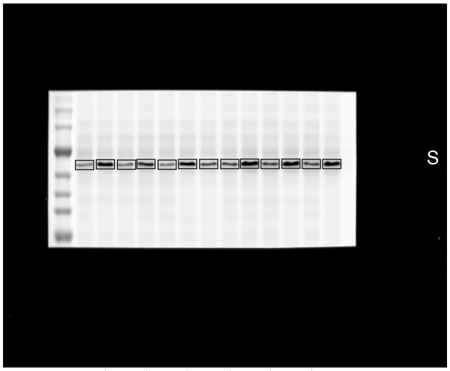


1 3 5 7 9 11 13
2 4 6 8 10 12 14

4.tif

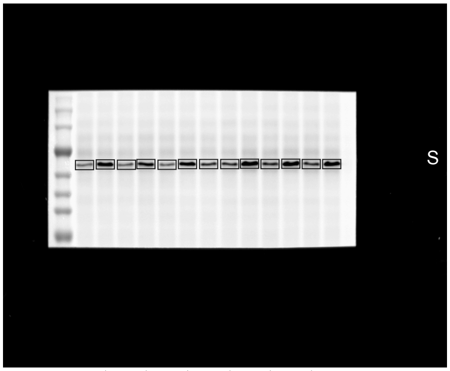


1 3 5 7 9 11 13
2 4 6 8 10 12 14

5.tif

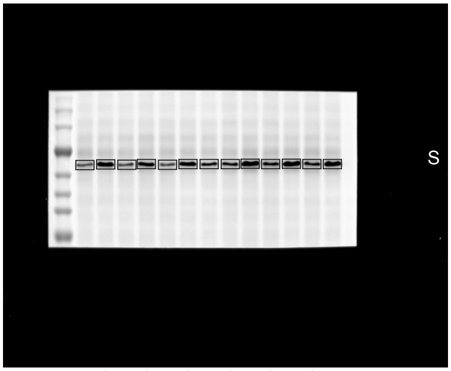


1 3 5 7 9 11 13
2 4 6 8 10 12 14

0.tif

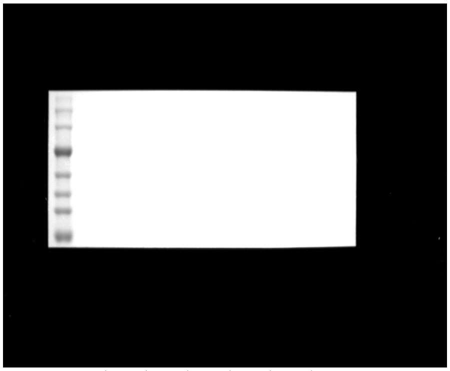

1 3 5 7 9 11 13
2 4 6 8 10 12 14

linear RGB

0 255

| Lane | Donor | Format | Treatment | Lane type |
|---|---|---|---|---|
| 1 | - | - | Marker | marker |
| 2 | D01 | 2D | Vehicle | sample |
| 3 | D01 | 2D | TGF-beta1 | sample |
| 4 | D01 | cast | Vehicle | sample |
| 5 | D01 | cast | TGF-beta1 | sample |
| 6 | D01 | printed | Vehicle | sample |
| 7 | D01 | printed | TGF-beta1 | sample |
| 8 | D01-D06 pool | 2D | TGF-beta1, T72 h | calibration pool |
| 9 | D02 | 2D | Vehicle | sample |
| 10 | D02 | 2D | TGF-beta1 | sample |
| 11 | D02 | cast | Vehicle | sample |
| 12 | D02 | cast | TGF-beta1 | sample |
| 13 | D02 | printed | Vehicle | sample |
| 14 | D02 | printed | TGF-beta1 | sample |
| 15 | - | - | loading buffer | empty |

2.tif *

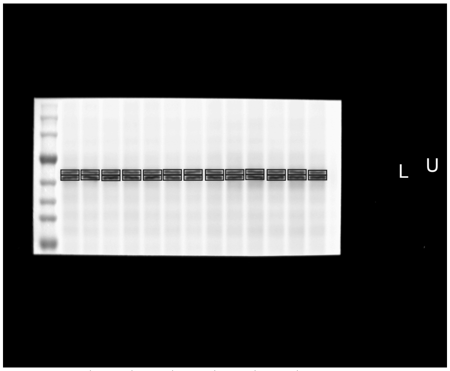


1 3 5 7 9 11 13
2 4 6 8 10 12 14

1.tif

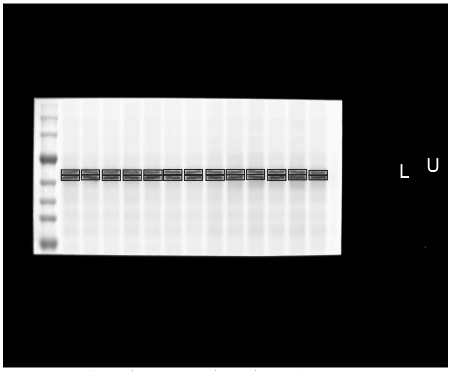


1 3 5 7 9 11 13
2 4 6 8 10 12 14

3.tif

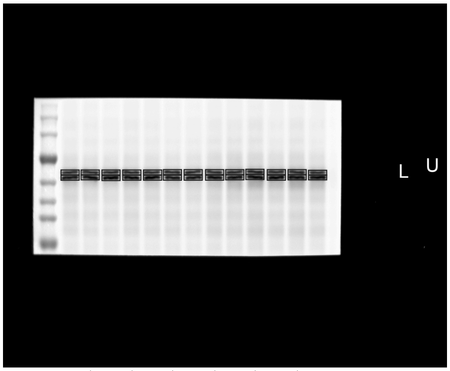


1 3 5 7 9 11 13
2 4 6 8 10 12 14

4.tif

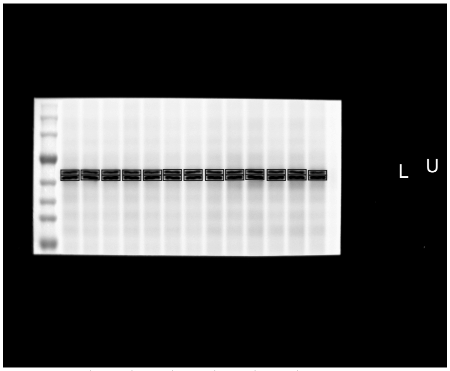


1 3 5 7 9 11 13
2 4 6 8 10 12 14

5.tif

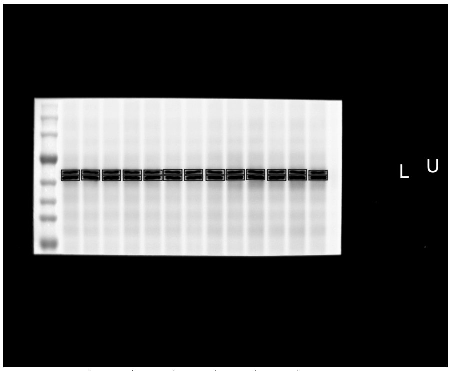


1 3 5 7 9 11 13
2 4 6 8 10 12 14

0.tif

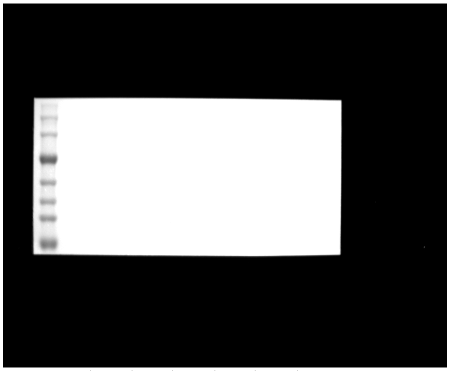

1 3 5 7 9 11 13
2 4 6 8 10 12 14

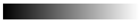

linear RGB

0 255

| Lane | Donor | Format | Treatment | Lane type |
|---|---|---|---|---|
| 1 | - | - | Marker | marker |
| 2 | D01 | 2D | Vehicle | sample |
| 3 | D01 | 2D | TGF-beta1 | sample |
| 4 | D01 | cast | Vehicle | sample |
| 5 | D01 | cast | TGF-beta1 | sample |
| 6 | D01 | printed | Vehicle | sample |
| 7 | D01 | printed | TGF-beta1 | sample |
| 8 | D01-D06 pool | 2D | TGF-beta1, T72 h | calibration pool |
| 9 | D02 | 2D | Vehicle | sample |
| 10 | D02 | 2D | TGF-beta1 | sample |
| 11 | D02 | cast | Vehicle | sample |
| 12 | D02 | cast | TGF-beta1 | sample |
| 13 | D02 | printed | Vehicle | sample |
| 14 | D02 | printed | TGF-beta1 | sample |
| 15 | - | - | loading buffer | empty |

2.tif *

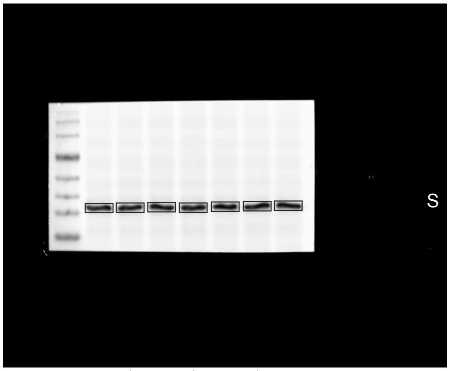


1 3 5 7
2 4 6 8

1.tif

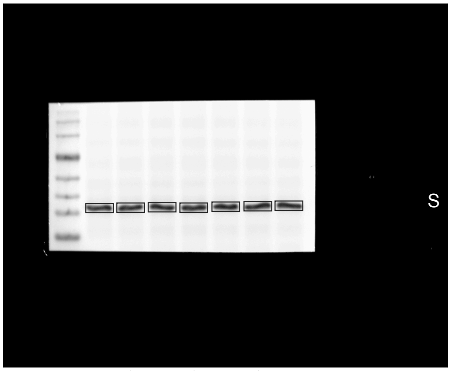


1 3 5 7
2 4 6 8

3.tif

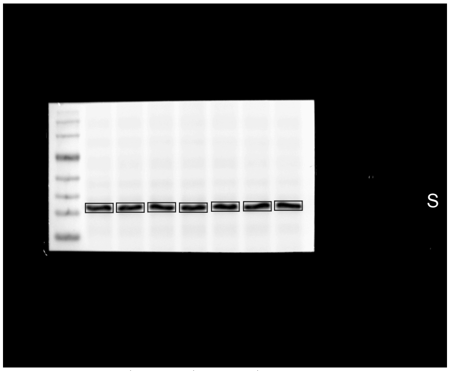


1 3 5 7
2 4 6 8

4.tif

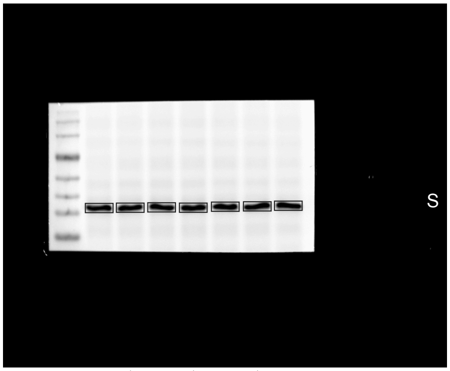


1 3 5 7
2 4 6 8

5.tif

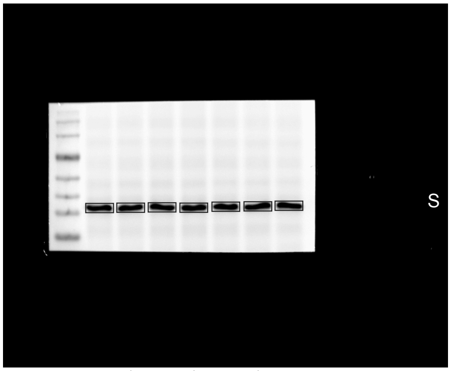


1 3 5 7
2 4 6 8

0.tif

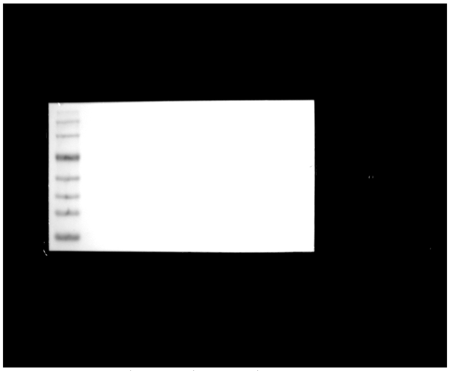

1 3 5 7
2 4 6 8

linear RGB

0 255

| Lane | Donor | Format | Treatment | Lane type |
|---|---|---|---|---|
| 1 | - | - | Marker | marker |
| 2 | D04 | 2D | Vehicle | sample |
| 3 | D04 | 2D | TGF-beta1 | sample |
| 4 | D04 | cast | Vehicle | sample |
| 5 | D04 | cast | TGF-beta1 | sample |
| 6 | D04 | printed | Vehicle | sample |
| 7 | D04 | printed | TGF-beta1 | sample |
| 8 | D01-D06 pool | 2D | TGF-beta1, T72 h | calibration pool |
| 9 | - | - | loading buffer | empty |
| 10 | - | - | loading buffer | empty |
| 11 | - | - | loading buffer | empty |
| 12 | - | - | loading buffer | empty |
| 13 | - | - | loading buffer | empty |
| 14 | - | - | loading buffer | empty |
| 15 | - | - | loading buffer | empty |

1.tif *

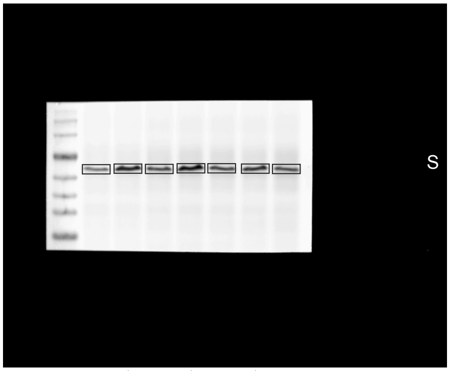


1 3 5 7
2 4 6 8

2.tif

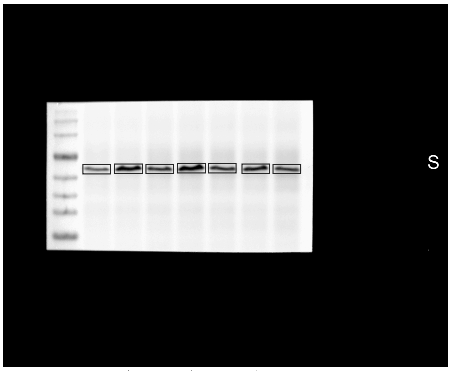


1 3 5 7
2 4 6 8

3.tif

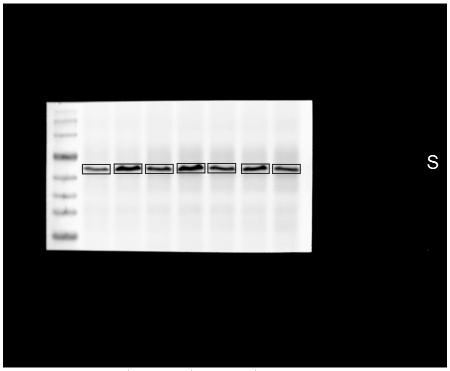


1 3 5 7
2 4 6 8

4.tif

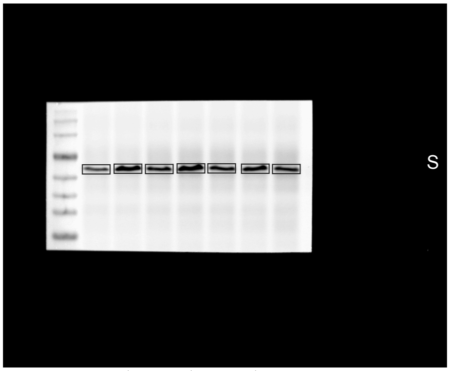


1 3 5 7
2 4 6 8

5.tif

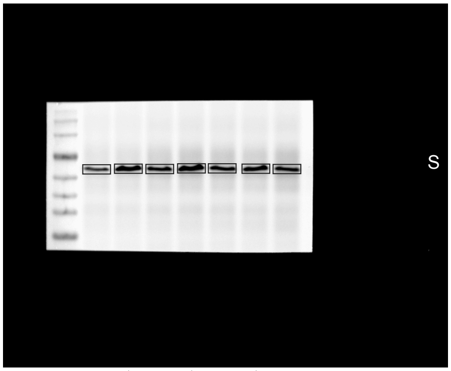


1 3 5 7
2 4 6 8

0.tif

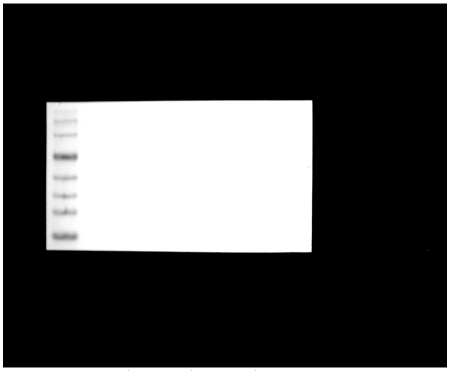

1 3 5 7
2 4 6 8

linear RGB

0 255

| Lane | Donor | Format | Treatment | Lane type |
|---|---|---|---|---|
| 1 | - | - | Marker | marker |
| 2 | D04 | 2D | Vehicle | sample |
| 3 | D04 | 2D | TGF-beta1 | sample |
| 4 | D04 | cast | Vehicle | sample |
| 5 | D04 | cast | TGF-beta1 | sample |
| 6 | D04 | printed | Vehicle | sample |
| 7 | D04 | printed | TGF-beta1 | sample |
| 8 | D01-D06 pool | 2D | TGF-beta1, T72 h | calibration pool |
| 9 | - | - | loading buffer | empty |
| 10 | - | - | loading buffer | empty |
| 11 | - | - | loading buffer | empty |
| 12 | - | - | loading buffer | empty |
| 13 | - | - | loading buffer | empty |
| 14 | - | - | loading buffer | empty |
| 15 | - | - | loading buffer | empty |

2.tif *

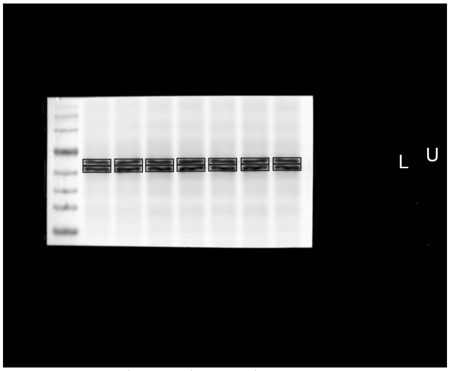


1 3 5 7
2 4 6 8

1.tif

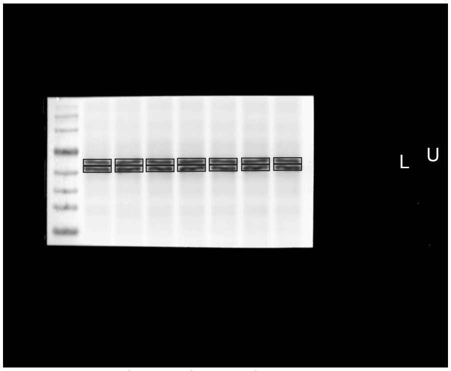


1 3 5 7
2 4 6 8

3.tif

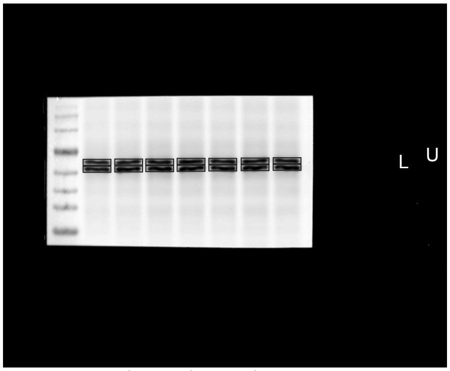


1 3 5 7
2 4 6 8

4.tif

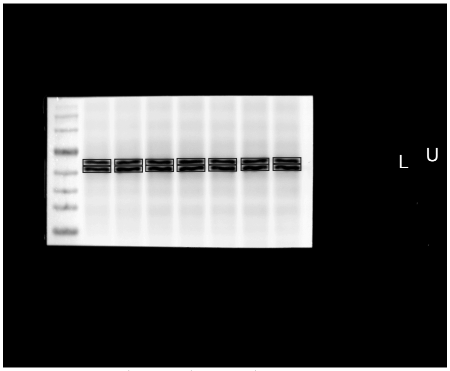


1 3 5 7
2 4 6 8

5.tif

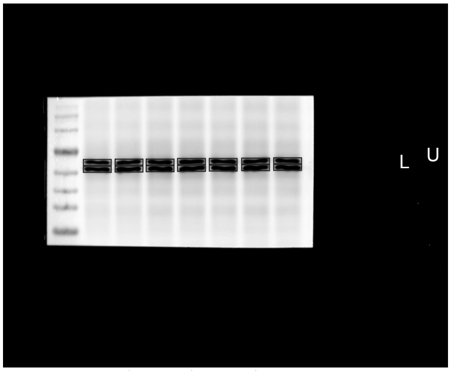


1 3 5 7
2 4 6 8

0.tif

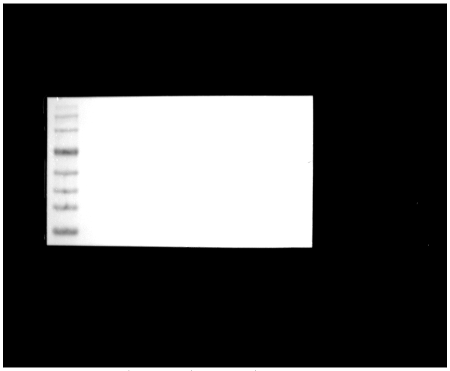

1 3 5 7
2 4 6 8

linear RGB
0 255

| Lane | Donor | Format | Treatment | Lane type |
|---|---|---|---|---|
| 1 | - | - | Marker | marker |
| 2 | D04 | 2D | Vehicle | sample |
| 3 | D04 | 2D | TGF-beta1 | sample |
| 4 | D04 | cast | Vehicle | sample |
| 5 | D04 | cast | TGF-beta1 | sample |
| 6 | D04 | printed | Vehicle | sample |
| 7 | D04 | printed | TGF-beta1 | sample |
| 8 | D01-D06 pool | 2D | TGF-beta1, T72 h | calibration pool |
| 9 | - | - | loading buffer | empty |
| 10 | - | - | loading buffer | empty |
| 11 | - | - | loading buffer | empty |
| 12 | - | - | loading buffer | empty |
| 13 | - | - | loading buffer | empty |
| 14 | - | - | loading buffer | empty |
| 15 | - | - | loading buffer | empty |

1.tif *

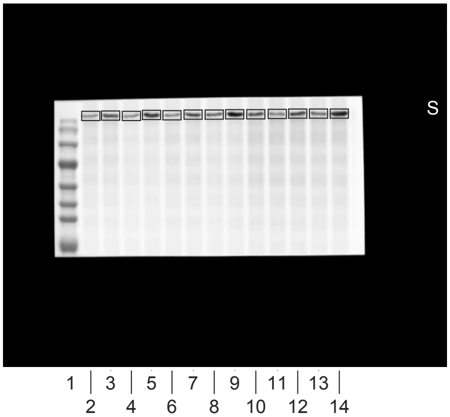


2.tif

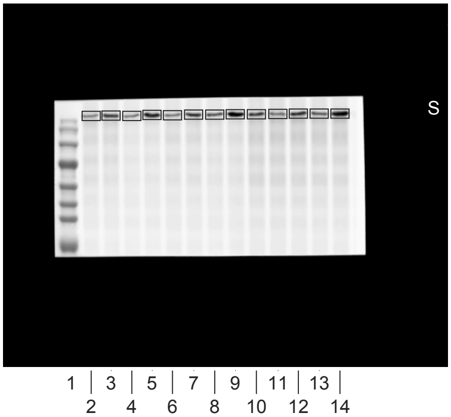


3.tif

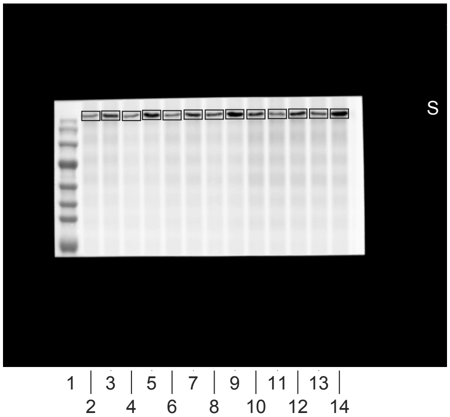


4.tif

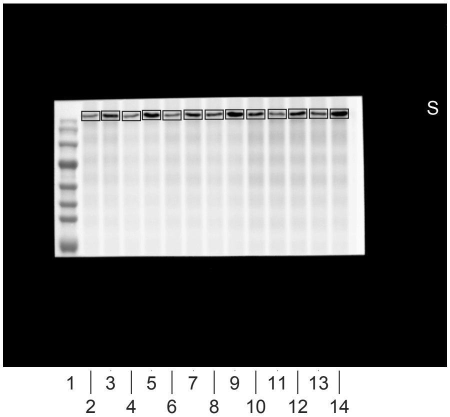


5.tif

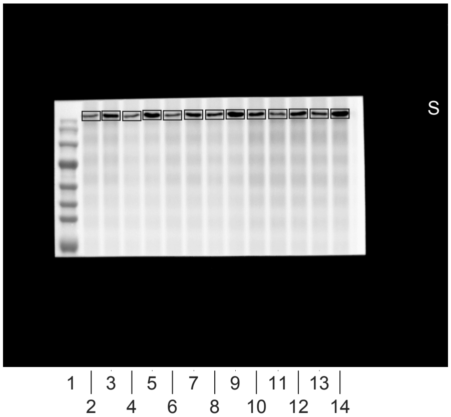


0.tif

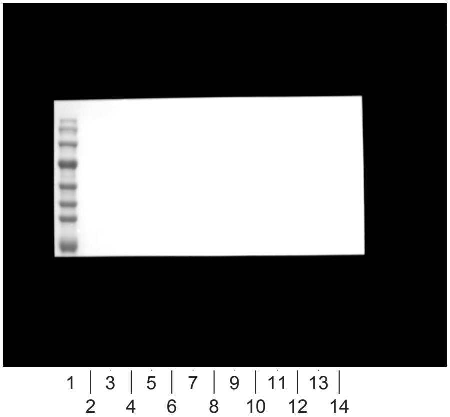


linear RGB

0 255

| Lane | Donor | Format | Treatment | Lane type |
|---|---|---|---|---|
| 1 | - | - | Marker | marker |
| 2 | D01 | 2D | Vehicle | sample |
| 3 | D01 | 2D | TGF-beta1 | sample |
| 4 | D01 | printed | Vehicle | sample |
| 5 | D01 | printed | TGF-beta1 | sample |
| 6 | D02 | 2D | Vehicle | sample |
| 7 | D02 | 2D | TGF-beta1 | sample |
| 8 | D02 | printed | Vehicle | sample |
| 9 | D02 | printed | TGF-beta1 | sample |
| 10 | D01-D06 pool | 2D | TGF-beta1, T72 h | calibration pool |
| 11 | D03 | 2D | Vehicle | sample |
| 12 | D03 | 2D | TGF-beta1 | sample |
| 13 | D03 | printed | Vehicle | sample |
| 14 | D03 | printed | TGF-beta1 | sample |
| 15 | - | - | loading buffer | empty |

1.tif *

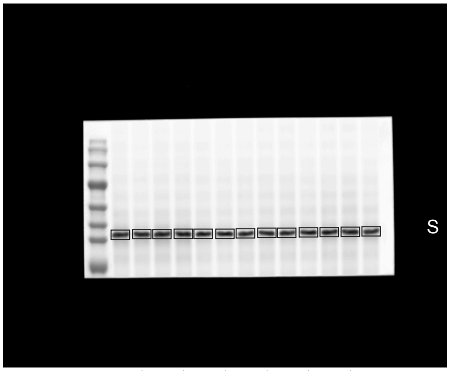


1 | 3 | 5 | 7 | 9 | 11 | 13 |
2 4 6 8 10 12 14

2.tif

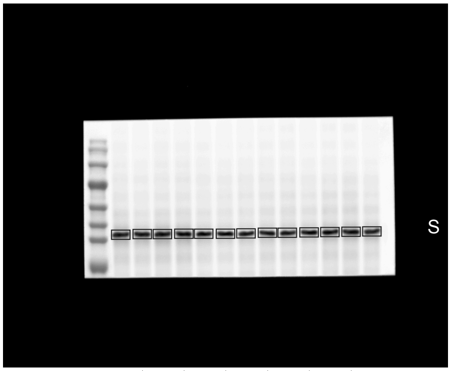


1 | 3 | 5 | 7 | 9 | 11 | 13 |
2 4 6 8 10 12 14

3.tif

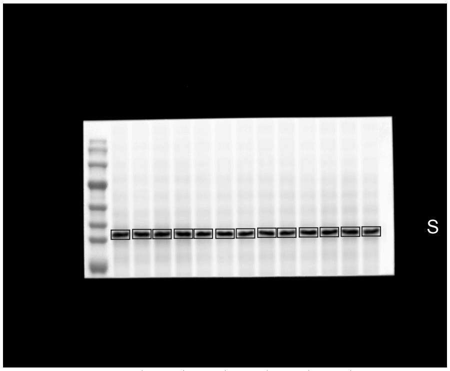


1 | 3 | 5 | 7 | 9 | 11 | 13 |
2 4 6 8 10 12 14

4.tif

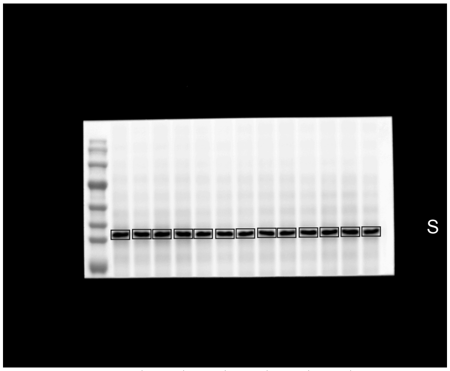


1 | 3 | 5 | 7 | 9 | 11 | 13 |
2 4 6 8 10 12 14

5.tif

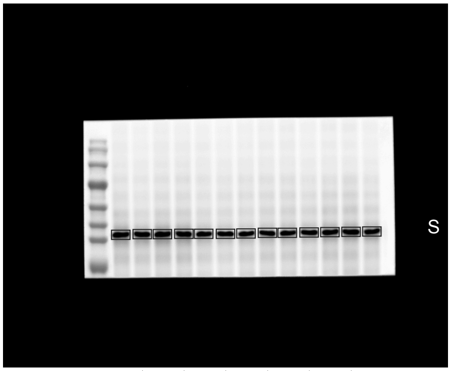


1 | 3 | 5 | 7 | 9 | 11 | 13 |
2 4 6 8 10 12 14

0.tif

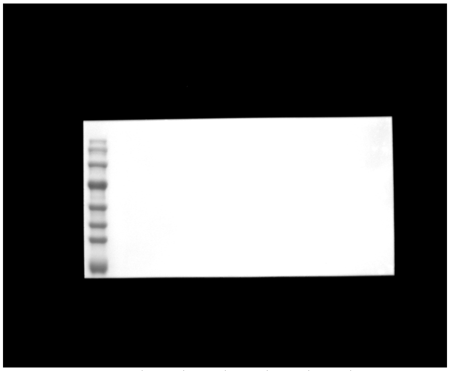

1 | 3 | 5 | 7 | 9 | 11 | 13 |
2 4 6 8 10 12 14

linear RGB
0 255

| Lane | Donor | Format | Treatment | Lane type |
|---|---|---|---|---|
| 1 | - | - | Marker | marker |
| 2 | D01 | 2D | Vehicle | sample |
| 3 | D01 | 2D | TGF-beta1 | sample |
| 4 | D01 | printed | Vehicle | sample |
| 5 | D01 | printed | TGF-beta1 | sample |
| 6 | D02 | 2D | Vehicle | sample |
| 7 | D02 | 2D | TGF-beta1 | sample |
| 8 | D02 | printed | Vehicle | sample |
| 9 | D02 | printed | TGF-beta1 | sample |
| 10 | D01-D06 pool | 2D | TGF-beta1, T72 h | calibration pool |
| 11 | D03 | 2D | Vehicle | sample |
| 12 | D03 | 2D | TGF-beta1 | sample |
| 13 | D03 | printed | Vehicle | sample |
| 14 | D03 | printed | TGF-beta1 | sample |
| 15 | - | - | loading buffer | empty |

1.tif

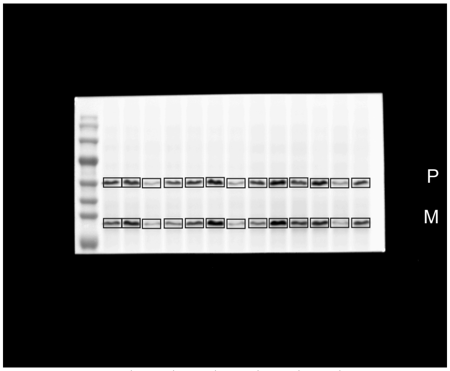


1 3 5 7 9 11 13
2 4 6 8 10 12 14

2.tif

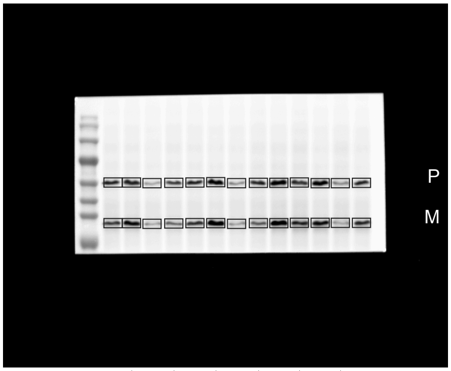


1 3 5 7 9 11 13
2 4 6 8 10 12 14

3.tif

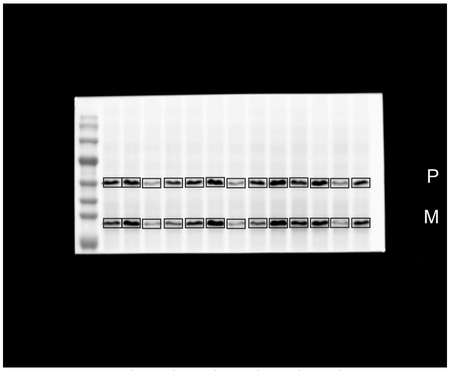


1 3 5 7 9 11 13
2 4 6 8 10 12 14

4.tif

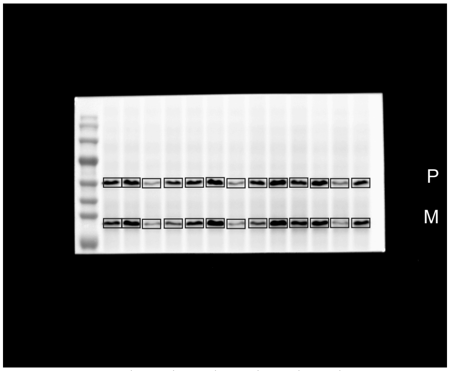


1 3 5 7 9 11 13
2 4 6 8 10 12 14

5.tif

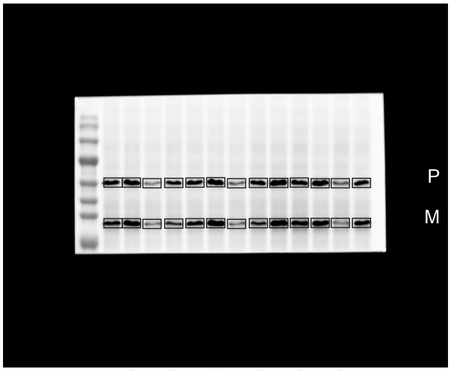


1 3 5 7 9 11 13
2 4 6 8 10 12 14

0.tif

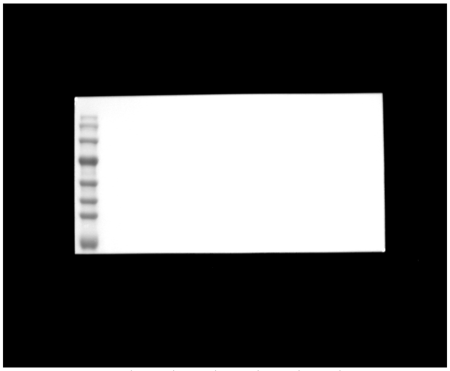

1 3 5 7 9 11 13
2 4 6 8 10 12 14

linear RGB

0 255

| Lane | Donor | Format | Treatment | Lane type |
|---|---|---|---|---|
| 1 | - | - | Marker | marker |
| 2 | D01 | 2D | Vehicle | sample |
| 3 | D01 | 2D | TGF-beta1 | sample |
| 4 | D01 | printed | Vehicle | sample |
| 5 | D01 | printed | TGF-beta1 | sample |
| 6 | D02 | 2D | Vehicle | sample |
| 7 | D02 | 2D | TGF-beta1 | sample |
| 8 | D02 | printed | Vehicle | sample |
| 9 | D02 | printed | TGF-beta1 | sample |
| 10 | D01-D06 pool | 2D | TGF-beta1, T72 h | calibration pool |
| 11 | D03 | 2D | Vehicle | sample |
| 12 | D03 | 2D | TGF-beta1 | sample |
| 13 | D03 | printed | Vehicle | sample |
| 14 | D03 | printed | TGF-beta1 | sample |
| 15 | - | - | loading buffer | empty |

1.tif *

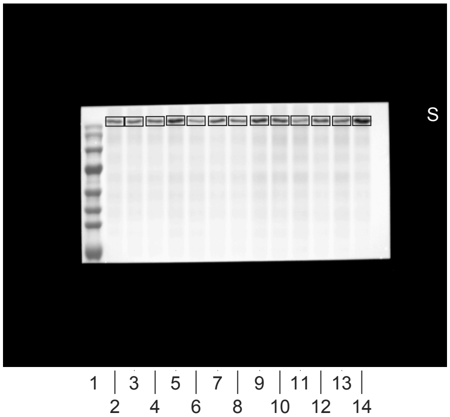


2.tif

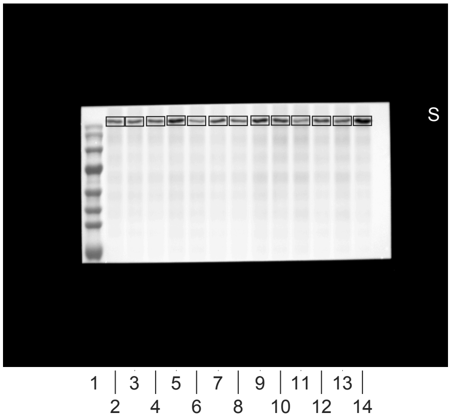


3.tif

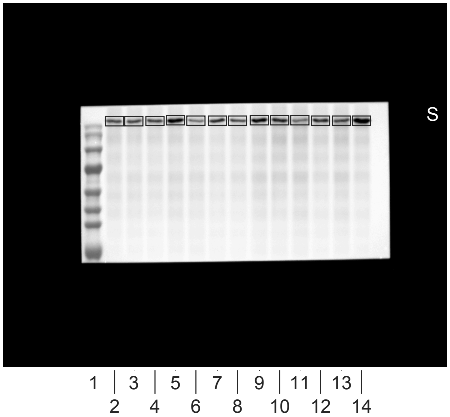


4.tif

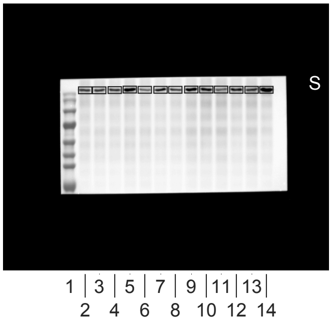


5.tif

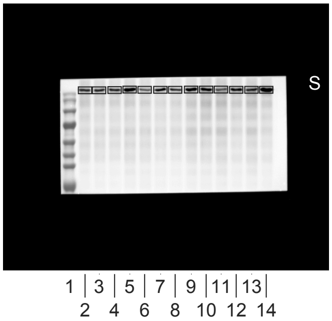


6.tif

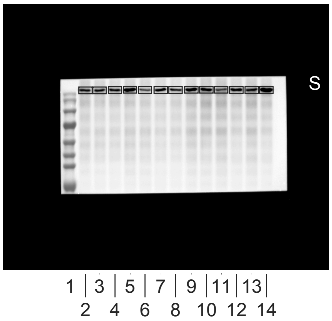


0.tif

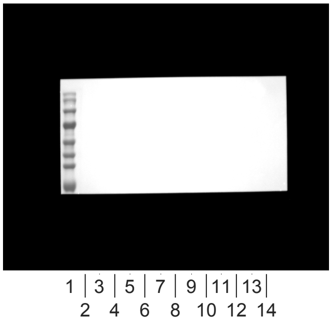


linear RGB
0 255

| Lane | Donor | Format | Treatment | Lane type |
|---|---|---|---|---|
| 1 | - | - | Marker | marker |
| 2 | D04 | 2D | Vehicle | sample |
| 3 | D04 | 2D | TGF-beta1 | sample |
| 4 | D04 | printed | Vehicle | sample |
| 5 | D04 | printed | TGF-beta1 | sample |
| 6 | D05 | 2D | Vehicle | sample |
| 7 | D05 | 2D | TGF-beta1 | sample |
| 8 | D05 | printed | Vehicle | sample |
| 9 | D05 | printed | TGF-beta1 | sample |
| 10 | D01-D06 pool | 2D | TGF-beta1, T72 h | calibration pool |
| 11 | D06 | 2D | Vehicle | sample |
| 12 | D06 | 2D | TGF-beta1 | sample |
| 13 | D06 | printed | Vehicle | sample |
| 14 | D06 | printed | TGF-beta1 | sample |
| 15 | - | - | loading buffer | empty |

2.tif *

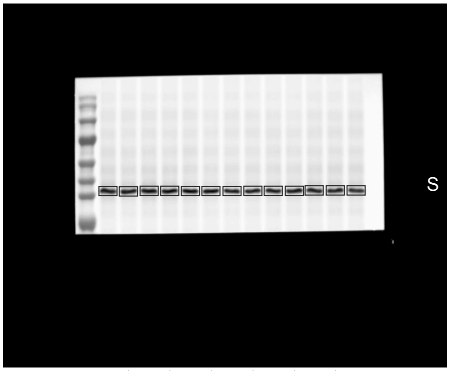


1.tif

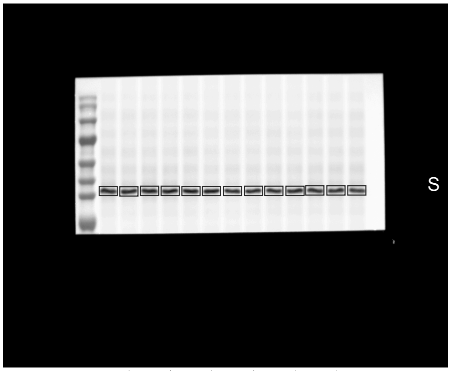


3.tif

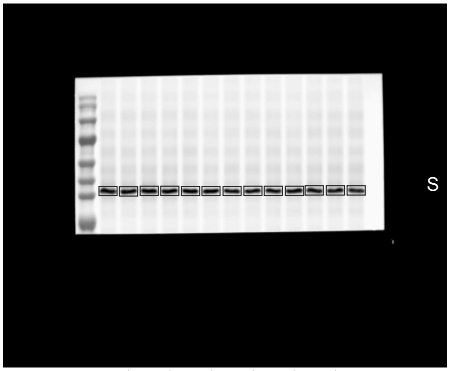


4.tif

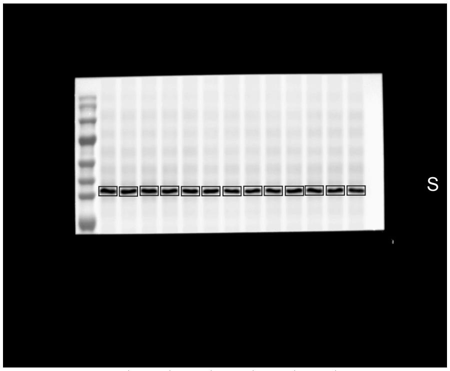


5.tif

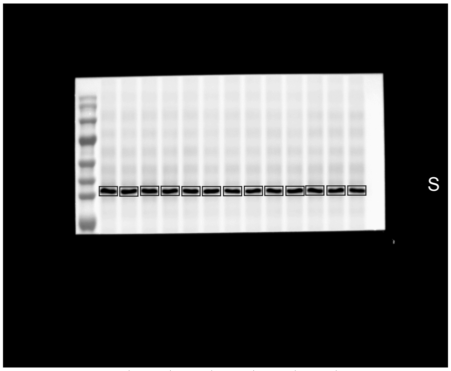


0.tif

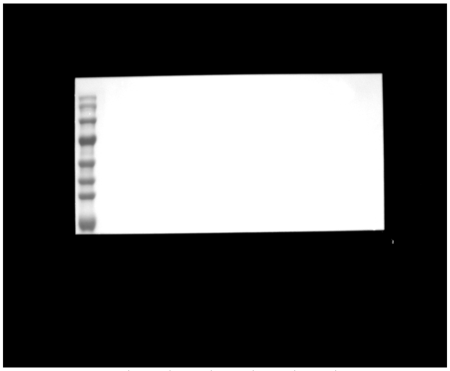


linear RGB
0 255

| Lane | Donor | Format | Treatment | Lane type |
|---|---|---|---|---|
| 1 | - | - | Marker | marker |
| 2 | D04 | 2D | Vehicle | sample |
| 3 | D04 | 2D | TGF-beta1 | sample |
| 4 | D04 | printed | Vehicle | sample |
| 5 | D04 | printed | TGF-beta1 | sample |
| 6 | D05 | 2D | Vehicle | sample |
| 7 | D05 | 2D | TGF-beta1 | sample |
| 8 | D05 | printed | Vehicle | sample |
| 9 | D05 | printed | TGF-beta1 | sample |
| 10 | D01-D06 pool | 2D | TGF-beta1, T72 h | calibration pool |
| 11 | D06 | 2D | Vehicle | sample |
| 12 | D06 | 2D | TGF-beta1 | sample |
| 13 | D06 | printed | Vehicle | sample |
| 14 | D06 | printed | TGF-beta1 | sample |
| 15 | - | - | loading buffer | empty |

1.tif

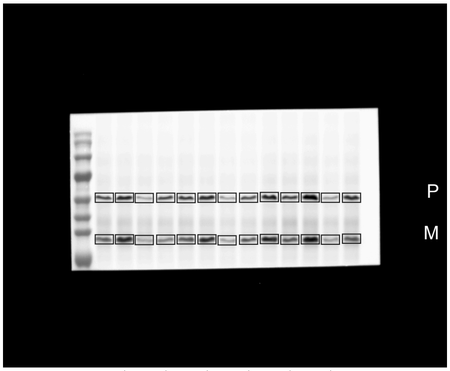


1 3 5 7 9 11 13
2 4 6 8 10 12 14

2.tif

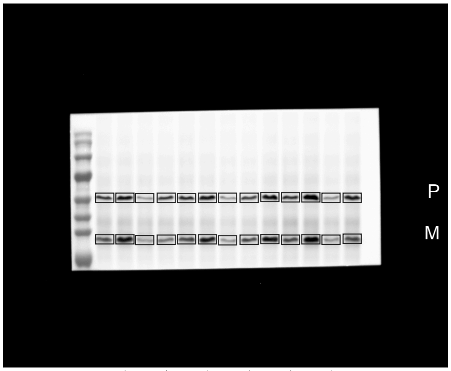


1 3 5 7 9 11 13
2 4 6 8 10 12 14

3.tif

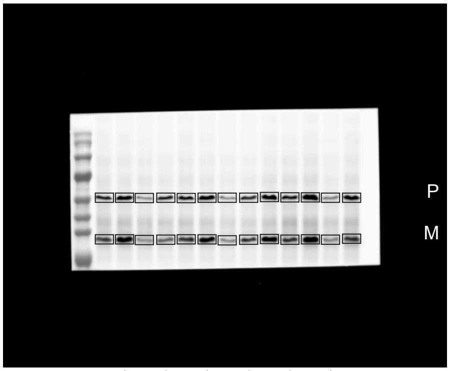


1 3 5 7 9 11 13
2 4 6 8 10 12 14

4.tif

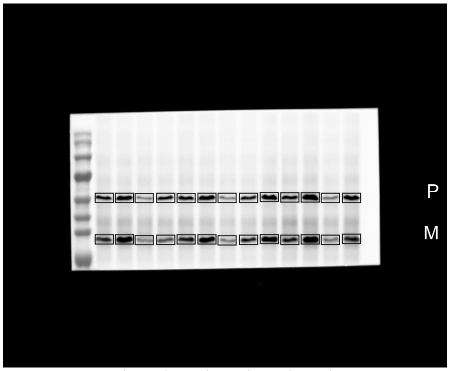


1 3 5 7 9 11 13
2 4 6 8 10 12 14

5.tif

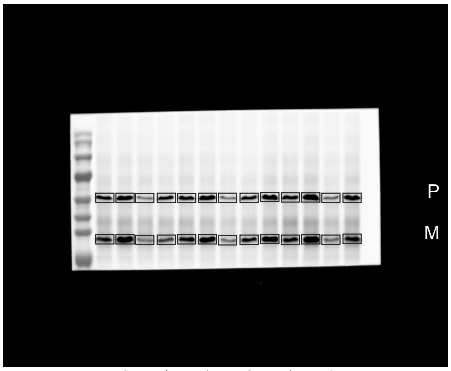


1 3 5 7 9 11 13
2 4 6 8 10 12 14

0.tif

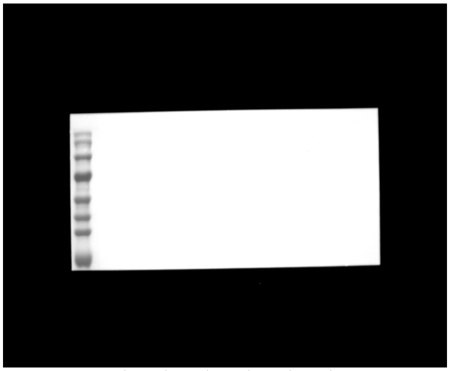

1 3 5 7 9 11 13
2 4 6 8 10 12 14

linear RGB
0 255

| Lane | Donor | Format | Treatment | Lane type |
|---|---|---|---|---|
| 1 | - | - | Marker | marker |
| 2 | D04 | 2D | Vehicle | sample |
| 3 | D04 | 2D | TGF-beta1 | sample |
| 4 | D04 | printed | Vehicle | sample |
| 5 | D04 | printed | TGF-beta1 | sample |
| 6 | D05 | 2D | Vehicle | sample |
| 7 | D05 | 2D | TGF-beta1 | sample |
| 8 | D05 | printed | Vehicle | sample |
| 9 | D05 | printed | TGF-beta1 | sample |
| 10 | D01-D06 pool | 2D | TGF-beta1, T72 h | calibration pool |
| 11 | D06 | 2D | Vehicle | sample |
| 12 | D06 | 2D | TGF-beta1 | sample |
| 13 | D06 | printed | Vehicle | sample |
| 14 | D06 | printed | TGF-beta1 | sample |
| 15 | - | - | loading buffer | empty |

3.tif *

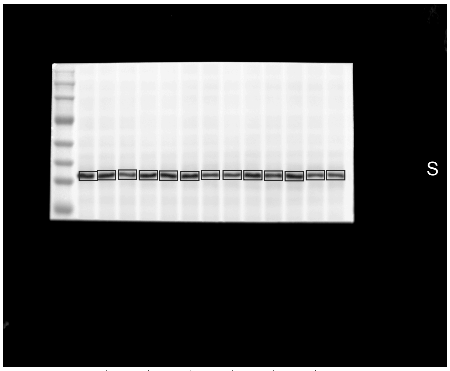


1 3 5 7 9 11 13
2 4 6 8 10 12 14

1.tif

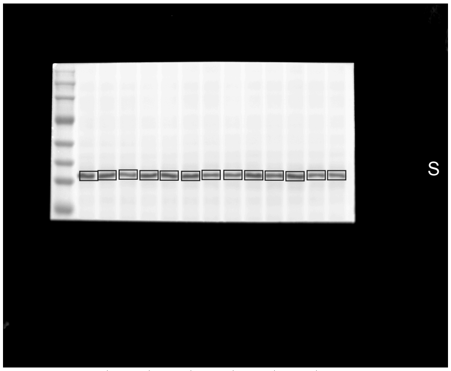


1 3 5 7 9 11 13
2 4 6 8 10 12 14

2.tif

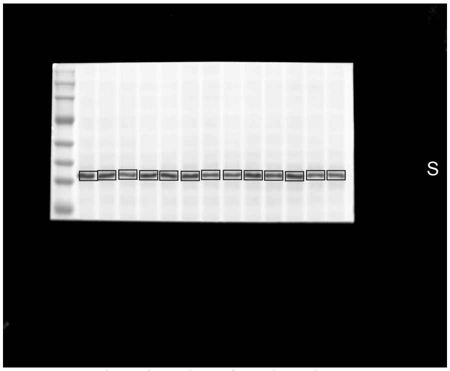


1 3 5 7 9 11 13
2 4 6 8 10 12 14

4.tif

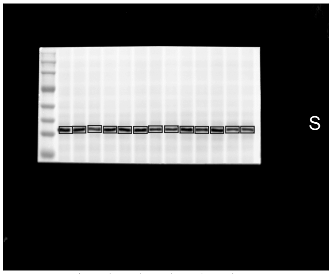


1 3 5 7 9 11 13
2 4 6 8 10 12 14

5.tif

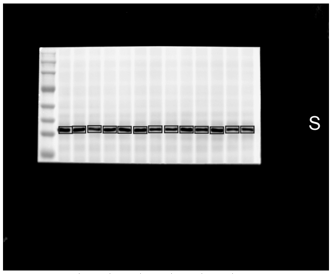


1 3 5 7 9 11 13
2 4 6 8 10 12 14

6.tif

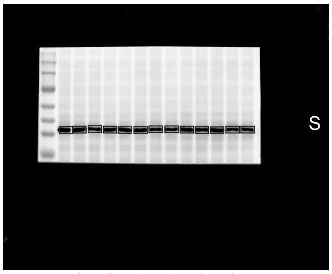


1 3 5 7 9 11 13
2 4 6 8 10 12 14

0.tif

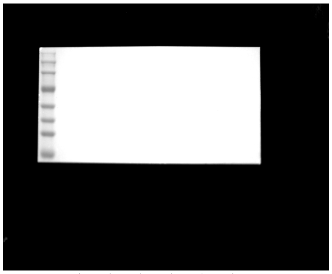

1 3 5 7 9 11 13
2 4 6 8 10 12 14

linear RGB
0 255

| Lane | Donor | Format | Treatment | Lane type |
|---|---|---|---|---|
| 1 | - | - | Marker | marker |
| 2 | D01 | 2D | Vehicle + spiked TGF-beta1 | sample |
| 3 | D01 | 2D | SB431542 + spiked TGF-beta1 | sample |
| 4 | D01 | printed | Vehicle + spiked TGF-beta1 | sample |
| 5 | D01 | printed | SB431542 + spiked TGF-beta1 | sample |
| 6 | D02 | 2D | Vehicle + spiked TGF-beta1 | sample |
| 7 | D02 | 2D | SB431542 + spiked TGF-beta1 | sample |
| 8 | D02 | printed | Vehicle + spiked TGF-beta1 | sample |
| 9 | D02 | printed | SB431542 + spiked TGF-beta1 | sample |
| 10 | D01-D06 pool | 2D | TGF-beta1, T72 h | calibration pool |
| 11 | D04 | 2D | Vehicle + spiked TGF-beta1 | sample |
| 12 | D04 | 2D | SB431542 + spiked TGF-beta1 | sample |
| 13 | D04 | printed | Vehicle + spiked TGF-beta1 | sample |
| 14 | D04 | printed | SB431542 + spiked TGF-beta1 | sample |
| 15 | - | - | loading buffer | empty |

2.tif *

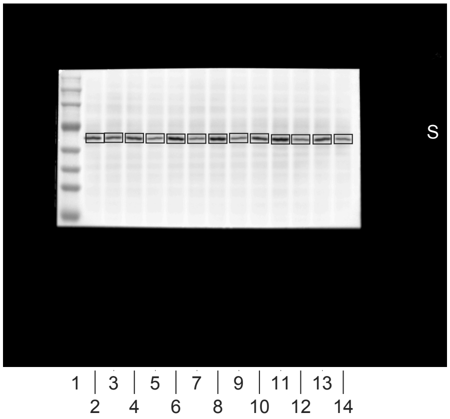


1.tif

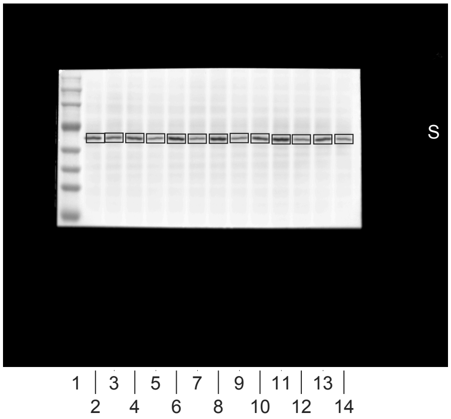


3.tif

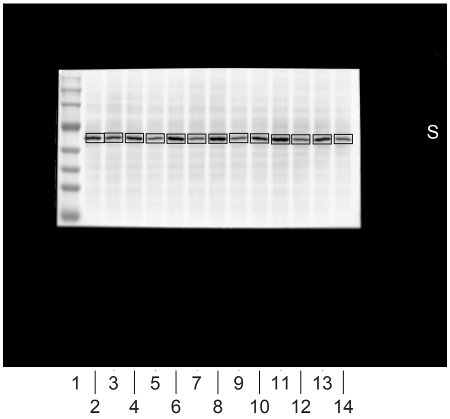


4.tif

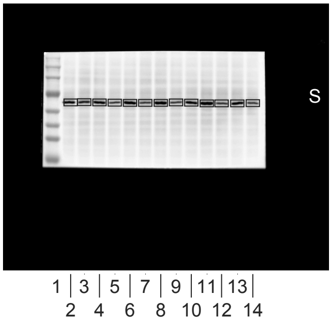


5.tif

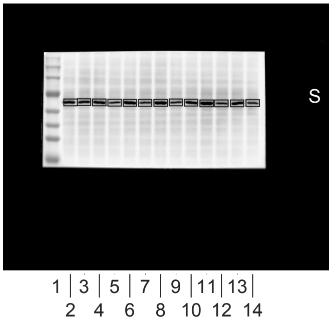


6.tif

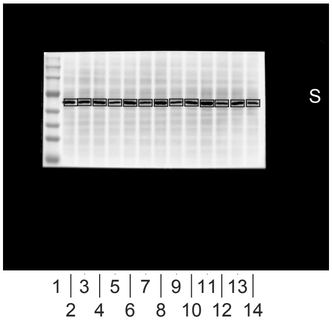


0.tif

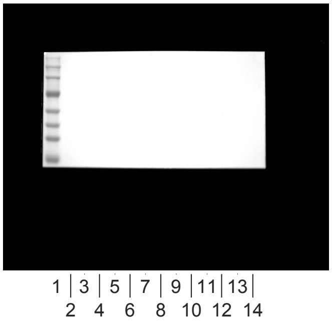


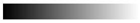
linear RGB

0 255

| Lane | Donor | Format | Treatment | Lane type |
|---|---|---|---|---|
| 1 | - | - | Marker | marker |
| 2 | D01 | 2D | Vehicle + spiked TGF-beta1 | sample |
| 3 | D01 | 2D | SB431542 + spiked TGF-beta1 | sample |
| 4 | D01 | printed | Vehicle + spiked TGF-beta1 | sample |
| 5 | D01 | printed | SB431542 + spiked TGF-beta1 | sample |
| 6 | D02 | 2D | Vehicle + spiked TGF-beta1 | sample |
| 7 | D02 | 2D | SB431542 + spiked TGF-beta1 | sample |
| 8 | D02 | printed | Vehicle + spiked TGF-beta1 | sample |
| 9 | D02 | printed | SB431542 + spiked TGF-beta1 | sample |
| 10 | D01-D06 pool | 2D | TGF-beta1, T72 h | calibration pool |
| 11 | D04 | 2D | Vehicle + spiked TGF-beta1 | sample |
| 12 | D04 | 2D | SB431542 + spiked TGF-beta1 | sample |
| 13 | D04 | printed | Vehicle + spiked TGF-beta1 | sample |
| 14 | D04 | printed | SB431542 + spiked TGF-beta1 | sample |
| 15 | - | - | loading buffer | empty |

1.tif *

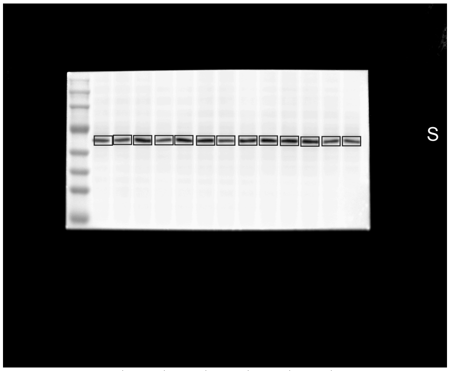


2.tif

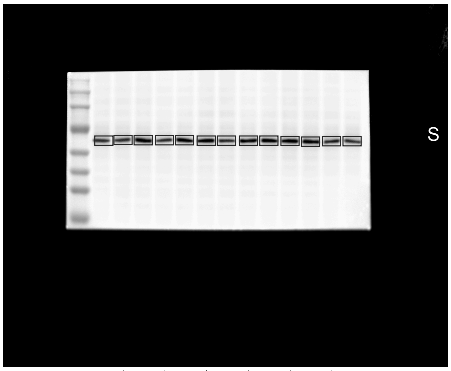


3.tif

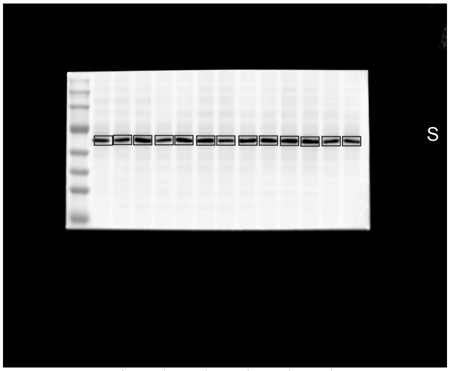


4.tif

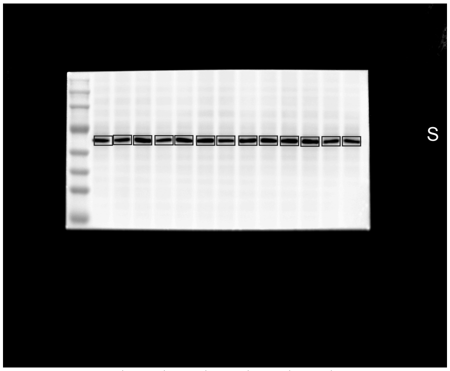


5.tif

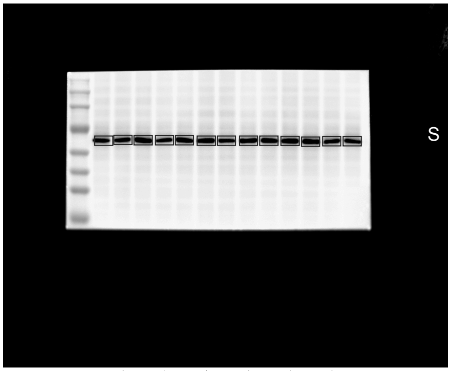


0.tif

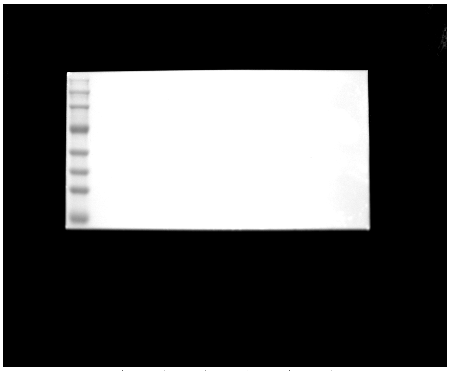


linear RGB
0 255

| Lane | Donor | Format | Treatment | Lane type |
|---|---|---|---|---|
| 1 | - | - | Marker | marker |
| 2 | D01 | 2D | Vehicle + spiked TGF-beta1 | sample |
| 3 | D01 | 2D | SB431542 + spiked TGF-beta1 | sample |
| 4 | D01 | printed | Vehicle + spiked TGF-beta1 | sample |
| 5 | D01 | printed | SB431542 + spiked TGF-beta1 | sample |
| 6 | D02 | 2D | Vehicle + spiked TGF-beta1 | sample |
| 7 | D02 | 2D | SB431542 + spiked TGF-beta1 | sample |
| 8 | D02 | printed | Vehicle + spiked TGF-beta1 | sample |
| 9 | D02 | printed | SB431542 + spiked TGF-beta1 | sample |
| 10 | D01-D06 pool | 2D | TGF-beta1, T72 h | calibration pool |
| 11 | D04 | 2D | Vehicle + spiked TGF-beta1 | sample |
| 12 | D04 | 2D | SB431542 + spiked TGF-beta1 | sample |
| 13 | D04 | printed | Vehicle + spiked TGF-beta1 | sample |
| 14 | D04 | printed | SB431542 + spiked TGF-beta1 | sample |
| 15 | - | - | loading buffer | empty |

1.tif *

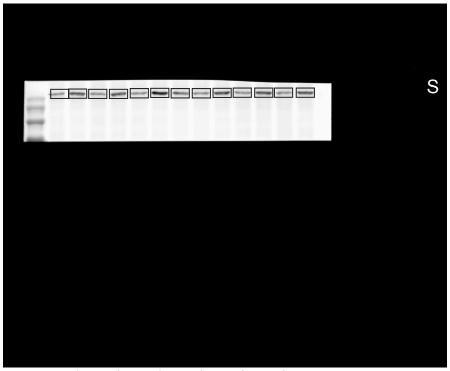


1 3 5 7 9 11 13
2 4 6 8 10 12 14

2.tif

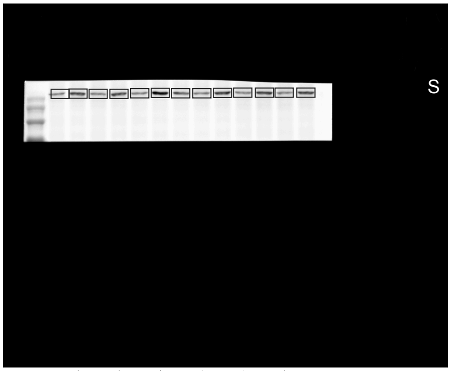


1 3 5 7 9 11 13
2 4 6 8 10 12 14

3.tif

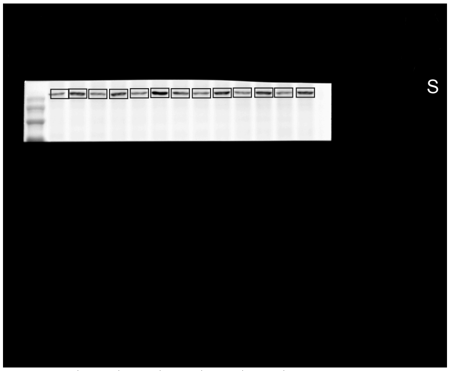


1 3 5 7 9 11 13
2 4 6 8 10 12 14

4.tif

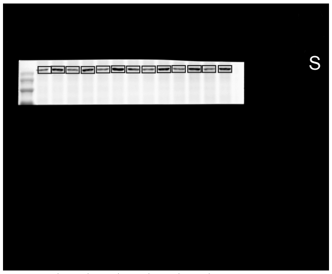


1 3 5 7 9 11 13
2 4 6 8 10 12 14

5.tif

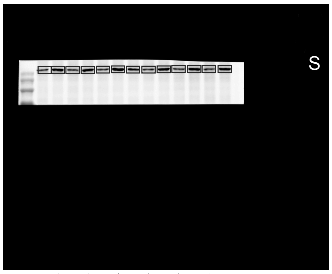


1 3 5 7 9 11 13
2 4 6 8 10 12 14

6.tif

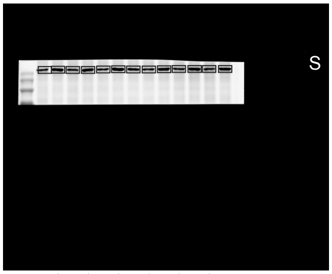


1 3 5 7 9 11 13
2 4 6 8 10 12 14

0.tif

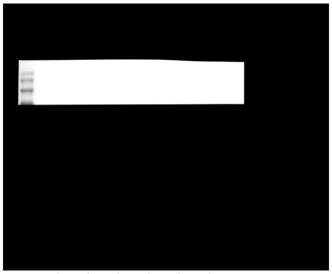

1 3 5 7 9 11 13
2 4 6 8 10 12 14

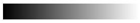

linear RGB

0 255

| Lane | Donor | Format | Treatment | Lane type |
|---|---|---|---|---|
| 1 | - | - | Marker | marker |
| 2 | D01 | printed | Vehicle history + delayed SB431542 | sample |
| 3 | D01 | printed | TGF-beta1 history + delayed SB431542 | sample |
| 4 | D02 | printed | Vehicle history + delayed SB431542 | sample |
| 5 | D02 | printed | TGF-beta1 history + delayed SB431542 | sample |
| 6 | D03 | printed | Vehicle history + delayed SB431542 | sample |
| 7 | D03 | printed | TGF-beta1 history + delayed SB431542 | sample |
| 8 | D01-D06 pool | 2D | TGF-beta1, T72 h | calibration pool |
| 9 | D04 | printed | Vehicle history + delayed SB431542 | sample |
| 10 | D04 | printed | TGF-beta1 history + delayed SB431542 | sample |
| 11 | D05 | printed | Vehicle history + delayed SB431542 | sample |
| 12 | D05 | printed | TGF-beta1 history + delayed SB431542 | sample |
| 13 | D06 | printed | Vehicle history + delayed SB431542 | sample |
| 14 | D06 | printed | TGF-beta1 history + delayed SB431542 | sample |
| 15 | - | - | loading buffer | empty |

1.tif *

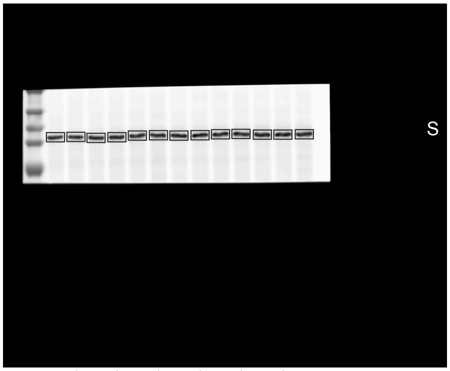


1 | 3 | 5 | 7 | 9 | 11 | 13 |
2 4 6 8 10 12 14

2.tif

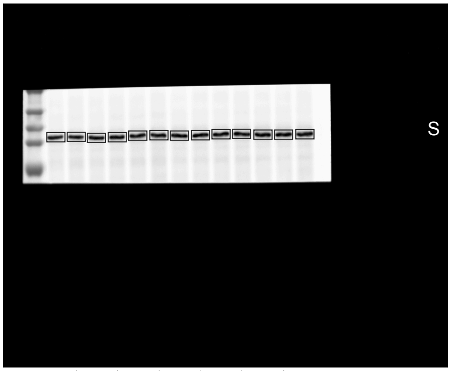


1 | 3 | 5 | 7 | 9 | 11 | 13 |
2 4 6 8 10 12 14

3.tif

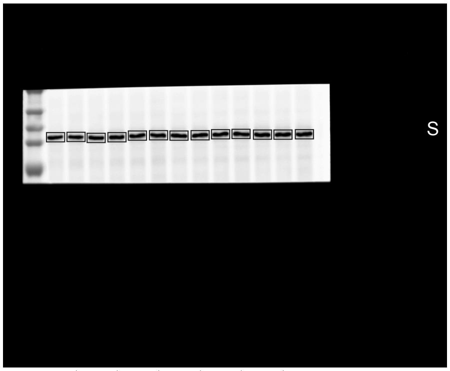


1 | 3 | 5 | 7 | 9 | 11 | 13 |
2 4 6 8 10 12 14

4.tif

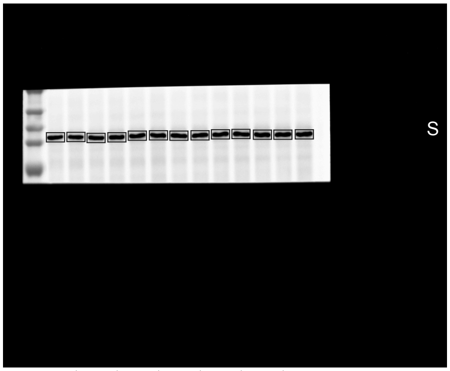


1 | 3 | 5 | 7 | 9 | 11 | 13 |
2 4 6 8 10 12 14

5.tif

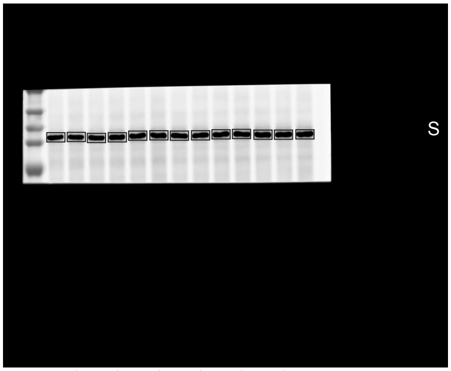


1 | 3 | 5 | 7 | 9 | 11 | 13 |
2 4 6 8 10 12 14

0.tif

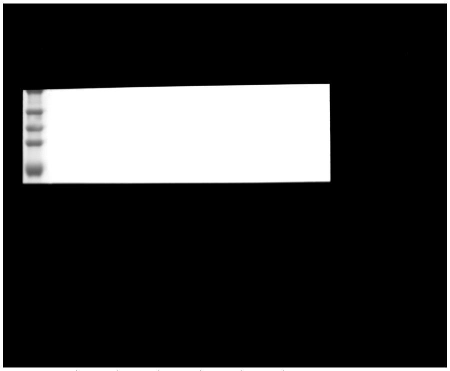

1 | 3 | 5 | 7 | 9 | 11 | 13 |
2 4 6 8 10 12 14

linear RGB
0 255

| Lane | Donor | Format | Treatment | Lane type |
|---|---|---|---|---|
| 1 | - | - | Marker | marker |
| 2 | D01 | printed | Vehicle history + delayed SB431542 | sample |
| 3 | D01 | printed | TGF-beta1 history + delayed SB431542 | sample |
| 4 | D02 | printed | Vehicle history + delayed SB431542 | sample |
| 5 | D02 | printed | TGF-beta1 history + delayed SB431542 | sample |
| 6 | D03 | printed | Vehicle history + delayed SB431542 | sample |
| 7 | D03 | printed | TGF-beta1 history + delayed SB431542 | sample |
| 8 | D01-D06 pool | 2D | TGF-beta1, T72 h | calibration pool |
| 9 | D04 | printed | Vehicle history + delayed SB431542 | sample |
| 10 | D04 | printed | TGF-beta1 history + delayed SB431542 | sample |
| 11 | D05 | printed | Vehicle history + delayed SB431542 | sample |
| 12 | D05 | printed | TGF-beta1 history + delayed SB431542 | sample |
| 13 | D06 | printed | Vehicle history + delayed SB431542 | sample |
| 14 | D06 | printed | TGF-beta1 history + delayed SB431542 | sample |
| 15 | - | - | loading buffer | empty |

2.tif *

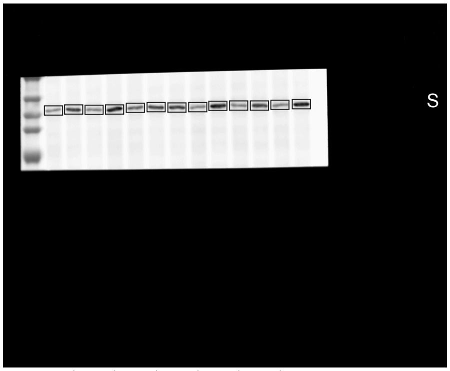


1.tif

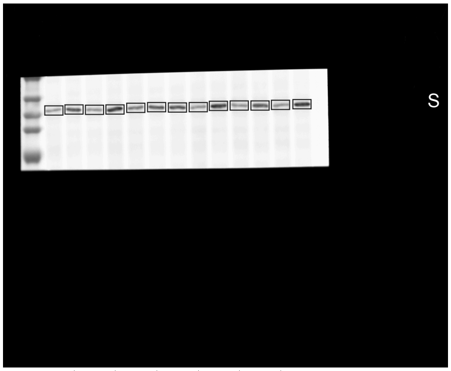


3.tif

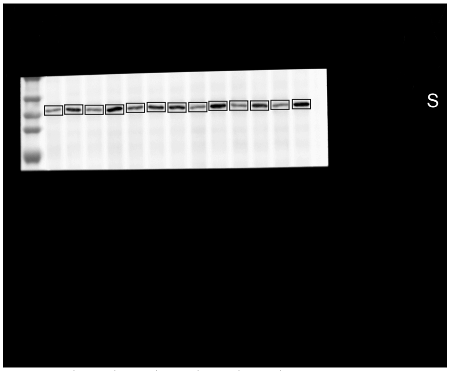


4.tif

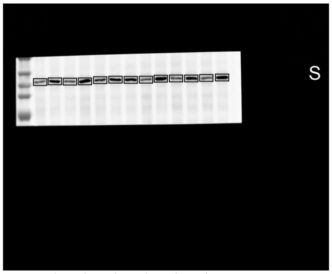


5.tif

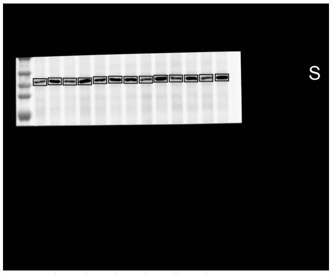


6.tif

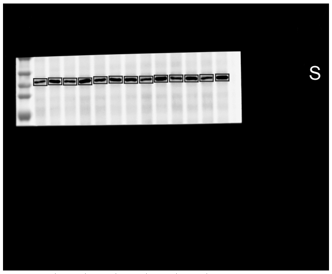


0.tif

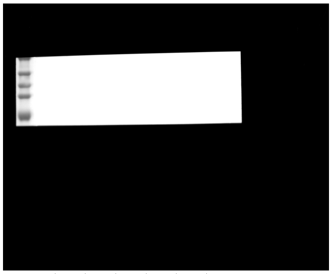


linear RGB
0 255

| Lane | Donor | Format | Treatment | Lane type |
|---|---|---|---|---|
| 1 | - | - | Marker | marker |
| 2 | D01 | printed | Vehicle history + delayed SB431542 | sample |
| 3 | D01 | printed | TGF-beta1 history + delayed SB431542 | sample |
| 4 | D02 | printed | Vehicle history + delayed SB431542 | sample |
| 5 | D02 | printed | TGF-beta1 history + delayed SB431542 | sample |
| 6 | D03 | printed | Vehicle history + delayed SB431542 | sample |
| 7 | D03 | printed | TGF-beta1 history + delayed SB431542 | sample |
| 8 | D01-D06 pool | 2D | TGF-beta1, T72 h | calibration pool |
| 9 | D04 | printed | Vehicle history + delayed SB431542 | sample |
| 10 | D04 | printed | TGF-beta1 history + delayed SB431542 | sample |
| 11 | D05 | printed | Vehicle history + delayed SB431542 | sample |
| 12 | D05 | printed | TGF-beta1 history + delayed SB431542 | sample |
| 13 | D06 | printed | Vehicle history + delayed SB431542 | sample |
| 14 | D06 | printed | TGF-beta1 history + delayed SB431542 | sample |
| 15 | - | - | loading buffer | empty |

2.tif *

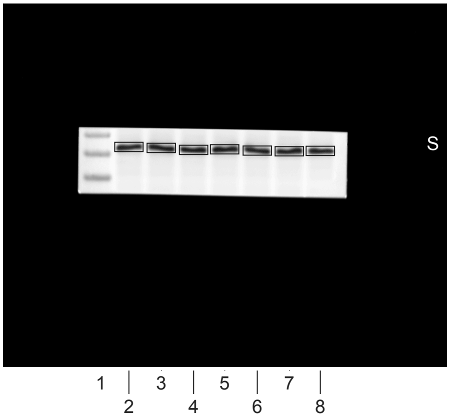


1.tif

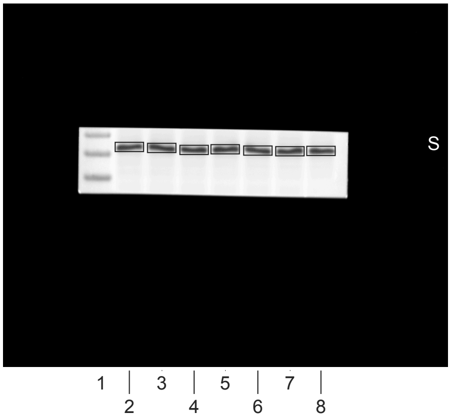


3.tif

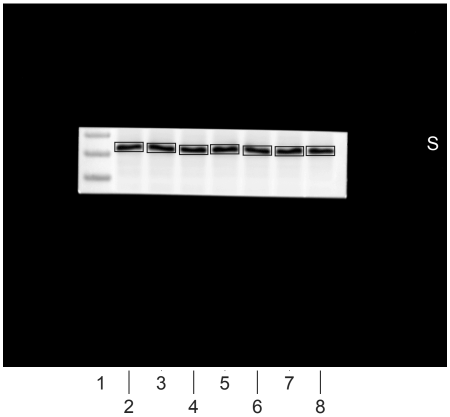


4.tif

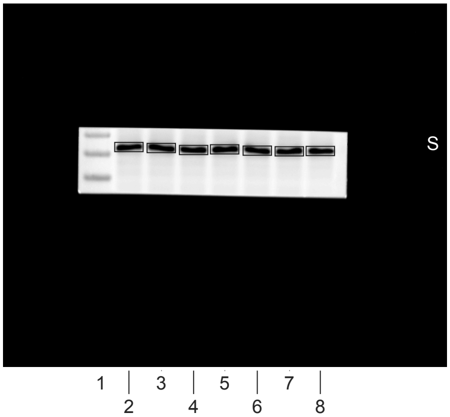


5.tif

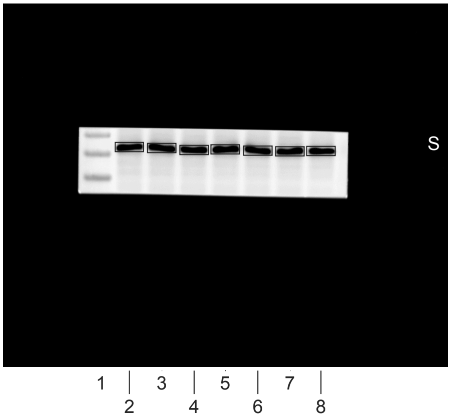


0.tif

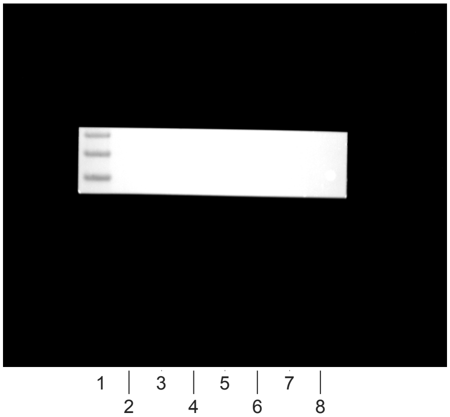


linear RGB
0 255

| Lane | Donor | Format | Treatment | Lane type |
|---|---|---|---|---|
| 1 | - | - | Marker | marker |
| 2 | D01 | printed | TGF-beta1 history, pre-dose | sample |
| 3 | D01 | printed | TGF-beta1 history, ~1 h post-dose | sample |
| 4 | D02 | printed | TGF-beta1 history, pre-dose | sample |
| 5 | D02 | printed | TGF-beta1 history, ~1 h post-dose | sample |
| 6 | D01-D06 pool | 2D | TGF-beta1, T72 h | calibration pool |
| 7 | D04 | printed | TGF-beta1 history, pre-dose | sample |
| 8 | D04 | printed | TGF-beta1 history, ~1 h post-dose | sample |
| 9 | - | - | loading buffer | empty |
| 10 | - | - | loading buffer | empty |
| 11 | - | - | loading buffer | empty |
| 12 | - | - | loading buffer | empty |
| 13 | - | - | loading buffer | empty |
| 14 | - | - | loading buffer | empty |
| 15 | - | - | loading buffer | empty |

1.tif *

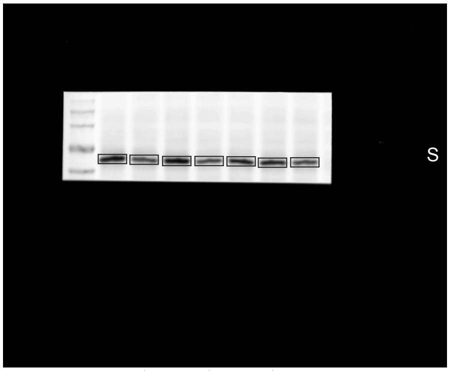


1 2 3 4 5 6 7 8

2.tif

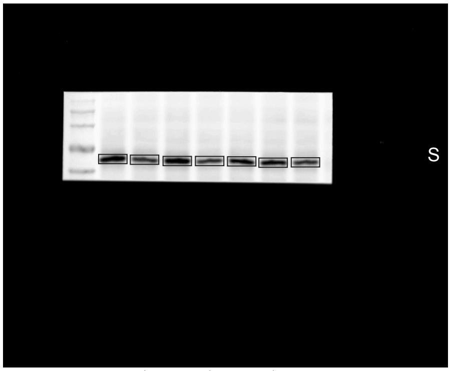


1 2 3 4 5 6 7 8

3.tif

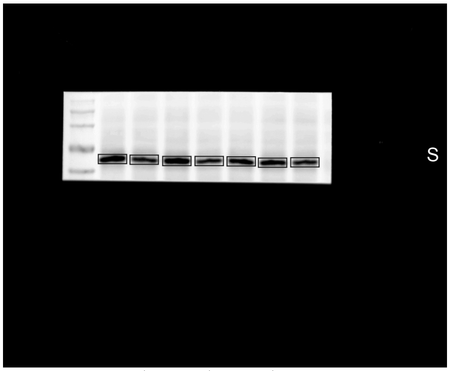


1 2 3 4 5 6 7 8

4.tif

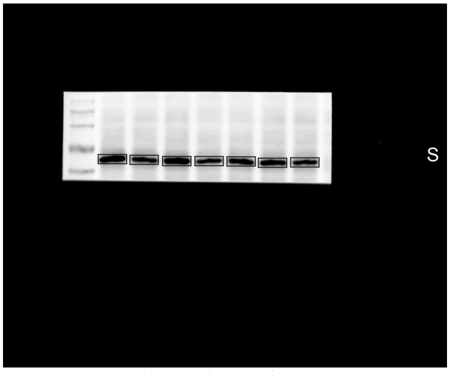


1 2 3 4 5 6 7 8

5.tif

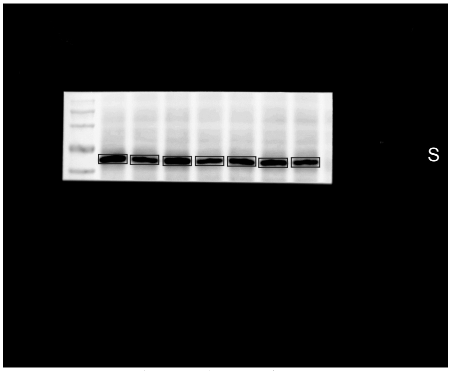


1 2 3 4 5 6 7 8

0.tif

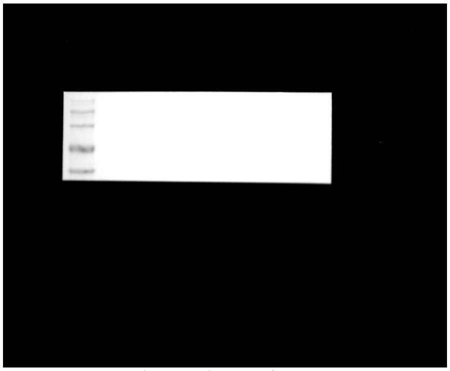

1 2 3 4 5 6 7 8

linear RGB

0 255

| Lane | Donor | Format | Treatment | Lane type |
|---|---|---|---|---|
| 1 | - | - | Marker | marker |
| 2 | D01 | printed | TGF-beta1 history, pre-dose | sample |
| 3 | D01 | printed | TGF-beta1 history, ~1 h post-dose | sample |
| 4 | D02 | printed | TGF-beta1 history, pre-dose | sample |
| 5 | D02 | printed | TGF-beta1 history, ~1 h post-dose | sample |
| 6 | D01-D06 pool | 2D | TGF-beta1, T72 h | calibration pool |
| 7 | D04 | printed | TGF-beta1 history, pre-dose | sample |
| 8 | D04 | printed | TGF-beta1 history, ~1 h post-dose | sample |
| 9 | - | - | loading buffer | empty |
| 10 | - | - | loading buffer | empty |
| 11 | - | - | loading buffer | empty |
| 12 | - | - | loading buffer | empty |
| 13 | - | - | loading buffer | empty |
| 14 | - | - | loading buffer | empty |
| 15 | - | - | loading buffer | empty |

1.tif *

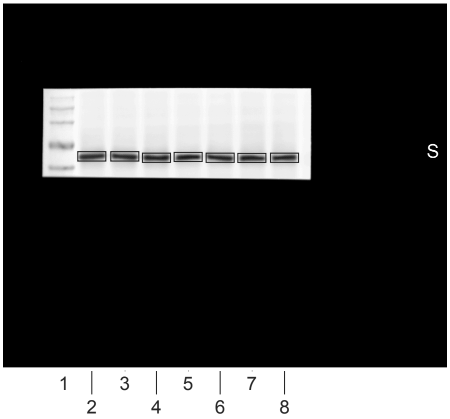


2.tif

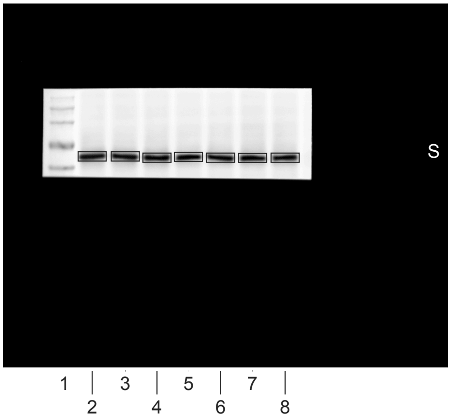


3.tif

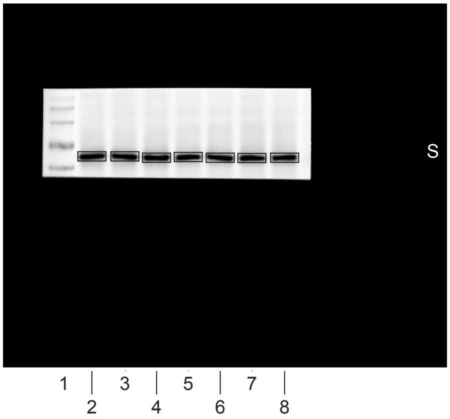


4.tif

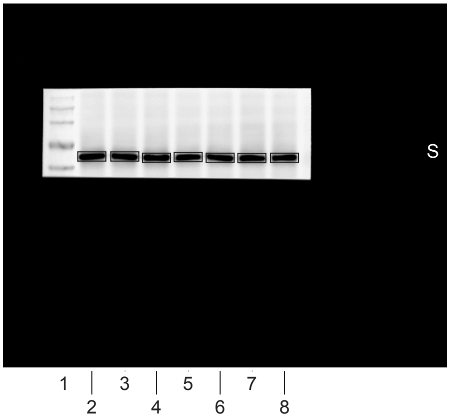


5.tif

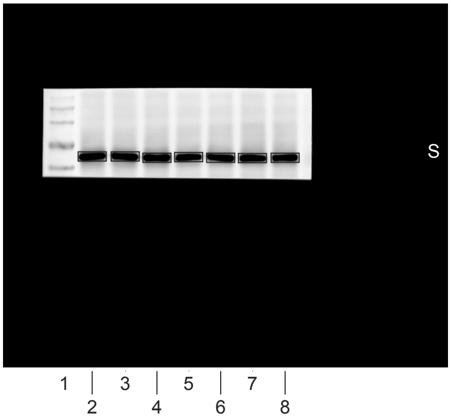


0.tif

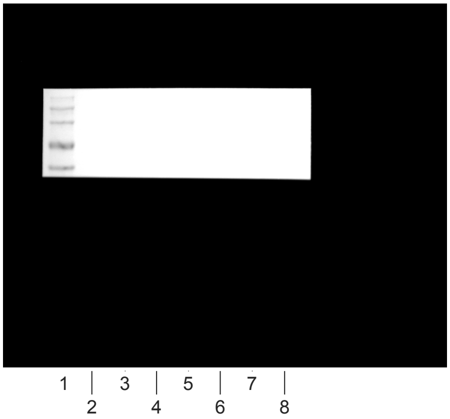


linear RGB
0 255

| Lane | Donor | Format | Treatment | Lane type |
|---|---|---|---|---|
| 1 | - | - | Marker | marker |
| 2 | D01 | printed | TGF-beta1 history, pre-dose | sample |
| 3 | D01 | printed | TGF-beta1 history, ~1 h post-dose | sample |
| 4 | D02 | printed | TGF-beta1 history, pre-dose | sample |
| 5 | D02 | printed | TGF-beta1 history, ~1 h post-dose | sample |
| 6 | D01-D06 pool | 2D | TGF-beta1, T72 h | calibration pool |
| 7 | D04 | printed | TGF-beta1 history, pre-dose | sample |
| 8 | D04 | printed | TGF-beta1 history, ~1 h post-dose | sample |
| 9 | - | - | loading buffer | empty |
| 10 | - | - | loading buffer | empty |
| 11 | - | - | loading buffer | empty |
| 12 | - | - | loading buffer | empty |
| 13 | - | - | loading buffer | empty |
| 14 | - | - | loading buffer | empty |
| 15 | - | - | loading buffer | empty |

m01_whole.jpg

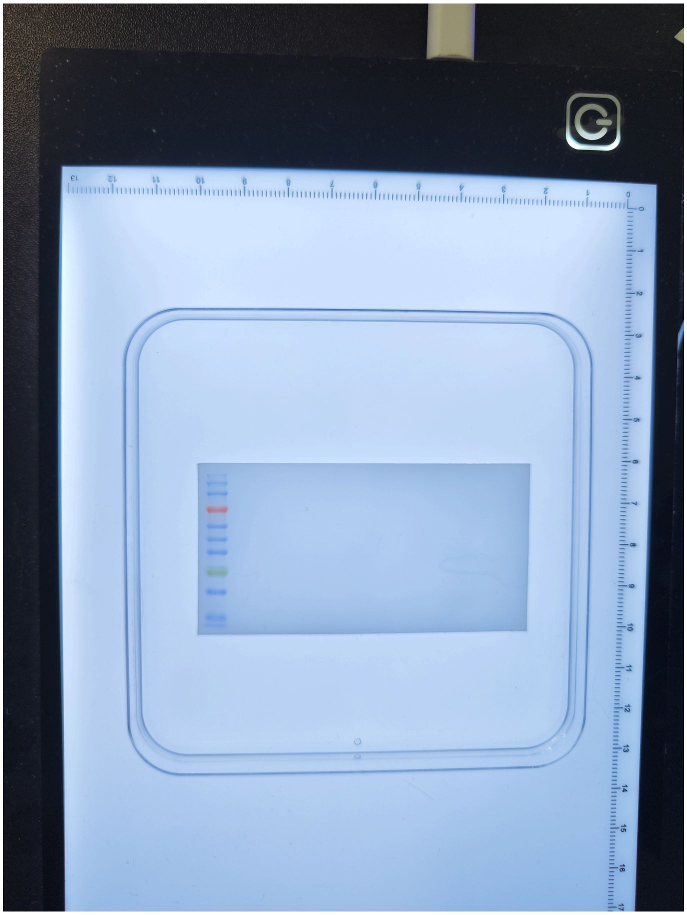

7f0a2825d18b3dddc84c

m01_cut.jpg

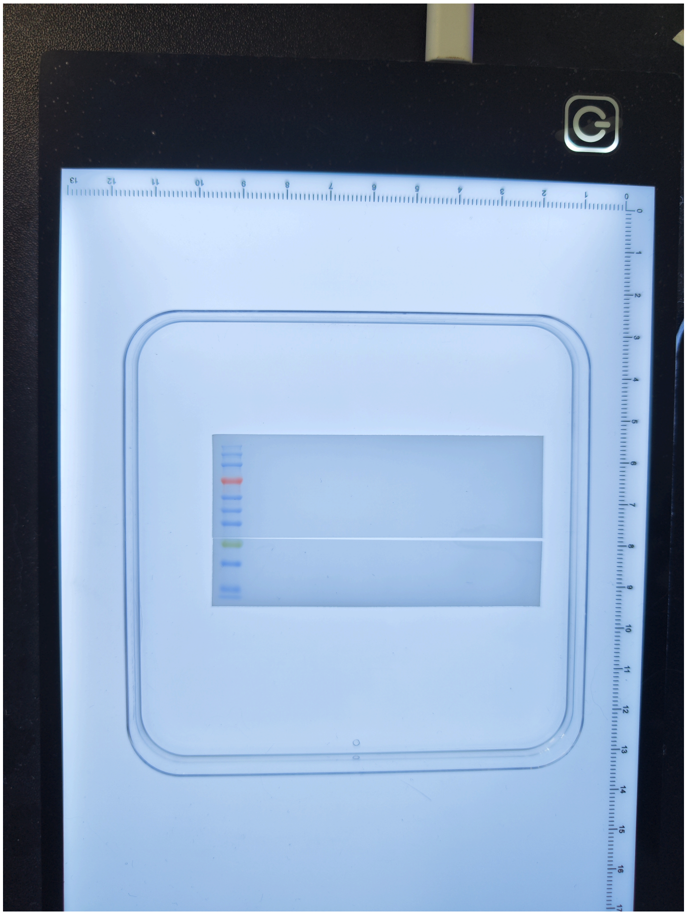

aa8630e93ed3f16f49ac

| Lane | Donor | Format | Treatment | Lane type |
|---|---|---|---|---|
| 1 | - | - | Marker | marker |
| 2 | D01 | 2D | Vehicle | sample |
| 3 | D01 | 2D | TGF-beta1 | sample |
| 4 | D01 | cast | Vehicle | sample |
| 5 | D01 | cast | TGF-beta1 | sample |
| 6 | D01 | printed | Vehicle | sample |
| 7 | D01 | printed | TGF-beta1 | sample |
| 8 | D01-D06 pool | 2D | TGF-beta1, T72 h | calibration pool |
| 9 | D04 | 2D | Vehicle | sample |
| 10 | D04 | 2D | TGF-beta1 | sample |
| 11 | D04 | cast | Vehicle | sample |
| 12 | D04 | cast | TGF-beta1 | sample |
| 13 | D04 | printed | Vehicle | sample |
| 14 | D04 | printed | TGF-beta1 | sample |
| 15 | - | - | loading buffer | empty |

m03_whole.jpg

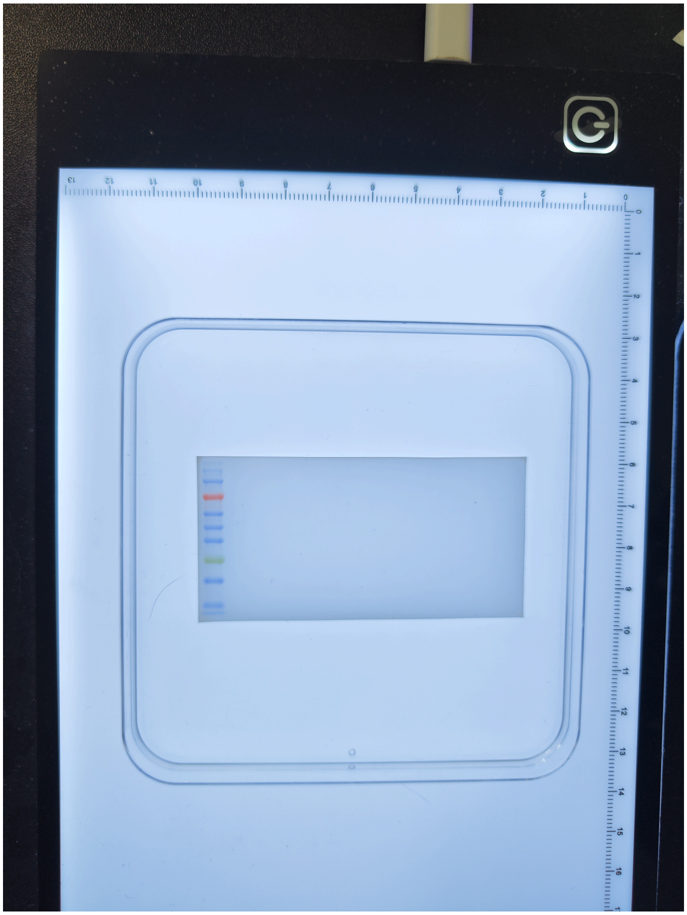

fd1a8627a3fa5b1fe1c0

m03_cut.jpg

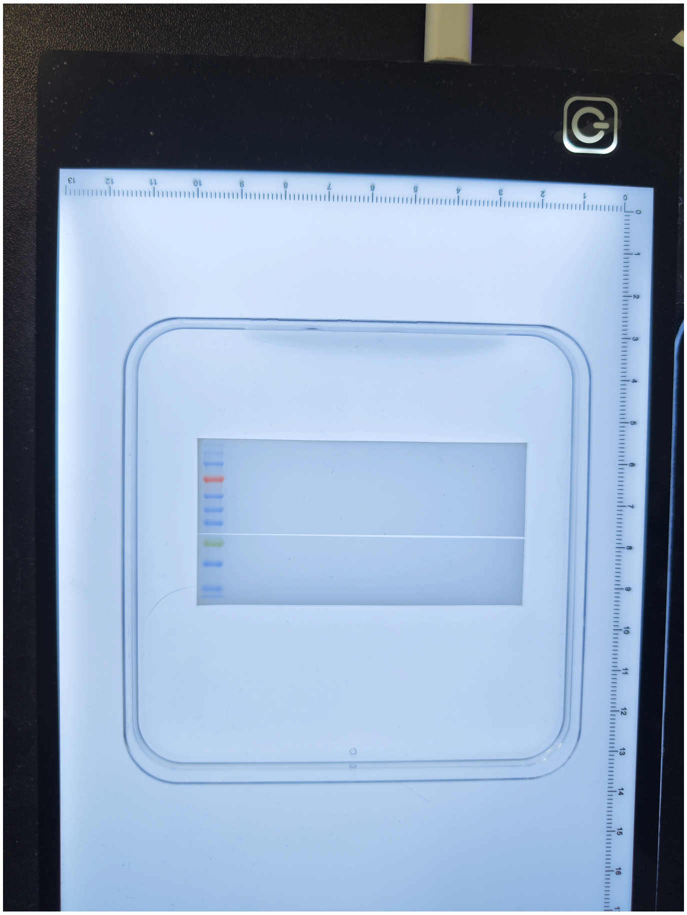

b8b15a9edffc0cdb2261

| Lane | Donor | Format | Treatment | Lane type |
|---|---|---|---|---|
| 1 | - | - | Marker | marker |
| 2 | D02 | 2D | Vehicle | sample |
| 3 | D02 | 2D | TGF-beta1 | sample |
| 4 | D02 | cast | Vehicle | sample |
| 5 | D02 | cast | TGF-beta1 | sample |
| 6 | D02 | printed | Vehicle | sample |
| 7 | D02 | printed | TGF-beta1 | sample |
| 8 | D01-D06 pool | 2D | TGF-beta1, T72 h | calibration pool |
| 9 | D05 | 2D | Vehicle | sample |
| 10 | D05 | 2D | TGF-beta1 | sample |
| 11 | D05 | cast | Vehicle | sample |
| 12 | D05 | cast | TGF-beta1 | sample |
| 13 | D05 | printed | Vehicle | sample |
| 14 | D05 | printed | TGF-beta1 | sample |
| 15 | - | - | loading buffer | empty |

m12_whole.jpg

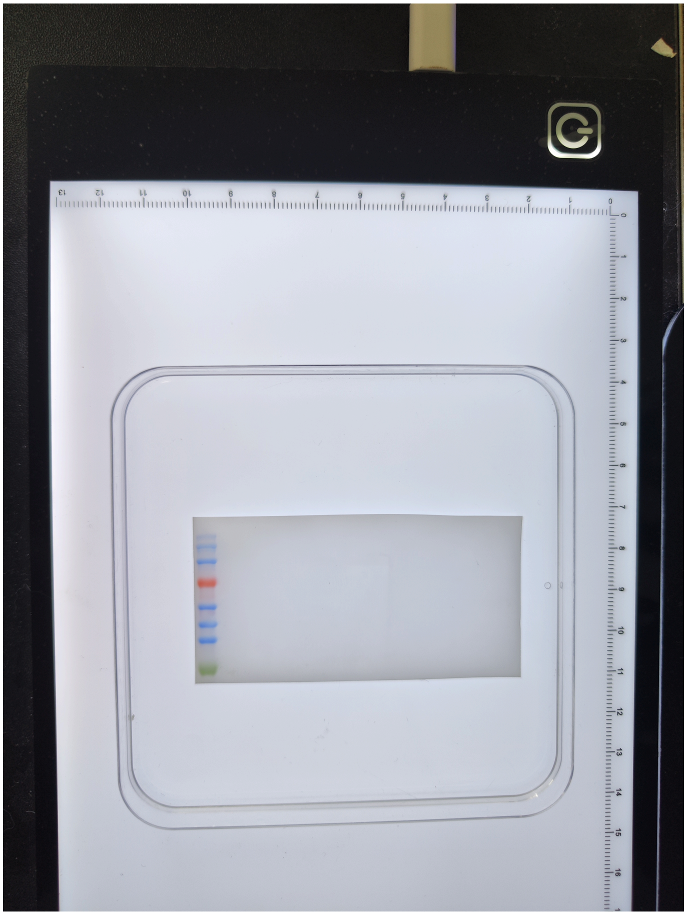

6083ec760dd6bef9f36f

m12_cut.jpg

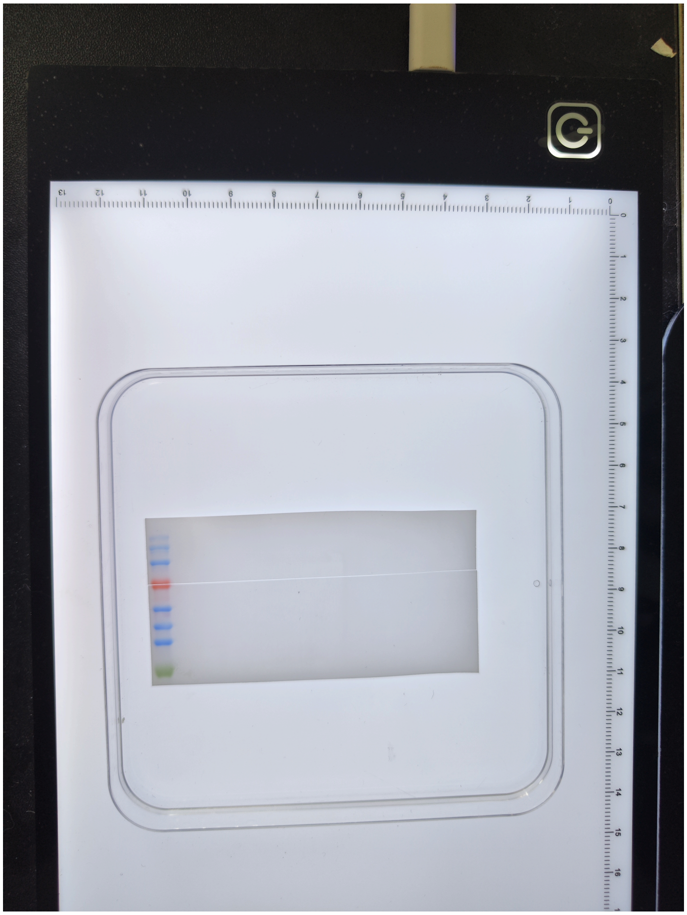

aee07c5d03f363b6057c

| Lane | Donor | Format | Treatment | Lane type |
|---|---|---|---|---|
| 1 | - | - | Marker | marker |
| 2 | D01 | printed | Vehicle history + delayed SB431542 | sample |
| 3 | D01 | printed | TGF-beta1 history + delayed SB431542 | sample |
| 4 | D02 | printed | Vehicle history + delayed SB431542 | sample |
| 5 | D02 | printed | TGF-beta1 history + delayed SB431542 | sample |
| 6 | D03 | printed | Vehicle history + delayed SB431542 | sample |
| 7 | D03 | printed | TGF-beta1 history + delayed SB431542 | sample |
| 8 | D01-D06 pool | 2D | TGF-beta1, T72 h | calibration pool |
| 9 | D04 | printed | Vehicle history + delayed SB431542 | sample |
| 10 | D04 | printed | TGF-beta1 history + delayed SB431542 | sample |
| 11 | D05 | printed | Vehicle history + delayed SB431542 | sample |
| 12 | D05 | printed | TGF-beta1 history + delayed SB431542 | sample |
| 13 | D06 | printed | Vehicle history + delayed SB431542 | sample |
| 14 | D06 | printed | TGF-beta1 history + delayed SB431542 | sample |
| 15 | - | - | loading buffer | empty |

m13_whole.jpg

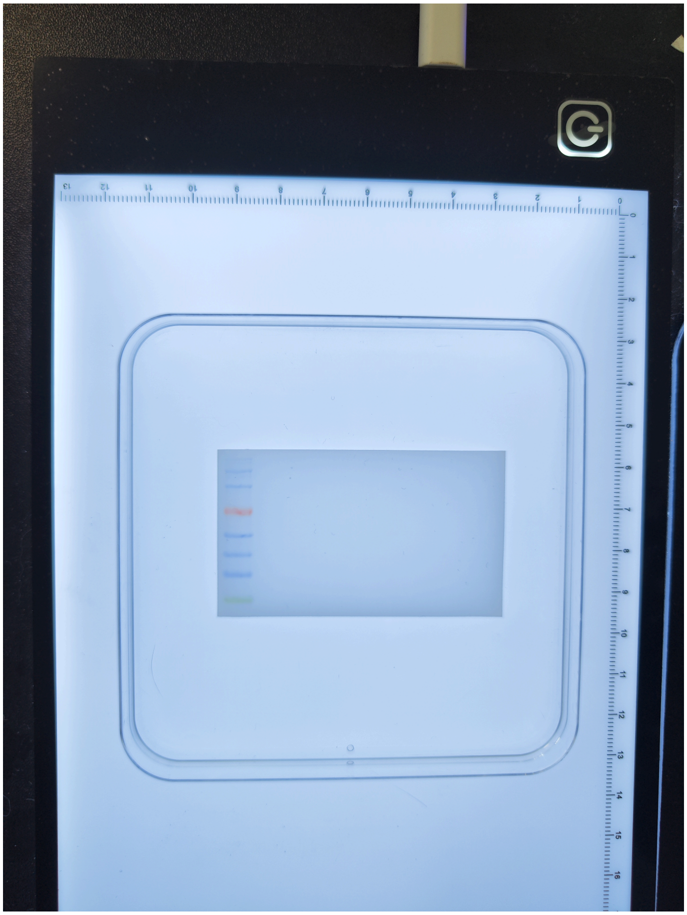

25bd9ae6b2e327322e17

m13_cut.jpg

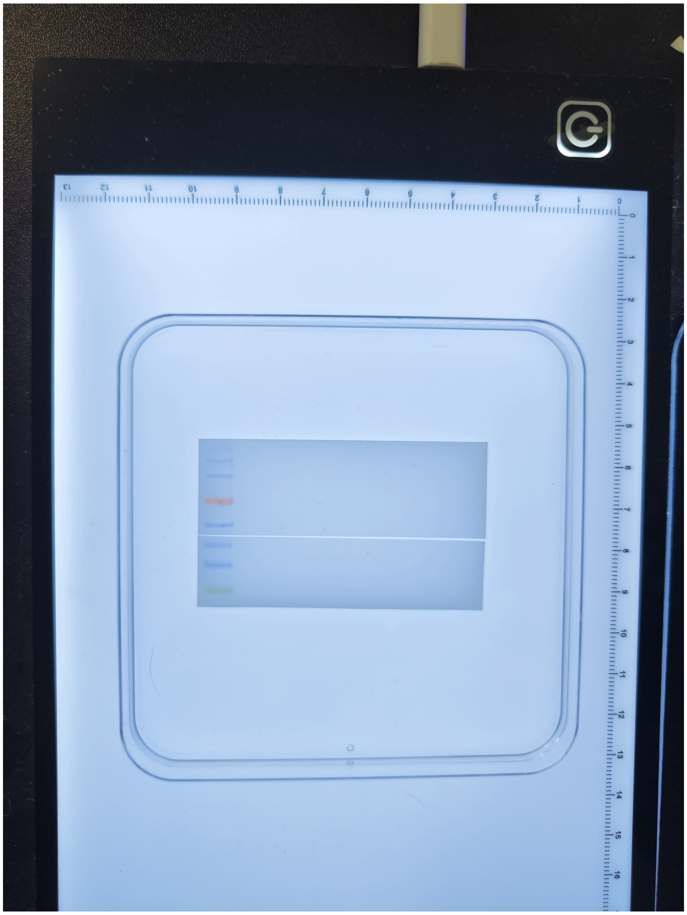

89e5e6cb060a1f3f0ce9

| Lane | Donor | Format | Treatment | Lane type |
|---|---|---|---|---|
| 1 | - | - | Marker | marker |
| 2 | D01 | printed | TGF-beta1 history, pre-dose | sample |
| 3 | D01 | printed | TGF-beta1 history, ~1 h post-dose | sample |
| 4 | D02 | printed | TGF-beta1 history, pre-dose | sample |
| 5 | D02 | printed | TGF-beta1 history, ~1 h post-dose | sample |
| 6 | D01-D06 pool | 2D | TGF-beta1, T72 h | calibration pool |
| 7 | D04 | printed | TGF-beta1 history, pre-dose | sample |
| 8 | D04 | printed | TGF-beta1 history, ~1 h post-dose | sample |
| 9 | - | - | loading buffer | empty |
| 10 | - | - | loading buffer | empty |
| 11 | - | - | loading buffer | empty |
| 12 | - | - | loading buffer | empty |
| 13 | - | - | loading buffer | empty |
| 14 | - | - | loading buffer | empty |
| 15 | - | - | loading buffer | empty |
